# Computational Microbiome Pharmacology Analysis Elucidates the Anti-Cancer Potential of Vaginal Microbes and Metabolites

**DOI:** 10.1101/2024.10.10.616351

**Authors:** Damilola C. Lawore, Smrutiti Jena, Alicia R Berard, Kenzie Birse, Alana Lamont, Romel D. Mackelprang, Laura Noel-Romas, Michelle Perner, Xuanlin Hou, Elizabeth Irungu, Nelly Mugo, Samantha Knodel, Timothy R. Muwonge, Elly Katabira, Sean M Hughes, Claire Levy, Fernanda L. Calienes, Florian Hladik, Jairam R. Lingappa, Adam D Burgener, Leopold N Green, Douglas K. Brubaker

## Abstract

The vaginal microbiome’s role in risk, progression, and treatment of female cancers has been widely explored. Yet, there remains a need to develop methods to understand the interaction of microbiome factors with host cells and to characterize their potential therapeutic functions. To address this challenge, we developed a systems biology framework we term the Pharmacobiome for microbiome pharmacology analysis. The Pharmacobiome framework evaluates similarities between microbes and microbial byproducts and known drugs based on their impact on host transcriptomic cellular signatures. Here, we apply our framework to characterization of the Anti-Gynecologic Cancer Vaginal Pharmacobiome. Using published vaginal microbiome multi-omics data from the Partners PrEP clinical trial, we constructed vaginal epithelial gene signatures associated with each profiled vaginal microbe and metabolite. We compared these microbiome-associated host gene signatures to post-drug perturbation host gene signatures associated with 35 FDA-approved anti-cancer drugs from the Library of Integrated Network-based Cellular Signatures database to identify vaginal microbes and metabolites with high statistical and functional similarity to these drugs. We found that *Lactobacilli* and their metabolites can regulate host gene expression in ways similar to many anti-cancer drugs. Additionally, we experimentally tested our model prediction that taurine, a metabolite produced by *L. crispatus,* kills cancerous breast and endometrial cancer cells. Our study shows that the Pharmacobiome is a powerful framework for characterizing the anti-cancer therapeutic potential of vaginal microbiome factors with generalizability to other cancers, microbiomes, and diseases.

## Introduction

The vaginal microbiota consists of a diverse range of beneficial microbes and potential pathogens that reside within the vaginal environment^1, 2^. Techniques like metagenomics, metatranscriptomics, metaproteomics, and metabolomics have laid the foundation for understanding the complexity of the vaginal microbiome and highlighted the importance of specific bacteria species like *Lactobacillus* species in maintaining a healthy vaginal environment^3, 4^. The vaginal microbiota varies during a woman’s life and the menstrual cycle, making dysbiosis of the vaginal microbiota challenging to diagnose. Nonetheless, vaginal microbiota lacking *Lactobacillus* species have been linked to a variety of disease states, including persistent HPV infection, an increased risk of sexually transmitted infection (STI), pelvic inflammatory disease, preterm birth, poor obstetric outcomes, infertility, and the development of gynecologic cancers such as cervical cancer, endometrial cancer, and ovarian cancer^5, 6, 7, 8^.

The vaginal microbiome has been the subject of increasing interest in the context of gynecological cancers, where studies have shown that dysbiosis may induce epithelial barrier dysfunction, immunological dysregulation, genotoxicity, and inflammation, resulting in a tumor-permissive microenvironment^9, 10^. Bacteria, like *Chlamydia trachomatis*, can trigger epithelial-to-mesenchymal transition (EMT) in infected cells^11^. This process can potentially lead to the loss of adhesion among epithelial cells and the downregulation of DNA damage responses. These effects are significant because they are believed to contribute to the development of carcinogenic conditions^11^. Also, several studies have reported the depletion of *Lactobacillus* species and a substantial increase in vaginal microbiota diversity in women with cervical cancer, endometrial cancer, and ovarian cancer^12^. Cervical cancer is strongly associated with infection with high risk types of human papillomavirus^13^, particularly in the presence of *Sneathia sanguinegens, Anaerococcus tetradius,* and *Peptostreptococcus anaerobius*^14^. An increase in *A. vaginae* and *Porphyromonas* species is associated with endometrial cancer^15^ while *Brucella, Mycoplasma*, and *Chlamydia* species, were found in 60%–76% of ovarian cancers^16, 17, 18^.

Furthermore, bacterial communities can potentially play a role in the development, severity, and treatment response of gynecological malignancies. Due to sparse information on modulatory effects, the intricate relationship between the vaginal microbiota and gynecological cancers is not fully elucidated^19^. One key challenge is the heterogeneity observed in vaginal microbiomes across individuals. The diversity and abundance of microbial species can vary significantly among women due to age, ethnicity, hormonal status, sexual practices, antibiotic use, and hygiene habits^20^. This inherent variability makes it challenging to establish a standard reference or define a healthy vaginal microbiome profile applicable to all women. While evidence suggests that alterations in the vaginal microbiota may contribute to an increased risk of certain cancers (e.g., cervical or endometrial cancer), the exact mechanisms behind this interplay remain unclear^15^.

The vaginal microbiome also plays a significant role in influencing the response to cancer therapy by regulating inflammation in the context of gynecologic cancers^21, 12, 22^. Additionally, intraperitoneal bacterial infections have been linked to adverse events in ovarian cancer patients^23^. It is essential to recognize that the disruption of the vaginal microbiome can have indirect consequences on the efficacy of pelvic cancer treatments and the healing process following surgery^24, 22, 25^. Moreover, the microbiome has the potential to modulate immune competency in distant sites, thus impacting immune cell functionality and the release of cytokines^26, 25^. To fully comprehend these effects, it is necessary to investigate the intricate interactions between the microbiome, the immune system, and host tissues within relevant cancer models.

In addition to understanding the role of the endogenous microbiome in gynecologic cancer and its response to chemotherapy, it may be possible to apply exogenous microbes as treatment adjuvants. Precision therapy could leverage specific microbes or microbial functions for a targeted and personalized treatments This method enhances efficacy by considering individual microbial variability, which affects drug response. However, precise timing and dosing are essential to avoid disrupting microbial balance, which could cause infections, resistance, or immune reactions. Understanding the complex microbial ecosystem is key to preventing adverse effects and optimizing outcomes^27^. Prospects for microbial-based cancer therapies could be developed using known metabolites from specific microbes. Understanding how microbial products influence immune function and cell signaling could lead to targeted cancer treatments. This framework could yield precise and safe therapeutic agents worth of further exploration.

Here, we propose a systems biology computational framework to characterize the therapeutic potential of microbes and their products, termed the *Pharmacobiome*. The Pharmacobiome is the characterization of potential effects of specific microbiome community and constituents in terms of known drugs and pathways by studying associations between the microbiome and disease states, pathways, and known drug functions (**Figure 1**). One application of this concept could be for the precision treatment of cancers using microbial products identified as clinically relevant by characterizing the *Anti-Gynecologic Cancer Pharmacobiome.* Here, we examine this concept by integrating vaginal microbiome multi-omics data from patients with and without bacterial vaginosis and with *in vitro* drug transcriptomics data to identify vaginal microbes and microbial products that may have potential therapeutic application for female reproductive cancers.

**Figure 1.**
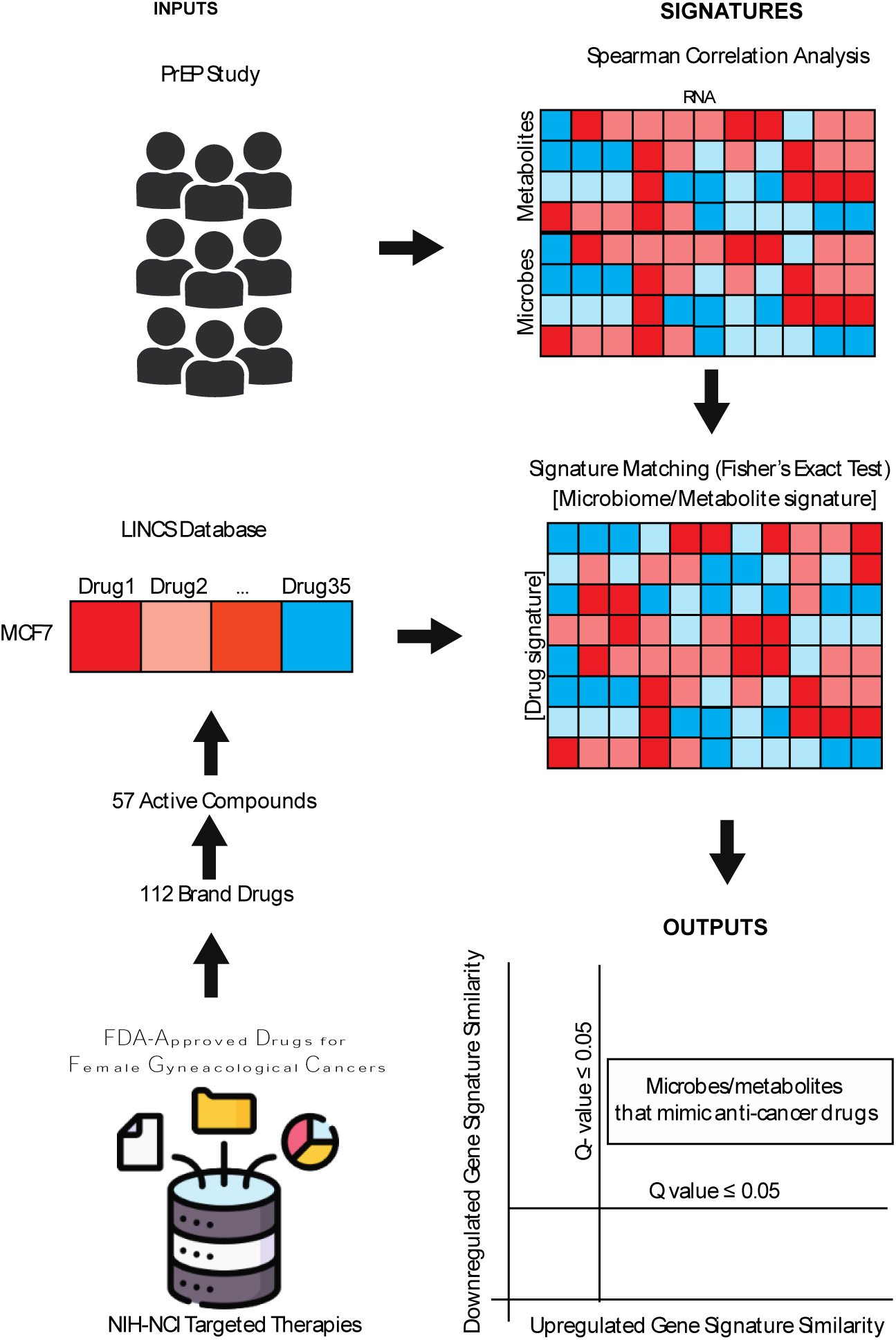
Pharmacobiome Workflow: Transcriptomics, Metabolomics and Microbiome composition data from the Partners PrEP Vaginal Microbiome Multiomics datasets were analyzed via Spearman correlation analysis to extract microbe-host gene and metabolite-host gene signatures. National Institutes of Health-National Cancer Institute drugs screening in the LINCS Small Chemical Perturbation datasets were extracted to identify drug-host gene signatures. Microbiome-gene and drug-gene up- and down-regulated signatures were compared via Fisher’s exact test to identify Anti-cancer Drug:Vaginal Microbe and Anti-cancer Drug:Vaginal Metabolite signature similarities.

## Materials and Methods

### Human vaginal microbiome multi-omics data

We obtained vaginal microbiome multi-omics data from a subgroup of 405 HIV-negative women enrolled in the Partners PrEP (Pre-Exposure Prophylaxis) study^28, 29^. The proteomics, metaproteomics, and metabolomics data were previously generated (as described)^28^ from two sets of samples collected from participants with diverse exposures to HIV-risk factors (*e.g.*, unprotected heterosexual sex, bacterial vaginosis, and other sexually transmitted infections): Group 1 included data from vaginal swabs collected from 315 participants aged 18-51 years; and Group 2 included transcriptomics data from vaginal biopsies and proteomics, metaproteomics, and metabolomics data from vaginal swabs collected from 90 consenting participants (aged 25-51). Data from Group 2 served to validate the functional pathways identified in Group 1 in terms of treatment, behavioral, clinical, or demographic factors between non-Lactobacillus dominant and Lactobacillus dominant women.

### Microbiome-Associated Host Signature Extraction

We used transcriptomics, microbiome composition, and metabolomics data from Group 2 participants. We calculated Spearman correlation coefficients between the microbiome composition or vaginal metabolomics data and host transcriptomics profiles using R studio (Version 2022.12.0+353) to generate microbe-RNA and metabolite-to-RNA correlation coefficients. For the microbiota composition analysis, we used Spearman correlation analysis, but for the metabolome analysis, we ued partial Spearman correlation analysis which allowed us to incorporate a covariate for microbiome composition (*Lactobacillus* dominant or not) in assessing metabolome-host gene correlation coefficients.

Coded identifiers were used to match between datasets to ensure data integrity. Spearman correlation analysis was carried out using the cor. test() function and the correction for multiple comparisons using the p.adjust() function, both from the R stats package(version 4.3.0). Initial gene lists for each correlation were generated for adjusted p-values less than or equal to 0.05. *Lactobacillus, Lactobacillus iners, Lactobacillus crispatus,* and *Gardnerella* were present at a relative abundance equal to or greater than 70% in the human samples.

### Identification of Post-Treatment Drug Transcriptomic Perturbation Data

The gene expression changes induced by 35 FDA-approved anti-cancer drugs were extracted from the Library of Integrated Network and Cellular Signatures (LINCS) Chemical Perturbation database^30, 31^. Using the the National Institutes of Health-National Cancer Institute database, we compiled a candidate list of 57 active compounds used as targeted therapies to treat various female gynecological cancers, including breast cancer, ovarian cancer, cervical cancer, vaginal cancer, vulvar cancer, and uterine cancer^30^. We compared the active compounds with the LINCS chemical perturbation transcriptomic database, identifying 35 critical (significant) drugs for further analysis^31^.

The LINCS Center for Transcriptomics catalogs gene expression profiles for cellular perturbagen molecules at different time points, doses, and cell lines using the L1000 assay. We obtained the LINCS data for the MCF7 breast cancer cell line because it serves as a prototypical hormone-responsive female cancer type for hypothesis generation and because it was the female cancer cell line with the largest number of drugs screened at a 10 uM dose over 24 hours, the most consistent criteria among the drugs analyzed.

### Comparison of Drug Gene Signatures with Microbiome Host Signatures

The microbes and metabolites signatures we obtained from Spearman correlations were compared to LINCS-derived signatures for similarities using a Fisher’s Exact test and corrected for multiple hypothesis testing using the Benjamini Hochberg method on R software. Each microbe and metabolite gene list was compared to the gene list of each drug, where upregulated and downregulated genes were compared separately. Fisher’s Exact Test was carried out using the fisher.test() function and the p-values correction using the p.adjust() function, both from the R stats package(version 4.3.0).

### Bacterial culture and sample preparation for metabolomics

*L. crispatus* ATCC 33820, *G. vaginalis* ATCC 14018, and *L. iners* ATCC 55195 were cultured for the metabolomics study. For *L. crispatus*, De Man–Rogosa–Sharpe media was used for suspension cultures^32^. NYC III media was used for growing *G. vaginalis* suspension^33^. *L. iners* suspension culture was grown in BHI^34^. All cultures were grown at 37^°^C, with 5% CO_2_ conditions for 24 hours. Culture supernatants were collected after centrifuging at 5000g for 10mins followed by syringe filtration with a 0.22μ filter to exclude all the bacterial cells. Samples were sent in triplicates to Metabolon, NC, USA, for non-targeted metabolomics^35^. Metabolomics profiling was performed using the method described previously^36^ and the data was stored in Metabolomics Workbench: http://dx.doi.org/10.21228/M82M8R

### Cell viability and Half-Maximal Inhibitory Concentration (IC_50_)

The Cell Counting Kit-8 (CCK8) assay was carried out to determine cell viability and the IC_50_ of Taurine. Literature has shown Taurine to reduce cell viability at concentrations of 130mM and above; hence, lower concentrations were tested in this study. Human endometrial cancer cells (HEC1A) and human breast cancer cells (MCF7) were grown to 80% confluency overnight at 37°C and 5% CO_2_ on separate 96-well tissue culture plates in McCoy’s 5A Medium and Eagle’s Minimum Essential Medium (EMEM) respectively. Then, one μL of Taurine at different concentrations (3 mM, 10 mM, 30 mM, 100 mM,130mM), 20 mM Cisplatin (positive control), and DMSO (vehicle control) were added in triplicates and incubated for 24 h at 37° C and 5% CO_2_. Next, 10uL CCK8 solution was added to each well, and the plate was rocked gently to ensure even mixing. The cells were incubated at 37° C and 5% CO2 for four hours. The absorbance was determined using a BioTek Gen5 microplate reader and imager (Agilent Technologies, Santa Clara, CA) at 450nm. A one-way analysis of variance (ANOVA) was performed to determine significance using Prism GraphPad Software (vs. 8.0, San Diego, CA, USA). p-values of ≤ 0.05 were considered significant.

### Data and Code Availability

All data has been supplied in Supplementary tables, including Spearman coefficient calculation results, Fisher’s exact calculation, gene lists, and gene counts. Published data from Partners PrEP Study is available at GEO with accession number GSE139655 for Transcriptomics data and MetaboLights with accession number MTBLS7087 for Metabolomics data. LINCS Chemical Perturbation data is available at https://maayanlab.cloud/sigcom-lincs/#/Download. The Bacterial Culture metabolomics data is available at http://dx.doi.org/10.21228/M82M8R. All original code has been deposited at GitHub: https://github.com/Brubaker-Lab/Characterizing-the-Anti-Cancer-Potential-of-Vaginal-Microbes-and-Metabolites.

## Results

### Linking vaginal epithelial and microbiome gene signatures to anti-gynecologic cancer drug gene signatures

In this study, we characterized the relationship between microbiome composition and RNA expression and the association between microbial metabolites and host vaginal epithelial RNA expression. Our analysis was conducted on microbiome composition, metabolomics, and vaginal biopsy transcriptomics data obtained from HIV-negative participants. For vaginal microbiota composistion and taxa, we performed Spearman correlation analysis between 51 measured microbes and 23,482 gene expression signatures from vaginal biopsies. The size of the coefficient determined the effect magnitude, while the sign indicated the proposed regulatory direction on gene expression (Table S1-S2). Gene lists for each microbe were generated (adjusted p < 0.05) and Fisher’s exact test was used to compare the microbe gene signatures to post-treatment gene expression perturbation signatures of 35 chemotherapeutic drugs (Table S3-S4).

We compared the microbe-gene and drug-gene signatures along two axes: Upregulated signature similarity being genes positively correlated with microbe abundance and up-regulated by drug treatment, and down-regulated signature similarity being genes negatively correlated with microbe abundance and down-regulated by drug. Doxorubicin, Fulvestrant, Etoposide, Raloxifene, and Exemestane had the highest significance for upregulated signature similarity, and Everolimus, Raloxifene, Fulvestrant, Lapatinib, and Rucaparib had the highest significance for downregulated signature similarity. **Figure 2A** shows the gene signatures of microbes that match three common anti-cancer drugs: Everolimus, Doxorubicin, and Raloxifene. Everolimus showed the strongest similarity with the microbes for downregulation; its mechanism of action is blocking growth-driven transduction signals from T and B cells^37^. Doxorubicin showed the strongest similarity with the microbes for upregulation; it interferes with the function of DNA by inhibiting the progression of topoisomerase II, which is responsible for the uncoiling of DNA for transcription^37^. Raloxifene showed high similarity in upregulation and downregulation, and it binds to estrogen receptors, activating estrogenic pathways and blockade in tissues that express estrogen receptors^37^. Across these three drugs, the *Lactobacillus* species, *Prevotella*, *Bifidobacterium*, and *Gardnerella* species are captured to be similar at different significance.

**Figure 2:**
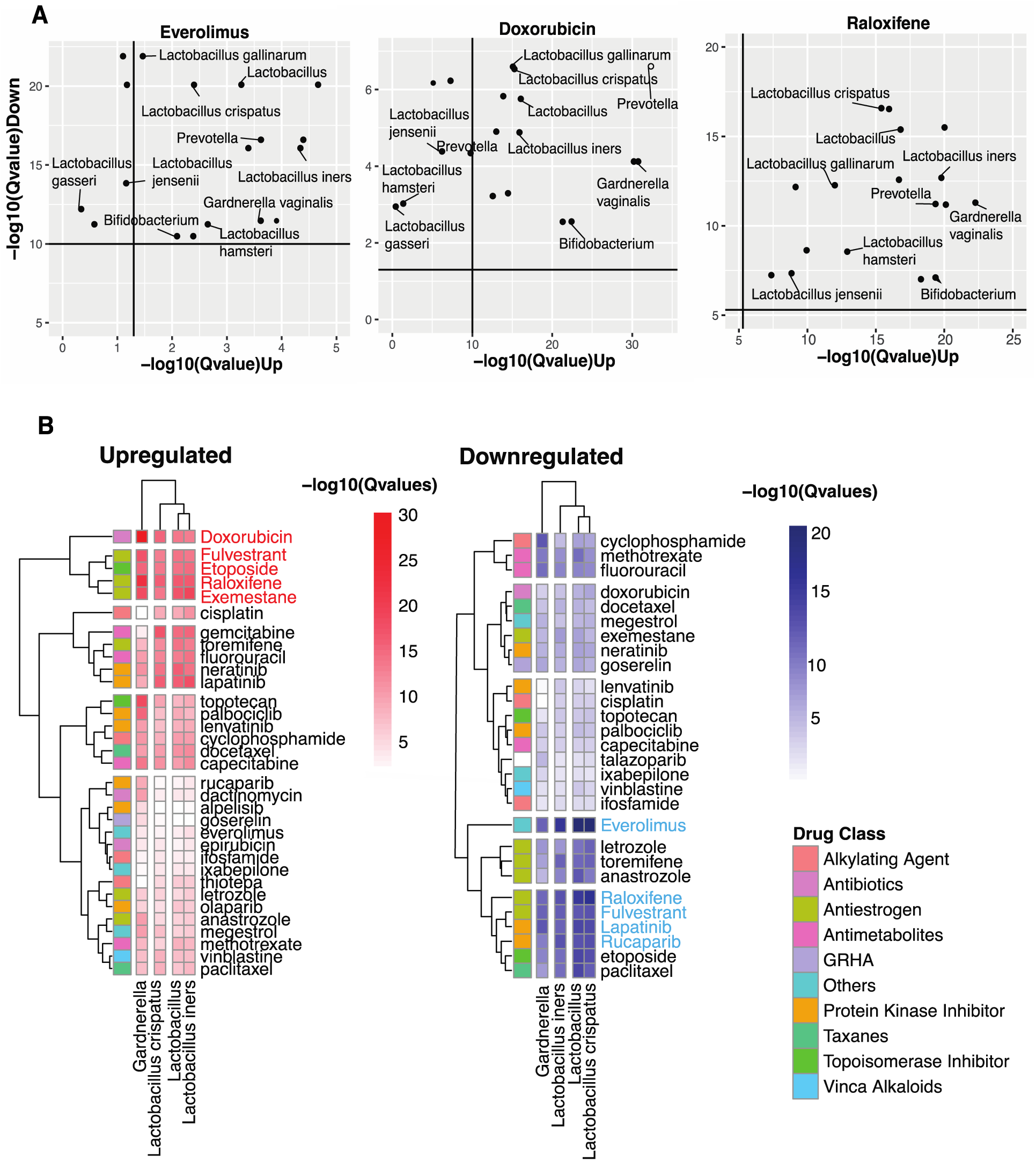
Identifying Vaginal Microbes with Anti-Cancer Therapeutic Potential. **(A)** Scatter plots of Everolimus, Doxorubicin, and Raloxifene show microbes with host-impacting gene signatures that significantly correspond to the drug induced changes in host up- and down-regulated genes (**B**) Heatmap showing similarities between microbe and anti-cancer drug in upregulated and downregulated gene signatures using the -log10(Qvalues). Drug Class key shows the different classes to which anti-cancer drugs belong.

We focused on four key taxa present in greater than 70% of human samples: *Lactobacillus, Lactobacillus iners, Lactobacillus crispatus,* and *Gardnerella* (**Figure 2B**). *Lactobacillus* species showed the highest number of similarities with drugs that belong to the antiestrogen, and protein kinase inhibitor drug class, and high similarity with drugs in the antimetabolites, topoisomerase inhibitor, antibiotics, and alkylating agents drug class. While *Gardnerella* showed a some similarities to the *Lactobacillus* species, its highest number of similarities is with drugs in the protein kinase inhibitor drug class.

### Identification of vaginal metabolites with anti-cancer drug similarity based on gene signature alignment

Using the participant-derived vaginal metabolomics and vaginal epithelial transcriptomics data, 23 codedIDs matched, and we calculated partial correlation coefficients between 99 metabolites and 23,482 host genes while holding for Lactobacillus dominance across participants to generate metabolite-RNA partial Spearman correlation coefficients. Gene lists for 99 metabolites were generated (adjusted p < 0.05) and Fisher’s exact test was employed to compare the gene signatures of 99 metabolites identified through Spearman correlation to post-treatment gene expression signatures for the same 35 chemotherapeutic drugs we assessed against vaginal microbes. When analyzing the relationship between metabolites and drugs through gene similarity, a notable pattern emerges that aligns with observations from microbe-to-drug gene similarity analysis. In both cases, specific sets of drugs show the highest significance. For example, drugs such as Doxorubicin, Fulvestrant, Exemestane, Etoposide, and Raloxifene exhibit the greatest significance for upregulated gene similarity. indicating that these drugs activate genes in ways that resemble the action of certain metabolites. was observed where the same sets of drugs have the highest significance **(Figure 3A)**. While, drugs like Everolimus, Raloxifene, Fulvestrant, Etoposide, and Fluorouracil are most significant for downregulated gene similarity, suggesting that they suppress gene expression similarly to certain metabolites (**Figure 3B**). More than 40 metabolites showed significant similarities in host gene regulation to 24 anti-cancer drugs (adjusted p <0.05), further highlighting the intricate relationship between drug action and metabolite-induced gene regulation (**Figure 3A & 3B**).

**Figure 3:**
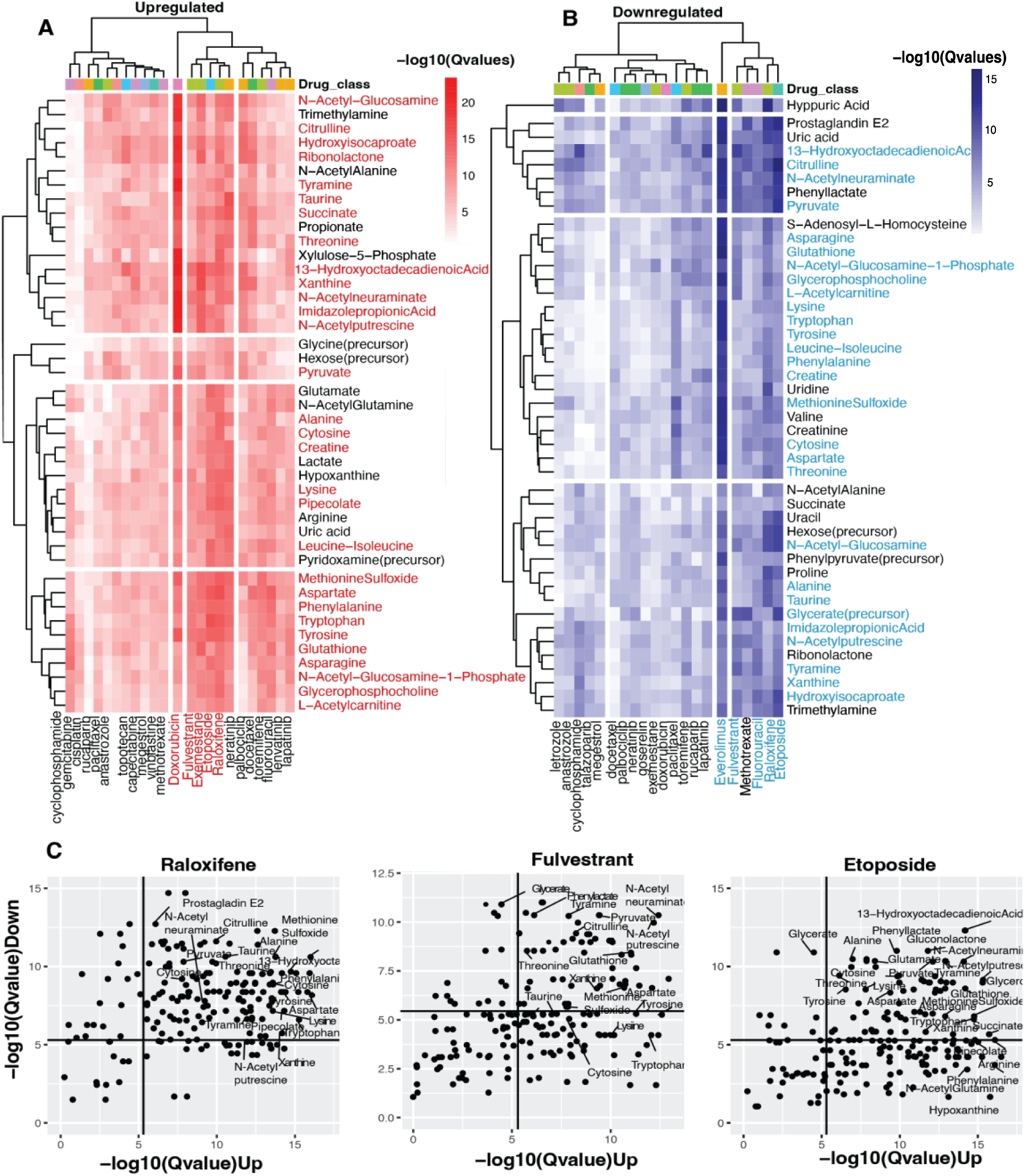
Identifying Vaginal Metabolites with Anti-Cancer Potential. **(A)** Heatmap showing similarities between metabolite and anti-cancer drug in downregulated gene signature. **(B**) Heatmap showing similarities between metabolite and anti-cancer drug in upregulated gene signature. Drug Class key shows the different classes to which anti-cancer drugs belong. **(C**) Scatter plots of Raloxifene, Fulvestrant, and Etoposide show metabolites with host-impacting gene signatures that significantly correspond to the drug induced changes in host up- and down-regulated genes.

We identified the various metabolites in drug clusters of drugs including Fulvestrant, Raloxifene, and Etoposide, showing both upregulation and downregulation signatures. These metabolites include essential amino acids like Threonine, Phenylalanine, Tryptophan, Lysine, and Leucine-Isoleucine, as well as non-essential amino acids like Alanine, Methionine Sulfoxide, Taurine, Asparagine, Tyrosine, Aspartate, and Citrulline. Additionally, other compounds such as peptides (Creatine, Glutathione), sugars (N-Acetylglucosamine, N-Acetylneuraminate), and unsaturated fatty acid (13-hydroxyoctadecadienoic acid) were identified. Moreover, metabolites involved in the synthesis or metabolism of amino acids and sugars were found including N-Acetylglucosamine-1-phospahate, L-Acetylcartinine, Hydroisocaproate, Imidazole Propionic acid, and Pipecolate. Other significant compounds include amines (N-Acetylputrescine, Tyramine), esters (Glycerophosphocholine, Glycerate), nucleic acids (Xanthine, Cytosine), and keto-acids such as Pyruvate **(Figure 3A** & **3B, Table S5**).

Fulvestrant, Raloxifene, and Etoposide were drugs with significant similarities across metabolites for upregulation and downregulation of host gene signatures. Raloxifene and Fulvestrant are antiestrogens that bind to estrogen receptors, where Raloxifene activates estrogenic pathways and blockades tissues that express estrogen receptors^37^. While Fulvestrant activates antiestrogenic effects, it inhibits receptor dimerization and enhances estrogen receptor degradation^38^. Conversely, Etoposide is a topoisomerase inhibitor; it inhibits DNA synthesis by forming a complex with DNA and topoisomerase II and hinders replication^39^ (**Figure 3C**).

### *Lactobacillus crispatus*-produced Taurine inhibits the growth of endometrial cancer cells

Several studies have shown *L. iners*, *L. crispatus*, and *G. vaginalis* as major vaginal microbes. Dominance in *L. crispatus* is associated with healthy conditions*, L. iners* is associated with a transitionary, less stable healthy condition, and *G. vaginalis* is associated with sub-optimal conditions^35, 34, 40^. To understand the relationship between vaginal microbes and metabolites that showed similarities with anti-cancer drugs, we cultured each of these microbes *in vitro* in suspension culture and extracted the conditioned media for metabolomics analysis. The objective was to identify vaginal strains capable of producing, or potentially consuming, the 28 anti-cancer metabolites identified through our analysis. Eighteen key metabolites were measured in the bacterial metabolomics, indicating these can be vaginal microbe-derived (**Figure 4A**). Of these, *L. crispatus* uniquely produced Taurine and Cytosine, with *L. iners* showing evidence of Taurine and Cytosine consumption, while *G. vaginalis* neither produces or consumes them (**Figure 4B**). These data show that *L. crispatus* is selectively producing metabolites with potential anti-cancer activity, *G. vaginalis* neither produces or consumes these metabolites, while *L. iners* consumes them, potentially reducing their effectiveness in inhibiting cancer activity.

**Figure 4:**
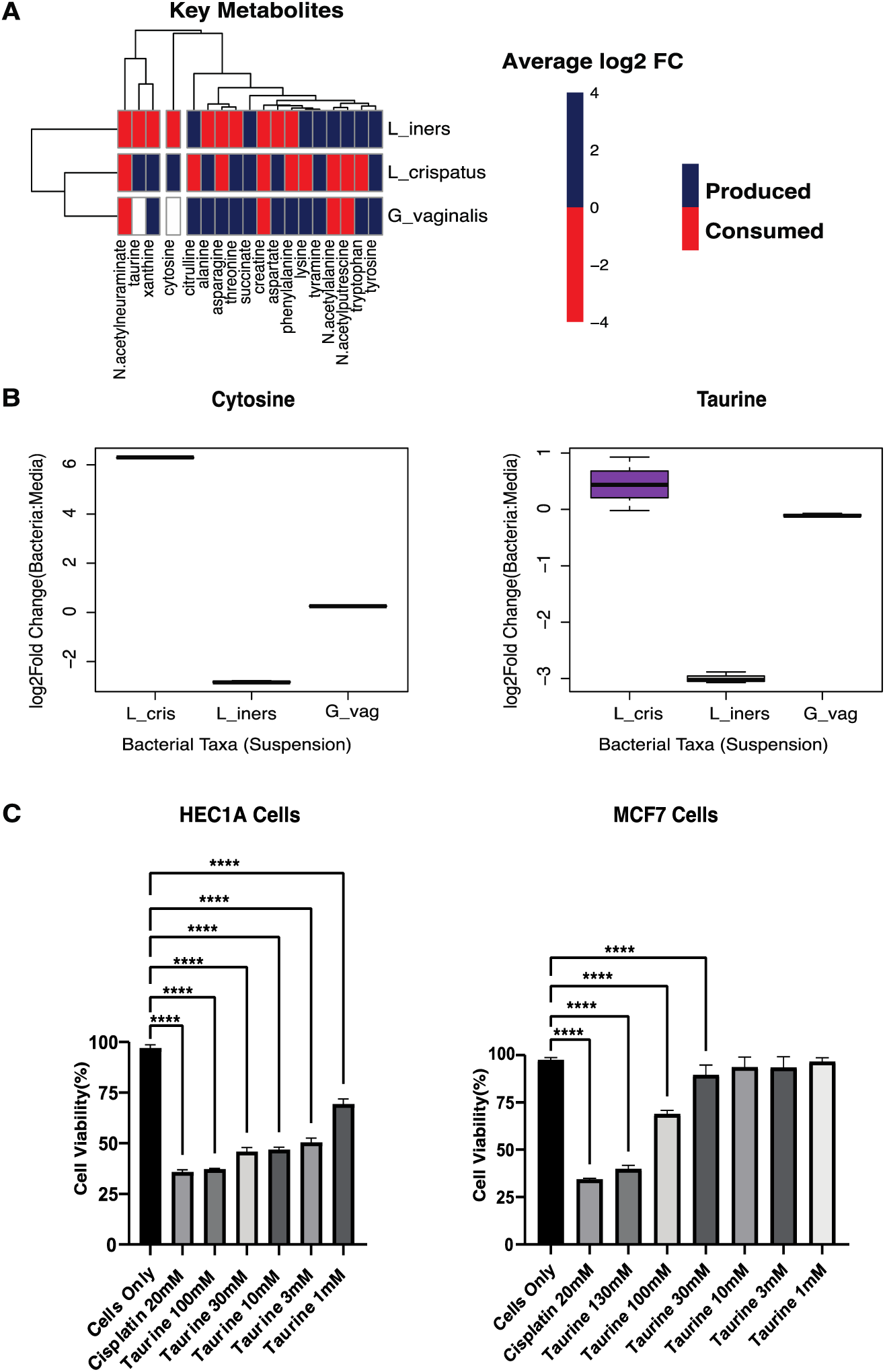
Metabolomics of Cultured Bacteria. **(A)** Heatmap showing the production/consumption of 18 Key Metabolites by L_crispatus = *Lactobacillus crispatus*. L_iners = *Lactobacillus iners*. G_vaginalis = *Gardnerella* vaginalis. **(B)** Boxplots showing the production of Cytosine and Taurine by L_cris = *Lactobacillus crispatus*. L_iners = *Lactobacillus iners*. G_vag = *Gardnerella vaginalis*. **(C)** Cell viability was determined using a CCK8 assay. Human endometrial cancer cells (HEC1A) and Human breast cancer cells (MCF7) were treated with Taurine at various concentrations 100mM, 30mM, 10mM, 3mM, 1mM for 24 h and spectrophotometrically measured at 450 nm. Cisplatin(positive control; chemotherapeutic drug) and all concentrations of Taurine significantly lowered the cell viability of HEC1A cells while only the 30mM, 100mM and130mM concentrations of Taurine significantly lowered the cell viability of MCF7 cells when compared to Cells Only (no treatment). A one-way ANOVA was used for statistical analysis. A p-value of < 0.05 was considered significant. **** p<0.0001. ns – not significant.

In order to assess whether these vaginal microbe-derived metabolites showed anti-cancer activity, we screened Taurine in an *in vitro* cell viability assay against MCF7 cells, the cell line used in the LINCS database for gene signature generation, and HEC1A cells, an endometrial cancer cell line. Taurine was selected for assessment due to studies that indicated that it might possess anti-cancer properties^41, 42^, though these studies were done in MCF7 breast cancer cells and not in gynecologic cancers. Other studies have also shown that Taurine aids the synthesis and healthy detoxification of estrogen^43^. HEC1A meets our criteria for a gynecologic cancer type in which Taurine has not been tested as an anti-cancer therapy.

The viability of HEC1A and MCF7 cells was assesed following treatment with Taurine at different concentrations (130mM, 100mM, 30mM, 10mM, 3mM, 1mM and 20 mM) for 24 hours. The positive control, Cisplatin, was also applied under the same conditions. Cell viability was measured using the CCK8 assay, which provides quantitative data on cell metabolic activity and survival. Cisplatin and all the concentrations of Taurine significantly (P< 0.001) lowered HEC1A cell viability while only 30mM, 100mM, and 130mM concentrations of Taurine significantly lowered MCF7 cells when compared to cells Only (no treatment) (**Figure 4C**). The results indicate that the vaginal microbiome-derived *L. crispatus* metabolite Taurine reduces the viability of HEC1A cells and is more potent in that gynecologic cancer context than in the MCF7 breast cancer cell line context.

## Discussion

We developed a novel systems biology approach to characterize microbes and microbial products’ function relative to known drugs, which we termed microbiome pharmacology analysis: Pharmacobiome. To explore this concept, we integrated vaginal microbiome multi-omics data from HIV-negative study participants and *in vitro* drug screening transcriptomics data to identify vaginal microbes and microbial products with therapeutic potential against female reproductive cancers. *Lactobacillus, Lactobacillus iners, Lactobacillus crispatus*, and *Gardnerella* emerged as top microbes detected in above 70% of patient samples. This agrees with findings in a study carried out by Ravel *et al.* that established the five major types of vaginal microbiota termed community state types (CST), where four are *Lactobacillus* dominant species; *Lactobacillus crispatus, Lactobacillus gasseri, Lactobacillus iners* and *Lactobacillus jensenii* (CST-I, II, III, and while the fifth is non-*Lactobacillus* dominant but instead dominated by polymicrobial obligate anaerobes like *Gardnerella* (CST-IV)^20^.

Our results showed that 28 metabolites and three microbes, *L.crispatus, L. iners,* and *Gardnerell*a, were significantly similar in their host-associated gene signatures to host gene sigantures of one or more anti-cancer drugs. We performed untargeted metabolomics on cultured *L.crispatus, L. iners,* and *G.vaginalis* to identify which of the micribes produce anti-cancer metabolites. Since *L. crispatus* is generally associated with an optimal, beneficial vaginal microbiota, information on the metabolites produced might better inform how this microbe associates with cancer protection and anti-cancer properties. Similarly, consumption of metabolites with anti-cancer potential by *G. vaginalis* or other bacteria may hinder the effectiveness of cancer treatments. Given these potential consequences, it’s essential to monitor and understand the interactions between bacteria and possible anti-cancer metabolites in the context of cancer treatment to ensure the best possible outcomes for patients.

The convergence of our computational and experimental bacterial metabolomics analysis pointed toward *L. crispatus* produced Taurine as a potential vaginal microbiome-derived anti-cancer agent. Taurine is one of the most abundant intracellular amino acids in humans^44^, and numerous studies have highlighted the antioxidant and anti-inflammatory properties of Taurine^45, 46^. Nonetheless, sparse research examines its impact on tumors, and any antitumor mechanism remains elusive. To test our hypothesis, the anti-cancer activity of Taurine was assessed by treating HEC1A endometrial cancer and MCF7 breast cancer cell lines with varying concentrations of Taurine for 24h. Taurine concentrations within cells, particularly in excitable tissues, can reach up to 70 mM, such as in the retina’s photoreceptor cells^47^. However, at artificially elevated concentrations ranging from 130 mM to as low as 1 mM, taurine significantly reduced the viability of HEC1A endometrial cancer cells. While, MCF7 breast cancer cells only showed significant reduction in cell viability at concentrations of 30 mM or above. This indicates that taurine’s cytotoxic potential is cell-type dependent.

While taurine is essential in various physiological functions like redox balance and cell volume regulation, high concentrations may disrupt cell metabolism and growth, triggering protective responses or promoting cell death^48^. This suggests that while taurine has vital roles at normal physiological levels, excessive amounts can exert a detrimental effect on specific cell lines like HEC1A. Further studies may need to be conducted to determine if Taurine induces apoptosis in other gynecological cancers, and in primary non-cancerous cells, to understand the molecular mechanism and differences in cytotoxicity in detail.

The result of our analysis shows that *L. crispatus* induces host gene signatures similar to known chemotherapeutic drugs, specifically Raloxifene, Fulvestrant, Etoposide and Lapatinib and is capable of producing metabolites like taurine that have anti-cancer potential. This opens up an avenue for *Lactobacillus crispatus* in cancer therapy. Using *Lactobacillus crispatus* or microbial products may have advantages over currently accepted probiotics or drug administration. Vaginal microbiota-derived molecules are emerging as a potential novel approach for the treatment of gynecologic cancers, presenting several advantages over conventional chemotherapeutic drugs. Firstly, unlike broad-spectrum chemotherapy agents which can cause harm to healthy cells, these molecules may exert targeted effects^49^. They have the potential to selectively target specific mechanisms involved in cancer development or progression. Secondly, owing to their origin from the host’s microbiome, they are likely to be better tolerated with fewer side effects compared to traditional chemotherapy drugs^50^. Thirdly, considering the role of the vaginal microbiome in immune function, these molecules may have the capacity to stimulate the immune system to combat cancer cells^51, 52^.

It is crucial to acknowledge that this field of research is still in its infancy. Further investigations are necessary to ascertain the efficacy and safety of these molecules in the treatment of gynecologic cancers^53^. Nevertheless, the notable advantages they offer underscore the potential of vaginal microbiota and derived molecules as a promising avenue for future cancer therapy.

In conclusion, this study shows that commensal microbes like *Lactobacillus crispatu*s and metabolites like Taurine are capable of inducing cytotoxic effects on cancer cells comparable to chemotherapeutic drugs. This quantitative link made possible by our Pharmacobiome approach opens the door to a multitude of potential opportunities for therapy in cancer patients. The use of microbial products for cancer therapy is not limited to the bacteria and metabolites discussed within this study, and it is a field with massive potential for future growth. In that effort, Pharmacobiome analysis based on profiling the therapeutic actions of human microbiome communities has great potential to discover novel therapeutic strategies in gynecologic cancers and other conditions.

## Supporting information

Table S3- gene list

Table S4- gene list

## Acknowledgments

The authors would like to thank the Partners PrEP-BV Study Team and the participants whose data was used in this study. We are grateful for the opportunity to work with the team to advance other areas of gynecologic health.

## Funding

This work was supported by a grant from the Good Ventures Foundation, startup funding from Purdue and Case Western Reserve University, and the Eunice Kennedy Shriver National Institute of Child Health & Human Development of the National Institutes of Health under Award Number R01HD110367. Generation of the omics data used for this study was supported through funding from R01 AI111738. The Partners PrEP Study was supported through a research grant (ID #47674) from the Bill and Melinda Gates Foundation. The content is solely the responsibility of the authors and does not necessarily represent the official views of the National Institutes of Health.

## Authors Contribution

Conceptualization: DCL & DKB; Methodology, DCL, SJ, DKB; Visualization: DCL.; Writing – original draft, DCL & DKB.; Writing – review & editing: DCL, ARB, KB, AL, RDM, LNR, MP, XH, EI, NM, SK, TRM, EK, SMH, CL, FLC, JMB, CC, FH, JL, LNG, DKB.

## Notes

### Competing Interest Statement

The authors have declared no competing interest.

### Summary of Updates

Author names and affiliations were updated in the revised paper

## References

(1) Smith, S. B.; Ravel, J. The Vaginal Microbiota, Host Defence and Reproductive Physiology. J Physiol 2017, 595 (2), 451–463. 10.1113/JP271694.

(2) Gajer, P.; Brotman, R. M.; Bai, G.; Sakamoto, J.; Schütte, U. M. E.; Zhong, X.; Koenig, S. S. K.; Fu, L.; Ma, Z. S.; Zhou, X.; Abdo, Z.; Forney, L. J.; Ravel, J. Temporal Dynamics of the Human Vaginal Microbiota. Sci Transl Med 2012, 4 (132), 132ra52. 10.1126/scitranslmed.3003605.

(3) Chen, X.; Lu, Y.; Chen, T.; Li, R. The Female Vaginal Microbiome in Health and Bacterial Vaginosis. Front Cell Infect Microbiol 2021, 11, 631972. 10.3389/fcimb.2021.631972.

(4) Barrientos-Durán, A.; Fuentes-López, A.; de Salazar, A.; Plaza-Díaz, J.; García, F. Reviewing the Composition of Vaginal Microbiota: Inclusion of Nutrition and Probiotic Factors in the Maintenance of Eubiosis. Nutrients 2020, 12 (2). 10.3390/nu12020419.

(5) Martin, D. H.; Marrazzo, J. M. The Vaginal Microbiome: Current Understanding and Future Directions. J Infect Dis 2016, 214 Suppl 1 (Suppl 1), S36–41. 10.1093/infdis/jiw184.

(6) Lewis, F. M. T.; Bernstein, K. T.; Aral, S. O. Vaginal Microbiome and Its Relationship to Behavior, Sexual Health, and Sexually Transmitted Diseases. Obstet Gynecol 2017, 129 (4), 643–654. 10.1097/AOG.0000000000001932.

(7) Nunn, K. L.; Forney, L. J. Unraveling the Dynamics of the Human Vaginal Microbiome. Yale J Biol Med 2016, 89 (3), 331–337.

(8) Lev-Sagie, A.; De Seta, F.; Verstraelen, H.; Ventolini, G.; Lonnee-Hoffmann, R.; Vieira-Baptista, P. The Vaginal Microbiome: II. Vaginal Dysbiotic Conditions. J Low Genit Tract Dis 2022, 26 (1), 79–84. 10.1097/LGT.0000000000000644.

(9) Rajagopala, S. V.; Vashee, S.; Oldfield, L. M.; Suzuki, Y.; Venter, J. C.; Telenti, A.; Nelson, K. E. The Human Microbiome and Cancer. Cancer Prev Res (Phila) 2017, 10 (4), 226–234. 10.1158/1940-6207.CAPR-16-0249.

(10) Garrett, W. S. Cancer and the Microbiota. Science 2015, 348 (6230), 80–86. 10.1126/science.aaa4972.

(11) Zadora, P. K.; Chumduri, C.; Imami, K.; Berger, H.; Mi, Y.; Selbach, M.; Meyer, T. F.; Gurumurthy, R. K. Integrated Phosphoproteome and Transcriptome Analysis Reveals Chlamydia-Induced Epithelial-to-Mesenchymal Transition in Host Cells. Cell Rep 2019, 26 (5), 1286–1302.e8. 10.1016/j.celrep.2019.01.006.

(12) Ventolini, G.; Vieira-Baptista, P.; De Seta, F.; Verstraelen, H.; Lonnee-Hoffmann, R.; Lev-Sagie, A. The Vaginal Microbiome: IV. The Role of Vaginal Microbiome in Reproduction and in Gynecologic Cancers. J Low Genit Tract Dis 2022, 26 (1), 93–98. 10.1097/LGT.0000000000000646.

(13) Walboomers, J. M.; Jacobs, M. V.; Manos, M. M.; Bosch, F. X.; Kummer, J. A.; Shah, K. V.; Snijders, P. J.; Peto, J.; Meijer, C. J.; Muñoz, N. Human Papillomavirus Is a Necessary Cause of Invasive Cervical Cancer Worldwide. J Pathol 1999, 189 (1), 12–19. 10.1002/(SICI)1096-9896(199909)189:1<12::AID-PATH431>3.0.CO;2-F.

(14) Mitra, A.; MacIntyre, D. A.; Lee, Y. S.; Smith, A.; Marchesi, J. R.; Lehne, B.; Bhatia, R.; Lyons, D.; Paraskevaidis, E.; Li, J. V.; Holmes, E.; Nicholson, J. K.; Bennett, P. R.; Kyrgiou, M. Cervical Intraepithelial Neoplasia Disease Progression Is Associated with Increased Vaginal Microbiome Diversity. Sci Rep 2015, 5, 16865. 10.1038/srep16865.

(15) Walther-António, M. R. S.; Chen, J.; Multinu, F.; Hokenstad, A.; Distad, T. J.; Cheek, E. H.; Keeney, G. L.; Creedon, D. J.; Nelson, H.; Mariani, A.; Chia, N. Potential Contribution of the Uterine Microbiome in the Development of Endometrial Cancer. Genome Med 2016, 8 (1), 122. 10.1186/s13073-016-0368-y.

(16) Banerjee, S.; Tian, T.; Wei, Z.; Shih, N.; Feldman, M. D.; Alwine, J. C.; Coukos, G.; Robertson, E. S. The Ovarian Cancer Oncobiome. Oncotarget 2017, 8 (22), 36225–36245. 10.18632/oncotarget.16717.

(17) Chan, P. J.; Seraj, I. M.; Kalugdan, T. H.; King, A. Prevalence of Mycoplasma Conserved DNA in Malignant Ovarian Cancer Detected Using Sensitive PCR-ELISA. Gynecol Oncol 1996, 63 (2), 258–260. 10.1006/gyno.1996.0316.

(18) Ibrahim, Y.; Emara, M.; Vyas, V.; Awadi, S.; Jaroslav, N.; Khodry, A.; Essam, T.; Amanguno, H.; Purohit, P. Synchronous Occurrence of Brucellosis and Ovarian Cancer-A Case Report. Austral - Asian Journal of Cancer 2007, 6, 257–259.

(19) Chase, D.; Goulder, A.; Zenhausern, F.; Monk, B.; Herbst-Kralovetz, M. The Vaginal and Gastrointestinal Microbiomes in Gynecologic Cancers: A Review of Applications in Etiology, Symptoms and Treatment. Gynecol Oncol 2015, 138 (1), 190–200. 10.1016/j.ygyno.2015.04.036.

(20) Ravel, J.; Gajer, P.; Abdo, Z.; Schneider, G. M.; Koenig, S. S. K.; McCulle, S. L.; Karlebach, S.; Gorle, R.; Russell, J.; Tacket, C. O.; Brotman, R. M.; Davis, C. C.; Ault, K.; Peralta, L.; Forney, L. J. Vaginal Microbiome of Reproductive-Age Women. Proc Natl Acad Sci U S A 2011, 108 *Suppl 1* (Suppl 1), 4680–4687. 10.1073/pnas.1002611107.

(21) Łaniewski, P.; Ilhan, Z. E.; Herbst-Kralovetz, M. M. The Microbiome and Gynaecological Cancer Development, Prevention and Therapy. Nat Rev Urol 2020, 17 (4), 232–250. 10.1038/s41585-020-0286-z.

(22) Kudela, E.; Holubekova, V.; Kolkova, Z.; Kasubova, I.; Samec, M.; Mazurakova, A.; Koklesova, L. Vaginal Microbiome and Its Role in HPV Induced Cervical Carcinogenesis. In Microbiome in 3P Medicine Strategies: The First Exploitation Guide; Boyko, N., Golubnitschaja, O., Eds.; Springer International Publishing: Cham, 2023; pp 43–86. 10.1007/978-3-031-19564-8_3.

(23) Manyam, M.; Stephens, A. J.; Kennard, J. A.; LeBlanc, J.; Ahmad, S.; Kendrick, J. E.; Holloway, R. W. A Phase 1b Study of Intraperitoneal Oncolytic Viral Immunotherapy in Platinum-Resistant or Refractory Ovarian cancer11Part of This Study Was Presented at the 54th Annual Meeting of the American Society of Clinical Oncology (ASCO), June 1–5, 2018, Chicago, IL, USA. Gynecologic Oncology 2021, 163 (3), 481–489. 10.1016/j.ygyno.2021.10.069.

(24) Kyrgiou, M.; Moscicki, A.-B. Vaginal Microbiome and Cervical Cancer. Seminars in Cancer Biology 2022, 86, 189–198. 10.1016/j.semcancer.2022.03.005.

(25) Sipos, A.; Ujlaki, G.; Mikó, E.; Maka, E.; Szabó, J.; Uray, K.; Krasznai, Z.; Bai, P. The Role of the Microbiome in Ovarian Cancer: Mechanistic Insights into Oncobiosis and to Bacterial Metabolite Signaling. Molecular Medicine 2021, 27 (1), 33. 10.1186/s10020-021-00295-2.

(26) Chambers, L. M.; Kuznicki, M.; Yao, M.; Chichura, A.; Gruner, M.; Reizes, O.; Debernardo, R.; Rose, P. G.; Michener, C.; Vargas, R. Impact of Antibiotic Treatment during Platinum Chemotherapy on Survival and Recurrence in Women with Advanced Epithelial Ovarian Cancer. Gynecologic Oncology 2020, 159 (3), 699–705. 10.1016/j.ygyno.2020.09.010.

(27) Fernandes, M. R.; Aggarwal, P.; Costa, R. G. F.; Cole, A. M.; Trinchieri, G. Targeting the Gut Microbiota for Cancer Therapy. Nature Reviews Cancer 2022, 22 (12), 703–722. 10.1038/s41568-022-00513-x.

(28) Berard, A. R.; Brubaker, D. K.; Birse, K.; Lamont, A.; Mackelprang, R. D.; Noël-Romas, L.; Perner, M.; Hou, X.; Irungu, E.; Mugo, N.; Knodel, S.; Muwonge, T. R.; Katabira, E.; Hughes, S. M.; Levy, C.; Calienes, F. L.; Lauffenburger, D. A.; Baeten, J. M.; Celum, C.; Hladik, F.; Lingappa, J.; Burgener, A. D. Vaginal Epithelial Dysfunction Is Mediated by the Microbiome, Metabolome, and mTOR Signaling. Cell Rep 2023, 42 (5), 112474. 10.1016/j.celrep.2023.112474.

(29) Baeten, J. M.; Donnell, D.; Ndase, P.; Mugo, N. R.; Campbell, J. D.; Wangisi, J.; Tappero, J. W.; Bukusi, E. A.; Cohen, C. R.; Katabira, E.; Ronald, A.; Tumwesigye, E.; Were, E.; Fife, K. H.; Kiarie, J.; Farquhar, C.; John-Stewart, G.; Kakia, A.; Odoyo, J.; Mucunguzi, A.; Nakku-Joloba, E.; Twesigye, R.; Ngure, K.; Apaka, C.; Tamooh, H.; Gabona, F.; Mujugira, A.; Panteleeff, D.; Thomas, K. K.; Kidoguchi, L.; Krows, M.; Revall, J.; Morrison, S.; Haugen, H.; Emmanuel-Ogier, M.; Ondrejcek, L.; Coombs, R. W.; Frenkel, L.; Hendrix, C.; Bumpus, N. N.; Bangsberg, D.; Haberer, J. E.; Stevens, W. S.; Lingappa, J. R.; Celum, C. Antiretroviral Prophylaxis for HIV Prevention in Heterosexual Men and Women. N Engl J Med 2012, 367 (5), 399–410. 10.1056/NEJMoa1108524.

(30) Targeted Therapy Drug List by Cancer Type - NCI. https://www.cancer.gov/about-cancer/treatment/types/targeted-therapies/approved-drug-list (accessed 2023-10-23).

(31) Todd Golub, A. S. L1000 Dataset - Small Molecule, Nucleic Acid Perturbagens - LINCS Phase 2 (December 2021). 2021.

(32) Reid, G. The Scientific Basis for Probiotic Strains of Lactobacillus. Appl Environ Microbiol 1999, 65 (9), 3763–3766. 10.1128/AEM.65.9.3763-3766.1999.

(33) Swidsinski, A.; Mendling, W.; Loening-Baucke, V.; Ladhoff, A.; Swidsinski, S.; Hale, L. P.; Lochs, H. Adherent Biofilms in Bacterial Vaginosis. Obstet Gynecol 2005, 106 (5 Pt 1), 1013–1023. 10.1097/01.AOG.0000183594.45524.d2.

(34) Petrova, M. I.; Reid, G.; Vaneechoutte, M.; Lebeer, S. Lactobacillus Iners: Friend or Foe? Trends Microbiol 2017, 25 (3), 182–191. 10.1016/j.tim.2016.11.007.

(35) Jimenez Nicole R.; Maarsingh Jason D.; Łaniewski Paweł; Herbst-Kralovetz Melissa M. Commensal Lactobacilli Metabolically Contribute to Cervical Epithelial Homeostasis in a Species-Specific Manner. mSphere 2023, 8 (1), e00452–22. 10.1128/msphere.00452-22.

(36) Evans, A. M.; DeHaven, C. D.; Barrett, T.; Mitchell, M.; Milgram, E. Integrated, Nontargeted Ultrahigh Performance Liquid Chromatography/Electrospray Ionization Tandem Mass Spectrometry Platform for the Identification and Relative Quantification of the Small-Molecule Complement of Biological Systems. Anal Chem 2009, 81 (16), 6656–6667. 10.1021/ac901536h.

(37) LiverTox: Clinical and Research Information on Drug-Induced Liver Injury; Bethesda (MD), 2012.

(38) Carlson, R. W. The History and Mechanism of Action of Fulvestrant. Clin Breast Cancer 2005, 6 *Suppl 1*, S5–8. 10.3816/cbc.2005.s.008.

(39) Montecucco, A.; Zanetta, F.; Biamonti, G. Molecular Mechanisms of Etoposide. EXCLI J 2015, 14, 95–108. 10.17179/excli2015-561.

(40) Castro, J.; Machado, D.; Cerca, N. Unveiling the Role of Gardnerella Vaginalis in Polymicrobial Bacterial Vaginosis Biofilms: The Impact of Other Vaginal Pathogens Living as Neighbors. ISME J 2019, 13 (5), 1306–1317. 10.1038/s41396-018-0337-0.

(41) Zhang, X.; Lu, H.; Wang, Y.; Liu, C.; Zhu, W.; Zheng, S.; Wan, F. Taurine Induces the Apoptosis of Breast Cancer Cells by Regulating Apoptosis-Related Proteins of Mitochondria. Int J Mol Med 2015, 35 (1), 218–226. 10.3892/ijmm.2014.2002.

(42) Shennan, D. B.; Thomson, J. Estrogen Regulation and Ion Dependence of Taurine Uptake by MCF-7 Human Breast Cancer Cells. Cellular & Molecular Biology Letters 2007, 12 (3), 396–406. 10.2478/s11658-007-0011-4.

(43) Li, L.; Lu, C.; Zhang, D.; Liu, H.; Cui, S. Taurine Promotes Estrogen Synthesis by Regulating microRNA-7a2 in Mice Ovarian Granulosa Cells. Biochemical and Biophysical Research Communications 2022, 626, 129–134. 10.1016/j.bbrc.2022.07.084.

(44) Lourenço, R.; Camilo, M. E. Taurine: A Conditionally Essential Amino Acid in Humans? An Overview in Health and Disease. Nutr Hosp 2002, 17 (6), 262–270.

(45) Jong, C. J.; Sandal, P.; Schaffer, S. W. The Role of Taurine in Mitochondria Health: More Than Just an Antioxidant. Molecules 2021, 26 (16). 10.3390/molecules26164913.

(46) Baliou, S.; Adamaki, M.; Ioannou, P.; Pappa, A.; Panayiotidis, M. I.; Spandidos, D. A.; Christodoulou, I.; Kyriakopoulos, A. M.; Zoumpourlis, V. Protective Role of Taurine against Oxidative Stress (Review). Mol Med Rep 2021, 24 (2), 605. 10.3892/mmr.2021.12242.

(47) Seidel, U.; Huebbe, P.; Rimbach, G. Taurine: A Regulator of Cellular Redox Homeostasis and Skeletal Muscle Function. Molecular Nutrition & Food Research 2019, 63 (16), 1800569. 10.1002/mnfr.201800569.

(48) Centeno, D.; Farsinejad, S.; Kochetkova, E.; Volpari, T.; Gladych-Macioszek, A.; Klupczynska-Gabryszak, A.; Polotaye, T.; Greenberg, M.; Kung, D.; Hyde, E.; Alshehri, S.; Pavlovic, T.; Sullivan, W.; Plewa, S.; Vakifahmetoglu-Norberg, H.; Monsma, F. J.; Muller, P. A. J.; Matysiak, J.; Zaborowski, M. P.; DiFeo, A.; Norberg, E.; Martin, L. A.; Iwanicki, M. Modeling of Intracellular Taurine Levels Associated with Ovarian Cancer Reveals Activation of P53, ERK, mTOR and DNA-Damage-Sensing-Dependent Cell Protection. Nutrients 2024, 16 (12). 10.3390/nu16121816.

(49) Lin, X.; Zheng, W.; Zhao, X.; Zeng, M.; Li, S.; Peng, S.; Song, T.; Sun, Y. Microbiome in Gynecologic Malignancies: A Bibliometric Analysis from 2012 to 2022. Transl Cancer Res 2024, 13 (4), 1980–1996. 10.21037/tcr-23-1769.

(50) Cocomazzi, G.; Del Pup, L.; Contu, V.; Maggio, G.; Parmegiani, L.; Ciampaglia, W.; De Ruvo, D.; Faioli, R.; Maglione, A.; Baldini, G. M.; Baldini, D.; Pazienza, V. Gynecological Cancers and Microbiota Dynamics: Insights into Pathogenesis and Therapy. Int J Mol Sci 2024, 25 (4). 10.3390/ijms25042237.

(51) Rizzo, A. E.; Gordon, J. C.; Berard, A. R.; Burgener, A. D.; Avril, S. The Female Reproductive Tract Microbiome-Implications for Gynecologic Cancers and Personalized Medicine. J Pers Med 2021, 11 (6). 10.3390/jpm11060546.

(52) Xia, C.; Su, J.; Liu, C.; Mai, Z.; Yin, S.; Yang, C.; Fu, L. Human Microbiomes in Cancer Development and Therapy. MedComm (2020) 2023, 4 (2), e221. 10.1002/mco2.221.

(53) Han, M.; Wang, N.; Han, W.; Ban, M.; Sun, T.; Xu, J. Vaginal and Tumor Microbiomes in Gynecological Cancer (Review). Oncol Lett 2023, 25 (4), 153. 10.3892/ol.2023.13739.

