## Supplementary material for "Computational Microbiome Pharmacology Analysis Elucidates the Anti-Cancer Potential of Vaginal Microbes and Metabolites": Table S3- gene list

V1 "V2" "V3" "V4" "V5" "V6" "V7" "V8" "V9" "V10" "V11" "V12" "V13" "V14" "V15" "V16" "V17" "V18" "
1 c("HIST1H2BG", "HIST1H2BH", "HIST1H2BD") c("CD55", "ANXA3", "MYC", "EZR") c("ADD3", "MMP10"
2 c("BOP1", "CDKN3", "CIT", "CYP51A1", "EIF5A", "HIST1H2BD", "HIST1H2BG", "HIST1H2BH", "ITGA6", "
S100A2, "S100A7", "S100P", "SQLE", "STAT3", "TCERG1", "TCN1", "TEAD4", "THBS1", "TMEM158", "TNF
SCNN1A, "SDC1", "SEC61B", "SERPINB1", "SH3BGRL3", "SLPI", "STAT3", "TARS", "TBC1D8", "TEAD4", "TC
3 c("G3BP1", "TUBB", "EIF5A", "F12", "GSTO1", "JUP", "KLK5", "PROCR", "SLC38A1", "HIST1H2BD", "ITG
TRAPPC3, "CYP1A1", "EPHA2", "LGALS2", "ELMO1", "ITGB3BP", "MMP9", "RFTN1", "TWIST1", "IL1RN", "
CLDN4, "HMOX1", "GOLPH3L", "MUC4", "CANT1", "ST3GAL1", "NAGK", "NETO2", "CYP1A1", "EPHA2", "
MYO6, "TACSTD2", "GABRP", "S100A11", "PRKCD", "FCGBP", "BCL3", "CEACAM6", "ELF3", "RBM47", "O
4 c("SDF4", "HIST1H2BD", "F12", "LPL", "ITGA6", "KCTD5", "KLK10") c("SOX9", "SLC20A1", "EGR3", "HSP
PDE4B, "ARHGDIB", "TLR4", "PRNP", "KRT16", "RBP4", "CD53", "CLEC7A", "IFI16", "RRM2", "TACSTD2",
SLC16A1, "TMBIM1", "S100A10", "SH3BGRL3", "PLS3", "IFI16", "SERPINA3", "ANXA1", "APOLD1", "TACS
5 c("FGFR1OP", "HIST1H2BG", "HIST1H2BH", "EIF5A", "CREM") c("SLCO1B3", "SIK1") c("MID1", "CCL5")
6 c("JUP", "TNNT1", "SLC38A1", "PROCR") c("HSPB8", "MET", "TRIM16") c("ASPH", "HMGCS1", "GPR65"
7 SLC38A2 c("CSTB", "EGR2", "MED13", "OBSL1") c("ADD3", "TARP", "CYP1B1") c("ATXN7", "MVP", "TM
8 c("PSME3", "ITGA6", "L1CAM", "PPP1R9A", "HIST1H2BD", "KLK5", "SLC38A1", "EIF5A", "HIST1H2BG",
ADORA2B, "SLC7A7", "LYN", "CD84", "G0S2", "ID1", "SLC2A9", "NAMPT", "PLIN2", "PRNP", "S100A2", "
KRT5, "DDIT4", "SLC25A6") c("TBC1D8", "PF4", "DDX10", "CXADR", "FDPS", "ACOT7", "TFRC", "ELOVL6",
ITPR3, "OAS2", "AQP9", "TNFSF10", "GPRC5A", "LAPTM4B", "SPP1", "SPATS2L", "SLC16A1", "PLAT", "BS
9 c("HAT1", "SLC38A1", "HIST1H2BG", "JUP", "SLC38A2", "EIF5A", "L1CAM", "HIST1H2BD", "MBNL2", "R
BACE2, "SLC35F2", "TCN1", "SLC25A13", "UBE2H", "SIGLEC15", "CLDN4", "LIPG", "TRAPPC3", "F2RL1", "
PRSS3, "TWIST1", "HLA-DQB1", "RFTN1", "MAST4", "BMP2", "HLA-E") c("CSTB", "IRF6", "APOD", "ZFP36
DEFA4) c("ZFP36L2", "ADD3", "RABEP2", "FEZ1", "TSPAN15", "CCNL2", "ZMAT3", "SORL1", "DUSP3", "CI
GOLGB1, "TPP1", "ALDH7A1", "ANXA3", "LRP10", "CLDN7", "GRB7", "FOXO4", "RAB11FIP1", "HMOX1",
MARCKS, "RGS16", "GABRE", "HGD", "INPP4B", "SERPINB1", "LACTB2", "TSPAN31", "TXNDC9", "SPP1", "
MICALL1, "NINJ1", "SSH3", "ALDH1A3", "CDKN1A", "SERPINA3", "ALDH1L1", "DAPK1", "CEACAM6", "GC
FDPS, "RBM47", "PLAT", "VNN1", "C4BPA", "HGD", "INPP4B", "SERPINB1", "LACTB2", "TSPAN31", "AREC
) c("ZFP36L2", "SOX9", "KCNK3", "GRIA2", "REG3A", "TRIM16", "FAR2", "VAMP1", "OBSL1", "YPEL5", "KI
IDE, "PDCD6", "BAG1", "CTNND1", "MED28", "PPT2", "GLUD2", "AQP3", "KLK7", "CMTM6", "MMP1", "N
BAG1, "ERAP1", "CTNND1", "CD59", "LAMC2", "PPARG", "PPA1", "LRMP", "DKK1", "BMP2") c("FBP1", "
THBD, "UCHL3", "PRPF4", "ZC3H15", "SERPINE1", "PRNP", "LDHA", "OPN3", "ANO1") c("RARRES1", "KR
CMTM6, "LTA4H", "STK38") c("PTGDS", "MFAP2", "VGLL4", "EPB41L4B", "STC2", "PIK3R3", "GLUL", "CD
EDNRB, "TIMP1", "HIP1", "CLSTN2") c("CSTA", "PPIC", "OAZ3", "COL9A3", "TSPO", "SCNN1A", "TJP2", "T
) c("ITGBL1", "HEBP2", "HOPX", "RARRES1", "BGN", "SORBS3", "ITGA5", "STXBP2", "STC2", "PIK3R3", "T
USP6NL, "PRODH") c("HEBP2", "HOPX", "GABRP", "PRR16", "GLUL", "ELF3", "SLC2A3", "PPFIA1", "GPX2"
MMD, "LSM5", "PKIA", "IKZF1", "GLCE", "TGFA", "CISD1", "PLAT", "NPL", "SCML1", "NAMPT", "CHI3L1",
CSRP2, "SCML1", "CYB5A", "INPP4B", "SCYL3", "TSEN2", "BCL2L1", "AUTS2", "TMEM126B", "SORBS1", "
C4BPA, "HOXB6", "MS4A1", "CCDC92", "TMEM126B", "KCTD3", "ZNF33B", "AP1S2", "CCDC6", "TRAK1",
PELI1, "SPP1", "LPXN", "ZSCAN18", "HSPA12A", "GALC", "CMTM6") c("GPX4", "PIK3R3", "ZNF217", "PRR
KIF5C, "CCDC92", "RAC2", "FAM193B", "PPARG", "MYH10", "ZNF331", "SLC39A4", "FKBP11", "IFITM1",
UBXN4, "GALC", "PPA1", "RABGAP1L", "LTA4H", "RBM34") c("CDR2L", "RARRES1", "MFAP2", "ISLR", "LX
) c("HOPX", "RARRES1", "STXBP2", "FGFR3", "S100A11", "ELF3", "TACSTD2", "S100A10", "PRKAR2B", "G
10 c("GRPEL1", "HIST1H2BG", "CYP51A1", "HIST1H2BK", "GSTO1", "HIST1H2BD", "ITGA6", "EIF5A", "NA
TRAPPC3, "ACTN1", "CDKN1A", "LAMC2", "S100A2", "FLNA", "SLC25A13", "CCL20", "KRT17", "PLAT", "A
BST2, "FCGBP", "LYN", "TNFRSF21", "TSPAN1", "VAMP8", "S100P", "DAPK1", "SLPI", "SPINK1", "TBC1D8
KLF10) c("CTSK", "PPIC", "LAMB2", "AEBP1", "ITGAV", "ASPN", "NRIP1", "RARRES1", "TPBG", "IRS2", "PC
11 c("HIST1H2BG", "LPL", "NUTF2", "HIST1H2BD", "TNNT1", "VARS", "BOP1", "JUP", "NEK7", "CHMP6",

ASPN, "LYN", "ZDHHC6", "VPS37B", "TUBB6", "PIP5K1A", "VNN1", "KRT16", "TLR4", "RRAS2", "DDX21",
APOL3, "PRPF4", "KRT16", "MUC1", "GBP1", "CX3CL1", "GSTP1", "IDO1", "MAL", "BLNK", "QPCT", "ANX
SLC7A5, "TAP2", "ACTN1", "NQO2") c("RUNX3", "SCARA3", "ABCA8", "RARRES1", "ADCY7", "FILIP1L", "L
12 character(0) c("B4GALT5", "NRIP1", "SPINK5", "MREG", "TRIM16") c("ASPH", "MS4A1") c("TMEM15
13 c("CDYL", "EIF5A", "KLK10", "FZD10", "TNNT1", "NPR3", "HIST1H2BG", "SLC38A1", "ITGA6", "POLR2I
SOD2, "NR4A2", "CEBPB", "CLEC7A", "LYPD1", "CDKN1A", "FPR1", "CXCL2", "S100A7", "TACSTD2", "KYN
MAL, "SOD2", "FCGBP", "CDKN1A", "S100A7", "TACSTD2") c("MPO", "TM4SF1", "VAV3", "GTF2IRD1", "
14 c("SLC38A2", "XPOT", "EIF5A", "ITGA6", "POLR2I", "NFKBIB", "ATF4", "SLC38A1", "KLK5", "L1CAM", "
IL1RN, "NAMPT", "ASPN", "LIMA1", "CYP1A1", "F2RL1", "KRT14") c("EREG", "INHBB", "ABCA8", "TRIM1
CEACAM1, "IFITM1", "SOD2", "LCN2", "GABRP", "SWAP70", "ACTN1", "IER5", "LRP10", "GSTK1", "KLK5"
15 c("HIST1H2BD", "MLF1", "LYL1", "NAPA", "ALG5", "NDUFA7", "NUTF2", "L1CAM", "EIF5A", "CHMP6",
EAF2, "DGAT1", "F2RL1", "LYPD1", "KLF10", "GK", "IL1RN", "MMP9", "CDK6", "LYN", "RAC2", "MAST4",
KRT5, "CEACAM6", "PRDX3", "CFB", "TPD52L1", "S100A7", "SWAP70", "CNIH4", "PITX1", "ATP6V0C", "S
16 c("SNRPA1", "KCNQ2", "HIST1H2BD", "L1CAM", "F12", "KBTBD11") c("CCNL1", "CEP170", "PAK2", "
TWIST1, "CXCL6", "BMP2", "PTGS2", "ID3", "RHOBTB3") c("DUSP3", "CEP170", "KCNQ2", "MED13", "GR
MRPL13, "APOBEC3A", "CNIH4", "KRT14", "CA2", "KRT5", "BLNK", "GRB7", "HLA-B", "CFB", "IFI44", "NA
17 POLR2I c("CD55", "SPRR3", "HSPB8", "EPHB2", "EGR3") TARP c("KRT17", "SAT1", "HLA-DRA", "BAG1"
18 c("SLC38A2", "SIGMAR1", "CYP51A1", "XPOT", "L1CAM", "KLK10", "F12", "LPL", "ITGA6", "MBNL2", "
SQLE, "SRGN", "XPO7", "MID1IP1", "CD46", "CNN2", "ANXA2", "UBE2H", "CEACAM1", "SLC25A5", "PTG
SLC16A1, "CA2", "NQO2", "RPL10", "MYO1B", "PDZK1IP1", "MAP2K2", "ALDH1A3", "QPCT", "HPRT1", "
19 c("CDKN3", "VARS", "ITGA6", "KLK10", "EIF5A") c("SLCO1B3", "HNRNPA2B1", "EIF3A") c("MRPL18", "
20 c("PRPF6", "TNNT1", "CDYL", "NPR3", "EIF5A") c("LUZP1", "UGCG", "SPRR3", "SLC20A1", "MREG", "C
21 c("PPP1R9A", "NPR3", "CREM", "POLR2I", "LPL", "BOP1", "L1CAM", "HIST1H2BH", "HIST1H2BK", "CH
22 HAT1 c("OBSL1", "SPINK5", "SIK1", "SPRR3", "FOSL1") character(0) c("KRT16", "THBD", "TUBB6", "RR
23 c("SLC38A1", "HIST1H2BD", "KCTD5", "SET", "HIST1H2BK", "JUP", "MLF1", "SLC38A2", "KLK5", "MBN
SMPDL3A, "S100P", "TTC9", "KRT14", "SLC6A14", "OSBPL10", "ARHGDIB", "NAMPT", "BPGM", "CLDN7",
MYO6, "CA2", "KRT19", "FOXO4", "FOXC1", "KDM5B", "GSTZ1", "GOLGB1", "FZD6", "MBOAT7", "ATN1"
PHLDA1, "MGST3", "RAB5B") c("TACSTD2", "IFITM1", "GABRE", "DNASE1L1", "TSPAN31", "FERMT1", "G
SLC25A6, "IFNAR2", "CLCA2", "ACSL1", "PSMD10", "TBC1D8", "PDLIM1", "SH3BP4", "ID1", "BTN3A3", "
PALMD, "KRT16", "PAPSS1", "H2AFY2", "STOM", "DUOX2", "CAPNS1", "FCGBP", "GABRP", "RAP2C", "H
PHF11, "UBE2L6", "THBD", "ANXA2", "DYNLT1", "TBC1D8", "PDLIM1", "ID3", "ID1", "IL1RN", "IL13RA1",
REC8, "MAST4", "PHLDA1") c("TSPAN31", "CHI3L1", "SEC16A", "YWHAE", "SLC35D2") c("DHRS7", "CTSK
MGLL, "TPBG") c("HEBP2", "PSMD5", "GPX2", "SET", "BCAM", "CLDN10", "GCH1", "FCGBP", "GABRP", "
TPBG) c("TACSTD2", "ALDH3B2", "TGIF1", "CDS1", "LXN", "STK38L", "CXADR", "TMPRSS2", "TNFSF10", "
USP6NL, "SLC15A2", "LY6D", "KYNU", "CLCA2", "PHF11", "RNASE6", "MS4A4A", "TBC1D8", "IL13RA1", "
BCL2L2, "NCOA2", "GNLY", "TPD52", "LSM5", "TP63", "KRT15", "PCCB", "KLHDC10", "LAMP1", "DYSF", "
C4BPA, "CFH", "CD163", "TP63", "KRT15", "WASL", "LHFPL2", "FKBP11") c("SSR4", "HIST1H2BD", "MSRE
IDS, "RGL2", "LYPD3", "CASP4", "CERK", "NDUFA3", "CD163", "CDR2L", "PRKAR2B", "GALNT1", "TPBG")
PCCB, "CDO1", "MAST4", "IFRD1", "EPHX1", "FKBP11", "PHLDA1", "DYSF", "MGST3", "NUDT9", "CXCL12
WASL, "AREG", "LRRC15", "RAB5B", "TAPBP") c("ALDH3B2", "LXN", "THBS2", "ZNF140", "SNAI2", "IGF1"
24 c("SLC38A1", "KLK10", "L1CAM", "ENSA", "SPDEF", "POLR2I", "HIST1H2BK", "JUP", "HIST1H2BD") c("
FCGR2B, "FCGR2C", "RGS1", "GZMA", "G0S2", "KIF5B", "IL1R1", "SLC7A7", "KYNU", "GZMB", "CYBB", "P
SLC24A3, "ELF3", "EYA2", "CLDN4", "SERPINB2", "POLD4", "PFKFB3", "CEACAM5", "CLIC3", "TPD52L1", "
IFITM1, "NAT1", "LAP3", "IFITM3", "ID1", "SCNN1A", "HGD", "STAT1", "GLUD1", "ARMCX5", "ZDHHC4",
E2F2, "SPP1") c("PLAT", "S100A11", "RARRES1", "MVP", "KRT7", "THOC7", "PIK3R3", "KLF10", "SEMA3C
25 c("NAPA", "MLF1", "G3BP1", "ITGA6", "SLC38A1", "POLR2I", "EIF5A", "LPL", "CDKN3", "VEGFC") c("H
CDC25B, "CA2", "SIGLEC15", "IER5", "RBM47", "KLK13", "ALDH1A3", "TNFRSF1A", "LRP8", "S100A7", "N

ZFAND5, "NDUFA4L2", "ENG", "TMEM147", "ANXA3", "FABP4", "TACSTD2", "EYA2", "SCFD1", "USP6NL"
TCN1, "SUB1", "BCAT1", "RGS16", "FZD6", "SDC1", "FMO2", "SPP1") c("ITM2B", "NAPA", "NUSAP1", "UG
MMP3, "HNRNPD", "PRPF4", "NQO2", "THBD", "CXCL5", "G0S2", "TPD52L1", "THOC7", "CA2", "UBE2L6"
HGD, "KRT14", "KIF14", "TUBB6", "LTF", "FOXC1", "IRS1", "CCDC86") c("ASPN", "ITM2C", "TCF7", "CTSK"
26 c("CIT", "SDF4", "KLK5", "LPL", "MELK", "PPP1R9A", "HIST1H2BD", "HIST1H2BK", "L1CAM", "MLF1", "
THBD, "TMEM158", "AQP3", "MLF1", "KMO", "TIAM1", "KYNU", "RBM47", "GLUD2", "PRNP", "CA2", "A
BLNK, "NETO2", "SCGB2A2") c("TACSTD2", "GCLC", "TMX1", "EIF4A3", "MARCKSL1", "ACSL1", "TCN1", "
ANXA4, "SLC16A1", "RBM47", "AREG", "S100A10", "CA2", "TSPAN31", "ANXA1", "LRP10", "MYO6", "GP
27 c("SLC38A1", "L1CAM", "SLC38A2", "JUP", "MGAT4B", "POLR2I", "EIF5A", "TNNT1", "HIST1H2BD", "K
CLCA4, "PDLIM1", "TPD52L1", "IGFBP2", "MMP10", "SPINT1", "S100A10", "TACSTD2", "ERBB3", "EPHA2
EPHA2, "SMPDL3A", "TNFRSF21", "ATOX1", "KRT17", "PAPSS1", "PLEK2", "SLC6A14", "GPRC5A", "FBL", "
28 c("KLK10", "SERPINB7") c("MMP3", "YME1L1", "ANXA3", "HIPK3", "S100A10") c("HSD17B4", "YME1L
29 c("TNNT1", "FGFR1OP", "GSTO1", "ITGA6", "NUTF2", "EIF5A", "KLK10", "NDUFA7", "ACP1", "PPP1R9
STOM, "GLUD2", "MYO1B", "CLEC7A", "KLK10", "MMP12", "G0S2", "TPPP3", "TLR4", "GFPT1", "ATP2C1
S100A11, "EHBP1", "ASRGL1", "CAPNS1", "HEBP2", "GPRC5A", "ZNF536", "CFH", "ITIH2", "MUC4", "CFI"
OAS2, "ISG15", "CAPN2", "AQP9", "ATP1B3", "KCNJ16", "CXCL5", "KRT5", "SDC1", "MARCKSL1", "MTHF
LCN2, "VNN1", "WSB2", "SH3BGRL3", "IL1B", "STOM", "FERMT1", "TSPAN1", "AZIN1", "MYO1B", "ST14"
C4BPA, "LYN", "NDRG1", "ARPC2", "HMOX2", "PSMC3", "OSBPL10", "HS3ST1", "ACTR2", "SMS") c("SLC2
ARFGAP3, "GZMB") c("PSPH", "RFC2", "CXCL13", "BANK1", "PAICS", "CPS1", "NUP93", "ZIC1", "SLC5A6",
SMS) c("CXADR", "TLE2", "TAGLN2", "FAM3C", "HOPX", "TM9SF2", "TFPI2", "PLAC8", "SSR3", "DNAJB6",
CTSE, "SCD", "GNLY", "PDHX") c("MEIS1", "IKZF1", "EXPH5", "MAP7", "GALNT1", "TM9SF2", "PPA1", "ZI
ARHGAP6, "RAC2", "CD83", "FKBP14") c("WWTR1", "MRPS16", "HOPX", "GPM6A", "CLDN10", "ITPR1", "
30 c("PPP1R9A", "HIST1H2BD", "EIF5A", "LPL", "SLC38A2", "CDKN3", "HIST1H2BH", "TNNT1", "VANGL1"
PSTPIP2, "POLR1C", "EPHA2", "ALOX5AP", "LCP1", "CEACAM1", "TPST2", "KLK6", "RFTN1", "TMPRSS4",
KRT16, "PSTPIP2", "EPHA2", "CLIP1", "TMBIM1", "GBP1", "CEACAM1", "RAB25", "ID3", "POLR1D", "PLS
31 c("EIF5A", "POLR2I", "CREM", "PPP1R9A") c("TPM4", "FOSB", "CCL20", "S100A10", "MET") c("IL7R", "
32 c("NPR3", "LPL", "L1CAM", "TNNT1", "NEK7", "KLK10", "EIF5A", "SDF4") c("SHB", "TOP1", "KCNN4", "
HK2, "CCL20", "KRT14", "LTF", "TOMM34", "PLAC8", "JAKMIP2", "SLC43A3", "FCGR2C", "CD46", "BAG1"
LAPTM4B, "DUOX2", "SLPI", "KRT14", "GABRP", "PLAC8", "SLC35A2", "ADAR", "TMED9", "TAF9B", "FAM
33 c("VANGL1", "POLR2I", "SET", "TNNT1") c("SPRR3", "MREG", "UPP1", "TPM4", "CFI", "ANXA3", "PDLI
34 c("CARM1", "EIF5A", "NUDT15", "LPL", "SERPINB7") c("RRAS2", "SEC14L1", "S100A10", "MET", "TRIM
35 c("PROCR", "GSTO1", "NUTF2", "MLF1", "BOP1", "NCAPH", "SNRPA1", "F12", "TNNT1", "MELK", "PPP
MELK, "KMO", "CD3G", "PITPNC1", "ZYX", "S100A2", "CYBB", "ELK3", "ADRB2", "CDK6", "AQP3") c("TRIM
MMP12, "NAT1", "MMP3", "NDRG1", "C1GALT1", "IFI30", "CFTR", "FCGBP", "CFB", "MT1E", "WDR91", "

'V19" "V20" "V21" "V22" "V23" "V24" "V25" "V26" "V27" "V28" "V29" "V30" "V31" "V32" "V33" "V34" "'
) c("HLA-DQA1", "EHHADH", "PLA2G7", "LGALS2", "SLC39A14", "STOM", "ELMO1", "LYZ", "CEACAM1", "
KCTD5", "KLK10", "NUTF2", "OLA1", "POLR2I", "RPL26L1", "TNNT1", "YARS") c("ANXA3", "BMI1", "CCL2(
RSF1B", "TPST2", "TRAPPC3", "TUBA1A", "TUBB6", "TWIST1", "VNN1", "VPS37B", "ZYX") c("ADO", "BMI
GFA", "TPD52L2", "TSPAN1", "TUBA1A", "VNN1", "WSB2") c("AREG", "CYP3A5", "IL18", "IL20RA", "MAP3
A6", "HIST1H2BG", "LPL") c("PDLIM5", "YME1L1", "USP7", "RYBP", "STX3", "ANXA3", "SIK1", "UGCG", "S
LY96", "S100A7") c("PDLIM5", "YME1L1", "PPWD1", "USP7", "INHBB", "RYBP", "LYPLA1", "SPATS2", "RH
PHLDA1", "RAB11FIP1", "IL1RN", "AREG", "DYNLT1", "CLIC3", "GPRC5A") c("SAMM50", "MAT2B", "IFITM
SBPL10", "ANXA2", "LRP10", "RNF19B", "ZC3H12A", "GOLPH3L", "QPCT", "CDKN1A", "TSPAN1", "DAPK1
B8", "CCNL1", "FOSL1", "FOSB", "SIK1", "HBEGF", "EGR2", "OBSL1", "MMP3", "PCDH9", "TPM4", "EPHB2
"KCTD5", "KLF10", "SLC10A3", "LCP1", "FCER1G", "RGS1", "CYP1A1", "KLK10", "CD14", "CD74", "RALGDS
TD2", "KCTD5", "P4HB", "POLE3", "RAB25", "HLA-A", "HLA-B", "TREM1") c("FXYD3", "GSTM3", "HSPB8",
c("CCDC81", "FCER1G", "AQP3", "MMP9", "PLAC8", "EAF2", "CXCL6", "SAT1", "LYZ", "CDK6", "SOD2", "K
, "MMP10") c("ITGA3", "CD46", "SRGN", "EDEM1", "ITGB2", "CSF1R", "CYBB", "LAPTM5", "KRT17", "MS
PRSS4", "LIMA1", "LPXN", "KLK3", "MID1IP1", "TRAPPC3", "ME2", "CDKN1A") c("DNMBP", "SLC19A2", "
LYRM2") c("MREG", "CD55", "G3BP2", "B4GALT5", "YME1L1", "LAMC2", "SIK1", "EGR3", "SOX9", "EGR2
LOX5AP", "IER5") c("CD55", "SEMA6A", "YME1L1", "RPS6KA3", "MRPS30", "KLHL21", "ADO", "DUSP3",
, "MCTS1", "H2AFY", "EMX2", "UCHL3", "HOPX", "FKBP1A", "TXN", "NME1", "HELLS", "STT3A", "CYB5R2"
T2", "PHLDA1", "DAPK1", "AKR1A1", "KCNJ16", "LYN", "HEG1", "IL1A", "S100A11", "G0S2", "ID1", "SLC2
RAD", "CYP51A1", "KLK10", "HIST1H2BH", "LPL", "SERPINB7") c("SPRR3", "UPP1", "CSTB", "SOX9", "S10(
CSF1R", "HLA-DQA1", "ASPN", "ARHGDIB", "ADAM8", "HSD17B2", "ATP6V1G1", "CD74", "TMEM165", "
L2", "KCNK3", "GRIA2", "PTPN12", "POGK", "PIM1", "TRIM16", "EPHB2", "CAB39", "OBSL1", "KLHL21", "
DEC", "ZNF83", "SFRP1", "TOB2", "PTX3", "RIF1") c("HOPX", "DHRS3", "H1F0", "ERBB3", "GDE1", "GTF2I
"KRT19", "TPD52L1", "CLDN4", "MT1E", "COBLL1", "CTSH", "BAD", "CYP4F3", "HMGCL", "ANXA1", "CTN
"STAT1", "ISG15", "PRPF4", "CDC37L1", "HLA-DMB", "FAM3C", "MMP1", "IFITM1", "XPNPEP1", "MYOF",
LPH3L", "RAB38", "BCL2A1", "CNDP2", "FCGBP", "MUC1", "SLC6A14", "BST2", "IER5", "LRP10", "CLDN7"
G", "SPP1", "CX3CL1", "IDE", "STAT1", "SLPI", "TAPBP", "S100A2", "KRT14", "HDDC2", "LAMC2", "THBD",
LHL21", "RNF103", "ANXA3", "FUT8", "FOXA1", "ATF3", "ADO", "RRAS2", "MYC", "PCDH9", "PELI1", "NRI
MAST4", "BMP2", "HLA-E") c("GRHL2", "AMD1", "UAP1L1", "SLC5A6", "DLD", "RXRA", "HAT1", "AACS", "
SKAP2", "ZNF217", "CLDN8", "SEMA4D", "GPA33", "CASP10", "FAR2", "CCNL2", "ZMAT3", "TOB1", "OPH
T7", "BGN", "PPIC", "STXBP2", "GPX4", "FGFR3", "SDC4", "CTSK", "CD46", "PTPN12", "LTBR", "APOLD1", "
C42", "PDGFD", "ERBB2", "SCD5", "DACH1", "IRX5", "SH3GLB2", "MUC1", "ALDH3B2", "KYNU", "XIST", "
NFSF10", "NCOA4", "CCNG1", "KIAA0232", "ZFAND6", "PPP2R5E", "TRAF5", "MAP3K8", "KCNJ2", "SAT1"
BL1X", "CPT1A", "CYP1B1", "CLU", "TACSTD2", "S100A10", "EFHD1", "EPS8L3", "NDFIP1", "TMPRSS2", "S
", "NDFIP1", "PPP2R5E", "FCGBP", "MUC1", "FBXO11", "MBNL2", "TCN1", "SLC25A13", "FOXA1", "TGFA"
"SPP1", "SLPI", "KCNJ15", "REL", "USP6NL", "MCM6", "CLDN10", "PPFIBP2", "LGR4", "MAP7", "DNMT3A
CTNND1", "GTPBP8", "SCD", "MYL9", "TRAK1", "CDH1", "STK38") c("PEX11A", "DSP", "COL9A3", "MEIS1
"PDCD4", "GPM6B", "FKBP11", "PTGER3", "LGR4", "PRKCH") c("LRRC1", "HOPX", "SLC5A6", "PPIC", "PR
16", "S100A11", "ERBB2", "PRKAR2B", "MUC16", "TNFSF10", "SLC12A2", "MVP", "PRRG1", "ZFAND6", "
"PODXL", "OLFML2A", "TIMP1", "PRODH", "MAST4") c("HOPX", "GLB1", "BAZ2B", "RTN1", "PTPN12", "N
N", "SIRT3", "EFHD1", "PLCB3", "STXBP6", "SNAP25", "MVP", "DDB2", "HUWE1", "TANC2", "ALDH3B2",
PX2", "TMPRSS2", "TNFSF10", "STK38L", "TMPRSS4", "MVP", "PRRG1", "ITGAV", "MUC1", "ALDH3B2", "
PA", "TNNT1", "ALG5", "LPL", "L1CAM", "MGAT4B", "F12") c("SOX9", "NRIP1", "PSPC1", "MREG", "YME1
TP6V1G1", "ITGA4", "RFTN1", "HLA-DRA", "MYC", "EAF2", "PRSS3", "PWP1", "TRPV2", "PLS1", "HLA-DQ
", "TJP3", "GABRP", "ACTN1", "CDKN1A", "LAMC2", "KRT5", "PHLDA1", "S100A2", "SLC25A13", "CX3CL1"
GF2", "KRT7", "RNASE6", "APPL2", "ADAM28", "FABP4", "FILIP1L", "IL17RB", "SCGB1D2", "SDC4", "LAM
"ITGA6", "POLR2I", "NPR3", "HIST1H2BH", "EIF5A") c("SLCO1B3", "ANXA3", "SPAG1", "PNN", "FOSL1", "

"DAXX", "RRS1", "AQP3", "DNAJB6", "CCR7", "BTN3A2", "S100P", "ADRB2", "PRMT1", "MLC1", "PTGS2",
A1", "TPD52L1", "S100P", "HLA-B", "IFITM1", "PLEK2", "SERPINB1", "NFKBIE", "CXCL5", "ACTN1", "GPRC
MO2", "FBLN5", "GPM6A", "FGFR3", "PLEKHJ1", "NEK7", "FEZ2", "ASPN", "GPX4", "VPS37B", "LAMA5",
", "AHCYL1", "KLK6", "NAMPT", "HS3ST1", "BPGM", "ALOX5AP", "S100A2", "CCR7", "PLAT", "KRT17", "C
", "OLA1", "XPOT", "GSTO1", "HIST1H2BH", "JUP", "HIST1H2BK") c("GPA33", "ZNF24", "TRIM16", "EPHB
U") c("ZNF24", "TRIM16", "EPHB2", "USP7", "GRIA2", "CACNA2D2", "ADO", "TERF2", "AGER", "JUND", "
RAP1GAP", "SLC7A8", "PHGDH", "IDH1", "ESR1", "FXYD3", "CACNA2D2", "CRABP2", "SCNN1A", "ZNF586"
F12") c("EGR2", "CFI", "SYNPO", "FOSB", "TRIM16", "KCNN4", "BMI1", "PIM1", "EPHB2", "YME1L1", "CA
", "MCL1", "KIFAP3", "BMI1", "NFKBIA", "PIM1", "EPHB2", "ZSCAN18", "YME1L1", "RRAS2", "APOD", "IL
, "CLCA2", "SLC6A14", "MICALL1", "KRT17", "IL1RN", "KRT5", "KRT14") c("JAG1", "GMPR2", "MTUS1", "
, "TCP1", "HAT1", "TNNT1") c("EGR3", "EGR2", "POLR2H", "PELI1", "EPHB2", "SOX9", "STX3", "ARL14", "
"MGAM", "PLS1") c("ZFP36L2", "NEDD4L", "GRIA2", "FAM131A", "INHBB", "EPHB2", "CCL8", "MRPS30",
LC6A14", "GSTP1", "PDZK1IP1", "DNAJC1", "PTP4A1", "ANXA4", "DHCR24", "RAPGEF5", "DAPK1", "SLPI"
MED13", "CCL20", "SLC20A1", "DMTF1", "HSPB8", "NRIP1", "DFFA", "SHB", "MREG", "GPA33", "MMP3", "
IA2", "DMTF1", "RAB21", "ZSCAN18", "IL17RB", "KLHL21", "LYPLA1", "SHC1", "SEMA6A", "MMP3", "EPH
T1", "HLA-F", "KCNJ16", "STK38", "MMP3", "PRPF4", "ID3", "CLCA4", "PSMB8") c("SLC7A8", "CAST", "SC
", "TTC9", "WTAP", "TMPRSS4", "S100A2", "ISG20", "MS4A6A", "SMPDL3A", "RHOBTB3", "TCN1", "ALOX
KCNQ2", "NUTF2", "CDKN3", "BOP1", "CIT", "POLR2I", "KBTBD11", "SET", "CBX3", "OLA1") c("TRIM16",
S2", "PITPNC1", "HS3ST1", "FSCN1", "RBM47", "ID1", "TRAPPC3", "CCR7", "ID3", "PLAT") c("EIF1", "TRIM
MT1E", "FDPS", "CEACAM6", "IDE", "DDOST", "CD46", "ANXA2", "TMBIM1", "CEACAM1", "FERMT1", "N
GNL2", "BRD2", "CCL5", "PEG3") c("PRNP", "HMGB3", "TUBB6", "HSPH1", "LMNB2", "EPHA2", "KLK10",
FI", "B4GALT5", "KCNK3") c("IL7R", "CYP1B1") c("MAST4", "FARSA", "CCR7", "LRP8", "SPHK1", "MLC1", "
MP6") c("MTHFD2L", "TOP1", "HSPB8", "CCL20", "ARG2", "NRIP1") c("CD8A", "SOAT1", "PEG3", "CCL5",
M2", "LY96", "NR4A2", "EPHA2", "G0S2", "S100A7", "LYZ", "KRT17", "ALDH1A3", "S100P", "ID1", "ALOX
L2", "ADAM17", "NEK7", "HIST1H2BG", "CHMP6", "FZD10", "MAEA", "LPL") c("OBSL1", "SOX9", "IL1A", "
, "CYP1A1", "RRAGC", "TLR4", "TRAPPC3", "KYNU", "S100A2", "EFR3A", "IRF6", "FGFR3", "THBD", "ISG20
, "MPPED2", "MAOA", "TPP1", "SQSTM1", "EAPP", "S100A10", "ANXA3", "ECHDC2", "CANT1", "TAX1BP
STK1", "INPP4B", "CDH3", "MMP12", "ALOX5", "MMP7", "DPAGT1", "ADA", "BID", "STEAP1", "COMMD
PIC", "GPATCH2", "ECHDC1", "PTPRK", "PPT1", "KRT5", "CXCL5", "WARS", "NFKBIE", "MMP1", "ISG15")
AC1", "SLC9A3R1", "ELF3", "CDKN1A", "CLCA4", "PLAC8", "MYO6", "CA2", "CMTM6", "PRDX3", "FOXC1
"LRP10", "PPIC", "PMM2", "STAU2", "UBE2E3", "SH3BGRL3", "MICALL1", "CASP4", "EHD4", "C4BPA", "C
", "MMP2", "SLC2A6", "C7", "RIPK2", "FAM149A", "MAN1C1", "ZMYND8", "HSPA8", "IRF4", "ITM2C", "G
TCN1", "ELF3", "PPP2R5E", "PLAC8", "PRDX3", "MUC1", "RAB9A", "MBNL2", "FOXA1", "HOPX", "TTC9", "
SAT1", "GCH1", "RARRES1", "ST7", "CLDN4", "TCN1", "PRRG1", "MVP", "ELF3", "CDC42SE1", "PLAC8", "F
KCNJ15", "CPPED1", "POLI", "IDS", "MUC16", "MNAT1", "RUNX3", "MRPS16", "E2F2", "MAN2B2", "CERK
SPG11") c("HOOK2", "ZIC1", "CDS1", "SPRY2", "RUNX1", "TNFSF10", "MEIS1", "CYP2J2", "DAAM1", "EDF
2", "ITGB1BP1", "BID", "PDCD6", "TTR", "LACTB2", "PAPSS1", "LRRC1", "SLC5A6", "PPP2R5E", "TAC1", "
c("ALDH3B2", "ERBB2", "TNFSF10", "SLC12A2", "S100A11", "PRRG1", "MVP", "PLAC8", "PIK3R3", "FOXC
") c("DUSP22", "STK38L", "AUTS2", "SET", "BCAM", "CLDN10", "LAMA5", "GCH1", "SLC12A2", "FMR1", "
, "RARRES1", "VPS13B", "MVP", "FGFR1", "SULT1A1", "LY6E", "LIMK2", "INHBB", "JAM2", "SELE", "SERP
KCNN4", "LAMC2", "S100A10", "OBSL1", "FOSB", "ANXA3", "SOX9", "FOSL1", "SIK1", "EGR3", "PRKCB", "
FN1", "CLDN4", "MMP9", "DNAJB6", "MGAM", "CEACAM1", "S100A7", "IL1RN", "RBM47", "CXCL2") c("I
ERBB3", "TPP1", "MAL", "IL1RN") c("HOMER1", "SH3BP4", "GPR87", "MMP7", "INPP4B", "KRT5", "CAP
"DDX60", "IFI44", "KCNJ16", "G0S2", "PAK1IP1", "FBXO34", "UTP14A", "RAB38", "MSRB2", "ELF3", "MA
", "PLAC8", "SLC2A1", "BCL3", "KLK6", "PRSS21", "TFPI2", "KYNU", "SERPINB2", "DNAJB6", "GPX2", "S10
SPA4L", "EGR3", "SPRR3", "HSPB8", "S100A10", "SMOX", "ARHGAP5", "EPHB2", "ARG2", "CD55", "TRIM
AMPT", "MID1IP1", "EMP2", "LY96", "KRT14", "KIF14", "STAM", "TUBB6", "TJP2", "C1QB", "UGCG", "LA

', "FOXC1", "KRT19", "PSCA", "FZD6", "MBOAT7", "GABRP") c("NES", "SLC25A13", "SLC25A28", "MCM3",
CHL3", "PI3", "NRBF2", "AKR1B10", "BTG3", "SPRR2B", "SPRR1B", "HOPX", "GSTP1", "CEACAM6", "HNRN
", "IER5", "AQP9", "NETO2", "RBM47", "ALDH1A3", "PRDX1", "AREG", "S100A7", "ANXA1", "SLC12A8", "
', "GFPT2", "GEM", "FGFR3", "PRPS2", "FBXO11", "KYNU", "SDC4", "SLC2A1", "SERPINB2", "SPRED2", "TH
'EIF5A", "SPDEF", "INTS6", "TNNT1", "SLC38A1") c("EGR2", "STX3", "MED13", "TRIM44", "MYC", "YME1L
TP2A2", "SLC43A3", "TUBB6", "KCNS3", "LIMA1", "PLAT", "DAXX") c("QKI", "SMAD7", "MED13", "ADO",
HLA-F", "PTK2", "KRT5", "QPCT", "MMP7", "FZD6", "ARCN1", "PLAU", "MYO1B", "MLXIP", "ALOX5", "STA
RC5A", "GBP1", "IFIH1", "DDX60", "STIP1", "BLNK", "PLAT", "NETO2") c("SIRT1", "TKTL1", "STX3", "GSTM
LK10", "HIST1H2BK") c("PRKCB", "SHB", "SYNPO", "OBSL1", "ANXA3", "CD55", "SPRR3", "S100A10", "SIK
2", "ATP9A", "GPRC5A", "DUSP2", "HES1", "CAPNS1", "SGPL1", "CLIC3", "CEACAM5") c("ANXA4", "TP53T
"YIF1A", "HES1", "DHRS7", "CAPNS1", "SGPL1", "SLC25A6", "AIM2", "KRT16", "INPP4B", "IFITM1", "TNFS
.1", "RAB11FIP2", "PEG3", "MMP10", "ASPH", "TARP") c("SLC43A3", "TRAPPC3", "LRRC8B", "BSG", "KLK1
A", "XPOT", "SERPINB7", "NAPA", "NPR3", "JUP", "POLR2I", "SIGMAR1") c("HSPB8", "CD55", "CA8", "SIK
", "GARS", "ME2", "CARS2", "IRF6", "APOBEC3A", "HMGA1", "CXCL6", "PRMT1", "CA2", "EFHD2", "MINK
', "NDRG1", "VCAM1", "OSBPL10", "GNLY", "LST1", "ABHD2", "CRAT", "CLIC3", "EAPP") c("TLE2", "PLEK",
D1", "MLXIP", "CTSC", "TCN1", "HEBP1", "TSPAN6", "UBE3C", "EPX", "ZNF586", "NME1", "ACSL1", "ARPC
', "PLS3", "DHCR24", "MMP12", "UGP2", "SERPINB1", "XPNPEP1", "LTA4H", "TGFA", "G0S2", "FAM120A"
25A14", "HSPB8", "CHCHD7", "HIST1H1C", "SCNN1A", "MAP3K1", "CSNK2A2", "EPHX2", "CCDC6", "CDK7
, "CETN2", "IGF2BP3", "NDUFB2", "EHF", "MN1", "RBBP8", "ACO1", "UQCRQ", "SNRPF", "PHGDH", "NME
, "STOM", "TEK", "PRKAR2B", "SWAP70", "PTPN13", "S100A7", "SERPINB2", "S100A11", "FBXO11", "MA
C1", "MAL", "ACADM", "LSM5", "SSBP1", "FAIM", "TP53TG1", "LTA4H", "NDUFA3", "CDS1", "EDF1", "PEF
"AKAP9", "TM9SF2", "PI3", "CSAD", "LSM6", "LY75", "LIMS1", "MDFIC", "HIP1", "DUSP22", "NDUFA3", "V
', "RNF6", "ITGA6", "KLK10", "BOP1", "HIST1H2BG") c("TRIM44", "TPM4", "KCNN4", "TRIM16", "TNFRSF
"DNAJB6", "ID3", "AQP3", "MMP12", "PTP4A1", "KLK10", "ID1", "CDC42SE1", "KLK3", "HLA-DRA", "ALDH
3", "MMP12", "PTP4A1", "ANXA1", "ID1", "CLPTM1", "LAPTM4B", "ALDH1A3", "UTP14A", "CXCL5", "UBE
"MEA1", "ADD3", "PEG3", "MID1") c("LYPD1", "GLUD2", "IL1RN", "PEA15", "BTN3A2", "PCSK1", "S100A2
'ALDOA", "SLC20A1", "PELI1", "CFI", "EGR2", "PCDH9", "EGR3", "UPP1", "ETNK1", "NRIP1", "CCL20", "LR
', "ATP6V1G1", "SQLE", "TUBB3", "DDX3X", "CASP7", "BSG", "ME2", "FCGR2B", "SLC6A14", "ASPN") c("T
1120A", "CD46", "ADIPOR1", "RAPGEF2", "ATP6V1G1", "CLTA", "GSTP1", "UGP2", "RBPMS", "CTSC", "CA
M5", "HSPB8", "S100A10") c("DAPK1", "SOAT1", "CYP1B1", "ZNF32", "GRB14", "PEG3") c("OLFML2B", "
/16", "YME1L1", "PCDH9", "CCL20", "HNRNPA2B1", "ASRGL1", "HSPB8") c("KLC1", "SOAT1", "SLC5A6", "
P1R9A", "HIST1H2BK", "HIST1H2BD", "NPR3", "PA2G4", "ITGA6", "EIF5A") c("TRIM16", "GPAA1", "BMI1"
M16", "BMI1", "CYP4F11", "FGL1", "NRIP3", "GRIA2", "QKI", "NEDD4L", "CD1D", "MMP3", "EPHB2", "KLH
'PPP1R9A", "LAP3", "PITPNC1", "FAM120A", "EHD3", "RAB25", "PRPF4", "UBA6", "RPN2", "TARS", "S100

V35" "V36" "V37" "V38" "V39" "V40" "V41" "V42" "V43" "V44" "V45" "V46" "V47" "V48" "V49" "V50" "\
MMP9", "ASPN", "SAT1", "PBLD", "USP18", "EAF2", "MYC", "NR4A2", "ISG20", "PTP4A1", "LTF", "SMPDL
)", "EGR2", "EGR3", "FOSL1", "HBEGF", "HIPK3", "HSPA4L", "LAMC2", "LRRC8D", "MET", "PDLIM5", "PPP
1", "DUSP3", "DVL1", "EREG", "FGL1", "FSCN1", "INHBB", "KLHL21", "PDLIM5", "PPP3CA", "PRNP", "PTPI
3K1", "MT1H", "OSTF1", "RAP1GAP", "S100A10", "SCNN1A") c("ADO", "ANXA3", "IFRD1", "KLHL21", "PDI
HB", "SPATS2", "CFI", "RHBDF1", "EZR", "SOX9", "ETNK1", "PPP3CA", "GPA33", "TRAF4") c("MMP10", "P
HBDF1", "APOD", "IRF6", "PPP3CA", "NFKBIA", "SEMA6A") c("PLSCR4", "SAMM50", "P2RX4", "GRB14", "
/11", "ANXA4", "MED24", "G3BP1", "MMP7", "STAT1", "TMED5", "PSMD10", "ADAMDEC1", "UBE2N", "IS
.", "LCN2", "TBC1D8", "CX3CR1", "HSD17B1", "NETO2", "EPHA2", "MPZL2", "PHLDA1", "EIF4EBP1", "IL1R
2", "S100A10", "FRMD4B") c("MAGEF1", "GPR65", "IL7R", "SLC5A6", "CCL5", "PRF1", "LCP1") c("TNFRSF1
5", "FPR1") c("GRIA2", "IER3", "NR4A1", "KLHL21", "ADO", "OBSL1", "CCL8", "MMP3", "NRIP3", "IL17RB"
, "TM4SF1", "MT1H", "VAMP1", "JAG1", "S100A13", "ECM1", "TKTL1", "AREG", "MKNK2", "S100A10", "S
YNU", "TMEM158", "F2RL1", "ISG20", "TCN1") GRIA2 c("CAMK2N1", "GCH1", "SERINC5", "MOCS2", "PSI
4A6A", "CLEC7A", "FPR1", "C1QB", "CXCL1", "GZMA", "ARHGDIB", "BPGM", "CD14", "ALOX5AP", "LTF", "
CSTB", "MED13", "OBSL1", "ADO") c("DAAM2", "HIGD2A", "IMPDH2", "PRTN3") c("PRSS21", "DNMBP",
2", "HIF1A", "UMOD", "LRRFIP1", "CCL20", "KCNK3", "PAQR6", "KCNN4", "TNFRSF12A", "RRAS2", "FOSL1
"TERF2", "POGK", "DUSP7", "KCNK3", "NFKBIA", "CASK", "IL17RB", "TNFRSF12A", "LYPLA1", "RRAS2", "P
', "GINS2", "SLC11A2", "AKR1B10", "TRIM22", "GSTP1") c("YIF1A", "HCCS", "ATR", "FERMT1", "FDPS", "S
A9", "ARPC3", "PSCA", "MARCKS", "ADA", "PLS3", "KRT5", "S100A2", "GSTP1", "POLR1D", "MICALL1", "D
0A10", "TRAF4", "GPA33", "KCNK3", "PIM1", "TRIM16", "CCL20", "EPHB2", "TSPYL1", "OBSL1", "NRIP1",
IFI16", "DDX21", "LDLR", "RBM47", "PLAT", "TPPP3", "TRAF3IP2", "VNN1", "KLK10", "WTAP", "CXCL2", "I
DUSP3", "DDX17", "SEMA6A", "NFKBIA", "LYPLA1", "PPP3CA", "SMAD7", "PXDN", "GCNT1", "CHSY1", "A
RD1", "NRCAM", "FHIT", "RAB5B", "CEACAM6", "CLDN7", "SAT1", "TSPAN6", "PRR15L", "NKX3-1", "TMC
NA1", "TRIOBP", "SNAP23", "FOXA1", "HOXC6", "MAOA", "FABP4", "ADM", "ATF3", "NPAS2", "SH3YL1",
, "HCP5", "STK38", "SLC25A6", "BTN3A3", "HLA-F", "HLA-E", "PSMB8") c("HOPX", "AACS", "ELOVL6", "ER
', "GRB7", "SAT1", "ANXA2", "ZC3H12A", "SDC1", "S100P", "KRT5", "AQP9", "SLC25A13", "TPD52L1", "N/
"VAMP8", "GTPBP8", "LAP3", "UCHL3", "RGS10", "PRPF4", "ACSL5", "REL", "CMTM6", "LDHA", "IFITM1'
P3") c("HDAC1", "BPGM", "YWHAZ", "ID1", "HIST1H2AE", "TACSTD2", "GNAQ", "LRP4", "PEA15", "CCL2(
STX4", "TSPYL1", "DAZAP2", "TMEM106C", "BANK1", "HNRNPH2", "HELLS", "ZIC1", "EHF", "MNAT1", "CE
N1", "HECA", "CITED2", "IGBP1", "CALML5", "GOLGB1", "FBXO11", "OGT", "TUFT1", "RAB11FIP1", "ZKSC
"STK38L", "PRRG1", "FAM13A", "ITGAV", "NRIP1", "LITAF", "IGF1R", "TPBG", "APLP2", "KYNU", "CYFIP2",
SCUBE2", "LY6D", "PDGFRL", "AKR1B10", "SLC25A13", "MLLT11", "PAX5", "ARPP19", "LRRC15", "BICC1",
', "CRABP2", "ZC3H12A", "COBLL1", "GSTT1", "PERP", "SRPK2", "ARMCX6", "HOXC6", "FCER1A", "PRKCZ'
TK38L", "MVP", "FBN2", "PRRG1", "CST3", "LTBP3", "FGB", "DAPK1", "IFIT1", "TPBG", "TANC2", "FMO5"
', "PLAT", "NAMPT", "TIMM9", "PDZD8", "AREG", "SPP1", "SLPI", "GALC", "USP6NL", "MCM6", "RAB1A",
A") c("HOPX", "RARRES1", "KRT7", "PIK3R3", "S100A11", "PRKAR2B", "GPX2", "MVP", "STOM", "CDC42SE
", "ERBB3", "TNFSF10", "TMPRSS4", "RAP2C", "TMBIM1", "RIN2", "TP53TG1", "PPP2R5E", "PIK3IP1", "SL
R16", "SPINT2", "HIST1H2AE", "PPFIA1", "RNF43", "NCOA4", "RAB5B", "PPP2R5E", "HIST1H2BD", "ZC3H
FGB", "GPR137B", "MUC1", "ALDH3B2", "KYNU", "CDKN1B", "CTSA", "FBXO11", "MBNL2", "TBC1D8", "N
IDFIP1", "STK38L", "SLC12A2", "DDB2", "MRPS16", "MBNL2", "ITPR1", "PDZD8", "AUTS2", "TXNDC9", "TI
"IFRD2", "IER5", "PDGFRL", "LRRC15", "PLAT", "SEC14L1", "CDH11", "WIPF2", "RRAS2", "SPP1", "SNAI2",
KYNU", "LPCAT1", "MAP3K8", "SAT1", "S100P", "HMOX1", "CLDN4", "TPK1", "MMD", "TGFA", "TMEM16
L1", "PYROXD1", "CFI", "OBSL1", "EGR3", "EGR2", "KCNN4", "LAMC2", "ARG2", "SLC20A1", "CCL20", "CD
B1", "KLF10", "TACSTD2", "IL1R1") c("IER3", "PRNP", "MRPS30", "CCL8", "KLHL21", "PSPC1", "POGK", "YI
", "KRT17", "PLAT", "SCNN1A", "DDIT4", "SP110", "ATP6V1G1", "DHRS7", "IRF9", "THOC7", "POLD4", "C\
A5", "THOC7", "RTP4", "STK38L", "IGF1R") c("MS4A4A", "HK1", "SCD", "RARRES1", "IFRD2", "ME2", "PIK
PPP3CA", "S100A10", "LAMC2", "PCDH9", "HSPB8", "EGR2", "LRRC8D", "TSC22D2", "PDLIM5", "EGR3", "

, "CCL20", "MPHOSPH6", "CNN2", "CXCL1", "ACTN1", "RAC2", "ALOX5", "CXCL2", "PLAC8") c("INHBB", "N
5A", "MARCKS", "NQO2", "PLAC8", "PSMB8") c("DST", "CYP3A5", "TIMM13", "MT1H", "CLDN8", "HIST1H
"GEM", "GABARAPL1", "KRT7", "SLC7A1", "SERPINB2", "DNAJB6", "EHD1", "ADRB2", "VASP", "ITM2C", "I
DK2", "LTF", "TACSTD2", "CEACAM1") c("APOD", "TRIM16") c("ASPH", "VPS45", "TPSB2", "TPSAB1", "AL
2", "USP7", "PLEKHB2", "SPRR3", "PRKCB", "ASRGL1", "PCDH9", "SOX9", "HSPB8", "NRIP1", "CFI", "SIK1",
ABCA8", "DUSP3", "QKI", "LYPLA1", "OBSL1") c("SLC3A2", "ADAM28", "NUP85", "ADRB2", "SUSD4", "NA
", "MARCO", "SH3BP4", "SH3YL1", "HIST1H1C", "NTM", "HSPB8", "PDZD2", "FGGY", "ECM1") c("ZNF24",
8", "EGR3", "RRAS2", "RHBDF1", "TRIM44", "OBSL1", "UGCG", "TSC22D2", "ANXA3") c("RAC2", "UGP2",
17RB", "RHBDF1", "OBSL1", "TSC22D2") c("SKIV2L", "BMP4", "DAAM2", "ARPC4", "ADAM28", "CAMK2N
HIST1H1C", "TM4SF1", "PPL", "AREG", "SH3YL1", "NBR1", "NAV2", "ZNF586", "MT1H", "ADAM12", "FAM
PPP3CA", "TSPYL1", "GPA33", "LAMC2", "S100A10", "BMP2K", "ETNK1", "TRA2A", "VRK2", "CD55", "ANX
"ZSCAN18", "PPP3CA", "APOD", "CD55", "CAB39", "OBSL1", "KLHL21") c("CRK", "HIST1H2BD", "NAV3", "
, "SLC16A1", "CHMP6", "ACSL5", "MED24", "ENO1", "PSMB8", "NAT1", "IL1RN", "IDO1", "CDK6", "LYN",
"EPHB2") c("CEP170", "BRD2", "AMPH", "RCN1", "HMGCS1", "CYP1B1", "TARP", "PEG3", "CD8A", "UGP2
B2") c("SNCG", "XRCC5", "NUP85", "ILVBL", "AIFM1", "PRKX", "SEMA4C", "HIST1H2BD", "TNFAIP2", "ASI
NN1A", "CSNK1G3", "CRABP2", "PGAP1", "SH3BP4", "ZNF586", "GMPR2", "JAG1", "BMP7", "VAMP1", "S
5", "TNFRSF21", "S100P", "LTF", "LYZ", "PLAC8") c("PXDN", "CD55", "EPHB2", "FOXO1") c("BAALC", "HM
"SPRR3", "SPINK5", "UPP1", "CD55", "TIPARP", "S100A10", "ANXA3", "GPAA1", "DMTF1", "PPP3CA", "PC
16", "ABCA8", "MCL1", "APOD", "CD55", "GRIA2", "KIFAP3", "RBPJ", "DVL3", "FGL1", "KCNQ2", "DMTF1
AT1", "VAMP8", "PITPNC1", "TFRC", "GSTK1", "TSPAN1", "HS3ST1", "APOLD1", "FSCN1", "RBM47", "ID1"
"CA2", "MPHOSPH6", "KLK6", "PLAC8", "FGFR3", "CLIC1", "ACTN1", "KIF14", "DAXX") c("PRNP", "GNL2",
BSG", "PLIN2", "SAT1", "UGCG", "CDC25B", "RGS1", "GRWD1", "TOMM34", "ID1", "TMPRSS4", "CD14",
"GRB14", "SKAP1", "HLA-DOB", "APIP") c("KL", "KRT24", "EHHADH", "S100P", "MGAM", "IL1RN", "GZM
5", "CA2", "NUSAP1", "CDKN1A", "KYNU", "ASPN", "S100A2", "PLAT", "FOSL1", "TCN1", "AQP3", "LYPD1
PIM1", "CSTB", "BMI1", "RHBDF1", "S100A10", "ANXA3", "TRIM16", "PELI1", "CFI", "CD55", "SPRR3", "P
", "LIPG", "ANXA2", "ID3", "ID1", "IL1RN", "NEK7", "XPO7", "ATP1A1", "PTGS2", "WTAP", "F2RL1", "C1Q
", "FOXA1", "TTC9", "OSBPL10", "RAB11FIP1", "SCNN1A", "USP6NL", "ZFAND5", "BCL6", "CLDN7", "EPS
3", "DDIT4", "MARCKSL1", "SDC1", "ANXA4", "SERPINB1", "LACTB2", "HOMER1", "MARCKS", "TP53TG1",
c("CYB5R2", "CXADR", "ENAH", "TRIM22", "EMX2", "PBLD", "EPHA4", "FAM129A", "COMMD10", "ERBB
", "RBM47", "MUC1", "VPS35", "KCNJ16", "GLUD1", "SLC35A2", "NLRP2", "KLK5", "UBE2I", "IRF9", "CD4
HMP6", "CNDP2", "KRT5", "SLPI", "GFOD1", "CXCL5", "WARS", "LAPTM4B", "TNFRSF21", "ZNF593", "AR
PNMB", "MYOF", "GBP2", "PKN2", "RBL2", "BTN3A3", "MSRA", "ITPKB", "NDN", "CDO1", "CXCL9", "CHS
"NAMPT", "USP6NL", "CLCA2", "PHF11", "RNASE6", "LY75", "XPO7", "SH3BGRL3", "TIMM9", "NDUFA3",
IK3R3", "CMTM6", "PRDX3", "FOXC1", "MUC1", "RAB9A", "VPS35", "MBNL2", "HOPX", "TTC9", "NAMPT
", "LSM5", "NDUFA3", "SLPI", "PRKAR2B", "FZD10", "GALNT1", "MGST3", "ATP8A1", "TPBG") c("CXADR"
1", "MAL", "VAMP8", "TMBIM1", "TP53TG1", "RIN2", "ERBB3", "TCN1", "RAP2C", "COL9A3", "PPP2R5E"
RBM47", "RARA", "SEPHS2", "NLRP2", "SPINT2", "RNF43", "HOPX", "TAOK3", "SLC25A6", "SEL1L3", "PDZ
1", "MUC1", "RAB9A", "VPS35", "FGB", "ZNF217", "LIMK2", "SELE", "CTSA", "MBNL2", "GPR137B", "USP
AKAP9", "SLC35A3", "PCGF2", "RAB9A", "BAZ2B", "MBNL2", "TXNDC9", "HOPX", "USP6NL", "TM4SF1", "
NB9", "SIRT3", "S100A2", "MS4A4A", "IL13RA1", "XPO7", "SH3BGRL3", "SEC14L1", "MFAP2", "CERK", "C
HSPB8", "PNN", "TNFRSF12A", "PLK3", "TSPYL1", "SLC20A1", "TSC22D2", "EPHB2", "MED13", "G3BP2", "
NHBB", "ADO", "IRF6", "OBSL1", "DUSP3", "NFKBIA", "APOD", "IL17RB", "MCL1", "SMAD7", "NRIP3", "PN
N2", "TCN1", "PLAU", "QPCT", "CEACAM6", "TACSTD2", "ALDH1A3", "DDIT4", "PTPRK", "CTSC", "PIK3R3"
RCKS", "POLD4", "RBPMS", "CLCA2", "ANXA4", "BST2", "CEACAM1", "TPD52L1", "FMO1", "S100A7", "M
0A7", "SRC", "SPP1") c("SMARCA2", "SCD", "PIK3IP1", "S100A11", "TBC1D9", "SHROOM2", "RUNX1", "M
16", "SIK1", "CCL20", "UPP1", "HBEGF", "MMP3", "SOX9", "NRIP1", "CYP4F11", "PCDH9", "B4GALT5", "N
MP3", "RFTN1", "LTF", "TACSTD2", "PTP4A1", "ALOX5AP", "ADCY7", "TCN1", "KRT17", "PLAC8", "SLC7A7

, "ANXA4", "ALOX5", "SLC16A1", "IRF9", "IMPA1", "CHI3L2", "PLAU", "CTSC", "MARCKS", "ID1", "IER3IP1
IPL", "USP1", "FAM129A", "SAP30", "TXN", "ANP32E", "DNAJA3", "ADCY7", "POP5", "DNMT3A") c("SLC2
HGD", "ZFAND5", "PVR", "KRT14", "TMEM147", "APOLD1", "TACSTD2", "SCAMP3", "PTP4A1", "SCFD1",
HOC7", "SLC7A1", "NRIP1", "TPBG", "S100A7", "KRT7", "TXNRD1", "APOLD1", "FABP4", "ADCY7", "USP6N
.1", "IL6R", "SHB", "NRIP1", "MREG", "EGR3", "PRKCB", "S100A10", "OBSL1", "SOX9", "SLC20A1") c("MM
"SLC19A2", "KLHL21", "POGK", "MYC", "YME1L1", "GCNT1", "ZFP36L2", "SHC1", "PRNP", "ZSCAN18", "R
AM", "TFF1", "INPP4B", "ANPEP", "STOM", "ADCY3", "GRWD1", "ID1", "IFITM1", "NRIP1", "CXCL5", "AZIN
13", "FHIT", "ARMCX6", "ECHDC2", "NTM", "ADAM12", "TLK2", "ESR1", "H1F0", "SCNN1A", "SH3YL1", "S
(1", "SEC14L1", "PIM1", "CSTB", "NRIP1", "EGR3", "EGR2", "MYC", "KCNN4", "EPHB2", "SOX9") c("BCAR3
G1", "AHCY", "MBTPS1", "MMP12", "DDIT4", "BCL2A1", "QPCT", "IDH1", "HEBP1", "TCN1", "MYOF", "TS
F10", "KRT14", "SPP1") c("SLC7A8", "MAP1LC3B", "SLC24A3", "TP53TG1", "SH3YL1", "FBXW4", "AREG",
.0", "LYN", "LAMP3", "S100A2", "MSN", "AQP3", "IL1RN", "MMP1", "KCNS3", "CASP7", "LCK", "ASPN", "
1", "CDK7", "SEC14L1", "CCL20", "TRA2A", "SPAG1", "FOSB", "ARG2", "ANXA3", "S100A10", "SPRR3", "M
(1", "S100A7", "CXCL2", "IL1RN", "AGPAT2", "PDIA6", "TCN1", "PCSK1", "POLR2L", "TUBB6", "SLC43A3",
"MBTPS1", "MT1M", "CCL4", "IFITM1", "MYL4", "FAM3C", "ALOX5", "IFT57", "CXCL10", "PSME2", "NQC
2", "LST1") c("CXADR", "GSTP1", "DBI", "CYB5R2", "MYL4", "TFRC", "HOPX", "PSME2", "GALNT1", "PI3",
", "SPATS2L", "ARPC3", "ACO1", "IFI44", "S100A10", "OAS2", "PPP1R9A", "AQP9", "APOBEC3A", "CA2", "
", "ST6GALNAC2", "AREG", "SH3YL1", "ECM1", "MEIS2", "TP53TG1", "GSTM3", "DST", "NR1D2", "S100A1
E4", "IARS", "PSAT1", "WDFY3", "TMEM160", "SEC61G", "HELLS", "SAE1", "PCSK1", "DLD", "SMC1A", "XP
PRE2", "CASC3", "PRSS21", "HAPLN1", "SRC") c("CXADR", "IKZF1", "NF1", "GALNT1", "TM9SF2", "OLFM4
R3", "HOOK2", "CHFR", "SERPINB2", "BEX1", "GBE1", "BACH2", "AKTIP", "TCN1", "NME1", "CALML4", "H
VASP", "BEX1", "NDUFB3", "BACH2", "POLR2I", "ACOX2", "POLR2K", "NDFIP1", "NPM1", "RAD23B") c("W
12A", "FOSL1", "PCDH9", "SLCO1B3", "FRMD4B", "HBEGF", "SIK1", "SOX9", "PRKCB", "S100A10", "PPP3C
H1A3", "PLA2G7", "TUBB6", "KARS", "MYO5A", "RGS1", "CCL18", "CXCL2", "TNFRSF21", "ADRB2") c("PRN
E2E3", "S100A10", "TNFRSF21") c("IL18", "AREG", "MTUS1", "SCNN1A", "HIST1H1C", "POLR1D", "S100A1
2", "AHCYL1", "DPH2", "BACE2", "G0S2", "GOT2", "IL1R1", "CD74", "HMGA1", "RBP4", "CCL20", "FLNA", "
RFIP1", "ERGIC2", "CD55") c("DAPK1", "IL7R", "CD8A", "MS4A1", "CYP1B1", "ETS1", "GTF2H2B", "TARP",
OP1", "CCL8", "DNAJA1", "EREG", "DGKZ", "ABCA8", "KLHL21", "SHC1", "CHSY1", "RBM26", "RBPJ", "CD5
SP7", "TREM1", "SLC25A6", "SLC6A14", "LAP3", "C4BPA") c("ESPL1", "FXYD3", "HIST1H1C", "MARCO", "T
LYPD1", "CHIC2", "TEAD4", "HTR2B", "CLEC7A", "F2RL1", "KLK6", "KIF14", "TNFRSF1A", "S100A2", "FSCN
YME1L1", "PRF1", "PTGER2", "CD8A", "GRB14", "TARP") c("EAF2", "SLC43A3", "HK2", "PPP2R1B", "EIF4E
, "SHB", "CYP4F11", "ARG2", "PRKCB", "CD2AP", "ASRGL1", "PELI1", "CCL20", "SLC20A1", "HIPK3", "CFI",
HL21", "ABI3BP", "DGKZ", "SPATS2") c("BAALC", "ASAH1", "PLSCR4", "RBP1", "ITGAE", "EFHC1", "KIF11",
DA2", "NQO2", "GPRC5A", "KCNJ16", "PAX8", "GALNT2", "LAPTM4B", "CDK6") c("KLRC3", "FXYD3", "MTU

/51" "V52" "V53" "V54" "V55" "V56" "V57" "V58" "V59" "V60" "V61" "V62" "V63" "V64" "V65" "V66" "V
.3A", "PLAC8") c("CD55", "MYC", "APOD") c("DAAM2", "MTMR11", "CAMK2N1", "TPSB2", "TPSAB1", "HI
'3CA", "S100A10", "SLCO1B3", "SOX9", "SPAG1", "TNFRSF12A", "TPM4") c("CACYBP", "CCL5", "CD8A", "C
N12", "SPAG1", "TNFRSF12A") c("CALR", "GRB14", "HIST1H2BD", "PCCA", "RBP1", "SAMHD1", "SERINC5"
LIM5", "SOX9", "STK17A") c("ADAM10", "ALDH1A3", "AQP3", "ARPC3", "AXIN1", "CCL20", "CLIC1", "CTN
EG3", "P2RX4", "GRB14", "SPCS3", "RAC2", "YME1L1", "USP7", "MS4A1", "RCN1", "RAB11FIP2", "PGF", '
TFDP1", "CAMK2N1", "F12", "PHKA1", "NUP85", "GSTT1", "ANXA6", "SORT1", "AP1G2", "EFHC1", "TPSAI
G15", "ADA", "OAS2", "SUB1", "TSPAN6", "FZD6", "COX7A2", "PDLIM1", "MYOF", "CLCA2", "DDIT4", "FA
N", "RRAGA", "LSM1", "RGS4", "SNX3", "MUC1", "AREG", "SERPINA3", "DYNLT1", "S100A7", "GPRC5A")
LB", "RBM47", "DDX21", "CCRL2", "CDK6", "LPXN", "SAT1", "TJP2", "ID3", "NXT1", "SRGN", "LYN", "ALDH
, "IRF6", "PRNP", "ABCA8", "EPHB2", "IDS", "SMAD7", "SLC19A2", "MCL1", "NFKBIA") c("SUSD4", "TUFM
LC7A8", "HIST1H1C") c("SOX9", "GRIA2", "IFRD1", "CCNL1", "FOSB", "SIK1", "ATF3", "KLHL21", "NR4A2",
MB9") character(0) c("GAD1", "NRCAM", "CEACAM6", "SAT1", "HIST1H1C", "SERPINB4", "HOPX", "PSME
TLR4") c("INHBB", "TRIM16") c("ASPH", "TNFAIP2", "CAMK2N1", "ADAM28", "PTHLH", "ALOX5AP") c("S
"ADD3", "MED13", "TARP") c("ZBED1", "ATXN7", "KLK3") c("ANXA4", "TRIM22", "RGS4", "CYP3A5", "FXY
.", "ASRGL1", "CFI", "PELI1", "IL1A", "FOSB") c("SOAT1", "PEG3", "FAM98A", "CCL2", "YME1L1", "IL7R", "
XDN", "PRNP") c("IMPDH2", "MPZL1", "CAMK2N1", "RBP1", "PPP1R9A", "GPD1L", "HIST1H2BD", "ATP1/
LC25A32", "TIMM9", "DHCR24", "AREG", "HIF1A", "FASN", "MCTS1", "HK2", "NME1", "SMYD2", "LGALS3
DIT4", "IER5", "SLC25A6") c("HCCS", "GSTM3", "TM4SF1", "AREG", "FHIT", "CLDN8", "SCNN1A", "GPD1L
"ANXA3", "ETNK1", "FOSL1", "PPP3CA", "EZR", "MET", "ARG2", "RCN2", "HSPB8", "EGR2", "SEC14L1", "S
NAMPT", "RRAS2", "GARS", "SRGN", "MYC", "RAC2", "PLIN2", "BAG1", "S100A2", "KRT14", "LPXN", "PPT
ADO", "RRAS2", "MYC", "HGSNAT", "USP7", "NRIP3", "JUND", "INHBB", "ABCA8", "ZSCAN18", "RHBDF1",
5", "BBX", "ARMCX6", "SERPINA5", "TGFA", "RBM47", "COMMD8", "PHLDA2", "NFATC1", "TSEN2", "LAF
"ABHD2", "CANT1", "MBOAT7", "FBXO7", "SERPINB1", "SCGB2A2", "AREG", "IDE", "SH3TC1", "CD59", "I
BB3", "DSTN", "RAB5B", "TRIM22", "CEACAM6", "HELLS", "ABCA12", "AKR1B10", "E2F2", "ENAH", "CKS2
AT1", "GSTK1", "MT1E", "DDX42", "TBC1D8", "DGKD", "ANXA1", "NRGN", "UBE2E3", "CX3CR1", "NLRP2"
', "RGS4", "SWAP70", "XPNPEP1", "GYG1", "GSTP1", "LTA4H", "MAST4", "STK38", "SLC25A6", "HLA-F", "I
)", "TJP2", "TMPRSS4", "CDK6", "CDC42SE1", "BAG3", "DAZAP2", "LYPD1", "ALDH1A3", "DKFZP586I1420
ETN2", "ELF5", "CXCL13", "IGF2BP3", "TMEM123", "FAM57A", "TIPRL", "TSEN2", "KCTD3", "CASP1", "DN
CAN1", "TACC2", "PPM1A", "ACOX3", "YPEL1", "RAB4A", "SASH1", "CCR2", "NRP1", "RCN2", "SH3YL1", "F
, "FBXO11", "MAP3K8", "LAMB2", "MBNL2", "KHNYN", "PERP", "DGKD", "CSF1R", "ASPN", "FABP4", "FIL
"HGSNAT", "CCDC85B", "BCL2L1", "S100A2", "ARMCX1", "MTMR2", "MCM6") c("HOPX", "STK24", "STX
", "TMEM165", "CSRP2", "COMMD8", "RAB2A", "SLCO4A1", "MYBL1", "AREG", "REL", "USP6NL", "UBE2E
, "MUC1", "ALDH3B2", "ENPP1", "BST2", "LPCAT1", "VPS13C", "GAS6", "AKAP12", "CFD", "GPX1", "CILP"
"CLDN10") c("HEBP2", "HOPX", "STXBP2", "PRR16", "S100A11", "BAZ2B", "NDFIP1", "OLFM4", "MVP", "
E1", "ITGAV", "KYNU", "FBXO11", "MAP3K8", "MBNL2", "SEMA3C", "F5", "ARPP19", "MMD", "PLAT", "SF
C15A2", "ZIC1", "TP63", "TCN1", "SLC25A13", "MNAT1", "CDS1", "WFDC2", "RUNX1", "LSM5", "IKZF1", "
12A", "PGAM1", "GSTT1", "TMC5", "NLRP2", "PAPSS1", "PDZK1IP1", "TGFA", "RGS16", "RBM47", "GSTA4
MMD", "PLAT", "SCML1", "BCL2L1", "SPP1", "TMEM126B", "TAPBP", "USP6NL") c("SLC2A6", "CPA3", "LTI
MEM126B", "RAD23B", "BEX1", "USP6NL", "CLDN10", "HIP1") c("HOPX", "STXBP2", "YWHAE", "ZNF217",
, "INHBB", "S100A2", "LPXN", "FHL1", "SULT1A1", "THBS2", "MYL9", "HSPA12A", "FZD7", "SERPINB9") c(
5", "PLAT", "SCML1", "NAMPT", "PDZD8", "SPP1") c("TPI1", "MFAP2", "HOXA10", "SERINC5", "CYB561",
D55", "MYC", "ETNK1", "ANXA3") c("SOAT1", "PTGER2", "PEG3", "HMGCS1", "RCN1", "CCL2", "HYAL2", "
ME1L1", "ADO", "OBSL1", "GRIA2", "IL17RB", "MEAF6", "NFKBIA", "RUFY2", "DUSP3", "LYPLA1", "CD55",
YB5R1", "SERPINB1", "RTP4", "ANXA4", "INPP4B", "PLS1", "TACSTD2") c("GPD1L", "POLR1D", "AREG", "E
3R3", "BCAP31", "PSPC1", "YME1L1", "TFPI2", "MAN2A1", "SMYD2", "CHFR", "S100A2", "RARA", "PEX11
RRAS2", "MET", "EZR", "CCL20", "TRIM16", "PRKCB") c("MS4A1", "CD8A", "TARP", "CCL5", "GRB14", "PK

NRIP3", "SPAG1", "ABCA8", "EREG", "APOD", "FSCN1", "PNN", "DVL1", "PPP3CA", "DGKZ", "ADO", "TSC2
H1C", "MKNK2", "MTUS1", "S100A10", "ARMCX6", "HSPB8", "ECM1", "TSC22D2", "SH3BP4", "ZNF586", "
FGF9") c("MS4A4A", "MRPL19", "RARRES1", "SMYD2", "PLCB3", "XPO7", "PRR16", "RBM3", "CCDC47", "
LOX5AP", "CAMK2N1") c("SFRP1", "ZMAT3") c("B4GALT5", "SERPINB4", "SCEL", "TSPAN6", "PHLDA2", "H
, "OBSL1") c("USP7", "HMGCS1", "ADD3", "TARP", "ATP10A", "SKAP1", "PQBP1", "DAPK1", "PKP4", "PTG
AA", "DIXDC1", "RTN3", "CRK", "NDRG3", "RACGAP1", "ALDH7A1", "SERINC5", "DAAM2", "CAMK2N1", "
"TRIM16", "GRIA2", "ADO", "DUSP1", "PCDH9", "SOX9", "REG3A", "FUT8", "CFI", "NR4A2", "SIK1", "OBS
"TP53", "MEA1", "DAPK1", "SLC5A6", "YME1L1", "CTSW", "PRF1", "CD8A", "COL15A1", "CCL5", "PEG3",
N1", "SIL1", "NAAA", "MTMR11", "CDIPT", "TNFAIP2", "SHMT2", "IMPDH2", "DEFA4", "F12") c("NUAK1",
102A", "MKNK2", "GSTM3", "LPCAT4", "IKBKE", "CLDN8", "DST", "DGKA", "FXYD3", "TSC22D2", "VAMP1
A3", "MTHFD2L", "OBSL1") c("HSD17B4", "KCNQ1", "CYP1B1", "PQBP1", "ADD3", "MMP10", "SPCS3", "
"HSD17B4", "TBC1D12", "IFT57", "GSTT1", "BAALC", "NAAA", "GCH1", "ITGAE", "SLC3A2", "CCDC28B", "
"GNG10", "KCNJ16", "MAST4", "NQO2", "LSM1", "ANXA1", "PLS1") c("ERLIN2", "HCCS", "HUWE1", "MT
", "ADD3", "ASPH", "MTRR", "MMP10", "SHC1") c("HLA-DQA1", "KCNS3", "HRK", "SNRPA1", "PAK2", "IL
PH", "HMMR", "F12", "PTHLH", "DEFA4", "CNR1", "VWA5A", "STK38", "TPSAB1", "TPSB2", "PSMB9", "BN
LC24A3", "TLK2", "DST", "HSPB8", "IKBKE", "AREG", "RAD54L", "MPO", "TOP2A", "SH3YL1", "CD40") c("
MR", "ITGAE", "GCH1", "PSMB9", "RBP1") c("HLTF", "TARP", "SFRP1", "SORL1") c("SAT1", "SERPINB4", "
CDH9", "SOX9") c("HMGCS1", "RCN1", "EPPK1", "PRF1", "DENND2D", "CTSW", "CCL5", "AMPH", "GMPS"
", "PPP3CA", "EREG", "ADO", "LYPLA1", "FSCN1", "IL17RB", "SHC1") c("TPSAB1", "CTBP1", "ADRB2", "RA
, "UBE2E3", "DYRK3", "SLC25A6", "ID3", "PLAT") c("CRABP2", "NR1D2", "HIST1H1C", "PHGDH", "TLK2", "
, "EREG") c("RBP1", "RTN3", "FGF9", "IARS2") c("CES3", "FGF9", "PTX3") c("LST1", "SERPINB6", "PHLDA2"
"HS3ST1", "CCDC86", "CAMK1", "RHOBTB3", "NAMPT", "RAD23A", "EHBP1L1", "PEA15", "KIF5B", "CLDN
A", "CCL20", "PITPNC1", "HMGB3", "ISG20", "LYN", "ELF4", "CEACAM1", "DNAJB6", "RBP4", "IFI16", "CY
", "TACSTD2") c("OBSL1", "FDX1", "NR4A1") c("GAMT", "RBP1", "FBXO21") c("SGCB", "FEZ1") c("FDX1", "
KN2", "TPM4", "PNN", "PPP3CA", "SEC14L1", "USP7", "HBEGF", "MTHFD2L") c("MMP10", "TP53", "TMX
B", "CDC25B", "CCL18", "CXCL2", "HBEGF", "TIAM1", "THBS1", "TNFRSF21", "MMP9", "MAST4", "RRS1",
BL2", "CYP1A1", "SLC15A2", "CFI", "PHF11", "DYNLT1", "PDLIM1", "IL1RN", "CTDSP1", "LRP10", "NCOA1"
, "FMR1", "COMMD10", "PAPSS1", "H2AFY2", "STOM", "TCN1", "HDAC1", "DAZAP2", "HEBP1", "CLCA4",
3", "HMGN1", "DNAJA3", "LIMK2", "AKR1B10", "ABCA12", "CARHSP1", "HOPX", "CEACAM6", "DSTN", "T
6", "RAB38", "S100A10", "DDX42", "SMPDL3A", "S100P", "CEACAM6", "TTC9", "KRT14", "RBPMS", "SLC6
EG", "GALNT1", "MAST4", "PHLDA1", "GYG1", "PSME1", "HLA-B", "RAB5B", "TAPBP", "NFKBIE") c("PPL",
T2", "PLA1A", "LIPA") c("HES1", "SMAD3", "TACSTD2", "TGIF1", "RALGDS", "DNASE1L1", "SIDT1", "COM
"SLPI", "AREG") c("HEBP2", "CXADR", "BCAM", "S100A11", "TCN1", "MVP", "OLFM4", "PLAC8", "PRDX3"
", "KYNU", "CLCA2", "PHF11", "RNASE6", "IRF6", "TBC1D8", "IL13RA1", "IDS", "CASP4", "CERK", "NDUFA
, "GPX2", "KRT7", "S100A7", "SEMA3C", "DYNC2LI1", "LAMA5", "S100A11", "RARRES1", "STOM", "MVP"
, "FBXO2", "MAT2B", "VPS35", "MPPED2", "PIK3IP1", "TTC9", "STK38", "SLC15A2", "IL13RA1", "NCOA2",
K1IP1", "SEC11A", "GSTT1", "HIST1H2AE", "PPIC", "TPM4", "NME3", "VAMP5", "SLPI", "SUV39H1", "KLH
6NL", "KYNU", "CLCA2", "TBC1D8", "IL13RA1", "DMD", "IDS", "MUC16", "CERK", "PRKAR2B", "HTR3A", "
"CLCA2", "PHF11", "RAD23B", "LY75", "MRPS16", "NDUFA3", "WASL", "BEX1") c("TACSTD2", "DUSP1", "
DR2L", "EDN1", "GALNT1", "LRRC15", "FHL1") c("TACSTD2", "GABRE", "CTSH", "GPX2", "MAP3K4", "FGG
"PCDH9", "SPRR3", "EGR2", "TRIM16") c("DENND2D", "CYP1B1", "HERPUD1", "COL15A1", "PEG3", "DEC
NN", "TNFRSF12A", "PLK3", "TSC22D2", "GRIA2", "EPHB2", "MED13", "PLAGL2", "DNMBP", "HGSNAT", "
, "NFKBIE", "TFF1", "TXNDC9", "MARCKSL1", "ISG15", "IRF9", "IFITM1", "CXCL10", "ID1", "ZNF586", "HG
AL", "IL1RN", "RBM47", "PDZK1IP1", "SPP1") c("CAST", "FBXW4", "DGKA", "ECM1", "SH3BP4", "SH3YL1",
INPP1", "FMO5", "ZIC1", "IKZF1", "SLC27A2", "PCCB", "ZNF586", "ZBTB38", "WNT5B", "TMX4", "OLFM4
MET", "UGCG", "ANXA3", "OBSL1", "EZR") c("DAPK1", "CYP1B1", "CD7", "MMP10", "PRF1", "MID1", "COL
", "TYROBP", "CCDC86", "LAPTM5", "CD53", "RRS1", "RGS1", "IFI16", "ARHGDIB", "PLAT", "SRGN") c("D

", "EPS8L3", "UBE2I", "ERH", "VPS35", "ADORA2B", "G3BP1", "MMP12", "KRT5", "PLEK2", "SERPINB1", "
5A13", "PMM2", "ANXA4", "ADRM1", "IER3IP1", "HK2", "CTSE", "ERH", "HCCS", "LGALS3", "TXNDC9", "T
"GBP1", "KRT17", "PAK1IP1", "STIP1", "PLAC8", "FOXC1", "PSCA", "SDC1", "SERPINA3", "IFI16", "LMO2",
NL", "AEBP1", "LMO2", "SPP1") c("FAM102A", "MCM3", "NUSAP1", "SPIB", "SLC39A14", "CCDC47", "S10
IP10", "PRF1", "CD8A", "CYP1B1", "COL15A1", "SGCD", "PKP4", "YME1L1", "MAPK9", "TP53", "DAPK1", "
ALGAPA1", "OBSL1", "RBPJ", "CCL8", "APOD") c("SLC25A4", "PGD", "DIXDC1", "HMMR", "RAB33A", "IMI
N1", "TSPAN6", "HOMER1", "ANXA4", "SLC16A1", "TSPAN31", "STEAP1", "RGS16", "CXCL10", "DDX60", "
T6GALNAC2", "TM4SF1", "EPHX2", "SLC35A3", "JAG1", "AKAP7", "IQCG", "TMEM70", "AREG", "S100A10
", "SLC5A6", "PEG3", "SKAP1", "DENND2D", "MMP10", "HLA-DOB", "HSD17B4", "MID1", "RAC2") c("SAT
PAN31", "PLAU", "ALDH1A3", "TFF1", "ISG15", "CAPN2", "CXCL10", "SERPINB1", "CLCA4", "PDLIM1", "C
"JAG1", "RAP1GAP", "GTF2IRD1", "SCNN1A", "MKNK2", "CROT", "KCTD3", "IDH1", "SH3GLB2", "MPO", "
CDH3", "ALDH1A3", "S100A7") c("MMP3", "YME1L1", "HGSNAT", "CACNA2D2", "NRIP3", "ADO") c("HSD
IMP3", "ASRGL1", "PCDH9", "SLCO1B3", "STX3", "CFI") c("MID1", "TARP", "CCL2", "TFDP2", "SLC5A6", "P
"TMEM158", "ARHGAP26", "PTP4A1", "DDX21", "H2AFZ", "SLC6A15", "CDC25B", "LYN", "RAC2", "CD19"
D2", "CDK6", "NETO2", "TM9SF2", "EIF4A3", "ADA", "CEACAM6", "ID1", "PRPF4", "SUZ12", "DSG2", "PDL
"CEACAM6", "ADK", "ATF1", "ANP32E", "CCNA2", "NDUFB2", "MSN", "EGF", "BTG3", "STT3A", "HNRNPL
PSCA", "TMED9", "SWAP70", "HERC6", "PITX1", "S100A7", "ARNTL2", "KCNJ16", "CXCL5", "KRT5", "MT1
10", "PHGDH", "PEBP1", "HMGA1", "SLC2A10", "OSTF1", "MPO", "MKNK2", "MT1H", "HIPK2", "CACNA2I
O1", "NME1", "AKAP7", "VDAC1", "TYRO3", "HMOX2", "SCD") c("RHOBTB3", "APBA2", "SIK1", "GABPB1"
", "SUZ12", "ZIC1", "MICA", "MAL", "LSM5", "RBBP4", "ATP10B", "SLC35E3", "SLC39A4", "TP53TG1", "ND
APLN1") c("CXADR", "WWTR1", "MEIS1", "VAMP5", "PGK1", "S100P", "CYB5A", "ABLIM1", "ALDH6A1", "
WTR1", "HOPX", "SAT1", "TM9SF2", "PDE4B", "DUSP1", "LSM6", "KLF9", "EREG", "LIMS1", "RBBP5", "KL
A") c("RAB11FIP2", "PRF1", "IL7R", "PEG3", "TARP", "RCN1", "TCERG1", "ADD3", "CDC42EP3", "LCP1", "
NP", "TRIM16", "TNFRSF12A", "PXDN", "DGKZ", "INHBB", "ABCA8", "EREG", "DNAJA1", "GRIA2", "APOD",
.0", "VAMP1", "MT1H", "TIMM13", "CLDN8") c("KCNN4", "TRIM16", "PCDH9", "SIK1", "SOX9", "GRIA2",
"ALDH1A3", "ID1", "MSN", "FGFR3", "NAMPT", "LYN", "MMP1", "PBLD", "RHOBTB3", "HLA-DQA1", "SOD
"PRF1", "PEG3", "SHC1", "MMP10", "CTSW", "CHSY1", "UGP2") c("KIF14", "HSPH1", "S100P", "PAF1", "
55") c("RAB20", "TNFAIP2", "MBOAT7", "ADAM28", "IARS2", "PRR11", "NAV3", "PTHLH", "SERINC5", "CR
TMEM70", "AREG", "TOP2A", "MCM7", "ZNF365", "SH3BP4", "SLC25A14", "MTUS1", "CAST", "PPFIBP2",
1", "TPPP3", "EMP2", "TPM3", "LAMP3", "PPP4R1", "TACSTD2", "IL1R1", "S100A7", "TLR4") c("MCL1", "
E2", "ID3", "RRAS2", "MSN", "KLF10", "ID1", "MID1IP1", "TNPO1", "CNN2", "HSPH1", "SRGN", "PLIN2", "
"MMP3", "GPA33", "EPHB2", "PCDH9", "IL6R", "SPATS2", "UMOD") c("ASPH", "ETS1", "CDC42EP3", "PE
"NCAPH", "IFT57", "F12", "ASPH", "PTHLH", "DIXDC1", "GCH1", "PPP1R9A", "BPHL", "ADAM28", "HIST1
JS1", "TKTL1", "HIST1H1C", "TOP2A", "CD2AP", "SLC7A8", "TLK2", "NAV2", "MEIS2", "PPFIBP2", "CRABP2

'67" "V68" "V69" "V70" "V71" "V72" "V73" "V74" "V75" "V76" "V77" "V78" "V79" "V80" "V81" "V82" "V8
IST1H2BD", "DEFA4") c("ADD3", "GADD45B", "SORL1") c("SAT1", "HIST1H1C") PEX11A c("PLK2", "HBD",
CTSW", "GRB14", "HMGCS1", "IL7R", "PEG3", "PKP4", "PTGER2", "SOAT1", "TARP", "TCERG1") c("ACTN1
', "SUSD4", "TPSAB1", "TPSB2") c("ATP2A1", "CLIP2", "DUSP3", "EMP3", "FEZ1", "NUAK1", "OSBPL9", "P
ND1", "EPHA2", "EXOC5", "GOT1", "GOT2", "GZMB", "HIST1H2AE", "HIST1H2BD", "HNRNPL", "ID1", "IL7
'SPATS2", "DAPK1", "BCAR3", "HMGCS1", "COL15A1") c("KMO", "OLFML2B", "G3BP1", "TFDP1", "SMPD
31", "TPSB2", "RNASET2", "CSNK1D", "GPD1L", "GRN", "HIST1H2BD", "CRK", "TNFAIP2", "PBXIP1", "DDR
AM3C", "STOM", "PPIC", "KRT5", "BCL2A1", "CAP1", "NEU1", "ALOX5", "HDAC1", "PIK3R3", "TACSTD2", "
c("GSTM3", "IKBKE", "CDKN1B", "STX3", "TM4SF1", "MTUS1", "TKTL1", "JAG1", "PPL", "ST6GALNAC2", "
I1A3", "DDX10", "PLAC8", "ADORA2B", "FOSL1", "HLA-DQA1", "FLNA", "HBEGF", "BMP2", "THBD", "TYRO
I", "HIST1H2BD", "CDK4", "ARHGAP1", "GCH1", "F12", "DAAM2", "MOCS2", "DFFB", "BPHL", "FCGR2A")
"ADO", "VAMP1", "OBSL1", "CCL8", "NRIP3", "PCDH9", "GADD45B") c("RBM47", "DDX21", "CDK6", "SA
E2") c("ARHGEF3", "CYP3A5", "TRIM22", "NRCAM", "IGFBP2", "FXYD3", "MEIS1", "CXCL10") c("HS2ST1",
GCB", "SORL1") c("HIST1H1C", "TSPAN6", "HOPX") c("SLC16A4", "KCNN3", "MEIS1", "ARL6IP5", "FAM13
'D3") PITRM1 c("MVP", "NCOA4", "DDIT4", "PSCA", "SLC25A6") c("FILIP1L", "ENPP1", "RBBP7", "MRPS16
HMGCS1", "MMP10", "UTRN", "LCP1", "SLC5A6", "CDC42EP3", "CYP1B1", "DAPK1", "MID1", "ZNF32") c(
A1", "TNFAIP2", "BAALC", "NUP85", "ADRB2", "IGBP1", "ADAM28", "PIGB", "ALOX5AP") c("G3BP2", "MR
3", "SNRPD2", "S100A11", "SLC1A3") c("AREG", "ACOX2", "DHRS2", "PDZD2", "GINS2", "CAST", "SCAND1
", "NAV2", "PDZD2", "ZNF586", "CAST", "ARMCX6", "PARP2", "LRMP", "DHTKD1", "JAG1", "POLR1D") c("
STX3", "RRAS2", "MYC", "USP7", "PCDH9", "PELI1", "LRRC8D", "LAMC2", "RHBDF1", "SLC20A1") c("SLC5A
2", "LAMC2", "ISG20", "THBD", "CD14", "PTGS2", "GLUD2", "AQP3", "TMEM158", "ALOX5AP", "KLK7", "
"SHC1", "PRNP", "CACNA2D2") c("TNFAIP2", "TPSB2", "TPSAB1", "SERINC5", "NUP85", "CAMK2N1", "NI
P3", "HIST1H1C") c("CDR2L", "ARHGEF3", "ANXA4", "PEX11A", "PPCS", "GLB1", "TACSTD2", "FTH1", "MEI
PTGS2", "RGS10", "PPA1", "USP6NL", "QDPR", "HPGD", "CLIC3", "CNPY3", "NDUFA4L2", "HIST1H1C", "IG
2", "TBC1D8", "FKBP1A", "PPIB", "FAM129A", "DAD1", "DNAJA3", "TMEM165", "FDPS", "COMMD8", "GT
', "EHD4", "PAPSS1", "GRB2", "PMM2", "LMO2", "PDZK1IP1", "ATP6V1G1", "TGFA", "IFI16", "NPTX2", "N
HLA-E", "HLA-B", "PSMB8") c("H1F0", "IDH1", "CLDN8", "S100A10", "BMP7", "TKTL1", "RAP1GAP", "IKBK
", "HNRNPH2", "TRIM37", "BCL2A1", "ALDH7A1", "HIST1H2BD", "PPBP", "IER5", "RALGDS", "SAT1", "ATI
AJC7", "SCD", "FKBP5", "GPSM2", "SMYD2", "SMC1A", "PNKP") c("SPRR3", "CSTB", "TKTL1", "CTBP2", "L
KBP14", "SORBS2", "ADO", "JAG1", "CFLAR", "KCTD3", "CD59", "MTMR2", "WWC3", "RIF1", "GTF2I", "PI
IP1L", "LMO2", "TGFA", "NPTN", "CDR2", "SPRED2", "RGS19", "TSPAN31", "SLCO4A1", "SPP1", "JUND", "
BP2", "YWHAZ", "CDC42", "COL9A3", "PLCB3", "MT1F", "NCOA4", "MVP", "RNF103", "TPP1", "IER5", "RI
E1", "PPFIBP2") c("HEBP2", "HOPX", "KRT7", "SLC5A6", "PIK3R3", "ZNF217", "S100A11", "ELF3", "ERBB2"
', "TBC1D8", "MMD", "PLAT", "SCML1", "CNN3", "WIPF2", "TIMM9", "SPA17", "CCDC92", "SPP1", "INHBI
'MUC1", "MBNL2", "TCN1", "TBC1D8", "PLAT", "SCML1", "WIPF2", "NAMPT", "TIMM9", "PDZD8", "SPP1
P1", "FAM3C", "SWAP70", "ICAM1") c("PEX11A", "PRR16", "S100A11", "HOXA5", "ATP10B", "OLFM4", "
'TMEM123", "RAB2A", "TSEN2", "VAMP8", "GTPBP8", "BEX1", "REL", "PPA1", "TRAK1", "MAP7", "LTA4H
I", "ITGAE", "CR2", "TMEM123", "LACTB2", "TAC1", "RAC2", "SLPI", "TNNC1", "PDCD6", "EI24", "P2RX4",
F", "CTSZ", "TRAF4", "TFF3", "SPINK1", "MT1F", "RAB15", "AKR1C2", "TSPAN15", "CFB", "CGA", "SERPINA
, "IRF6", "SOX9", "TACSTD2", "SLC2A3", "RTN1", "PTPN12", "NDFIP1", "TJP2", "MUC4", "TMPRSS2", "TM
"DDR2", "SLC9A3R1", "GABRP", "ID4", "IQGAP2", "GLB1", "CPA3", "TACSTD2", "FTH1", "PLEKHB1", "MA
"PLXNB1", "BMP7", "SPINK1", "MAGEA3", "COG5", "DACH1", "SORT1", "TM7SF2", "BCAS3", "KAT2B", "
GRB14", "YME1L1", "SLC5A6", "GCDH", "CYP1B1", "DAPK1", "ADD3", "CDC42EP3", "RAB11FIP2") c("CDC
, "MYC", "APOD") c("SAMHD1", "MPZL1", "ZMPSTE24", "GPD1L", "HIST1H2BD", "CNR1", "ECHDC3", "RB
SR1", "TM4SF1", "HMGA1", "ECHDC3", "NAV2", "SH3BP4", "TIMM13", "CCDC6", "SH3YL1", "GSTM3", "C
.B") c("PSTPIP2", "AREG", "RARRES1", "PIK3R3", "ITPR3", "DCAF10", "LPCAT1", "YME1L1", "TFPI2", "LAM
(P4", "BRD2", "PEG3", "DAPK1", "ASPH", "RAB11FIP2", "TMX4", "STAC", "RAC2") c("CLIC1", "AP3D1", "G

2D2", "KLHL21", "PDLIM5", "DUSP3", "RRAS2", "TRIM16") c("ALDH7A1", "PNPLA6", "TPSAB1", "HIST1H2
'AKAP7", "PEBP1", "PPL", "VAMP1", "ESR1", "UXS1", "GSTM3", "GPD1L", "ALDH3B1", "TKTL1") c("IFRD1"
EHD3", "SCML1", "RFC5", "SERPINB2", "POLR2I", "TFPI2", "SNRPF", "ZC3H4", "PLAC8") c("ERBB2", "PDZD
OPX", "MGST3", "NRCAM") c("FXYD3", "SLC16A4", "CXCL10", "TRIM22", "NRCAM", "TACSTD2") c("TAOK
ER2", "CYP1B1", "ETS1", "PEG3", "SLC5A6") c("CLDN7", "KLK7", "ATP2A2", "KLK10", "HMGB3", "KRT16",
'RNASET2", "GRN", "MBOAT7", "ALOX5AP", "SPOCK2", "TNFAIP2", "FCGR2A", "BPHL") c("ADD3", "TARP"
L1", "ATF3") c("KLK7", "ATP2A2", "VAPA", "BCL3", "PPBP", "ID1", "PUF60", "NIP7", "UQCRC1", "TBC1D8
"CYP1B1", "MS4A1", "HLA-DOB", "GTF2H2B") c("PTGS2", "ID1", "RAC2", "PLIN2", "LCK", "CIB1", "G0S2",
"SORL1", "EMP3", "CLDN15", "HLTF", "SFRP1", "ZMAT3", "PDE8B", "ZNF83", "FEZ1") c("SERPINB6", "CE
", "TLK2") c("CFI", "SYNPO", "ATF3", "FOSB", "IER2", "TRIM16", "KCNN4", "RRAS2", "YPEL5", "KLRK1", "
CD8A", "DAPK1", "MS4A1", "RAC2") c("OLFML2B", "PSMD3", "KIF14", "MNDA", "ALOX5", "KLK7", "RBM
PNPLA6", "SKIV2L", "FGFR1", "ADAM28", "NUP85", "TPSB2", "TPSAB1") c("ZFP36L2", "MRPS30", "ADD3"
US1", "SASH1", "HIST1H1C", "STX3", "S100A10", "TM4SF1", "GCHFR", "CD40", "SLC2A10", "SH3BP4", "Z
1R1", "CDC42SE1", "CXCL2", "CYP24A1", "GARS", "EIF4A3", "CCL20", "PWP1", "THBD", "ASNSD1", "EMP
MP4", "FGFR1") c("DUSP3", "CEP170", "MED13", "KCNK2", "TARP", "NUAK1", "ADD3", "STMN2", "MMP3
CNL1", "PAK2", "MED13", "OVGP1", "GRIA2", "VAMP1", "KLHL21", "PFKFB3", "NR4A2", "REG3A") c("DK
SCEL", "CEACAM6") c("REXO2", "CXCL10", "FXYD3", "TRIM22", "ATF3") c("MRFAP1L1", "PLN", "CDKN2C"
, "TARP", "MS4A1", "SSR2", "CDC42EP3", "ASPH", "MID1", "COL15A1", "SHC1") c("ADRB2", "IER2", "CXC
BAC1", "TPSB2", "DENND2D", "FGF9", "RAB20", "CAMK2N1", "CNR1", "ARHGAP6", "F12", "RBP1", "RAP
"AVPI1", "HUWE1", "NAV2", "ESR1", "TOP2A", "TKTL1", "SCNN1A", "TIPARP", "S100A10", "ESPL1", "MPC
", "RANBP9") c("RGS4", "CYP3A5", "CA2", "PSRC1", "MEIS1", "FTH1") c("AREG", "CCK", "CA2", "MEX3D",
N4", "PRMT1", "C1QB", "ALDH1A3", "LTF", "TCN1", "TMEM158", "GNA15", "ALOX5", "TNFRSF21") c("MC
P1A1", "ID3", "CARD10", "IL1R1", "CDC25B", "F2RL1", "ID1", "S100A7", "MMP1") c("IDS", "ZSCAN18", "T
"DHCR24") c("MEIS1", "TRIM22", "PTS", "LAMB3", "ARHGEF3", "CA2", "FXYD3", "TACSTD2") c("PIP", "PL
4", "DENND2D", "ZNF32", "DAPK1", "SLC5A6", "GRB14", "CYP1B1", "PEG3", "AMPH", "CCL5", "HSD17B4
"SIGLEC15", "MMP1") c("OBSL1", "ZFP36L2", "NFKBIA", "SEC16A", "PIM1", "HGSNAT", "CSTB", "SMAD7
, "CYP4F3", "PTGS2", "GNLY", "AMDHD2", "TRIOBP", "CFH", "GRN", "CXCL5", "HBEGF", "JUN", "AREG", "
"PIK3R3", "MAT2B", "FZD6", "VPS35", "KCNJ16", "ZNF586", "IDH1", "UBE2I", "ZMPSTE24", "IRF9", "DD
BC1D8", "IL13RA1", "E2F2", "BTG3", "GALNT1", "PSAT1", "DBI", "RAB5B") c("SSH3", "IRAK1", "TSPAN31"
A14", "DHCR24", "OSBPL10", "STK38", "SCNN1A", "TAPBPL", "ZFAND5", "SLC25A6", "IFNAR2", "CLDN7"
"MEIS2", "PPFIBP2", "GPD1L", "CRABP2", "SH3YL1", "FXYD3", "FGGY", "TKTL1", "JAG1", "TP53TG1", "SL
", "GPRC5A", "JUNB", "CD9", "SOX9", "TNFAIP6", "YPEL5", "SLC12A2", "PPARG", "HOMER1", "MLF1", "I
, "MUC1", "RAB9A", "BAZ2B", "SELE", "MBNL2", "STXBP2", "HOPX", "TTC9", "NAMPT", "USP6NL", "CLCA
3", "PRKAR2B", "AREG", "GALNT1") c("HOOK2", "LXN", "HIST1H2BD", "ERBB2", "CD9", "DAAM1", "EIF5"
, "CDC42SE1", "PLAC8", "PIK3R3", "ZMPSTE24", "MBNL2", "HOPX", "KYNU", "F5", "IDS", "CERK", "ARPP
, "MNAT1", "LSM5", "NDUFA3", "TP63", "GALNT1", "MGST3", "BEX1") c("TACSTD2", "CDH1", "TRAF5", "
DC10", "MAEA", "WARS", "TECR", "TNNC1", "RAB5B", "NFKBIE") c("TACSTD2", "ALDH3B2", "LXN", "HEB
GPX4", "GALNT1", "TAPBP") c("FLNA", "GABRE", "CFB", "SPINK1", "LTF", "SLC2A6", "BNC2", "AKR1C2", "
TMPRSS2", "SOX9", "SAT1", "GCH1", "TJP2", "FMR1", "CLDN4", "DPYD", "TPK1", "MUC1", "RAB9A", "ZNF
Y", "ZNF226", "ID4", "FTH1", "DYNC2LI1", "FRZB", "MAGEA3", "PLEKHB1", "ABCA8", "RHOD", "ABCC5",
R1", "MAN1A2", "SLC5A6", "GPR65", "PRF1", "TMX4", "CCL2") c("TRIB1", "KLK10", "TMPRSS4", "FLNA", "
IUND", "TRIM16") c("DENND2D", "CAMK2N1", "GRN", "TNFAIP2", "GCH1", "PRKX", "TSPYL4", "GSTT1", "
D", "STAT1", "WNT5B", "GZMH", "CSTF2T", "ARMCX5", "AIMP2", "RFC5", "DDX60", "TMEM106B", "KCN
, "CRABP2", "JAG1", "S100A10", "MKNK2", "RAP1GAP", "FXYD3", "ST6GALNAC2", "AREG", "S100A13", "S
", "LIMK2", "PEG10", "MAL") c("SRPX", "S100A11", "IL17RB", "SLCO4A1", "TSPYL5", "EGR3", "TSPYL4", "I
15A1", "PEG3", "ASPH", "SLC5A6", "MS4A1", "HMGCS1", "PKP4", "UTRN", "GTF2I", "CACYBP") c("ASPN"
GKZ", "RPS6KA3", "DUSP3", "SLC19A2", "ZSCAN18", "KLHL21", "SMOX", "TERF2", "PLAGL2", "PRNP", "EP

'ATP1B3", "TXNDC9", "GABRE", "FERMT1", "CEACAM6", "TLE2", "HNRNPD", "HCK", "STEAP1", "ZNF586",
'UBA4A", "FERMT1", "SLPI", "PRPF4", "NQO2", "THOC7", "SLC1A3", "SLC7A1", "PRDX1", "AREG", "CD58"
"GABRP", "BST2", "PLAT", "SPP1") c("PARP2", "FAM102A", "HSPB8", "CRABP2", "SASH1", "S100A10", "
0A2", "KLF5", "SERPINB2", "CYP11A1", "USP1", "SLC38A1", "POLR2I", "RAB33A", "CDK5RAP2", "RFC5", "
'NUDT1", "ASPH", "SHC1", "PEG3", "CCL5") c("GTF2B", "PCSK1", "MAP3K8", "IL1RN", "TMEM165", "HM
PDH2", "ARCN1", "PPP1R9A", "AIFM1", "VWA5A", "PABPC4", "HIST1H2BD", "PDLIM2", "MCM6", "THBS
NETO2") c("TMEM165", "TIMELESS", "H2AFY", "CXADR", "TUBA1A", "BCAP31", "HOPX", "ERBB3", "PBL
", "MEIS2", "CD40", "DST", "CRABP2") c("FOXA1", "ATF3", "MED13", "KLRK1", "ADO", "KLHL21", "FUT8"
T1", "CDKN1A", "SLC7A7", "IER5", "EDEM1", "TMPRSS4", "CDC42SE1", "MMP12", "S100P", "HS3ST1", "T
YP3A5", "TACSTD2", "NR4A2", "SH3BP4", "PAPSS1", "PLEK2", "CCL4", "YIF1A", "SLC25A6", "NRIP1", "PLE
'ECHDC2", "GSTM3", "FXYD3", "NBR1", "DST", "CRABP2", "LGR4", "HIST1H1C", "SLC2A10", "BMP7", "S1
17B4", "PLA2G4C", "CNR1", "GGCX", "RBP1", "ASPH") c("MMP3", "ZFPM2", "WWOX", "TARP") c("RBKS",
'EG3", "HP1BP3", "GTF2I", "SPIN1", "ADD3", "HSD17B4", "UGP2", "SOAT1", "PRF1", "SHC1", "HMGCS1",
, "KLK3", "PLA2G7", "DIAPH1", "ARFGAP3", "GZMB", "OSBPL10", "HS3ST1", "CCDC86") c("CD55", "KLHL
IM1", "ADAM9", "CHI3L2", "ANAPC13", "TMEM106B", "MMP1", "SSBP1", "STEAP1", "ANXA4", "QPCT", "
", "DNAJB6", "GFPT1", "FAM129A", "PSAT1", "E2F2", "SLC25A46", "SPRR1B", "CARHSP1", "SEC61G", "H
E", "SDC1", "IL1RN", "MMP3", "SLC25A11", "S100A11", "CTSC", "ATOX1", "CAPNS1", "GPRC5A", "UBE3C
D2", "VAMP1", "IKBKE", "STX3", "ZNF586", "AKAP7", "TKTL1", "ERLIN2", "CAST") c("CD55", "PFKFB3", "S
", "NR4A2", "PARM1", "ALDH1A2", "LAMB1", "IGFBP1", "SPRR3", "KRT23", "FLRT2", "HMGN3", "TCL1A",
UFA3", "CDS1", "CSDE1", "UQCRQ", "ZBTB38", "SEC61G", "NDRG2", "S100A11", "MOCS2", "HMG20B",
'GALNT1", "SAT1", "PDE4B", "PI3", "CCND3", "PDZK1IP1", "LSM6", "LIMS1", "METTL5", "KLF6", "EHF", "
F6", "MDM4", "ATP6V0B", "G0S2", "AQP9", "IRF6", "YWHAE", "HMGA1", "VASP", "ARNTL2", "GBE1", "
SIGLEC1", "CTSW", "CCL5", "PTGER2") c("THBS1", "CYBB", "PRNP", "IER5", "CLEC7A", "OSBPL10", "SOD2
"PPP3CA") c("PPP1R9A", "CAMK2N1", "RNASET2", "CTBP1", "HIST1H2BD", "RBP1", "SERINC5", "CHERP
"IFRD1", "VAMP1", "TIMM13") c("SPR", "IER5", "PEA15", "HIST1H2BD", "PRMT1", "EDEM1", "IL7R", "M
D2", "ME2", "TMEM158") c("CACNA2D2", "ZSCAN18", "DVL1", "EREG", "GRIA2") c("GAMT", "CTBP1", "R
LAMP3", "AGPAT2", "PLA2G7", "THBS1", "ITGB3BP", "KYNU", "HSD17B2", "HMGB3", "FPR3", "ALOX5", "
K", "DFFB", "RNASET2", "PNPLA6", "ANXA6", "CNR1", "CDC34") c("SIX2", "TARP", "NUAK1", "EMP3", "P
"MKNK2", "CRABP2", "ESR1") c("KCNN4", "CCL8", "PELI1", "CFI", "PCDH9", "KLHL21", "ATF3", "LRRFIP1"
FSCN1", "PDLIM5") GRB14 PCSK5 c("UBE2G1", "TGFA", "GRAMD1C", "NRCAM", "HOPX") c("SAMD4A", "
RBP4", "KLHL2", "HLA-DQA1", "TLR4", "POLR2L", "LY96", "GOT2", "CHCHD3", "VCP", "CLDN7", "PDE4B",
G3", "ADD3", "SLC5A6", "CD3G", "BRD2", "SPATS2") c("PBLD", "HTR2B", "TPPP3", "CXCL6", "ADORA2B",
H2BD", "CAMK2N1", "CNR1", "PTK2B", "ADRB2", "IARS2", "NUP85", "PPP1R13B") c("GGA1", "BBC3", "M
", "ESR1", "SCNN1A", "NR1D2", "MT1H", "GSTM3", "DGKA", "MARK1", "CD40") c("TRIM16", "GPAA1", "

33" "V84" "V85" "V86" "V87" "V88" "V89" "V90" "V91" "V92" "V93" "V94" "V95" "V96" "V97" "V98" "V9
"TMEM38B", "ASPN", "FUCA1", "LTF", "PEX11A") c("MARCKS", "NOSIP", "S100A9") c("TMEM176A", "CE
", "ADAM8", "ALDH1A3", "ANXA2", "AQP3", "C1QB", "CCL18", "CCL20", "CD14", "CDC25B", "CLIC1", "CN
TX3", "RABEP2", "SFRP1", "SGCB", "SORL1", "TARP") c("CEACAM6", "DHCR24", "DHRS3", "FERMT1", "GA
'R", "KCTD5", "KIF5B", "KLK6", "KLK7", "LAMB3", "LYPD1", "MSN", "NIP7", "PACSIN3", "PDE4B", "PRMT1
L3A", "RAC2", "TMED2", "PSMD3", "USP18", "LTF", "HLA-DQA1", "PLAC8", "HTR2B", "TMEM184B", "THE
GK1") c("ZFPM2", "EIF3H", "PDE8B", "PTX3", "RABEP2", "SORL1", "PCSK5", "ZMAT3") c("ANGPT1", "LAP
'TCN1", "HEBP1", "CEACAM6", "MARCKSL1", "COG5", "IDH1", "TFF1", "PLAU", "PKP2", "QPCT", "SNRPG"
'FXYD3", "SLC25A14", "SCNN1A", "CROT", "MEIS2", "GPD1L", "HIST1H1C", "IDH1", "RAP1GAP", "ERLIN2"
DBP", "ID1", "NR4A2", "S100A2", "PLAT", "IL1RN", "KRT14", "KRT17", "SLC43A3", "CDKN1A", "PEA15", "L
c("SFRP1", "MMP3", "FEZ1", "PTX3", "GADD45B") c("RBM47", "SAT1", "PHLDA2", "BCAP31", "LAP3", "H
T1", "FST", "TJP2", "ALDH1A3", "HIST1H2BD", "BMP2", "ID1", "NR4A2", "PLAT", "PEA15", "BSG", "LAMB
"RBPMS", "TFF3", "IGFBP2", "CDK6") c("VAMP8", "HBB", "GAD1", "SLCO1B3", "TP53TG1", "CEACAM6", "
A", "TRIM22", "CFH", "CXCL10") c("ZDHHC3", "PLK2", "FUCA1", "LAT2", "PIP", "LTF") c("ITGA3", "HOXB2
5", "ACADVL", "LXN") c("SULF1", "PITRM1", "HOXC6", "DUSP2", "DGKA", "FOXC1", "SSH3", "PSCA", "VCA
("IPO7", "SLC35F2", "TOMM34", "NR4A2", "CCDC86", "MRTO4", "TJP2", "EFHD2", "SAT1", "DDX10", "MI
PS30", "DUSP3", "ERCC1", "ZMAT3", "FEZ1") c("FERMT1", "SAT1", "B4GALT5", "MAL", "DHCR24", "PHLD
.", "POLR1D") c("YIF1A", "TBC1D8", "ATR", "PNPLA4", "FERMT1", "SAT1", "ANXA1", "FDPS", "CFB", "CIB1
'NR4A2", "CD55", "MRPS30", "SIK1", "KLHL21", "SOX9", "ADO", "LRRFIP1", "FAR2", "KCNK3", "KCNN4", "
A6", "CYP1B1", "ADD3", "DENND2D", "ASPH", "DAPK1", "TP53", "HMGCS1", "COL15A1", "HSD17B4", "SS

DRG3", "MTMR11", "GPD1L", "SORT1", "BBS1", "DENND2D", "GRN", "ARHGAP6", "TMEM106C", "ASPH"
IS1", "RAB15", "NRCAM", "CALCOCO1", "ARL6IP5", "FAM13A", "TRIM22", "PLOD2", "FXYD3", "SLC38A6"
FBP2") c("TPI1", "PPIC", "ADAMDEC1", "ANXA4", "CAPZA2", "ATP1B3", "DDIT4", "PIK3R3", "HDAC1", "ID
F2H5", "EI24", "TXN", "UCHL3", "CYB5R2", "HLA-DQB1", "GSTP1", "EMX2", "DNMT3A") c("LRRC1", "GLU

E", "SCNN1A", "GPD1L", "GTF2IRD1", "GCHFR", "FHIT", "SLC24A3", "CROT", "VAMP1", "ECHDC2", "TP53
P1A1", "ZC3H12A", "BACE2", "TCN1", "ADAM9", "UBE2H", "PGAM1", "ARPP19", "ASPN", "HSD17B2", "R
RP4", "RAB15", "TOX3", "CYB5R1", "PRRG1", "DDX17", "PKP2", "KRT23", "FCER1A", "ADAM8", "SMARCA
PFIBP2", "CDH1", "SFTPD", "NCK2", "TRIM38", "CACNA2D2") c("AMD1", "GAS2L1", "SLC5A6", "GLUD1",
'SCGB1D2", "ABCA8", "USP6NL", "ADAM28", "MYCN", "HIP1") c("HEBP2", "RARRES1", "STXBP2", "SLC38
NF144A", "ZC3H12A", "KRT23", "TCN1", "TRIP6", "FIS1", "TBC1D8", "SSR2", "UBE2E3", "ARMCX6", "GRB
', "TSPO", "MUC16", "PEA15", "TNFSF10", "SLC12A2", "NCOA4", "MVP", "CCNG1", "CDC42SE1", "ZFAND

", "BEX1", "USP6NL") c("HOPX", "RARRES1", "ZNF217", "PRR16", "GLUL", "S100A11", "TACSTD2", "NDFIF
'SHROOM2", "KIAA0232", "TBC1D9", "TP53TG1", "PIK3IP1", "SLC15A2", "FMO5", "ZIC1", "SLC25A13", "C
I", "STK38") c("CSTA", "HSD11B2", "STXBP2", "PPCS", "KRAS", "CD9", "TACSTD2", "MEIS1", "RNF43", "TN
, "DCK", "SEL1L3", "RGS4", "SLC25A6", "PSMB9") c("HEBP2", "HOPX", "RARRES1", "STXBP2", "PIK3R3", "
A1", "IER5", "KRT15", "STC1", "HSD17B4", "PON2", "TGFBI", "WFDC2", "ADIPOQ", "NLRP2", "PMM2", "A
IPRSS4", "DPYD", "GBP2", "IGBP1", "MUC1", "LPCAT1", "SAT1", "ZC3H12A", "AQP9", "CLDN4", "TBC1D8"
GEA3", "OVOL2", "GPX2", "COG5", "RAB5B", "ABCC5", "RHOD", "CCNG2", "HEXB", "CCNI", "ANXA3", "FU
MCCC1", "PIK3C2B", "GULP1", "RPS6KA5", "CFD", "CRISP3", "VAV3", "PGD", "VLDLR", "MICALL2", "IL20F
P1", "PNO1", "PDSS1", "TIAM1", "MPZL1", "PRNP", "PSTPIP2", "LIPG", "CEACAM1", "ASPN", "OSBPL10",
P1", "HYAL2", "CTSD", "GRB14", "CRK", "TNFAIP2", "ZNHIT1", "GSTT1", "SPOCK2", "BAALC", "ADAM28",
YP3A5", "MT1H", "ECM1", "SCNN1A", "RUFY2", "KCTD3", "ST6GALNAC2", "CD40", "JAG1") c("SOX9", "G
IB3", "CDS1", "JAK1", "SLC6A14", "VAMP8", "S100P", "HOPX", "TCN1", "TBC1D8", "BCL2L1", "PLAT", "RF
0S2", "MMP9", "LTF", "CCDC86", "KIF5B", "ITGB3BP", "C1QB", "TWIST1", "ZYX", "THBS1", "KRT17", "FLN

BD", "TPSB2", "GRB14", "PPP1R16B", "CHERP", "NUP85", "CTBP1", "DEFA4", "CRK", "CDC34", "RBP1", "
", "KLRK1", "TIMM13", "ANXA3", "NRIP3", "PNN", "PCDH9", "ADO", "PFKFB3", "TSC22D2", "KLHL21", "PI
D8", "TBC1D8", "RARRES1", "PRKAR2B", "RGS1", "HMOX1", "ADCY7", "RAPGEF5", "S100A10", "TUBD1",
K3", "HLA-F", "TFF3", "AREG", "EMP1", "LTF", "TACSTD2") c("DDIT4", "S100A9", "IRF9", "ZNF586", "FMO
, "PSTPIP2", "OLFM1", "BCL3", "EMP2", "CDC25B", "ADRB2", "KRT14", "ID1", "FCGR2B", "UQCRC1", "CRE
", "ZFPM2", "PRSS21", "FEZ1", "DUSP3", "SIX2", "ZMAT3", "SORL1") c("CLDN7", "GTF2IRD1", "DHCR24",
B", "ARHGAP22", "XPO7", "TJP2", "BPGM", "IL10RB", "GZMB", "RBM47", "AGER", "SERPINB1", "ALDH7A
, "PEA15", "THBD", "ARPC4", "ARHGDIB", "RBM47", "TJP2", "HS3ST1", "S100P", "DUSP6", "KYNU", "IL1R
ACAM6", "TMC5", "SLC25A46", "HIST1H1C", "RBM47", "HOPX", "CXCL5", "SAT1", "PHLDA2", "NRCAM", "
OBSL1", "TSC22D2", "VAMP1", "ANXA3") c("ID1", "RPA3", "LCK", "PEA15", "RBM47", "SPR", "TJP2", "DUS
47", "AIF1", "CA2", "SMPDL3A", "LAMP3", "CLEC7A", "PSTPIP2", "BMP2", "OLFM1", "THBD", "LIMA1", "
", "FEZ1", "WWOX", "PRSS21", "NUAK1", "EMP3") c("PSMB5", "IL13RA1", "RBM47", "HIST1H1C", "NFAT
NF586", "AKAP7") c("ZFP36L2", "GRIA2", "PELI1", "SOX9", "CCL8", "FAR2", "MRPS30", "CD55", "ANXA3",
2", "CDK6", "SQLE", "S100A2", "OLFM1", "QTRT1", "KRT17", "LSR", "MAPKAPK2", "OASL", "CYP1A1", "SN
3") c("RAF1", "DHCR24", "GGCT", "TGFA", "COMMD8", "SEMA3F", "NRCAM", "UBE2A", "SLC25A46", "CL
FZP586I1420", "CDC42SE1", "CXCL2", "XRCC5", "EIF4A3", "RAC1", "CCL20", "PWP1", "ASNSD1", "CDK6",
", "FUCA1", "EMP1", "LTF", "DNAJA3") c("ALAS2", "HBB", "TCN1", "CEACAM6", "ALOX5", "PSCA", "S100A
L6", "IL1RN", "RBP4", "PCSK1", "DAXX", "S100P", "S100A2", "ME2", "GARS", "KRT16", "G0S2", "KRT14",
1GDS1", "ITGAE", "BAALC", "DVL3", "PRR11", "IMPDH2", "MCM2", "DEFA4", "MCM6", "BMP4", "BPHL",
D", "MCM7", "IDH1", "MT1H", "SH3YL1", "GMPR2") c("IFRD1", "EIF1", "IER2", "TRIM16", "AVPI1", "ATF3
"PLK2") c("SLCO1B3", "BFAR", "KLK10", "MARCKS") c("TEX2", "CRIP2", "DIRAS3", "ENPP1", "EFHD1") c("
TP2", "TULP3", "KCNK3", "GRIA2") c("CALR", "ITPKB", "ANXA6", "IMPDH2", "MBOAT7", "ADORA3", "RBI
OP1", "KLHL21", "DNMBP", "MCTP2", "APOD", "EREG") c("GSTT1", "PPP1R9A", "RABAC1", "ITGAE", "PD
K2", "CA2", "ASPN", "TACSTD2") c("TBC1D8", "RRM2", "S100A9", "TP53TG1", "HOXB2", "PSCA", "ALOX5
", "PQBP1", "USP7", "ADD3") c("TACSTD2", "SLC10A3", "FLNA", "RALGDS", "DNASE1L1", "RFTN1", "CDH
", "ABCA8", "BMI1", "SLC19A2", "RHBDF1", "DDX17", "ZSCAN18", "INHBB", "APOD", "ADO", "MEAF6", "

", "FERMT1", "UGT2B4", "SET", "HOXB2", "LYPD1", "SDC1", "ANXA4", "VAMP8", "UQCRQ", "S100A11", "

C2A10", "RAP1GAP", "SLC24A3", "GSTM3", "GOLGA7", "SH3GLB2", "SLC35A3", "MKNK2", "HIST1H1C", "
TGB4", "CLDN4", "PRDX3", "FOXC1", "CTNND1", "TNFRSF10B", "KCNJ16", "GLUD1", "GPR137B", "FOXA1
A2", "TBC1D8", "XPO7", "TIMM9", "CERK", "NDUFA3", "GALNT1", "BEX1") c("TACSTD2", "ALDH3B2", "DU
", "BAX", "JAG1", "S100A11", "CDC42SE1", "CNOT7", "STAT6", "SLC35A2", "LIMK2", "TSPAN15", "STXBP2
19", "PRKAR2B") c("ZIC1", "CDS1", "TMX4", "SPRY2", "RUNX1", "CXADR", "HOXA5", "USP46", "DAAM1",
CST6", "COBLL1", "SRD5A1", "CXADR", "CD9", "TNFSF10", "SAT1", "MEIS1", "EPHA4", "PDCD6", "BBOX1"
P2", "STK38L", "PSMD5", "TMPRSS2", "S100A11", "RARRES1", "PRRG1", "MVP", "OLFM4", "PLAC8", "PIK
CPA3", "ADH1C", "CTNND1", "PON2", "NBR1", "NLRP2", "IRF9", "TSPAN15", "HSD17B4", "TGFBI", "RAB1
217", "YWHAE", "STXBP2", "HOPX", "TTC9", "NAMPT", "USP6NL", "GBP2", "CLCA2", "IRF6", "TBC1D8", "
"GABRP", "CPA3", "SLC9A3R1", "ADH1C", "CCNG2", "TAC1", "MBOAT7", "EYA1", "MAN2A1", "ANXA3", "
"LYZ", "LTF", "ZC3H12A", "RALGDS", "SMPDL3A", "CCL18", "SAT1", "ISG20", "S100A2", "JUNB", "LAMC2"
SEMA4C", "ZNF211", "ADAM28", "ARHGAP6", "SORT1", "TNS1", "TPSB2", "TPSAB1", "CRK", "HIST1H2BD
J16", "CCL4", "MARCKS", "CLCA2", "ANXA4", "GABRE", "FMO1", "SPP1") c("FAM129A", "TRIM22", "CEAC
SLC7A8", "DST", "GSTM3", "HSPB8", "TSC22D2", "ESR1", "SLC2A10", "SH3GLB2", "LGR4", "SCNN1A", "ZN
PIK3R3", "EPS8L2", "HSD17B2", "PLAC8", "CA2", "ADH1C", "PCCB", "HEXIM1", "CD9", "MYO5C", "SERPIN
", "ADAM8", "NOLC1", "DDX3X", "SLC25A13", "SLC25A28", "DAXX", "TWIST1", "CHIC2", "NUSAP1", "RBP
HB2", "CD55", "TRIM16", "FOXO1", "GRIA2", "MMP3", "CYP4F11", "INHBB", "EREG", "PVR", "OBSL1", "L

, "PRPF4", "NQO2", "CXCL5", "BID", "AQP9", "NETO2", "HOMER1", "RFC5", "CUTC", "ALDH1A3", "NRIP1'
', "TSEN2", "TIMM9", "HGD", "ITFG1", "SUB1", "KRT17", "LRRC1", "MIA", "SDC1", "DNMT3A") c("MCCC1
TM4SF1", "MTUS1", "ALDH3B1", "HCCS", "ECM1", "CRNN", "MT1H", "SCNN1A", "JAG1", "ECHDC3", "H1F
'LRP8", "SLC12A8", "SCML1", "NDRG2", "PLAC8", "GTF2I", "RBM3", "DNMT3A", "SPP1") c("FAM102A", "I
GB3", "BACH1", "TACSTD2", "CCDC86", "VNN1", "RANBP1", "RRAGC", "TLR4", "BAG3", "USP18", "EIF4A3
3", "FBXO21", "TNS1", "NUDT1", "ASPH", "SKIV2L", "PPP1R13B") c("SIX2", "CCNL2", "MED13", "ZFPM2",
)", "ITM2B", "CARHSP1", "HELLS", "GINS2", "DNAJA3", "DNAJB6", "FAM129A", "AKR1B10", "LIMK2", "EG
, "MYC", "PFKFB3", "ZFP36L2", "OBSL1", "SOX9", "CCL8", "WEE1") c("PCSK1", "PUF60", "BACH1", "TACST
HBS1", "BCL2A1", "UBE2H", "TCN1", "LYPD1", "PLAC8", "ALDH1A3", "ITGA4", "LYZ", "ME2", "PLAT", "CYI
K", "MARCKSL1", "INPP4B", "IFITM1", "MMP1", "SPP1") c("NDUFB2", "FAM129A", "TRIM22", "SLC39A6'
)0A10", "CYP3A5", "SH3BP4", "ST6GALNAC2", "IKBKE", "ESR1", "GMPR2", "PPL", "NOX4", "TM4SF1", "CI
, "NKX3-1", "TMEM183A", "ACAA1", "TGFA", "CXCL5", "CASP7", "NRCAM", "SERPINB4", "DHCR24", "HOI
"IL7R", "XPO1", "ASPH", "CCL5", "RAC2") c("BAG1", "CYBB", "SOD2", "BSG", "LIMA1", "HLA-DQA1", "RH
21", "EREG", "LSM12", "SPAG1", "ZSCAN18", "NR4A1", "IRF6", "NRIP3", "MMP3", "CACNA2D2", "TMEM
"TSPAN31", "MSN", "SLC16A1", "ALDH1A3", "NR4A2", "IL1B", "STOM", "FERMT1", "AZIN1", "TP53TG1",
ELLS", "NAPA", "SDC3", "AKTIP", "EHBP1", "AKR1B10", "SLCO1B3", "NME1", "SLC39A6", "H2AFZ", "LMAI

IK1", "KLHL21", "DUSP1", "NR4A2", "FOSB", "SV2A", "ANXA3", "NRIP3", "VAMP1", "PCDH9", "CFI", "GRIA
, "SMARCA1", "GLRB", "TKTL1") c("CSRP2", "SIK1", "CHD1", "DUSP1", "CCL2", "TREM1", "NR4A2", "FOS",
"XPO1", "ZNF586", "SCD", "TBC1D9") c("POLR2J", "HOPX", "CAV2", "TM9SF2", "DCTN4", "LSM6", "PDLIN
AIM", "RBP1", "SLC39A4", "ACPP", "CDH1", "TGFA", "COBLL1", "ARNTL2", "NDRG2", "HMGN3", "SRD5A
NDUFB3", "TUBB6", "POLR2I", "DPYD", "NDFIP1", "MUC4", "HAPLN1", "ACTR2", "ABHD2") c("HSPB8", "C
", "PEA15", "PRMT1", "EDEM1", "FPR1", "ACTN1", "PLEKHO1", "S100A7", "TMEM158", "BCL2A1", "AKIR
", "CRK", "DEFA4", "TPSB2", "TPSAB1", "ALOX5AP", "FCGR2A", "FGF9", "QRSL1", "HSPD1", "CNR1", "ARH
RPS2", "BCL2A1", "LRP4", "RBP1", "SERPINB1", "RRS1", "PLS1", "CAP1", "LYPD1", "RBM47", "FST", "CLIC
3P1", "PSMB9", "PDLIM2", "TPSAB1", "TPSB2", "PHKA1", "PPP1R9A", "EFHC1", "MTMR11", "ITGAE") c("
IFI30", "RBP4", "OLFM1", "IL1R1", "HLA-DQA1", "MMP12", "CA2", "ALDH1A3", "NUSAP1", "KLK6", "SPH
TX3", "PCSK5", "USP9X", "RBM26") c("DHRS3", "HIST1H1C", "RAF1", "SERPINB4", "HOPX", "NRCAM", "C
', "CD55") c("PAF1", "IL7R", "MRPS2", "AGPAT2", "HSD17B2", "ALDH1A3", "NUSAP1", "KLK6", "SPHK1", "
FAM107A", "P2RY14", "RAB15", "NRCAM", "TACSTD2") c("PACSIN3", "TACSTD2") c("ADAR", "SERPINB1"
"RFTN1", "TSFM", "LIPG", "LTF", "G0S2", "MMP12", "TRAPPC3", "CCL20", "HNRNPAB", "HTR2B", "GZME
, "ITGA4", "HSD17B2", "SLC39A14", "S100A7", "CCL18", "MSN", "ALDH1A3", "DGAT1", "MLF1", "CXCL1",
IMP3", "ADD3", "PTX3", "ZNF83", "HLTF", "SORL1") c("COX7A2", "SERPINB4", "SERPINB6", "CHMP2A", "I
'GPR19", "CYP4F11", "NRIP3", "GRIA2", "PELI1", "REG3A", "CFI", "KLHL21", "PCDH9") c("HTR2B", "TPPP3

9" "V100" "V101" "V102" "V103" "V104" "V105" "V106" "V107" "V108" "V109" "V110" "V111" "V112"
S2", "ZNF423", "LXN", "SUB1", "TMEM176B") c("AGL", "TMPRSS2", "GPRC5B", "GADD45B", "DUSP1", "A
N2", "DAXX", "DDX10", "ECT2", "EPHA2", "EXOC5", "F2RL1", "FGFR3", "FLNA", "FOSL1", "FSCN1", "G0S2
AD1", "GPC1", "HOPX", "IL1R2", "MAL", "PHLDA2", "SERPINA5", "TGFA", "TMC5", "TSEN2", "TSPAN6") c(
", "PTP4A1", "PUF60", "PYGM", "RBP1", "RRS1", "SERPINB1", "SERPINB2", "STAT3", "TCN1", "TUBB6", "
3D", "SAT1", "ASPN", "SIGLEC15", "CARD10", "UGCG", "ARFGAP3", "PBLD", "MVP", "STOM", "S100P", "K
3", "NKX3-1", "SLC25A46", "TSPAN6", "MGST3", "SAT1", "COX7A2", "SDHC", "COMMD8", "DHRS3", "ERE
, "TBC1D8", "NETO2", "RGS16", "RRAGA", "SNRPE") c("TMED2", "DBI", "GALNT1", "SLC25A46", "KRT6B"
, "VAV3", "GCHFR", "AREG", "CRABP2") c("PDLIM5", "ANXA3", "SIK1", "YPEL5", "CFI", "FOXA1", "SOX9",
SR", "BSG", "CHORDC1", "LRP8", "FEN1", "THBS1", "MS4A6A", "BCL2A1", "CCR7", "GZMA", "S100P", "SL
OPX", "FERMT1", "SERPINA5", "DHRS3", "RANBP9", "HIST1H1C") c("FXYD3", "TMED5", "ATF3", "P2RY14
3", "FEN1", "IL7R", "BCL2A1", "PYGM", "GZMB", "TCN1", "MMP1", "AQP3", "PDE4B", "FGFBP1", "TACST
S100A9", "CDK6", "TCN1", "PSME2") c("SUB1", "ATOX1", "CLU") c("IL15", "SCNN1A", "SQSTM1", "COBL
", "EDEM1", "GZMH", "VAMP8", "S100A9") c("TSKU", "LRIG1", "LPAR1", "TMEM9B", "VPS28", "ARL6IP5
M1", "MID1IP1", "EAPP") c("ANXA4", "EPS8L3", "DDX42", "PLEK2", "DDIT4", "QPCT", "CYP3A5", "SLC25
PZL1", "TIAM1", "NOLC1", "CIB1", "LAMC2", "CDCP1", "RFTN1", "SLC6A14", "SLC2A6", "F2RL1", "DDX21
A2", "FHIT", "TMPRSS3", "MCTS1", "RBM47", "HOPX", "ARMCX6", "SLC11A2") c("SAMD4A", "ABHD2", "
", "MAL", "LAMC2", "ACOT7", "SMS", "RGS4", "FMO1", "SLC6A14", "TSPAN3", "TFRC", "DHCR24", "ARE
RRAS2", "CFI", "PELI1", "ATF3", "FOSB") c("IPO7", "NR4A2", "PPBP", "TJP2", "EFHD2", "SAT1", "IL7R", "T
R2", "PEG3", "CHSY1", "IL7R", "USP7", "MS4A1", "HLA-DOB", "RAC2", "BCAR3", "P2RX4", "SHC1", "RIF1"

, "IGBP1", "RNASET2", "ITPKB", "SEMA4C", "QARS", "ALDH7A1", "HIST1H2BD", "ANXA6", "AP1G2", "ATP
, "MNAT1", "REXO2", "CKS2", "GMPR2", "SERPINA5", "ADM", "ATF3", "ABHD2", "SYNJ2", "NPL", "SOCS1
H1", "ID1", "PTPRK", "MMP7", "HEBP1", "MARCKSL1", "TACSTD2", "ALOX5", "EPS8L3", "LTBR", "MMP1
D1", "HK2", "PRDX1", "ANXA4", "CAPZA2", "PEX11A", "LGALS3", "S100A11", "LTBR", "BBS1", "CDK6", "

TG1", "MKNK2", "SH3GLB2", "GSTM3", "HUWE1", "DST", "ST6GALNAC2", "ZFHX3", "CDKN1B", "MTUS1
GS16", "DDX21", "FABP6", "RBM47", "PLAT", "TPPP3", "TRAF3IP2", "IL7R", "TNFAIP6", "ADAM10", "CXC
1", "PDK4", "UIMC1", "PPARG", "RAB1A", "ESM1") c("CSTB", "CLDN8", "ZFP36L2", "GRIA2", "PIM1", "A
"BAMBI", "ID1", "LTF", "TFF3", "STXBP6", "LTBR", "INSIG1", "FEZ1", "JUNB", "CYB5R1", "SLC9A1", "CDKN
A1", "PIK3R3", "PRR16", "S100A11", "MAPK1", "PLCB3", "IFRD2", "LY6D", "HSPA6", "RNF144A", "MNAT
2", "UBE2Q1", "PDZK1IP1", "TMEM165", "LEF1", "CR2", "PDCD6", "KCNJ15", "CDH1") c("HEBP2", "RARR
6", "MRPS16", "FGB", "ALDH3B2", "MBNL2", "UBQLN2", "SLC25A13", "CLDN4", "DIO1", "PERP", "TBC1D

1", "TNFSF10", "STK38L", "TMPRSS4", "SLC12A2", "PRRG1", "PPP2R5E", "MUC1", "ALDH3B2", "LPCAT1
DS1", "TBC1D8", "RUNX1", "ZBTB38", "LSM5", "IKZF1", "HMG20B", "NPL", "SCML1", "WIPF2", "FAM193
FSF10", "TMPRSS4", "MVP", "CORO1B", "ZMYND8", "PIK3IP1", "TRAF5", "IGBP1", "ALDH1L1", "SLC6A14
S100A11", "LXN", "TACSTD2", "SLC2A3", "NDFIP1", "TMPRSS2", "STK38L", "OLFM4", "MVP", "PRRG1", "
DAM8", "GABRE", "ICAM2", "NNMT", "CTNND1", "PTGS2", "PER1", "WIPI1", "MYCN", "SFTPD") c("ID3",
, "TPK1", "ABHD2", "PLAT", "NAMPT", "PDZD8", "LPXN", "USP6NL") c("HEBP2", "FBP1", "CHST15", "GAE
T8", "AKAP12", "GCNT3", "PKP2", "ARID4B", "MET", "CTSH", "GMPR2", "NRGN", "DPYSL2", "ITPR1", "TC
A", "GPATCH8", "CEACAM7", "GTF2H5", "MS4A1", "TSEN2", "PIR", "OSBPL3", "LRBA", "FOXJ3", "RNF12
"LTF", "SQLE", "SLC2A6", "KRT14", "ISG20", "ME2", "HMGA1", "GRWD1", "CHI3L2", "SLC35F2", "DKC1",
"CAMK2N1", "RARA", "F12", "TSPYL4") c("MAN1A1", "MRPS30", "SFRP1", "PSPC1", "ZMAT3", "PTX3", "
RPEL1", "MRPS30", "CCL8", "KLHL21", "TIMM13", "CFI", "ADO", "OBSL1", "GRIA2", "KCNN4", "CREB5", "
K", "THOC7", "RTP4", "GALC", "STK38L", "PLS1", "TACSTD2") c("MS4A4A", "UQCRH", "HK1", "FADS1", "T
A", "MYO5A", "KRT14", "GZMB", "KIF14", "CD14", "RGS1", "FSCN1", "ALDH1A3", "DKC1", "HNRNPM", "

PLSCR4", "PTHLH", "ASPH", "GPD1L", "ADRB2", "FGF9", "PSMB9") c("CES3", "EMP3", "TARP", "RABEP2",
DLIM5", "VAMP1", "RRAS2", "TRIM16") c("CLIC1", "ALDH7A1", "KIF5B", "ZYX", "HIST1H2BD", "GZMB", "A
"FGFR3", "TACSTD2", "HOPX", "VPS37B", "RIPK4", "CCDC47", "ARPP19", "MUC1", "SLC37A4", "SLC7A1",
2") c("BST2", "SUB1", "AKR1C1", "BBOX1", "TSKU", "VPS28") c("SCNN1A", "B4GALT5", "SHROOM2", "ECI
EB3", "LILRB2", "CCR1", "XPO7", "LYN", "TJP2", "CCL18", "BPGM", "IL10RB", "TWIST1", "VNN1", "GZMB",
"SERPINB6", "CHMP1B", "HIST1H1C", "RBM47", "KLK3", "ANP32E", "ANGPT1", "NKX3-1", "PLS1", "HOPX
.1", "KLK3", "METTL5", "GNAQ", "EPB41L1", "TNFRSF21", "KIF5B", "MID1IP1", "PLS1", "DDX21", "SERPIN
1", "CXCL2", "MMP12", "SAT1", "SLC35F2", "DDX3X", "IER2", "ID3", "PLAC8", "IRF1", "JUNB", "ME2", "CI
'ERBB3", "LPCAT4", "SMARCD2") c("GMPR2", "SLC25A46", "PDE1A", "ATF3", "TRIM22", "PLOD2", "ITGB
SP6", "ATP6V1D", "CXCL2", "SAT1", "DDX3X", "SERPINB1", "IRF1", "ALDH1A3", "PUF60", "TNFAIP6", "ZC3
TLR4", "ANXA2", "VNN1", "TRAF3IP2", "FGFR3", "ARL14", "KRT17", "MLF1", "CLDN4", "ISG20", "HK2", "R
C1", "COX7A2", "CEACAM6", "PON1", "DHCR24", "PLS1") c("FAM107A", "CXCL10", "PDE1A", "CA2", "LAM
"OBSL1", "KLHL21") c("HIST1H2BD", "KLK7", "RBM47", "LAMB3", "CTNND1", "BMP2", "LIMA1", "FST", "
MPDL3A", "LCK", "HTR2B", "S100P", "SLC39A1", "CLDN7", "TLR4", "OSBPL10", "AQP3", "LYZ", "ASPN", "R
DN7", "OIP5", "TSEN2") c("FAT4", "GMPR2", "CXCL10", "NRCAM", "MNAT1", "UBE2A", "SLC25A46", "PD
, "QTRT1", "ARPC3", "HIST1H2BD", "LCK", "HTR2B", "HNRNPH2", "PUF60", "AQP3", "ASPN", "MED28", "I
A9") c("TMEM176B", "ITIH5", "SPOCK1", "TMOD1") c("COBLL1", "FOXA1", "RHCG", "MAPK6", "RGS10", "
"S100A7", "AQP3", "LAMP3", "PLSCR3", "CCL18", "LIPG", "FLNA", "MAPKAPK2", "SLC2A9", "OASL", "SLC
"IGBP1", "ASPH", "CRK") c("EIF1", "RABEP2", "FEZ1", "FGF9", "GADD45B", "SFRP1", "PRSS21", "PCSK5",
", "CD55", "GRIA2", "GADD45B", "TIPARP", "FAR2", "PFKFB3", "REG3A", "FUT8", "ANXA3", "GPAA1", "AI
LST1", "CHST15", "GLIPR1", "ITIH2", "EIF3A", "MUC4", "AREG", "RANBP9", "EPHA2", "FZD6", "CA2", "FA
P1", "RNASET2", "ZFHX4", "ADAM28", "PPP1R13B", "IFT57") c("RABEP2", "GGA1", "PDE8B") c("SAT1", "H
LIM2", "GRB14", "RARA", "IGBP1", "RAB33A", "DFFB") c("ZNF83", "DNMBP") c("NDUFB8", "SLC35F5", "E
", "VAMP8", "TCN1") c("LXN", "BST2", "CRIP2", "GEM", "AKR1C1", "FOSL1") c("MAOA", "EPHA2", "TAX1
3", "ALOX5AP", "JUNB", "TMEM158", "S100A7", "MMP12", "ALOX5", "LTF", "KCTD5", "SAT1", "PBLD", "S
TRIM16", "RUFY2", "PXDN", "CD55", "IRF6", "CAB39", "DVL3", "PNN", "IDS", "PPP3CA", "USP7", "GFOD1

KRT17", "RIPK2", "PAPSS1", "CAPNS1", "LRRC1", "GLUD1", "NBR1", "LGALS3", "TXNDC9", "MIA", "DHCR

'SOX13", "STAT6", "NBR1", "ZNF586", "IDH1", "GSTM1", "S100A10", "ECHDC2", "IKBKE", "ZNF611", "SCN
.", "TRIM16", "BPGM", "CFI", "ID1", "IDS", "XPO7", "ATP1A1", "LRMP", "GNLY", "F2RL1", "EHD4", "FST", "
JSP22", "STK38L", "CXADR", "TNFSF10", "SAT1", "SLC12A2", "S100A11", "RARRES1", "TCN1", "PRRG1", "I
.", "TTC9", "MTMR11", "EXOC7", "STK38", "PDZK1IP1", "HIGD2A", "VAMP5", "TP63", "HSPA2", "SAMSN1
"MAL", "UQCRQ", "S100A11", "TP53TG1", "SLC39A4", "NDRG2", "SLC35A3", "OLFM4", "FBXO2", "MAT2
', "ALDH6A1", "RBP1", "TTR", "SLC39A4", "NDRG2", "HMGN3", "CORO1B", "RGS1", "MVP", "ZMYND8", "
3R3", "MUC1", "VPS35", "FGB", "STXBP2", "HOPX", "TTC9", "USP6NL", "F5", "MS4A4A", "TBC1D8", "FM
.5", "PMM2", "STK10", "PTGS2", "KRT15", "NNMT", "MUC5B", "TFF3") c("RARRES2", "PCGF3", "DHRS1",
'PKN2", "NEK7", "PPP1R16B") c("FBP1", "TACSTD2", "CST6", "DUSP1", "HEBP2", "GPRC5A", "DHRS3", "G
'FUT8", "GMPR2", "PHF11", "DCHS1", "PDLIM1", "ACSS3", "LRMP", "FAM107A", "SLPI", "WARS", "TMPR
', "EPHA2", "IRF6", "PLAT", "IER5", "SLC35F2", "TCN1", "ASPN", "AQP3", "MVP", "FOSL1", "TACSTD2", "C
)") c("SFRP1", "NUAK1", "DUSP3", "ZMAT3", "ZNF83", "MAN1A1", "FYCO1", "PRSS21", "MED13", "DNMI
CAM6", "SLC25A46", "AHCYL1", "KDELR2", "NDUFA4", "LIMK2", "DNAJB6", "EGF", "ERBB3", "PI3", "AKR1
F586", "SLC24A3", "ARMCX6", "MEIS2") c("ATF3", "KCNN4", "YPEL5", "ADO", "OBSL1", "FOSB", "ANXA3"
IB2", "SRC") c("WSB2", "TMPRSS4", "SAT1", "TGFA", "TCN1", "SLCO4A1", "TUBB6", "PIK3R3", "SLC6A14"
4", "IL1RN", "ELK3", "ID3", "ALOX5", "NRBF2", "KLK6", "CD37", "CHI3L2", "PCSK1", "SLC39A14", "LYN", "
YPLA1") c("PHKA1", "SPOCK2", "CD7", "PIGB", "RNASET2", "CDS2", "POLR3K", "SHMT2", "ECHDC3", "FB

", "CRABP2", "CLIC3", "TALDO1", "CADM1", "CRCT1", "EPHB2", "ALDH3B1", "ECM1", "GCNT3", "CRNN",
F0", "ZNF586", "GSTM3", "PHGDH", "CAST", "AREG", "PPL", "SH3BP4", "NAV2", "GPD1L", "PDZD2") c("KL
KLK6", "S100A10", "CLPX", "FGFR3", "MPZL2", "TMPRSS4", "CCDC47", "HOPX", "LAMB3", "KYNU", "HMC
", "OSBPL10", "MAPKAPK2", "CYP1A1", "TUBA1A", "FGFR3", "RHOBTB3", "TCN1", "LYZ", "TJP2", "MELK"
"FEZ1", "SORL1", "ZFP36L2", "TRIM13", "SFRP1") c("TIMELESS", "TMEM50A", "CREG1", "BCAP31", "HOF
F") c("SLC25A4", "LGALS3", "IRAK1", "CLDN16", "RAPGEF5", "REC8", "MCM6", "ACVR1B", "TRIP13", "GR
TD2", "ATP6V1D", "BAG3", "EIF4A3", "ARPP19", "TCN1", "TJP2", "EDEM1", "FST", "NDUFS2", "RRS1", "HI
P1A1", "IRF6", "MAP3K8", "S100A2", "S100A7", "TACSTD2", "NR4A2", "EPHA2", "SMPDL3A", "TNFRSF21
, "EMX2", "SPRR2B", "ERBB3", "HOPX", "SPRR1B", "PI3", "KRT6B", "E2F2") c("ANXA4", "R3HDM2", "ARE
D40") c("ATF3", "SYNPO", "OBSL1", "REG3A", "YPEL5", "ANXA3", "CD55", "NR4A2", "SIK1", "MYC", "GRIA
PX") NRCAM c("ACP6", "TMEM183A", "RBPMS", "ACAA1", "PIP", "CXCL5", "ASPN") c("KLK10", "MSN", "L
OBTB3", "TYROBP", "LAMP3", "ALOX5", "PRKCI", "S100P", "MMP9", "BMP2", "ALOX5AP", "CDK6", "LTF"
214", "SHC1", "PXDN", "GRIA2", "APOD") c("CDIPT", "PDE6D", "PDLIM2", "ADAM28", "IFT57", "ALOX5AP
"BID", "MYO1B", "CLEC7A", "PPT1", "ST14", "MMP12", "SERPINB1", "XPNPEP1", "SPATS2L", "NDUFAF3"
N1", "EMX2", "GTF2H5", "FKBP1A", "TXN", "ARPC2", "KLK3") c("FBXO3", "TNNT1", "RFC2", "SLC1A3", "PE

A2", "OVGP1") c("BAG1", "BSG", "LIMA1", "PRKCI", "BMP2", "CDK6", "GPR183", "SAT1", "TM9SF2", "EIF4
, "FOSB", "ARG2", "SV2A", "ANXA3", "CDS1", "COL4A3BP", "NRIP3", "PCDH9", "FBXO11", "GRIA2", "RGS2
M1", "PLAC8", "RCAN1", "ADM", "MYCBP", "DIO1", "TEK", "NDUFS7", "TSPYL5", "CA2", "SEC61G", "SERP
1", "NDUFB3", "AKTIP", "ASRGL1", "ECHS1", "PAX5", "MPDU1", "XPO1", "SLC6A15", "CAPZB", "ACTR2", "
LC", "CD1E", "SAT1", "TM9SF2", "CDK7", "DUSP1", "AREG", "SLPI", "DHRS3", "ACADM", "KRT19", "KLF6"
IN1", "KYNU", "FLNA", "FOSL1", "CD14", "RRS1", "LYN", "PLS1", "CAP1", "ITGB3BP", "LYPD1", "PLAC8", "
GAP6", "IARS2", "ADRB2") c("TARP", "EMP3", "PTX3", "SORL1", "ADD3", "ESF1", "FGF9", "SFRP1") c("GA
L", "SERPINB2", "PGAM1", "DDX3X", "KLK7", "PLAT", "SLC12A8", "DDX21", "METTL5", "POLR1C", "EPHA2
ADD3", "FEZ1", "SFRP1") c("UQCRQ", "ANP32E", "ACTR2", "NRCAM", "TGFA") c("MAN2B2", "KLF9", "NR
K1", "RRM2", "CD14", "S100A7", "NEK7", "CLEC7A", "RHOBTB3", "KLK7", "CYBB", "GZMA", "AQP3", "CD
XCL5", "GDE1", "CASP7", "LAP3") c("CKS2", "FXYD3", "ADM", "CLDN10", "MARCO", "CXCL10", "CA2", "FT
HNRNPL", "SERPINB2", "KLK7", "AQP3", "ATAD2", "PUF60", "ARPC5", "TRAF3IP2", "KMO", "MSN", "VPS
", "VAMP8") character(0) c("LYPD1", "CHIC2", "GSTZ1", "SERPINB1", "SPINT1", "RAB11FIP1", "CFI", "S100
3", "TCN1", "AKIRIN1", "RRS1", "ARAP2", "CHD1", "IER2", "KYNU", "NUSAP1", "ALOX5", "KLK7", "S100A2
, "LTF", "STOM", "F2RL1", "S100P", "PLAC8", "MMP1", "TACSTD2", "TYMP", "PPP2R1B", "BMP2", "FEN1"
MGST3", "CXCL5", "COX7B", "HIST1H1C", "GRAMD1C", "TMPRSS3", "MAP2K1", "LAP3", "SEMA3F", "HOI
", "HSD17B2", "MSN", "ALDH1A3", "HIST1H2AE", "RBP1", "DGAT1", "MLF1", "HNRNPL", "PUF60", "MMI

'V113" "V114" "V115" "V116" "V117" "V118" "V119" "V120" "V121" "V122" "V123" "V124" "V125" "V1
NXA3", "SLC15A2", "MMP10", "SERPINB2", "VCAM1", "CLCA4", "PEX11A", "HIST1H1C") c("MARCKS", "K
", "GARS", "GOT1", "GOT2", "GZMA", "GZMB", "HBEGF", "HIP1R", "ID1", "IFI30", "ITGA3", "ITGA4", "KCT
"ARHGEF3", "ARL6IP5", "CDR2L", "CLDN10", "COL8A1", "CYP3A5", "FTH1", "LAMB3", "PTS", "SAMD4A",
VPS37B", "ZYX") c("CACYBP", "CLSTN1", "COX6B1", "CYTIP", "GPSM2", "HSPA4L", "IGF2BP3", "LARS2", "C
RT14", "F2RL1", "BCL2A1", "CAP1", "ARHGDIB", "RBP4", "BAMBI", "HK2", "ALOX5", "HDAC1", "TACSTD2
B3", "SEMA3F", "PRR15L", "BBX", "CEACAM6", "HIST1H1C", "SERPINB6", "RBM47", "NRCAM", "ANP32E
, "COX7A2", "PBLD", "COMMD8", "LIMK2", "ERBB3", "CEACAM6", "DNAJA3", "TBC1D8", "ANP32E", "HS
"FAR2") c("KMO", "COL4A3BP", "G3BP1", "PUF60", "TFDP1", "MED28", "CLINT1", "TMED2", "HTR2B", "
C39A14", "G0S2", "S100A7", "GZMB", "TCN1", "CA2", "MMP12", "MMP1", "CCR1", "FGFR3", "AQP3", "I
", "LAMB3", "SOCS1", "PRDX4", "CXCL10", "CA2", "EEF1E1", "ANXA4", "COL6A3", "SERPINA5", "SLC16A2
D2", "KCTD5", "SLC10A3", "ZBTB43", "ARPP19", "GPR183", "RALGDS") c("IGF2BP3", "CYTIP", "MN1", "GI
L1", "MT1E", "GAD1", "SERPINB2", "DYNLT1", "TGIF1", "IGFBP2", "SLC15A2", "NETO2", "HIST1H1C", "TP
", "AKR1C1", "SUB1", "TMEM176B", "LXN", "CLU") c("SCGB2A2", "SULF1", "HIST1H1C", "CTSH", "DYNLT1
A6") c("ATXN7", "PPIB", "EPHA4", "SLC39A6", "TRIM22", "CYB5R2", "SLC38A10", "KLK3") c("ANXA4", "CT
, "OLFM1", "CXCL1", "CXCL2", "ITGA3", "DUSP7", "BMP2", "LCP1", "HK2", "TMEM158", "CCL20", "ATP1
IL6", "URB1", "RGS4", "SCARB1", "ARL6IP5", "PLOD2", "KLF9", "REXO2", "ATF3", "ADM", "TRIM22", "RA
G", "ENO1", "HIF1A", "NDRG1", "PPP1R9A", "ST14", "SCNN1A", "MCTS1", "HK2", "LTA4H", "CNDP2", "IN
IAM1", "RBP1", "NOLC1", "CDCP1", "DDX21", "CXCL2", "ST14", "BMP2", "HIST1H2BD", "BCR", "CCL20",
, "GTF2I") c("HK2", "BAMBI", "LY96", "HDAC1", "SLC2A6", "FGFR3", "ID3", "BPGM", "ID1", "IRF6", "LGAL

1A1", "CRK", "PBXIP1", "HSD17B4", "DIXDC1", "CPD", "ZNHIT1", "PTHLH", "GSTT1", "TSPYL4", "PGD", "N
", "RAB32", "CLDN10", "RGS4", "OLFML2A", "IGFBP2") c("LRRC1", "SLC9A3R1", "BAMBI", "PEX11A", "ND
2", "PEA15", "MT1M", "COG5", "STOM", "KCNJ16", "CDK6", "TP53TG1", "DAZAP2", "ALDH1A3", "TFF1",
CXR", "LYPD1", "SSH3", "CTSE", "SDC1", "SLC25A13", "HSPE1", "GSTT1", "CD58", "BAD", "PERP", "CTNN

", "NAV2", "SLC2A10", "CAST", "CRABP2", "FXYD3", "SH3BP4", "VAV3", "GMPR2", "ARMCX6", "SASH1", "

POLD1", "TOX3", "JUNB", "STK38L", "CD37", "YPEL5", "DDX17", "ANXA3", "FBXO11", "FUT8", "CDS1", "A
1A", "KLHL21", "HELLS", "IFRD2", "IER5", "MAP3K8", "LMNA", "SERPINB5", "RALGDS", "SFN", "SLC35F2"
1", "S100A6", "MEOX2", "SSR2", "NLRP2", "CISD1", "SCML1", "MAN2A1", "WIPF2", "HGSNAT", "SPP1", "
ES1", "SLC5A6", "SLC38A1", "DSP", "S100A11", "SDC4", "S100A10", "ASNS", "SEMA4D", "PPP2R5E", "SSH
8", "KLHL24", "PKIA", "TGFA", "DDX21", "SCML1", "PTX3", "BCL2L1", "AREG", "RAC2", "TNC", "RHOA", "I

, "SAT1", "MBNL2", "ZC3H12A", "TCN1", "TBC1D8", "TGFA", "PLAT", "SCML1", "WIPF2", "NAMPT", "PDZ
B", "SCD", "PEG10", "ZNF331", "SMARCA2", "SLC39A4", "ARL5A") c("HOPX", "SLC9A3R1", "ANKRD27", "
", "LPCAT1", "EHF", "SAT1", "S100P", "SLC25A13", "UBE2H", "COBLL1", "PAX5", "NKX3-1", "TBC1D8", "P
FGB", "FMO5", "MUC1", "ALDH3B2", "LPCAT1", "SLC25A13", "F5", "TBC1D8", "MMD", "PLAT", "SCML1",
"ZNF423", "MUC13", "PPFIA1", "RTN1", "MVP", "RAB14", "PLK2", "SDCBP", "LITAF", "APLP2", "ATP2A3",
RP", "KRT7", "HSD11B2", "GLUD1", "DHRS3", "H1F0", "KRAS", "S100A11", "CDC42", "CTSZ", "TACSTD2"
GFA", "CLIC5", "VLDLR", "MBOAT7", "STX3", "IL20RA", "GABRE", "MAN2A1", "CDH11", "NSDHL", "LMCD1
8", "ARMCX2", "DIAPH2", "ARG1") c("RHOB", "BAMBI", "IRF6", "HIST1H2AE", "MT1F", "PEA15", "AKR1C
, "TSTA3", "BSG", "IFI30", "POLR2L", "LARP4", "TLR4", "TYMP", "ITGA3", "G0S2", "HLA-DQA1", "LAMP3",
PRSS21", "ADD3", "DUSP3") c("BBX", "PHLDA2", "LAP3", "DHCR24", "NRCAM", "PPP1R7", "BCAP31", "TN
CD55", "MYC", "FUT8", "ANXA3") c("HIST1H2AE", "CDCP1", "PDSS1", "TIAM1", "TRIM37", "ASPN", "HIS
ARDBP", "IFITM2", "SCUBE2", "SQLE", "DHCR24", "PIK3R3", "PSPC1", "CXADR", "DACH1", "TFPI2", "PLTF
ADCY7", "HMGB3", "TPM3", "CARS2", "FOSL1", "XPO7", "ID1", "ITGA4", "THBD", "TNFRSF1B", "PEA15",

```
"PTX3", "SORL1", "PSMD12", "ATP2A1", "CNOT4", "SFRP1", "SGCB", "DUSP3", "HLTF", "FEZ1", "STAC", "
ALDH1A3", "DKC1", "EIF3D", "MRPS2", "PACSIN3", "ADCY7", "TPM3", "ADAM9", "XPO7", "ID1", "PEA15"
"SCML1", "TFPI2", "DNAJB6", "S100P", "ADRB2", "PLAC8") c("AP3D1", "ERBB2", "SCUBE2", "MS4A4A", "
HDC2", "FABP4", "MPPED2", "AREG", "SCGB2A2", "CLCA4", "CD59", "MGST3", "TACSTD2", "HPGD") c("T
, "RBM47", "MS4A6A", "CD14", "LYZ", "LTF", "TLR4", "GARS", "TPST2", "STAM", "IL1R1", "CCRL2", "KLK3
", "MAL", "ERBB3") c("CLDN10", "FXYD3", "IGFBP2", "MARCO", "SNRNP25", "ADM", "SLC38A6", "FAM1
B2", "DAZAP2", "NR4A2", "LYPD1", "CXCL2", "TACSTD2") c("DCTPP1", "DLD", "CASP1", "SCD", "HAX1", "
DKN1A", "ALDH1A3", "ZC3H12A", "CEACAM1", "PDE4B", "RRAS2", "CD19", "ISG20", "CXCL6", "HLA-DQA
1", "NRCAM", "KLF9", "LPCAT4", "ARL6IP5", "FXYD3", "SYCP2", "SAMD4A") c("C4BPA", "PEA15", "POLD4
3H12A", "PDE4B", "GZMB", "TRAF3IP2", "IER5", "HSD17B2", "AQP3", "NAMPT", "ASPN", "LIMA1") c("NU
NF19B", "RRAGC", "LAMC2", "LTF", "S100P", "VPS37B", "TCN1", "HSD17B2", "KRT14", "PLAC8", "ID1", "
MB3", "PLOD2", "OLFML2A", "STAB1", "CFH", "COX7A2", "PRDX4", "COL8A1", "NPL", "CALCRL", "ANXA4"
TRAF3IP2", "MLF1", "ACOX1", "VPS37B", "TCN1", "HSD17B2", "ID1", "AQP3", "BAG1", "LCK", "THOC6", "
FTN1", "MMP12", "HSPH1", "HSD17B2", "E2F8", "GNA15", "CYBB", "MTRR", "RANBP1", "KRT16", "C1QB
E1A", "ATP8B4", "IL6", "EEF1E1", "IGFBP2", "STAB1", "FTH1", "MAD2L1", "CA2", "ADM", "KBTBD11", "O
HSD17B2", "E2F8", "GNA15", "MTRR", "PCSK1", "PPBP", "HNRNPL", "NR4A2", "FGFBP1", "BMP2") c("ZN
CTSH", "SERPINB2", "CLCA4", "TTC9", "ANXA1", "ALDH1L1", "TPD52L1", "KRT19", "VCAM1", "PPA1", "FZ
43A3", "CCR1", "RFTN1", "IFI16", "TJP2", "KLK10", "RRS1", "TLR4", "TACSTD2", "POLR2L", "BCL3", "RHO
"STMN2", "CLIP2", "TARP", "MAN1A1", "CNOT4") c("SH3GL2", "MAL", "HIST1H1C", "TGFA", "HOPX", "T
DO", "PCDH9", "STK17A", "SOX9") c("FGFBP1", "SERPINB2", "PCSK1", "AQP3", "TRIM37", "TJP2", "RRS1",
BP4", "SCGB2A2", "PHLDA1", "FTH1") c("LST1", "TAF9B", "ACSL1", "GABRE", "HSPH1", "CYP3A5", "FZD6"
H1F0", "SNX11", "B4GALT5", "RAD23A", "ARMCX6", "NRCAM") c("OLFML2A", "FTH1", "FXYD3", "NRCAM
EFCAB2", "SERPINB6", "HOPX", "TGFA", "ZBED1", "COMMD8", "TSG101", "NKX3-1", "GAD1", "IL1R2", "P
BP3", "DGKA", "LAMB3", "FABP4", "PSCA", "GPRC5A", "CA2", "HPGD", "CTSH", "LYPD1", "TACSTD2", "SC
SLC2A6", "PLSCR3", "DUSP6", "BAMBI", "HS3ST1", "LYPD1", "HTR2B", "CXCL6", "HYOU1", "TJP2", "KRT1
", "CACNA2D2", "FOXO1") c("SNCG", "GPD1L", "DENND2D", "HIST1H2BD", "SERINC5", "ALOX5AP", "RN


24", "STK38", "PHF11", "ADI1", "GSTT1", "CTSE", "DYNLT1", "BBS1", "PERP", "PMM2", "GPATCH2", "SH3

NN1A", "RUFY2", "SASH1", "TM4SF1", "GMPR2", "PATZ1", "ZFHX3", "CAST", "VAV3", "MTUS1", "SH3BP4"
"TNFRSF21", "SPRED2", "AP1S1", "HNRNPH2") c("FBP1", "CDH1", "TUFT1", "PPFIBP2", "LIMCH1", "SH3Y
PPP2R5E", "PLAC8", "PRDX3", "MUC1", "RAB9A", "ZNF217", "LIMK2", "CARHSP1", "MBNL2", "HOPX", "T
", "ST3GAL5") c("AUTS2", "THBS2", "LAGE3", "MARCKSL1", "HGSNAT", "TPK1", "RAB9A", "ZNF217", "BA
2B", "PEG10", "VPS35", "TBC1D9", "ZNF586", "LIMK2", "PIK3IP1", "SLC15A2", "TBC1D8", "FMO5", "TMEN
GSTZ1", "RARA", "VPS35", "NLRP2", "LIMK2", "ABLIM1", "RNF43", "PIK3IP1", "S100P", "STXBP2", "SLC6
O5", "IDS", "TIMM9", "CERK", "NDUFA3", "GALNT1") c("MMP10", "ALDH3B2", "CDS1", "SPRY2", "TMPRS
"GPC4", "TMEM158", "STEAP1", "KRT17", "PENK", "MVP", "LITAF", "PLK2", "NLRP2", "SDCBP", "SELE", "
PX2", "KRT7", "SAT1", "GRB7", "GCH1", "SURF1", "MYCBP", "S100A11", "SMAD7", "FCGBP", "GABRP", "
SS3", "MGST3", "RAB5B", "LPL", "NFKBIE", "FGF13") c("TACSTD2", "ALDH3B2", "STK38L", "CXADR", "TM
DKN1A", "ALDH1A3", "S100P", "CCR1", "TRAF3IP2", "VNN1", "LYN", "ZYX", "TUBB6", "MAST4", "SLC6A1
BP", "G3BP2") c("DHX32", "SAT1", "TGFA", "CEACAM6", "SEMA3F", "PHLDA2", "SERPINA5", "SLC25A46",
B10", "E2F2") c("PERP", "WSB2", "S100A11", "AREG", "CTSE", "THOC7", "GSTT1", "TIMM9", "TXNDC9", "
", "SOX9", "SIK1", "NRIP3", "PNN", "PLK3", "TSC22D2", "GRIA2", "FUT8", "MED13", "PCDH9", "NUPL2", "
, "MUC16", "COL9A3", "TNFSF10", "EXPH5", "KYNU", "STC2", "SERPINB2", "LIMK2", "CLDN10", "CLCA2",
CXCL1", "MLF1", "TMEM158", "SOD2", "ID1", "FGFR3", "NXT1", "HK2", "IL1R1", "PRNP", "TMPRSS4", "A
XO21", "GCH1", "RAB33A", "ASPH", "SLC3A2", "HSPD1", "PGD", "GCLM", "RBP1", "TGOLN2", "DEFA4", "
```

"TMBIM1", "H1F0", "PHGDH", "CAST", "AREG", "STK39", "KIF14", "PDZD2") c("MLEC", "SLC25A13", "SLC
LHL21", "SMOX", "CD55", "TRIM16", "SIK1", "ATF3", "GRIA2", "REG3A", "SOX9", "FUT8", "CYP4F11", "PC
OX1", "S100P", "COL9A3", "GCH1", "H1F0", "THOC7", "SLC7A1", "ARPP19", "GLUL", "AREG", "SCEL", "S10
, "ELF4", "EDEM1", "MYO1B", "RRS1", "ALOX5", "STAM", "MYC", "DKC1", "KIF14", "PLIN2", "HLA-DQA1"
PX", "ERBB3", "FHIT", "ARMCX6", "CLDN16", "H1F0", "NRCAM", "ITM2B", "DHRS3", "CXCL5", "TGFA", "T
WD1", "SLC37A4", "PMM2", "ANXA4", "AREG", "TSPAN31", "TNNT1") c("CLIC3", "CIT", "STK39", "KIF20
IST1H2BD", "DKC1", "ATP13A3", "TNFRSF21", "TRIM37", "GRWD1", "ID1", "DNAJB6", "PITPNC1", "LCK",
", "PLSCR3", "PSMD3", "KLK7", "KRT17", "TRAPPC3", "SLC6A14", "SRGN", "KLK10", "MYC", "KLK6", "KRT
G", "LHPP", "RUSC1", "CTSE", "COX7B", "TSPAN31", "BAD", "LYPD1", "S100A11", "VAMP8", "NBR1", "M
A2", "KCNN4", "ADO", "SOX9") c("ALDH7A1", "SAT1", "COL4A3BP", "IER5", "LAMB3", "EDEM1", "TMPRS
LACTB2", "CASP7", "CDH3") c("NCAM1", "PKIG") c("SERPINB2", "ANXA3", "GSTZ1", "MMP10", "PLS3", "C
, "CD74", "THBD", "MGAM", "SAT1", "TM9SF2", "FGFR3", "EIF4A3", "CXCL1", "PDE4B", "CHD1", "ID1", "
", "ANXA6", "SKIV2L", "PTOV1", "DEFA4", "CAMK2N1", "NAV3", "PRTN3", "MTA2", "PPP3R1", "IARS2",

ERP", "NQO2", "CDK6", "PAICS", "PRDX1", "DERA", "ITFG1", "PRPF4", "SUZ12", "UBE2D1", "AREG", "SLPI

4A3", "PDE4B", "ID1", "ATF1", "ADAM9", "PAF1", "MMP1", "RBBP5", "CLIC1", "YWHAZ", "METTL5", "MS
2", "TNF") c("CSRP2", "SIK1", "CHD1", "DUSP1", "CCL2", "TREM1", "NR4A2", "FOS", "FOSB", "ARG2", "SV
INB2", "SMC6", "S100A11", "EHBP1", "ASRGL1", "SMARCA1", "SRPX", "PECR", "PPP2R5A", "MAPRE2", "
"ABHD2") c("RABEPK", "PPP2R2B", "MEIS1", "MAGEA3", "CYB5A", "GPM6B", "MAP3K1", "FKBP11", "FO
, "ADM", "LCN2", "FOS", "MYCBP", "SH3BGRL3", "CTSZ", "FOSB", "ANXA3", "TGFA", "KCNAB1", "GABAR
TCERG1", "RBP4", "TUBA1A", "RBM47", "MNDA", "RAB8A", "MPHOSPH6", "ARHGDIA", "ALOX5", "CLIC1
AD1", "NRCAM", "SERPINB6", "TSEN2", "PHLDA2", "PLS1", "ANP32E", "RBM47", "DHCR24", "ERBB3", "CE
2", "KLK6", "TMPRSS4", "DNAJB6", "AQP3", "PTP4A1", "ID1", "CDC42SE1", "HIST1H2AE", "KLK3", "ALDH1
CAM", "SLC16A4", "FXYD3", "MEIS1") c("GLUD2", "PEA15", "CDK2AP1", "C4BPA", "EMP1", "GALNT12", "
C25B", "CCL18", "S100A2", "IFI16", "ATAD2", "KRT17", "LY96", "KLK10", "TLR4", "APOBEC3A", "C1QB", "
TH1", "OLFML2A", "CALCRL", "CFH", "RGS4", "NRCAM", "PRDX4", "FCGR2C", "ATF3", "E2F5", "IGFBP2", "
37B", "CCL20", "BAG1", "TUBB3", "DDX3X", "CASP7", "BSG", "ASPN") c("SCD", "CXCL13", "MCM7", "RXR
0A11", "ANXA3", "MPPED2", "PPP4R1", "TACSTD2", "CLIC3", "ANXA1", "S100A10") c("ADAR", "BCAT1", "
", "LCK", "KLK6", "CDC25B", "CCDC86", "BSG", "VNN1", "AQP3", "S100A7") c("RRAS2", "PXDN", "FOXO1"
", "PPP4R1", "MCM10", "SNRPA1", "WTAP", "VNN1", "ALOX5", "HLA-DQA1", "TUBB3", "CCL20", "LYN", "
PX") c("SYCP2", "COX7A2", "FXYD3", "SLC38A6", "LYVE1", "TACSTD2", "ADM", "CD2AP", "FTH1", "MAD2
P1", "TACSTD2", "BMP2", "FST", "FEN1", "PGAM1", "SERPINB2", "TUBB3", "CCL20", "RP2", "TBC1D8B", "

:IT", "STOM", "GCLC", "IMPA1", "MATR3", "NR4A2", "PKP2", "SUB1", "MARCKSL1", "SPP1", "CLCA4", "SL
TD5", "KIF14", "KIF5B", "KLK10", "KLK6", "KLK7", "KRT14", "KRT16", "KRT17", "KYNU", "LAMC2", "LBR", "
"SERPINA5", "TARS") c("ANXA2", "AREG", "C4BPA", "COL8A1", "LTF", "P2RX1", "PACSIN3", "PIP", "PLK2"
OSBPL9", "PNKP", "PSAT1", "TBCD", "TCERG1", "TSEN2", "ZIC1") c("ADAM8", "CLIP2", "DAXX", "FCER1A",
", "CDC42SE1", "TCN1", "PLIN2", "BCL3", "RBM47", "OSBPL10", "IRF6", "RRAGC", "ANXA2", "TOMM34",
", "PHLDA2") c("COL4A3BP", "PLOD2", "ANXA4", "COL8A1", "PPP1CC", "TMED5", "ARL6IP5", "SLC25A46
D17B1", "FKBP1A") c("ANXA4", "LEPROTL1", "SLC25A46", "GSTT1", "SUB1", "BDH2", "COX7A2", "VAMP8
ATP6V1D", "SAT1", "ASPN", "ARFGAP3", "BCL2A1", "CAP1", "SPR", "HDAC1", "TACSTD2", "CDC42SE1", "

", "RAB15", "ADM", "PPT1", "TRIM22", "TACSTD2", "KLF9", "PDE1A", "TREM1", "OLFML2A", "IGFBP2") c
HITM", "SLC5A6", "CXCL13", "DLD", "FGFBP1", "CASP1", "SMC1A", "PSAT1", "ZIC1", "ZBTB43") c("SIK1", "
D52L1", "CYP4F3", "CEACAM5", "CTSH", "MUC4", "CROT", "BLNK") c("GCLC", "WFDC2", "CYP3A5", "MYO
L", "SEMA3A", "VCAM1", "FOXA1", "P2RY13", "SCNN1A", "TAX1BP3", "MYO6", "MMP10", "CFH") c("ITGA
SE", "MEX3C", "SSH3") c("POLR1D", "F5", "SSH3", "VAT1") c("PITRM1", "GALT", "ANXA4", "DDX42", "PO
A1", "RBM47", "ZYX", "CASK", "S100A7", "MYO1B", "TRAPPC3", "RHOBTB3", "HS3ST1", "DUSP6", "SMPD
B32", "FAM107A", "MEIS1") c("PPBP", "S100A6", "AREG", "NDRG1", "PPP1R9A", "ANXA2", "PLK2", "GST
PP4B", "PDZK1IP1", "IDO1", "UCHL3", "KLK5", "RBM47", "NRGN", "CX3CL1", "S100A7", "MYO1B", "IRF9"
"ATP1A1", "RBM47", "ZYX", "PUF60", "DUSP6", "ALDH1A3", "MMP1", "PLAT", "ARPP19", "ADAM10", "A
S2", "IFI30", "LTF", "TLR4", "TACSTD2", "STXBP6", "CD46", "ALOX5", "OSBPL10", "TMEM184B", "MMP1

MBOAT7", "ITGAE", "CHERP", "FGFR4", "DENND1A", "SAMHD1", "PPT2", "P2RX4", "ALOX5AP", "MCM6",
DRG1", "LTF", "TACSTD2", "TFF3", "OSBPL10", "SPINK1", "GDE1", "FUCA1", "PEA15", "CDK6", "PLK2", "M
"CEACAM6", "NRIP1", "BCL2A1", "NEU1", "SDC1", "STEAP1", "KRT5", "CAPN2", "PKP2", "PLAU", "TCN1",
A1", "MRFAP1L1", "PAPSS1", "PMM2", "FDPS", "LDLR", "HIST1H2BH", "HGD", "TIMM9", "TIPRL", "TSPAI

UBE2Q1", "SH3YL1", "HSPB8", "STX3", "IL20RA", "ECM1", "JAG1", "AREG", "KCTD3", "CCDC6", "PPFIBP2

RG2", "EVI2A", "ATF3", "CSRP2", "EGR2", "MYC", "PCDH9", "NRIP3") c("CSTB", "CLDN8", "ZFP36L2", "GR
, "AKR1B10", "LIPG", "CD58", "F2RL1", "PMM2", "NOP16", "ACAT2", "NPTX2", "NPTN", "CYP51A1", "DD
S100A2", "SCD", "SLC39A4", "GTF2I", "ST3GAL5", "SMYD2", "DNMT3A") c("GRHL2", "HOPX", "RARRES1"
H3", "FMO5", "ZIC1", "HIST1H2BD", "FBXO11", "MBNL2", "TP63", "SLC25A13", "S100A6", "SCGB1A1", "TI
PRPF4", "GALC", "REL", "USP6NL", "SMYD2", "CLEC11A") c("PIK3R3", "H1F0", "S100A11", "CDC42", "PRK

ZD8", "BCL2L1", "SPP1", "LPXN", "USP6NL") c("CSTA", "HOPX", "NUDT4", "H1F0", "SERINC5", "CYB561", "
PIK3R3", "S100A11", "CD9", "STXBP6", "APOLD1", "FZD5", "PRRG1", "ITGAV", "ALDH1L1", "DIO1", "DHR
PIB", "NLRP2", "PDZK1IP1", "TGFA", "TMEM165", "ABHD2", "CYB5A", "TMEM123", "NAMPT", "RAB2A",
"CYB5A", "WIPF2", "TIMM9", "SPP1", "USP6NL") c("PEX11A", "S100A11", "HOXA5", "ADAP1", "DNM3",
, "IER5", "RARRES2", "STEAP1", "KRT5", "DNAJC12", "DHRS1", "NLRP2", "SMARCA1", "HDAC2", "RGS16",
, "EFNB2", "GRIA2", "CD46", "GOLPH3", "APOLD1", "GPRC5A", "GPX2", "LCN2", "ECHDC2", "FCGBP", "M
1", "FGF13", "TAC1", "GTF2H5", "RAB2A", "OSBPL3", "FOXJ3", "SLPI", "BCAR3", "ABCA8", "LPL", "UCHL3"
2", "NCOA4", "CYB5R1", "PLK2", "PIK3IP1", "MDK", "ABCA12", "NAV2", "AP1G2", "PIP", "VPREB3", "TRIP
"ARPC5", "DDX21", "LYPD1", "SMPDL3A", "LIMA1", "SLC6A14", "VNN1", "PCSK1", "TJP2", "CYP1A1", "C
MPRSS3", "POLR2L", "SERPINB6", "HOPX", "SERPINA5", "ERBB3", "PLS1") c("SAMD4A", "CXCL10", "NRCA
T1H2BD", "PUF60", "GRWD1", "DKC1", "BSG", "RBP1", "POLR2L", "LARP4", "ARPC5", "DDX21", "LYPD1",
", "CST6", "GCDH", "AKR1B10", "CLTB", "RBBP8", "S100A2", "SLC25A13", "BCL2L1", "XIST", "E2F5", "FLG
IER5", "MAP7D1", "LBR", "TRAPPC3", "LAMC2", "PITPNC1", "ELK3", "RALGDS", "LYPD1", "F2RL1", "DIAP

'FGF9") c("AP3D1", "PHLDA2", "HIST1H1C", "NKX3-1", "CEACAM6", "ARMCX6", "HOPX", "ANGPT1", "PSN
, "PGAM1", "IER5", "PITPNC1", "RALGDS", "LYPD1", "PYGM", "TACSTD2", "BAG3", "RBP1", "HIST1H2AE"
'SCD5", "MRPL19", "HRAS", "DACH1", "PLEKHJ1", "PSMD11", "RBM3", "VPS37B", "EPB41L4B", "SMC2",
'AOK3", "NRIP1", "DDIT4", "DDX60", "PSMD10", "HLA-F", "SUB1", "TSPAN6", "IDH1", "CLCA4", "IRF9", "G
", "LAMP3", "LRFN4", "TNFRSF21", "KIF5B", "MID1IP1", "RBP4", "PLS1", "S100A2", "CSF1R", "DDX21", "C
07A", "SLC11A1", "TARS", "CFH", "CDR2L", "PLS1", "SAMD4A", "E2F5", "NPL", "FCGR2C", "CXCL10", "ATF
IDI1", "ZNF587", "ZIC1", "PHGDH", "GPSM2", "CYTIP", "CETN2", "FGR", "PSAT1", "CPS1", "AMD1", "RXRA
1", "GZMB", "SOD2", "TRAF3IP2", "SLC39A14", "CXCL1", "GZMA", "HIP1R", "SIGLEC15", "ACTN1", "MAPK
I", "GMPR2", "EMP1", "AREG", "SPINK1", "CXCL5", "NBR1", "DNAJA3", "ADAM12", "TFF3", "ZNF277", "H
IP93", "SMYD2", "CCDC90B", "CPS1", "ZIC1", "CASP1", "SLC5A6", "BANK1", "CXCL13", "DLD", "IGF2BP3")
AQP3", "TOMM34", "THBS1", "ZNF330", "S100A7", "BAG1", "TRA2A", "LCK", "VRK2", "KCNS3", "SRGN",
", "KLF9", "PLS1") c("HLA-B", "FUCA1", "IL13RA1", "CA2", "PER1", "ANXA2", "SLC9A3R1", "LTF", "SPINK1'
'LARP4", "ASPN", "PTP4A1", "PPBP", "EAF2", "DGAT1", "PUF60", "LYPD1", "CDK6", "ADAM9", "MAST4",

LFML2A", "CFH") c("ZNF304", "PWP1", "GGCT", "CDK6", "SPINK1", "GMPR2", "MORF4L1", "TFF3", "HBD
F587", "IDI1", "SCD", "MNAT1", "HNRNPH2", "PCCB", "SMC1A", "TSEN2", "CXCL13", "PCSK1", "CASP1", "
ZD6", "GPRC5A", "GABRP", "MUC4", "PSCA", "ATF3", "CEACAM5") c("CXCL10", "GPR87", "CLCA4", "IMPA
3TB3", "CYP1A1", "PLIN2", "SLC39A14", "ATAD2", "GRWD1", "POLG", "IFI30", "LYZ", "TCN1", "CA2", "PD
MPRSS3", "CREG1", "SCEL", "ERBB3", "POLR2L", "RBKS", "TSEN2", "LST1", "CEACAM6", "SERPINB6", "FEF
, "PUF60", "TACSTD2", "POLR2L", "DBP", "BCL3", "RBP1", "ATP6V1D", "ATAD2", "GRWD1", "PPBP", "POL
', "SEC13", "MARCKS", "CAPN2", "SH3BP4") c("SLCO1B3", "LSM4", "PSRC1", "SEC13", "SAP30", "CXADR"
", "IGFBP2") c("ADAM12", "GRWD1", "PPBP", "AREG", "DNAJA3", "PAX8", "TFF3", "PEA15", "LTF", "IGFB
HLDA2", "FERMT1", "HIST1H1C") c("ANXA4", "ARHGEF3") c("MYL6", "PPP1R9A", "CDKN2C", "PLK2", "EN
:NN1A") c("TBC1D8", "TSPAN31", "GPR87", "NR4A2", "TP53TG1", "KRT5", "SDC1", "ALDH1A3", "ID1", "A
7", "KRT16", "MLF1", "IL1R1", "KMO", "STOM", "HSD17B2", "CLDN4", "TCN1", "MNDA", "RGS1", "HDAC1
ASET2", "PABPC4", "RAB20", "CAMK2N1", "GCH1", "RBP1", "PELI2", "TNFAIP2", "ALDH7A1", "EFHC1", "G

BGRL3", "TIMM9", "SLPI", "CACNA2D2", "AREG", "REC8", "RHOA", "BDH2") c("FBP1", "SSH3", "HBP1", "

', "LRMP", "ARMCX6", "CACNA2D2", "PDZD2", "VAMP1", "AREG", "DGKA") c("OBSL1", "ZFP36L2", "DUSI
L1", "TOB1", "OPHN1", "MN1", "JAG1", "UCKL1", "HECA", "CITED2", "PPM1A", "ATP6V1D", "GOLGB1", "
TC9", "NAMPT", "USP6NL", "CLCA2", "TBC1D8", "IL13RA1", "CASP4", "GALNT1") c("IRAK1", "FERMT1", "
Z2B", "F5", "XPO7", "NDUFA3", "ST3GAL5") c("TACSTD2", "ALDH3B2", "STK38L", "CXADR", "CHAC1", "KF
M14B", "NCOA2", "LSM5", "NDUFA3", "PCCB", "GALNT1", "SPG11") c("DHRS1", "CD9", "CHL1", "SRPX", "
A14", "NAMPT", "STK38", "USP6NL", "PDZK1IP1", "ECH1", "TBC1D8", "CSTA", "ERMP1", "RUNX3", "EHF"
SS2", "HOXA5", "MAL", "NPAS2", "HECA", "S100A11", "PIGP", "MARCKS", "TP53TG1", "NDRG2", "PRRG1
ATP2A3", "DNAJC12", "ID3", "TMEM176B", "KRT5", "CXCL2", "AREG", "SAMD4A", "GYG1", "MMP1") c("
EYA2", "CLCA4", "KRT19", "MUC1", "IRF4", "GLUD1", "CD46", "ANXA3", "RAP2A", "ECHDC2", "NAMPT",
PRSS2", "TNFSF10", "GPX2", "SAT1", "GCH1", "S100A11", "RARRES1", "ST7", "CLDN4", "PCGF2", "PRRG1
4", "PSTPIP2", "ACTN1", "KLF10", "HSD17B2", "TJP2", "PLAC8", "HLA-DQA1", "SLC10A3", "AHCYL1", "PIG
, "LAP3", "ARMCX6", "ERBB3", "NRCAM", "MAL", "RBM47") c("ATF3", "MEIS1", "TRIM22", "FXYD3", "TA
'MAPK6", "LHPP", "ALAD", "SLC25A46", "CAPNS1", "MINPP1", "HGD", "GLUD1", "ZDHHC4", "ITFG1", "NI
'REG3A", "PFKFB3", "TRIM16") c("TMPRSS4", "ZC3H12A", "RALGDS", "SAT1", "EPHA2", "PLAT", "IER5", "
"SPP1") c("TMPRSS4", "S100A2", "PLAT", "S100A11", "RARRES1", "TCN1", "MVP", "ERBB2", "TACSTD2",
DORA2B", "G3BP1", "CD74", "MMP12", "PRSS3", "S100A2", "F2RL1", "ISG20", "KYNU", "CCL20", "HBEGF
TPSB2", "TPSAB1", "IFT57", "SCAMP3", "ALOX5AP", "SAMHD1", "GPD1L", "PDLIM2", "MBOAT7") c("DUS

C25A28", "PMM2", "HLA-B", "UCHL3", "DAPK1", "ANXA4", "NT5C2", "IL1RN", "ANKLE2", "ID3", "ADRM1"
DH9", "SNRK", "ANXA3", "OBSL1", "IFRD1") c("ASPN", "NOLC1", "DDX3X", "NUSAP1", "KLK6", "PCSK1", "
)0A7", "NAMPT", "SCML1", "TACSTD2", "ADCY7", "TCN1", "GALC", "PLAC8", "USP6NL", "FOXC1", "RGS1"
', "TNFRSF1A", "IFI16", "PBLD", "ATP13A3", "ATP6V1G1", "DIAPH1", "TNFRSF21", "STOM", "IL1R1", "GRV
SPAN6", "NFATC1", "RBM47") c("TACSTD2", "BECN1", "PLOD2", "PPP1CC", "ABHD2", "ATF3", "ADM", "R
\", "CADM1", "NTM", "H1F0", "KIF14", "FBP1", "GINS2", "FCER1A", "F5", "AREG", "CRABP2") c("IL1RN", "
"RPA3", "AQP3", "MLF1", "KMO", "TIAM1", "TMEM185B", "RBM47", "GLUD2", "ATP2A2", "RGS16", "TU
16", "RAC2", "KRT14", "DAXX", "MMP1") c("DUSP3", "APOD", "OBSL1", "IDS", "CD55", "IRF6", "NFKBIA",
APK6", "HOXB2", "TNNT1", "PERP", "KRT17", "PAPSS1", "FBL", "YIF1A", "CAPNS1", "AIM2") c("VAT1", "E
54", "CDC42SE1", "PUF60", "BCL2A1", "UBE2H", "TCN1", "RBP1", "LYPD1", "ALDH1A3", "PLAT", "SERPINE
QDPR", "S100A10", "HERC6", "IL1RN", "CXCL5", "MYO6", "IDH2", "MAOA") c("CAPN2", "GALNT2", "FAM3
SLC39A14", "ATF1", "ARHGDIB", "CEACAM1", "PLAC8", "CHI3L2", "CDKN1A", "PAF1", "PLAUR", "MMP1"
"RBP1", "NAAA", "EFHC1", "HSD17B4", "MAPKAPK3", "TNFAIP2", "PHKA1", "FGF9", "PPP1R9A", "UBE2M

", "SSBP1", "ANXA4", "TSPAN31", "WSB2", "ADI1", "SH3BGRL3", "RIPK2", "FERMT1", "DHCR24", "BAD",

N", "CCL20", "ASPN", "ALDH1A3", "RBP1", "HNRNPL", "NR4A2", "IL1B", "DNAJB6", "GLUD2", "ST14", "SE
2A", "ANXA3", "CDS1", "COL4A3BP", "NRIP3", "PCDH9", "FBXO11", "GRIA2", "RGS2", "TNF") c("AK1", "T
TXN", "RBBP7", "SRC", "RAD23B", "LST1") c("CXADR", "TFRC", "FAM3C", "PGK1", "PLCB3", "PRKCI", "HOF
SL2", "CCDC6", "CDK7", "CSRNP2", "CRIP2", "S100A8", "MAGT1", "AMFR", "UBA6", "GLI2", "SH3YL1", "M
APL1", "CD59", "SERPINB2", "GSTT1", "SMC6", "S100A11", "HIBCH", "HEBP2", "TUBB6", "GPRC5A", "TXN
", "MVP", "HMGB3", "S100P", "G0S2", "KRT14", "LTF", "MMP9", "CA2", "LAPTM5", "RHOBTB3", "DDX3X
EACAM6", "HIST1H1C", "SCEL", "RANBP9", "KLK3", "TMC5", "CXCL5", "ZC3H4") c("NRCAM", "ADM", "IGF
LA3", "TUBB6", "CXCL2", "TNFRSF21") c("HAS2", "DNAJC7", "TSEN2", "VDAC1", "BANK1", "TCERG1", "PS
'GSTP1", "PPP1R9A") c("PEA15", "NQO2", "FST", "TP53TG1", "MSN", "MT1M") c("CES2", "AXL", "DIRAS3

TREM1") c("CDKN2C", "NDUFS3", "AREG", "SPINK1", "CA2", "CFHR2", "PLN", "DNAJA3", "FADD", "MIA",
RA", "CASP1", "CYTIP", "RFC2", "SMC1A", "NME1") c("FLRT2", "KRT23", "TOX3", "LAMB1", "PPARG", "RH
'PPIC", "CLEC7A", "SERPINB1", "HNRNPD", "TACSTD2", "HOMER1") c("TFRC", "SPRR1B", "SPRR2B", "PI3"
', "EREG", "CCL8", "HGSNAT", "RBM26", "TRIM16", "YME1L1", "ADO", "SEMA6A", "MRPS30", "ABCA8") c
'FCGR2C", "SLC7A7", "RP2", "TNFRSF1B", "KRT14", "CD14", "CA2", "ID1", "SOD2", "HLA-DQB1", "DIAPH1
L1", "COL6A3", "FCGR2C", "KLF9", "PLOD2", "CA2", "SOCS1", "IL6", "TARS", "PDE1A") c("CDKN2C", "FUC
'ID1", "KMO", "PITPNC1", "FABP6", "HIST1H2BD", "ZYX", "MRPS2", "CDK6", "AQP3") c("NDUFB2", "PPP2

.C16A1", "IFITM1") c("GALNT1", "PBLD") c("AUH", "CD58", "HIST1H2BH", "SUB1", "PEX11A") c("KIF20A",
LMNB2", "LTF", "LYPD1", "MPHOSPH6", "MSN", "PDE4B", "PRMT1", "PRNP", "PRSS3", "PTP4A1", "RBP4
', "RUFY1", "TUBA1A", "UFM1") c("ARPC3", "CEACAM6", "COX6B1", "CTSC", "DDX10", "FERMT1", "GAD1
, "IGFBP1", "KRT23", "LAMB1", "PDLIM5", "PPARG", "TCL1A", "TOX3", "TPST2") c("ANXA3", "BMI1", "CSI
"RNF19B", "ZC3H12A", "BACE2", "CLDN4", "LYZ", "ISG20", "SLC35F2", "CDKN1A", "RABGGTB",
5", "COX7A2", "FXYD3", "TACSTD2", "FTH1", "NRCAM", "SLC16A2", "RAB15", "RGS4") c("COL8A1", "PLN"
3", "GLUD1", "PERP", "SLPI", "HK2", "S100A11", "FASN", "SNRPG", "RABGGTB", "NDUFC1", "ACVR1B", "F
TCN1", "BCL3", "HIST1H2BD", "RBM47", "ZC3H12A", "LRP4", "BACE2", "RGS16", "EPHA2") c("HAX1", "SA

:("PIP", "CDK6", "HLA-F", "S100A6", "GALNT12", "PEA15", "DNAJA3", "TFF3", "CA2", "AREG", "SPINK1", "
"PPARG", "NR4A2", "CYB5R1", "TOX3", "MYOT", "TKTL1", "GPR37", "KRT23", "RAB15", "GADD45B") c("G
OF", "SUB1", "GAD1", "CLCA2", "TP53TG1", "QPCT", "CEACAM6", "NETO2", "ADAMDEC1", "TFF1", "CDK
A3", "MYOF", "ADAM9", "GZMH", "CLCA2", "BTN3A3", "CCL4", "SUB1", "PSMD10", "P2RY13", "IFITM1",
DLR1D", "GTPBP8", "WBP2", "PLEK2", "FOXC1", "MSRB2", "DDIT4", "SSH3", "PSCA", "QPCT", "RGS4", "BA
DL3A", "ANXA2", "RRAS2", "PLAC8", "FOSL1", "ALDH1A3", "RBP4", "VNN1", "MMP1", "PLAT", "TLR4", "TH
P1", "IER5") c("TBC1D8", "PPBP", "FERMT1", "DDX10", "CIB1", "ACOT7", "ITGA3", "KLK5", "IRF9", "CTSC
', "HS3ST1", "SMPDL3A", "IFNGR2", "DDX42", "KLF7", "ANXA2", "CTSC", "PLAC8", "IFITM3", "RIN2", "IFIT
ATP6V1D", "ID1", "ARPC3", "NAMPT", "IER5") c("OPA1", "IGF2BP3", "SMC1A", "IDI1", "GLOD4", "SCD", "I
2", "IL1R1", "PEA15", "CCL20", "TJP2", "JUNB", "CYP1A1", "TMPRSS4", "MVP", "STOM", "LYZ", "CD37", "(

ANBA", "PPBP", "IER5", "ANXA2", "PIP", "MLLT11", "S100A6", "GMPR2", "APH1A", "MRFAP1L1", "PMM
, "AQP9", "GCLC", "ADAM9", "SH3BP4", "SLC25A13", "GSTK1", "TSPAN6", "WFDC2", "DDX42", "TBC1D8"
N31", "TSEN2", "TXNDC9", "AREG", "SLPI", "VAMP8", "RHOA", "PRPF4", "CDC37L1", "MCM6", "LDHA", "

IA2", "PIM1", "APOLD1", "TOX3", "JUNB", "STK38L", "CD37", "YPEL5", "DDX17", "ANXA3", "FBXO11", "F
X21", "LDLR", "STX3", "TPPP3", "TRAF3IP2", "NSDHL", "KLK10", "WTAP", "DDA1", "HGD", "RGS20", "PHL
', "MPZL2", "PIK3R3", "H1F0", "FGFR3", "IRF6", "GLUL", "S100A11", "ELF3", "COL9A3", "ERBB2", "TACSTI
BC1D8", "SSR2", "ZBTB38", "HMG20B", "CISD1", "MAN2A1", "CYB5A", "NSDHL", "TIMM9", "HGSNAT", "I
AR2B", "TRIB3", "STXBP6", "AKR1C3", "FZD5", "GIMAP4", "CFB", "ZNF160", "CALCOCO1", "FGL2", "RNF1

"CDC42", "MAPK1", "PIM1", "FHIT", "MBD2", "SIX1", "CETN2", "GLA", "TMC5", "CYFIP1", "ABHD2", "EPB
S1", "F5", "TPK1", "ASGR2", "LRRC15", "TSPYL5", "TSPYL4", "ADM", "HSD17B2", "SMARCA1", "HDAC2",
"MYBL1", "PDCD6", "LRRC8D", "LPXN", "PPT2", "REL", "USP6NL", "SLC39A4", "CDH1", "STK38") c("UAP1
"TMPRSS2", "OLFM4", "PRRG1", "HECA", "SHROOM2", "TBC1D9", "TP53TG1", "ANXA9", "PIK3IP1", "FG
"ADAM10", "PDK4", "SNX6", "CXCL2", "AREG", "TOR1A", "LAMC2", "TMEM158", "MMP1", "GYG1") c("
UC1", "ANXA3", "GRB7", "SAT1", "KRT19", "UBE2H", "SMAD7", "GSTT1", "CTNNA1", "RAP2A", "ADM", "

6", "SLC25A13", "ALDH2", "MMD", "ADAM8", "PTTG1IP", "DDX21", "TBX2", "NPL", "CR2", "KLK10", "LAC
D14", "THBS1", "CCR1", "HS3ST1", "LYN", "HSD17B2", "TNFRSF21", "S100P", "RBP4", "TCN1", "F2RL1", "
AM", "ARL6IP5", "LAMB3", "CFH", "PEX11A", "FAM107A", "CYP3A5", "PRDX4", "SOCS1", "MEIS1", "SERPI
"LAMB3", "LIMA1", "PPBP", "PCSK1", "TJP2", "HSD17B2", "TNFRSF21", "TCN1", "FST", "CCL20", "PLAT",
5") c("DHRS11", "UQCRH", "MFHAS1", "TSTA3", "RBP1", "FBXO2", "CDH1", "HOPX", "TCN1", "TBC1D8", "
PH1", "CA2", "FGFR3", "TACSTD2", "BAG3", "PBLD", "FOLR2", "STOM", "CEACAM1", "NEK7", "ARHGDIB",

MD11", "MAL", "ZC3H4", "CXCL5", "TMC5") c("CYP3A5", "PSRC1", "MYO5A", "FTH1", "ADCY7", "STAB1",
, "ASPN", "VPS37B", "TUBB6", "PIP5K1A", "AXIN1", "CLINT1", "ARPP19", "DDX21", "SERPINB2", "RRS1",
"ARPP19", "MUC1", "SLC37A4", "STC2", "POLR2I", "XIST", "TFPI2", "E2F5", "ADRB2", "PDCD5", "NPR3", "
GCLC", "CXCL10", "KRT5", "ZNF586", "COG5", "MARCKSL1", "NES", "TACSTD2", "FMO2") c("AHCYL1", "KR

F3", "TACSTD2", "MEIS1") c("R3HDM2", "PPBP", "DNAJA3", "IGFBP2", "TFF3", "FCGR2B", "ALAD", "HBD",
A", "PCCB", "DAZAP2", "SLC5A6", "MN1") c("CH25H", "SPRR3", "CTBP2", "TPST2", "IGFBP1", "FCER1A", "

BD", "IER5", "ASPN", "PLK2", "PIP") c("PLIN2", "CIB1", "PEA15", "CEACAM6", "POLD4", "HGD", "PSCA", "
) c("PDK4", "INSR", "IGFBP1", "TOX3", "PPARG", "LIF", "KRT23", "APBA2", "NUP210", "PAN2") c("EGR2",

", "PIP", "COL8A1", "SRBD1", "CR2", "GSTP1", "HBD", "DNAJC1", "ASPN", "DNAJA3", "PPBP", "DGAT1", "
"PLS1") c("FKBP5", "RXRA", "WDFY3", "PACSIN2", "ZIC1", "CYTIP", "TSPYL1", "TTLL12", "PNKP", "BANK1

", "OSBPL10", "AREG", "ENPEP", "ASPN", "NDRG1", "IGFBP2", "DNMT3A", "C4BPA", "DNAJA3", "CA2", "F
EFNA4", "FGFBP1") c("ILVBL", "FCER1A", "NR4A2", "NR3C2", "RHOBTB3", "PAN2") c("PAK2", "GRIA2", "
A1", "PSMD10", "HCP5", "BTN3A3", "FZD6", "TCN1", "CEACAM6", "TXNDC9", "ALOX5", "PSMB8") c("NDC
E4B", "TUBB3", "BSG", "SIGLEC15", "CD19", "MYO1B", "FCGR2B", "ALDH1A3", "RABGGTB", "TWIST1",
RMT1", "MAP2K1", "RBM47", "GDE1") c("ATF3", "PDE1A", "CXCL10", "OLFML2A", "MEIS1", "TACSTD2", "
LG", "TCN1", "PDE4B", "TUBB3", "BSG", "PGAM1", "ALDH1A3", "GLRB", "MTHFD1", "XPO7", "IDE", "MID
) c("TAF9B", "AREG", "PRDX1", "ADI1", "LSM4") c("SCARA3", "AREG", "DHRS2", "ZFHX3", "ADI1", "AURK
P2") c("MAST4", "SLC25A6", "PLIN2", "VAMP8", "GRWD1", "PPBP", "FST", "ZNF586", "PEA15", "TCN1", "
PEP", "TFF3", "ZNF277", "NBR1", "TSG101", "FUCA1", "AREG") c("CTSC", "ENPEP", "MPHOSPH9", "NQO
LOX5", "CLCA2", "TCN1", "TXNDC9", "TACSTD2") c("TBC1D8", "RRM2", "TRIM22", "SPRR2B", "CXADR", "
L", "MVP", "DAZAP2", "CDKN1A", "CDC42SE1", "PLAC8", "CA2", "CMTM6", "SRGN", "RBM47", "PLIN2", "
GRB14", "MBOAT7", "RARA", "FGFR1", "CDIPT", "ZMPSTE24", "AP1G2", "SEMA4C", "PLSCR4", "MTMR11

CLIC3", "CRABP2", "SCAND1", "UGT2B4", "MYL5", "SURF1", "TMBIM1", "SH3GLB2", "MCCC1", "HIST1H1

P1", "SOX9", "YPEL5", "ANXA3", "FUT8", "FOXA1", "ADO", "TRIM16", "PELI1", "CFI", "CD55", "REG3A", "F
ZNF217", "CALML5", "ADO", "RAB11FIP1", "SASH1", "CASP10", "CCR2", "TACC2", "DVL3", "ZMYM2", "SK
SERINC5", "EDF1", "BDH1", "PIM1", "CYB561", "RUFY1", "ATP6AP1", "TFAP2C", "NUDT4", "HOPX", "CSN
RT7", "CHI3L1", "CLDN10", "GCH1", "S100A11", "RARRES1", "ST7", "FCGBP", "TCN1", "PRRG1", "MVP", "
MYCBP", "S100A11", "MFAP4", "FMR1", "HSD17B2", "PRRG1", "SLC9A3R1", "TPK1", "ADH1C", "PLAC8",
, "VAMP5", "PCCB", "GALNT1") c("DNAJC4", "COL14A1", "HBP1", "GPD1L", "BTG2", "KLF2", "ERBB2", "L
", "OLFM4", "ANXA9", "MAT2B", "PEG10", "VPS35", "TBC1D9", "FGB", "ZNF586", "PIK3IP1", "TTC9", "FM
GSTK1", "KRT7", "SEMA3C", "PRKD1", "EMX2", "SLC39A4", "HEBP1", "CDKN1A", "ALCAM", "CAPN2", "M
"GABARAPL1", "ZNF611", "USP6NL", "CFI", "GSTT1", "CTDSP1", "PKN2", "IDS", "XPO7", "SH3BGRL3", "SL
", "MVP", "ELF3", "TPK1", "PLAC8", "PRDX3", "FOXC1", "MUC1", "RAB9A", "S100A10", "S100P", "STXBP

CSTD2", "SNRNP25", "SYNJ2", "SOCS1", "SERPINA5", "SLC25A46", "CA2", "SPDEF", "CXCL10", "FCGR2C",
DUFA4", "LGALS3", "ANXA4") c("CAST", "FBXW4", "ECM1", "CRABP2", "VAT1", "AREG", "S100A13", "DEC
TCN1", "ASPN", "AQP3", "TACSTD2", "ALDH1A3", "TRAF3IP2", "ZYX", "TUBB6", "MAST4", "HSD17B2", "T
, "S100P", "THOC7", "PIK3R3", "TIMM9", "MUC16", "MINPP1", "FMO5", "TNFSF10", "PRSS21", "ASXL2",
", "LYZ", "S100P", "MNDA", "TYMP", "CXCL6", "TOMM34", "BAG3", "EFHD2", "KLK7", "RAB8A", "WTAP"
P3", "SFRP1", "PTX3", "PDE8B", "CCDC69", "MMP3", "ZFPM2", "CNOT4", "STMN2", "ATP2A1", "SORL1"

', "SLC16A1", "NRBF2", "IRF9", "IMPA1", "HES1", "CHI3L2", "LYN", "S100A10", "CTSC", "MARCKS", "SOD2
MLF1", "MRPS2", "ID1", "TMPRSS4", "G3BP1", "SERPINB1", "ATP6V1D", "LAMB3", "FGFBP1", "CCL20", "
', "PLAT", "SPP1") c("ITM2B", "PBK", "ARMCX1", "SLC25A13", "PARP2", "FAM102A", "NUSAP1", "PLTP", "

EXO2", "NRCAM", "OLFML2A", "IGFBP2", "CALCOCO1", "SYCP2", "ANXA4", "COL6A3", "CA2", "SPDEF", "
TACSTD2", "VNN1", "HLA-B", "EIF4A3", "LCN2", "OSBPL10", "PLS3", "KLK5", "TUBA1A", "DUOX2", "HLA-
BB6", "LIMA1", "PLAT") c("PCSK1", "VDAC1", "BARD1", "TMEM123", "SMC1A", "PCCB", "HELLS", "AKAP
"INHBB", "SLC19A2", "PIM1", "CSTB", "MYC", "GRIA2", "ADO", "EPHB2") c("ALDH7A1", "BPHL", "GRN",
IF3L", "NINL", "FBXW4", "AREG", "SCAND1", "IDH1", "SH3GLB2", "MGST3", "CADM1", "DHRS2", "RND3"
31", "TACSTD2", "NR4A2", "EPHA2", "TNFRSF21", "KLK7", "HIST1H2BD", "GRHPR", "KLK6", "MMP1") c("
C", "AQP9", "INPP4B", "MSN", "CSF3R", "CXCL5", "STEAP1", "LACTB2", "MMP1", "CASP7", "CDH3", "ALD
, "RBBP5", "LYZ", "CLIC1", "PFN1", "PLSCR3", "MSN", "ITGA4", "CCL20", "MDM4", "ASPN", "TRA2A", "AL
1", "PLSCR4", "CALR", "GSTT1", "NR1H2", "ZNHIT1", "PTHLH", "MOCS2", "ITGAE", "PGD", "ASPH", "CNR1

"GGCT", "COX7B", "UQCRQ", "CARS2", "TUBA4A", "UBE2M", "SLC25A46", "GSTT1", "SDC1", "NDUFV2",

RPINB1", "ARPC3", "TPPP3", "COL4A3BP", "OAZ1", "PRMT1", "EFHD2", "CXCL2", "TNFAIP6", "SERPINB2"
CTN1", "AIDA", "NDUFB2", "CFD", "EIF2S3", "COX7A1", "FABP7", "LGALS1", "AKR7A3") c("MBTPS1", "MY
X", "CLDN10", "EXPH5", "MUC16", "GALNT1", "SAT1", "PDE4B", "MICA", "SLPI", "SSBP1", "RBP1", "WSB
METTL5", "NMU", "MN1", "ASAP2", "CFD", "CTNNB1", "AGPAT1", "HSD17B4", "UGP2", "PBX3", "ROBO1",
", "CFI", "GRIA2", "KLK11", "ABHD2") c("MID1", "PSME2", "CYB5A", "SPON1", "PAEP", "SELE", "GALNT1"

BP2", "FTH1", "GLRX", "PLS1", "STAB1", "CLDN10", "CA2", "ARHGEF3", "COL6A3", "MERTK", "ATP8B4", "
AT1", "AMD1", "DLD", "SMC1A", "FUBP1", "LARS2", "CLSTN1", "ELF5", "ZIC1", "ZC3H4", "PNKP") c("TCL1
", "ACADVL", "ALDH1A2") c("LYPD1", "IL1RN", "SERPINB2", "CYP4F3", "SEMA3A", "ACTN4", "SCNN1A", "

"CXCL5", "PIP", "SLC9A3R1", "LTF", "GDE1", "FAM120A", "IGFBP2", "CR2", "GSTP1", "RBPMS", "FCGR2B
OBTB3", "GPR37") c("TOP1", "CCL8", "TOX3", "EGR2", "PCDH9", "EGR3", "CSRP2", "ATF3", "FBXO11", "T
', "HOPX") c("LYPD1", "SLPI", "PERP", "FSCN1", "S100A11", "SET", "VAMP8", "TNNT1") c("UBE2G1", "SH3
c("PPP1R16B", "CALR", "CARM1", "NUP85", "CAMK2N1", "MAP4K1", "PABPC4", "CDIPT", "GSTT1", "PRT

A1", "CXCL5", "DGAT1", "LTF", "TACSTD2", "DNMT3A", "C4BPA", "PLK2", "SPINK1", "CA2", "NDRG1", "P
R1B", "IGF2BP3", "MLH1", "HAS2", "RBBP8", "CXCL13", "CASP1", "DLD", "SLC5A6", "EHF", "CPS1") c("AL

"BANK1", "HIST1H1C") c("CFB", "TMPRSS2", "MARCKS", "GALNT1", "POLB", "STOM", "C

, "APH1A", "TMEM38B", "LTF", "PLK2", "ASPN", "RBPMS", "FUCA1", "BAMBI", "TACST
RHOA", "ADI1", "SNRPE", "AREG", "DYNLT1") c("PARVB", "MGST3", "EIF3L", "GCNT3", "SLC

"CLEC7A", "TSPAN6", "TXNDC9", "ADAMDEC1", "CXCL10") c("AKR1B10", "GALNT1"

BDH2", "STK38", "SMYD2", "DNMT3A", "CACNA2D2") c("USP48", "GAS2L1", "FBP1"

PTX3", "NRIP3", "SPP1", "S100A2", "FAM193B", "SAMHD1", "DNAJC7", "THBS2",

B", "FMO5", "ALDH3B2", "GADD45A", "HMOX1", "SLC25A13", "CDS1", "NRGN", "NPA
USP48", "KRT7", "ID3", "HEBP1", "EPCAM", "CDC42", "EFNB2", "MUC13", "IGF2R
TGFA", "ABHD2", "HSPB8", "NAMPT", "AREG", "SPP1", "JUND", "SLPI", "PPT2", "

, "GSTP1", "LTF", "S100A6", "CFHR2", "CDK2AP1", "HS2ST1", "C4BPA", "TACSTD2

PPBP", "HLA-B", "HLA-F", "CDKN2C", "PIP") c("PSCA", "CYP24A1", "NPTX2", "RASL11B

80", "AKR1B10", "E2F2", "TRIM22", "CEACAM6", "DNAJA3") c("MRFAP1L1", "M

"ALOX5") c("ATOX1", "BST2", "FN1", "TSKU", "VPS28", "SLC24A3", "MACROD1",

C", "RARA", "BANK1", "IDH1", "CAPN1", "ZNF611", "TALDO1", "VAT1", "GMPR2",

K1A1", "CSTA", "IARS2", "PKN2", "BCL2L2", "LAMP1", "TIAM1", "SERPINB6", "ACA

"PIK3R3", "CA2", "HOPX", "EPS8L2", "F5", "PSMD10", "PDLIM1", "DIO1", "ANKRD27

O5", "NDUFA3", "PCCB", "MT1G", "GALNT1", "MGST3", "SPG11") c("ALDH3B2"

"SLC16A4", "CLDN10", "ANXA4", "NRCAM", "ABCB6") c("LTF", "PIP", "PLK2", "IER5",

FAM13A", "CXCL10") c("SLC25A4", "TACSTD2", "HLA-B", "OSBPL10", "TUBA1A", "HLA-F"

[illegible]H1A3") c("KDELR2", "PI3", "NDUFB2", "TMEM183A", "SPRR2B", "MSN", "KRT6B", "CASP7

", "ARHGAP6", "PCCA", "SERINC5") c("ZNF83", "SORL1", "TARP", "TFDP2", "HP1BP3

, "HSPG2", "KLF2", "PEBP1", "OSTF1", "COPS7A", "SMC6", "CCDC88A", "TM4SF20

'H1FX", "CAPNS1", "TMEM147", "HMGA1", "COBLL1", "PLS3", "ANXA1", "QDPR",
