## Supplementary material for "Computational Microbiome Pharmacology Analysis Elucidates the Anti-Cancer Potential of Vaginal Microbes and Metabolites": Table S4- gene list

V1 "V2" "V3" "V4" "V5" "V6" "V7" "V8" "V9" "V10" "V11" "V12" "V13" "V14" "V15" "V16" "V17" "V18"
1 c("SNAP25", "TNNC1", "GPC3", "ZNF586", "RUNX1", "LIMCH1", "LAMB1") c("PLLP", "MEIS2", "CST3",
2 c("ABLIM1", "ALDH1L1", "ATM", "BEX4", "CCDC28A", "CD99", "CYP27A1", "DUSP2", "ENPEP", "GPC3",
SLC2A10, "SORT1", "TCEAL2", "TM7SF2", "TMED3", "XIST") c("ABCG2", "ACO1", "APOL2", "ARL6IP5", "C
TSC22D3, "TXNIP") c("ACO1", "FASN", "HMGCS1", "PLA1A", "PRKDC", "RNF128", "SLC17A5", "SPINK1",
3 c("GOLM1", "STUB1", "PHKB", "ABLIM1", "TOMM20", "SATB2", "HMOX1", "UNG", "FAM173A", "ALDH
4 c("SLC24A3", "ZHX2", "VAT1", "ZNF586", "TSPAN6", "BMI1", "GOLM1", "SMARCA1", "LIMCH1", "AASS
5 c("CNN3", "LIMCH1", "GPC3", "IL17RB", "PLXNA2", "TNNC1", "SEMA5A") c("NVL", "NFIB", "UNC119B"
6 c("VAT1", "HMOX1", "IL17RB", "CEP68", "SLC43A1", "PTPN3", "GPC3", "ARHGEF2") c("MT1H", "RRAG
7 c("SNAP25", "CNN3", "HMOX1", "RND3") c("ACSL1", "ITGAE", "SPRR2B", "GDE1", "TMPRSS3") c("PCSK
8 c("TSPAN6", "ZNF586", "HMOX1", "APOD", "PIR", "ALDH5A1", "SASH1", "SMARCA1", "BCAS1", "SNAP
9 c("SNAP25", "RND3", "PDZD2", "ALDH5A1", "BLCAP", "RUNX1", "CALML3", "GLRX", "ENPEP", "GABRP
HS2ST1, "PPDPF", "ZMYM2", "MTA1", "CLIC3", "WNT5A", "MPHOSPH8", "AHI1", "SCNN1A", "OPTN",
FXYD6, "PDE8B", "LSP1", "CASC3", "GSTA1", "DSE", "PLTP", "ZNF423", "FGFR1", "SLIT2", "GNB5", "SPAR
TMED10, "GOLM1", "TBX2", "FGFR1", "ZEB2", "FLNA", "CYP26B1", "MCAM", "PCDH17", "MAP1A", "HG
MED13L, "SPR", "RIMS3", "SLC25A46", "UBE2G2", "CD93", "RPN2", "PECR", "POLR2K", "FLNA", "PSMD1
DHTKD1, "SOX4", "AEBP1", "SCPEP1", "IPW", "PEBP1", "GOLM1", "RGN", "BGN", "SLIT2", "FLRT2", "PRR
CD81, "SLIT2", "EDNRA", "SPARCL1", "MAOB", "COL4A6", "NOTCH1", "STC1") c("SNAP25", "ZMAT3", "R
FXYD6, "SELE", "CACNA2D3", "MATN2", "GYS1", "HUWE1", "PPID", "PARM1", "NCL", "IVD", "RGS5", "CD
ROGDI, "SYCP2", "INHBB", "PELI1", "ADAM28") c("PLAT", "RPS27L", "PCSK1", "PSMB9", "CA12", "ADM",
PLXND1, "PKP1", "MAOB", "COL4A6", "ATP2A3", "LONP1", "PDSS2", "SSX1", "ELOVL5") c("HSPA8", "RPS
ASCC3, "FN1", "PLTP", "MATN2", "GYS1", "HUWE1", "XPO7", "PPID", "ENG", "PDE4A", "MRPL18", "CPT1
NOTCH1) c("SNAP25", "NCAM1", "PRSS23", "MGLL", "FANCL", "ZNF337", "LOR", "SPEN", "IFIT1", "HSD1
COL4A6, "PDSS2", "NOTCH1") c("ZMAT3", "FDXR", "RET", "RRM2", "PRSS23", "RNF128", "MGLL", "HERC
EDNRA, "HGF", "PGF", "LAPTM4B", "ELOVL5", "FJX1", "TUBB3", "RANBP1", "HSPA6") c("SNAP25", "ZMA
GLI2, "CLIP2", "IGLL1", "FOXF1", "CCL21", "IGFBP5", "DACH1", "ATP6V0E2", "SYCP2", "INHBB", "PELI1")
ICAM3, "COL4A6", "SYCP2", "PELI1") c("BTG3", "FSTL1", "ADA", "IFIT3", "MFAP5", "FKBP11", "GCLM",
) c("SNAP25", "NR3C1", "ZMAT3", "MGLL", "EXO1", "FEN1", "IFIT1", "HERC6", "IFI44", "SERPINB9", "MS
GYS1, "HSPA5", "PPID", "CD81", "ENG", "EDNRA", "DPY19L1", "GLI2", "LAPTM4B", "COL4A6", "ATP2A3"
SPARCL1, "COL4A6", "NOTCH1") c("SACS", "ZMAT3", "NCAM1", "NOVA1", "SNCA", "FHOD3", "PRSS23",
ABCA3, "GDPD5", "IFITM2", "LTBP3", "TRAK2", "HBB", "PLTP", "DMXL2", "MATN2", "PBX3", "RNF130",
PLXND1, "MCAM", "NCOR2", "VWF", "COL6A3", "NDN", "PNMA1", "XIST", "LRRN3") c("SNAP25", "PRSS
CYB5R3, "IGFBP5", "COL4A6", "DPT", "COL4A5", "NOTCH1", "CTSB") c("SNAP25", "ZMAT3", "PRSS23",
EDNRA, "MRPL18", "STEAP3", "HGF", "MAOB", "COL4A6", "SSX1", "NOTCH1") c("SNAP25", "HP", "DTL",
MATN2, "ABCG2", "PPID", "CD81", "EPHA3", "FGFR1", "HGF", "SMPDL3A", "CPT1A", "SLC10A3", "LAPTM
IFITM2, "U2AF1", "ANXA5", "PLTP", "DMXL2", "LYZ", "MATN2", "PPID", "TBC1D1", "IVD", "RGN", "CD81
LSP1, "FN1", "LYZ", "MATN2", "PPID", "RGS5", "ACD", "ENG", "GLI2", "CLIP2", "IGLL1", "RCAN2", "CPT1
LHFPL2, "IFITM2", "FXYD6", "MSMB", "ANXA5", "AEBP1", "GYS1", "HSPA5", "ENG", "EDNRA", "DPY19L1
SPARCL1, "HGF", "NOTCH1", "XIST") c("SNAP25", "RET", "RPS27L", "PRSS23", "MGLL", "HERC5", "IFIT1",
PLTP, "MATN2", "GYS1", "PPID", "IVD", "CD81", "ENG", "EDNRA", "CPT1A", "FOXF1", "CCL21", "ANGPTL
) c("TUBB2A", "CAV1", "SLC39A6", "ALOX5", "AKAP12", "DIP2C", "SERPINA1", "IGFBP6", "IFI44L", "FSTL1
WSB1, "PTPN7", "CTNNAL1", "FN1", "ANXA5", "GYS1", "HSPA5", "CD81", "ENG", "EDNRA", "GLI2", "CPT
SLIT2, "EDNRA", "SPARCL1", "PRRX1", "COL4A6", "NOTCH1") c("TCN1", "SNAP25", "ZMAT3", "PRSS23",
ANXA5, "RPS6KA1", "ACOT9", "GYS1", "RNF10", "IVD", "CD81", "EDNRA", "MAOB", "COL4A6", "CCNE2",
GYS1, "RNF10", "PPID", "IVD", "CD81", "ENG", "EDNRA", "ROGDI", "PELI1", "SNCAIP") c("TUBB2A", "SLC
ABCG2, "NPDC1", "EDNRA", "PKP1", "COL21A1", "H1F0", "MAOB", "PADI2", "COL9A3", "CNR1", "MUC2
GYS1, "HUWE1", "PBX3", "PPID", "EDNRA", "PDE4A", "PRRX1", "HGF", "PPT1", "TCFL5", "NOTCH1") c("I

SERPINF1, "IRF7", "GPI", "LTBP3", "IL10RA", "TGFBR2", "MPDZ", "IDH2", "PLTP", "GLDC", "PDHX", "HS3S
CACNA2D3, "MPDZ", "CRIP1", "ASCC3", "FN1", "PLTP", "GYS1", "HUWE1", "RNF10", "PPID", "IVD", "CD8
DMXL2, "HUWE1", "PPID", "IVD", "CD81", "SLIT2", "MRPL18", "STEAP3", "COL4A6", "RNASE1", "STC1", "
GLIPR1) c("ZMAT3", "NCAM1", "TGFA", "SESN1", "NOVA1", "PDZD8", "PDGFC", "PRSS23", "TRIM14", "T
TBC1D1, "RGN", "RGS5", "SLIT2", "PDE4A", "ZEB2", "SPARCL1", "PRRX1", "MAP1A", "NR2F1", "RCAN2",
RCAN2, "RPS6KA2", "HLA-DPA1", "RNASE1", "CD14", "NOTCH1", "SNCAIP") c("ME2", "TCN1", "PLAT", "F
10 c("HMOX1", "PAK1", "TP53I3", "RND3", "HAUS4", "BAG2", "TSPAN6", "PIR", "ALDH5A1", "CNN3", "U
11 c("BEX4", "CD99", "ABLIM1", "CALML3", "HMOX1", "TSPAN6", "GPC3", "EFNB3", "SORT1", "DUSP2",
GABBR1, "SLC27A3", "MFAP2", "ASMTL", "RNF128", "GOLGB1", "HBP1", "PPP1R13B", "BNIP3", "FOSL2"
COBLL1, "MAGED1", "PPP1R13B", "BGN", "LBH", "COL15A1") c("UCHL1", "SPINK1", "ABCB6", "MTHFD2
RGS5, "HPRT1", "TMEM176B", "C1QB", "GBE1", "PON3", "RHOBTB1", "NR2F1") c("NQO1", "CLNS1A", "I
12 c("PRKACB", "DDX42", "PTPN3", "DUSP3") c("RGS4", "MT1M") c("COMP", "SAP30", "CHRNA3") c("SL
13 c("VAT1", "SMARCA1", "LAMB1", "DDX42", "SASH1", "UBB", "ALDH9A1", "CHN2", "CNN3", "APOD", "
14 c("SATB2", "SNAP25", "GUSB", "SEMA5A", "SOSTDC1", "GALNT11", "CACNA1D", "PAK1", "LIMCH1", "
15 c("GABRP", "ZNF586", "DUSP3", "LAMB1", "VAT1", "SATB2", "CST6", "SLC24A3", "SLC4A4", "CD99", "
16 c("APOD", "IVNS1ABP", "SORT1", "QDPR", "DUSP2", "PRKACB", "PDZD2", "TSC22D1", "ZNF365", "AD
PAM, "RNASET2") c("MYBPC1", "ZNF365", "USP13", "NRP1", "FLG", "TBCD", "BCHE", "UCHL1", "MACF1"
WT1, "MPDZ", "ITIH2", "PRELP", "OSR2") c("CXCL10", "SFRP1", "SORBS1", "DAB2", "SCG5", "LAMA4", "Z
17 c("PAK1", "UVRAG", "HGSNAT") c("CXADR", "TXNIP", "NVL", "ABCG1", "COL8A1", "CTDSPL") characte
18 c("BEX4", "SMARCD3", "EFNB3", "INPPL1", "NEO1", "LAMB1", "DUSP2", "CNN3", "IVNS1ABP", "PAK1
FAM162A, "PTOV1") c("OLFML2A", "ITGAE", "ALDH3A1", "AZGP1", "GPX7", "C3", "SCGB2A1", "LONP2",
19 c("STUB1", "SLC4A4", "RUNX1", "HMOX1", "PAK1") c("MEOX2", "ITGAE", "GNPDA1", "RRAGD", "CDII
20 c("APOD", "SCPEP1", "SNAP25", "GABRP", "CNN3", "PIR", "IVNS1ABP", "GPC3", "LIMCH1", "BEX4", "S
21 c("RUNX1", "ABLIM1", "TOMM20", "SATB2", "SCPEP1", "VAT1", "DDX42", "ADAP1", "PPL", "LIMCH1"
22 c("SLC4A4", "IL17RB", "GABRP", "CNN3", "TMBIM6", "OGT") c("STAT5B", "SLC1A3", "NPDC1", "EIF4B
23 c("IL17RB", "HMOX1", "SOSTDC1", "LASP1", "TNNC1", "APOD", "GLRX", "VAT1", "SATB2", "CX3CL1",
CELSR2, "LGR4", "GOLM1", "C2CD2", "DHRS2", "MSX1", "PKP1", "ATRX", "CYB561", "WNT5A", "ITIH5", "
ZNF281, "SEMA6A", "GNB5", "GSTM3", "HLTF", "RAD51C", "HSD17B11", "AFF1", "MAP2K6", "FOXO3", "
CSGALNACT1, "KHDRBS3", "IDE", "ST14", "BMP4", "ISLR", "SERPINF1", "CP", "BCAT1") c("CD200", "RTN3
SLC1A5, "PSMD10", "POLR2K", "ADI1", "PSMC4", "UTP6") c("TCEA2", "CXCL12", "SFRP1", "SCG5", "ITM2
FGF9, "ZNF232", "PKIA", "TFDP2", "BMP4", "AUTS2", "AEBP1", "ISLR", "SERPINF1", "FGFR2") c("FKBP1B"
GSTM3, "PARM1", "RET", "RGS5", "SERPINB9", "GBP1", "NCL", "SULF1") c("MGLL", "SHCBP1", "BIRC3", "
ABCC3, "SIRT3", "PKP1", "RET", "ERAP2", "CAPRIN1", "CSF3R", "PUF60", "SERPINB9", "GBP1", "CP", "CX
STXBP6, "SERPINB9", "CXCL10") c("MGLL", "ZNF318", "CXCL12", "PRSS23", "LDB2", "CKS1B", "CTDSPL",
PSMD11, "CXCL10") c("MGLL", "PELI1", "PRSS23", "CHORDC1", "NFAT5", "NTRK2", "PCMT1", "CORO1C"
ZSCAN18, "IFIT1", "SEMA6A", "HBB", "MSX1", "ALDH1A2", "EPHB2", "MYH11", "SLC22A5", "SLCO2A1", "
ABCG2, "DKK2", "KHDRBS3", "PCYOX1", "MS4A6A", "PSMC3") c("MGLL", "SIAH2", "PRSS23", "LGALS1",
KCNQ2, "LGALS3BP", "SIRT3", "ANXA1", "NOTCH1", "ZSCAN18", "IFIT1", "SERPINB9", "NXF1") c("NPY1R"
FCGR2A) c("MGLL", "TFF1", "PELI1", "HSP90AA1", "PIP4K2B", "VAT1", "RAP2A", "DOCK10", "RRP8", "SN
CD34, "ATF5", "COL6A2", "ABCC3", "KCNQ2", "FSTL1", "CDH2", "DACT1", "IFIT1", "FCER1A", "AEBP1", "I
IGFBP3, "SERPINB9", "NXF1", "CXCL10") c("TFF3", "NCALD", "PRSS23", "RBBP8", "BIRC3", "FXYD2", "SPA
FERMT2, "SNAP25", "IFIT1", "RPS6KA1", "OXA1L", "ADI1", "SERPINB9", "PSMD11", "CXCL10") c("MGLL",
MYH11, "TNFSF10", "WSB2", "CDS1", "NR2F2", "PGK1", "TMEM47", "SLCO2A1", "MAP9", "PDHX", "RGS
PKP1, "TMEM47", "RET", "CAPRIN1", "IGFBP3", "SERPINB9", "NR2F1", "CXCL10") c("CIT", "CXCL12", "M
RAD51C, "CAPRIN1", "SERPINB9", "PRMT5", "NGDN", "CXCL10") c("PELI1", "FARP1", "VAT1", "RAP2A", "
ZSCAN18, "TCTN1", "SEMA6A", "NRN1", "STAT5B", "MYH11", "PARM1", "NR2F2", "TMEM47", "OLFML2
MYH11, "RAD51C", "CANX", "TMEM47", "RET", "AGL", "RGS5", "HLA-DPA1", "SERPINB9", "CD14", "NR2

24 c("TSC22D1", "SLC4A4", "GOLM1", "ABLIM1", "CD99", "GLRX", "BAG2", "VAT1", "UNG", "QDPR", "M
TBC1D15, "MPZL2") c("SPC25", "GRWD1", "PRIM2", "NSL1", "USP13", "AARS", "FN1", "LAMA4", "LGALS
25 c("IL15RA", "RBM6", "SORT1", "SLC4A4", "ENPEP", "GALNT11", "APOD", "BEX4", "IVNS1ABP", "BMI1
CCNG1, "RNF128", "STUB1", "CKB", "BCHE", "MXI1", "CAST", "UBR5", "XIST", "ACAA2", "LDB3", "FGGY",
PRKACB, "NDRG2", "FBP1", "HLTF", "MAP2K6", "ZNF395", "CIRBP", "RNF43", "HMGCS2", "RHOB", "COB
FBN2, "COL6A1") c("LTBP1", "L1CAM", "NRP1", "PYCR1", "SCUBE2", "ZNF83", "LPAR1", "MGLL", "IFIT1",
PTPRD, "EMP3", "TPSB2", "COL6A1") c("PUF60", "LTBP1", "L1CAM", "ABCC3", "RBM6", "NRP1", "PCLO"
RCAN2, "HIST3H2A", "AHCY", "PEG3", "BTAF1", "TPM2", "PTPRD", "NRXN3", "TFPI") c("C7", "LTBP1", "T
OPHN1, "NRXN3", "TFPI") c("C7", "PODXL", "SLIT2", "SEMA6A", "MYH11", "DYNLRB1", "L1CAM", "ABCC
26 c("GOLM1", "SATB2", "BAG2", "SLC4A4", "PIR", "CST6", "CD99", "SMARCA2", "TPCN1", "STUB1", "RN
ANGPT1, "EFHC2", "NEBL", "XIST", "NDP", "NUDT11", "SESN1", "DHTKD1", "WIF1") c("SCGB2A1", "UBE2
RASAL1, "RHOBTB1") c("NT5E", "MGLL", "FTO", "LGALS1", "SERPINB9", "TFPI", "GALNT2", "CRIP1", "SAR
27 c("CNN3", "ZNF586", "CTDSP2", "GABRP", "APOD", "HMOX1", "PTPN3", "CALML3", "VPREB3", "SNAP
28 c("HMOX1", "DUSP2", "BLCAP", "ZNF586", "GABRP", "IVNS1ABP", "PIR") c("ID2", "FOXJ3", "THAP11"
29 c("APOD", "BEX4", "GABRP", "DUSP2", "WDR91", "IL17RB", "RBM6", "SQSTM1", "PIR", "PLXNA2", "N
GOLM1, "MOAP1", "CUTA", "STUB1", "UBE2Q1", "MLH3", "ERMP1") c("SCAND1", "FCGBP", "ARID1A", "
KIAA0232, "FAM13B", "ZBED5", "LRRC17", "MT1F", "LMO4", "CYP4B1", "SH3BGRL", "GOLM1", "PBX1",
ACOT9, "UGDH", "CADM1", "LRRC17", "INPP4B", "LMO4", "SPINT1", "RFX5", "CPA3", "ABCG2", "PODXL"
MT1F, "GDPD5", "FKBP11", "PHF10") c("TFF1", "SEMA4C", "TRIB2", "NUPR1", "ARL2", "NT5C", "SLC35A
PHF10) c("SFRP1", "TGFBR3", "UCHL1", "RGS5", "SPEN", "ARMCX1", "MGLL", "AHCY", "PSMD12", "SLC4
MAPKAPK5, "HLA-DRA", "CXADR", "RAMP1", "QPRT") c("SFRP1", "TGFBR3", "A2M", "TFPI", "SPEN", "TG
30 c("IL17RB", "TSPAN6", "CACNA1D", "GUSB", "CD99", "DUSP2", "UNG", "APOD", "ORAI3", "WDR13", "
ISL1, "ZBTB5") c("AKR7A3", "E2F2", "OLFML2A", "NDUFB6", "SKP2", "CLNS1A", "NDUFA4", "TNFSF10", "
31 c("DDX42", "GLRX", "FAM53C", "CSK", "PTPN3", "CALML3") c("FOXJ3", "TBP", "TAF5", "SMPDL3A", "
32 c("CALML3", "CNN3", "BEX4", "HMOX1", "QDPR", "LAMB1", "RND3", "PIR", "VAT1", "DUSP2", "CST6"
MEIS2, "PHACTR2", "PLXNB1", "FAM162A", "AGAP1") c("AZGP1", "SPRR1B", "SPRR2B", "ZNF512B", "OL
33 c("QDPR", "OPTN", "VPREB3", "HMOX1", "PRKACB", "CNN3", "PIK3IP1", "TTC3") c("NVL", "MT1M", "
34 c("PDZD2", "ABLIM1", "SLC24A3", "BCAS1", "GPC3", "CNN3", "BAG2", "ADCY6", "ENPEP", "RND3", "
35 c("STUB1", "PAK1", "TOMM20", "IL17RA", "DDX42", "GPC3", "RUNX1", "SMARCA2", "SATB2", "SNAP
SLC2A10, "ANG", "EIF3H", "HBP1") c("TRIM22", "CLNS1A", "ACSL1", "AZGP1", "SPATA20", "TCTA", "GRH

'V19" "V20" "V21" "V22" "V23" "V24" "V25" "V26" "V27" "V28" "V29" "V30" "V31" "V32" "V33" "V34" "
'DES", "RAB25", "ZC3HAV1", "SAC3D1") c("RAB25", "CAV2", "TIMM8B", "SLC7A5", "PRRG1", "SSH3") c("
, "HMOX1", "LMBRD1", "ORAI3", "PIK3IP1", "RNF103", "SALL2", "SLC24A3", "SLC4A4", "SMARCA1", "SM
3", "C4BPA", "CHGA", "CLNS1A", "ENPEP", "FAM117A", "FBXO21", "FCGBP", "GNPDA1", "HMGB2", "ID2
'UNG") c("AKAP12", "ARMCX2", "ARMCX3", "BST2", "C1S", "C7", "CLIC2", "CTNNB1", "CXCL10", "DAZAP
H5A1", "QDPR", "TNNC1", "PDZD2", "PLXNA2", "IL17RB", "GABRP", "RND3", "LIMCH1", "TSC22D1", "SOS
", "RND3", "ENPEP", "CNN3", "CCDC28A", "SIDT2", "SOSTDC1", "EFNB3", "CX3CL1", "APOD", "GLRX", "G
) c("NFATC3", "COMP") c("DEGS1", "CRYAB", "DDAH1", "SSPN", "ZNF91", "CUEDC1", "ELF5", "AGAP1", "
D", "MT1M", "PLLP") c("NFATC3", "LGALS3", "SLC36A1", "CHRNA3") c("MT1G", "RBBP8", "GATM", "HOX
(6", "TMPRSS3") c("CAST", "TPD52", "ESR1", "XIST", "CLU", "ANGPT1", "CXCL14", "CRNN", "DACH1", "M
25", "ALDH9A1", "VAT1", "PAK1", "PRKACB", "CACNA1D", "LIMCH1") c("ID2", "ITGAE", "AZGP1", "RAD51
", "HMOX1", "SQSTM1", "IL20RA", "ARHGEF2", "CX3CL1", "CNN3", "LIMCH1", "LAMB1", "DGKZ", "PRKA
MT1G", "PDCD4", "IRX5", "PRTN3", "RIMS3", "TCEAL2", "MBTPS1", "DHTKD1", "MUC5AC", "VAPB", "S10
CL1", "CHPT1", "COL14A1", "RPS6KA2", "SORBS3", "KRT1", "RNASE1", "ASTN2", "USP7", "GATAD2A", "H
F", "ESM1", "SPRY4", "RNASE1", "NDN", "SUSD5", "SERPINA3", "STC1", "TUBB3", "PLAUR") c("WIF1", "C
0", "SLC25A32", "LSM1", "NDN", "ENAH", "NDUFA7", "RANBP1") c("PDZD2", "CDH2", "ALDH5A1", "CXC
X1", "TFCP2L1", "MAST2", "GPC3", "ICAM3", "RHOB", "COX15", "DACH1", "NDN", "SALL2", "HMGCS2", "
ET", "AKAP12", "PRSS23", "MGLL", "HERC5", "BIRC3", "MCM2", "ZNF337", "IFIT1", "PCOLCE2", "COCH",
)81", "EDNRA", "SPARCL1", "GLI2", "MRPL18", "PDE8A", "SLC25A13", "SYCP2", "LRRC15", "PELI1", "RAN
, "MX1", "OAS2", "CST4", "IRS1", "SERPINB2", "SRGN", "STAT4", "MICAL2", "VPS37B", "DLAT", "KCTD5",
27L", "GRAMD1B", "AHSA1", "SPEN", "ZDHHC11", "ACAT1", "PODXL", "BET1", "MYC", "RRS1", "TEX10",
LA", "RPS6KA2", "COL4A6", "PDE8A", "RANBP1") c("SNAP25", "ZMAT3", "RPS27L", "PRSS23", "MGLL", "L
1B1", "WIF1", "OLFML3", "PON3", "SERPINB9", "MST1", "SFRP4", "LGALS1", "DNAJC10", "TSPAN7", "DIR
5", "MKI67", "MCM2", "SPEN", "IFIT1", "HSD11B1", "DSC3", "SERPINB9", "LY6D", "SLC39A14", "USP18",
T3", "PRSS23", "MGLL", "HERC5", "ZNF337", "LOR", "SPEN", "IFIT1", "HSD11B1", "SERPINB9", "NTRK2",
c("RET", "ALOX5", "CDH2", "HERC5", "HLA-DOB", "IFIT1", "IFI44L", "IFI6", "FSTL1", "HSD11B1", "IFI44", "
DIRAS3", "GSTO1", "CDH11", "MCOLN1", "PAEP", "BUB1", "NQO2", "TOX", "ZBTB16", "RUFY3", "EXOSC8
T1", "CYTIP", "GSTO1", "RRS1", "AGPAT5", "GRWD1", "IQCG", "OSBPL3", "MED27", "MTMR2", "HSP90A
, "SYCP2", "ELOVL5", "PELI1", "ADAM28") c("TFF3", "NCAM1", "PRSS23", "BIRC3", "ALDH3A1", "TGFBR3
"MGLL", "FANCL", "ZNF337", "SPEN", "IFIT1", "GNG11", "IGFBP6", "PCOLCE2", "FSTL1", "EFHD1", "OLF
'TBC1D1", "IVD", "RGS5", "SLIT2", "EDNRA", "ZEB2", "SPARCL1", "GLI2", "RCAN2", "CPT1A", "LAPTM4B",
23", "MGLL", "HERC5", "SPEN", "IFIT1", "HSD11B1", "SERPINB9", "FGF13", "DNAJC10", "DIRAS3", "GSTO
MGLL", "LOR", "SPEN", "IFIT1", "GAMT", "HSD11B1", "RCN1", "SERPINB9", "CGA", "KIF18A", "DNAJC10",
"CDH2", "CLIC5", "RPS27L", "AKR7A2", "CYBA", "POP5", "SDC3", "GNG11", "MMP9", "ESR1", "PTGER3",
M4B", "CADM1", "ICAM3", "ADIPOQ", "COL4A6", "SSX1", "SCO2", "GLIPR1", "UCK2") c("NR3C1", "PRSS2
", "ENG", "ZNF423", "CTBP2", "RPS6KA2", "RNASE1", "PDSS2", "APOC1", "TMEM176B", "NOTCH1", "GM
", "ESM1", "FOXF1", "CCL21", "TOX3", "IGFBP5", "SYCP2", "INHBB", "PELI1") c("NR3C1", "TFF1", "RPS27
", "CPT1A", "FOXF1", "CCL21", "FCER1A", "COL4A6", "DPT", "PDSS2", "COL9A3", "ELOVL5", "GMFB", "SE
"IFI6", "PCOLCE2", "HSD11B1", "PTGER3", "ZDHHC11", "SERPINB9", "MST1", "DNAJC10", "GSTO1", "CLI
2", "COL4A6", "RNASE1", "SYCP2", "INHBB", "PELI1") c("TFF3", "ZMAT3", "RBBP8", "PRSS23", "FAM129
L", "HSD11B1", "IFI44", "COL5A2", "PTGER3", "ANXA6", "CHSY1", "COL4A1", "TFAP2C", "JMJD6", "FKBP1
1A", "COL4A6", "DPT", "PDSS2", "SUSD5", "SSX1", "IL1B", "GMFB", "PSMA3") c("SNAP25", "ZMAT3", "N
"PBK", "MKI67", "DNAJC12", "LOR", "SPEN", "IFIT1", "IFI44L", "HSD11B1", "RCN1", "PON3", "SERPINB9",
, "ELOVL5", "SPHK1") c("SNAP25", "ZMAT3", "PRSS23", "MGLL", "ALB", "MRPS16", "FANCL", "LOR", "SPE
39A6", "NR3C1", "DIP2C", "IFIT1", "IGFBP6", "IFI44L", "FSTL1", "HSD11B1", "PTGER3", "ANXA6", "TFAP2
") c("PCNA", "SNAP25", "ZMAT3", "TFF1", "SNCA", "DIP2C", "GGH", "LOR", "PCOLCE2", "CCL19", "OLFM
FI27", "DDB2", "SHCBP1", "ALB", "DIP2C", "IFI44L", "FABP4", "IFI6", "FSTL1", "HSD11B1", "IFI44", "GLRX"

ST1", "MATN2", "HAPLN1", "NBEA", "B2M", "RGN", "RGS5", "RAMP1", "PGK1", "ZEB2", "FZD6", "S100P",
1", "EDNRA", "PDE4A", "STEAP3", "DPY19L2P2", "NR2F1", "RCAN2", "ESM1", "PKP1", "PROX1", "PPP2R
"CLMN", "RANBP1") c("SNAP25", "ZMAT3", "RNF128", "HERC5", "ZDHHC11", "CD83", "DLAT", "C7", "TB
GFBR3", "ZNF337", "SFRP1", "PCOLCE2", "OLFML3", "PON3", "TFPI", "SLC47A1", "MST1", "PEG3", "VIM"
"VWF", "H1F0", "NDN", "SDHA", "PNMA1") c("DDB2", "NCAM1", "RET", "RAD51C", "TGFBR3", "EFHD1",
RNF128", "MGLL", "LOR", "SPEN", "GAMT", "HSD11B1", "KIF18A", "SCML1", "DNAJC10", "CKS1B", "GRW
BB", "ZHX2", "SLC24A3", "NEO1", "LIMCH1", "PRKACB", "SLC4A4", "STAU2", "SATB2", "SMARCA2", "QD
"RNF103", "OGT", "VAT1", "TP53I3", "ANAPC13", "SLC24A3", "TSC22D1", "SNAP25", "TTC3", "CNN3", "
", "OXA1L") c("GNPDA1", "ALDH3A1", "RAB25", "TRIM22", "CLNS1A", "IGFBP2", "OLFML2A", "BTG1", "A
", "PRKDC", "OGT", "SHOX2", "AHCY", "HMGCS1", "SCUBE2", "SUSD4", "RNF128", "GOLGB1", "RRAGD",
MTHFD2", "SHMT1", "IFIT1", "SCG5", "IFITM1", "SMAD2", "MXRA5", "PLTP", "OGT", "ABCA3", "SLC29A1
CO3A1", "DDX42", "DACH1", "ACVR1B", "HOXA10", "SNAPC1", "ERP44", "UPF3A", "FKBP1B", "PTPN3", "
"SNAP25", "GOLM1", "ACE2", "IL20RA", "DUSP2", "UBL3", "PAK1", "GABRP", "PLXNA2", "GUSB") c("RRA
"PIR", "STUB1", "PRKACB", "TSPAN6", "CNN3", "TOMM20") c("BTG3", "KCNJ2", "RGS4", "CBX4", "CXADR
"IL17RA", "PLXNA2", "TNNC1", "CNN3", "CX3CL1") c("GAS1", "WNT5A", "RGS4", "MEIS2", "ACSL1", "DST
H5", "CNN3", "ZNF263", "SIDT2", "GOLM1", "SMARCA2", "HMOX1", "SLC4A4", "ZNF211", "GABRP", "CD
") c("CLIC4", "CXCL10", "RGS5", "KCNMB1", "SORBS1", "ZDHHC11", "ACTN1", "TOX3", "HSD11B1", "GM2
ZDHHC11", "COL4A6", "CD34", "NQO1", "HSD11B1", "USP13", "SIRT3", "NRP1", "LPAR1", "GM2A", "LGA
er(0) c("XIST", "ARMCX2", "DEGS1", "MFAP2", "ENO2", "ESR1", "FGFR2", "IRX5", "DDAH1", "ZFP36L1", "
", "BAG2", "ZNF586", "PIR", "ZNF211", "VPREB3", "RPRD2", "IL20RA", "TPCN1", "DGKZ", "VGLL1", "LIMC
"CYB561", "BTG1", "TNFSF10", "FCGBP", "IGFBP2", "TRIM22", "ID2", "BCL11B", "PYCARD", "IKZF1", "TB
PT", "BTG1", "WNT5A") character(0) c("STUB1", "CSTA", "CXCL14", "GATM", "MGLL", "HBP1", "COG5", "
SATB2", "HMOX1", "CX3CL1") c("VSNL1", "AZGP1", "RIMS3", "TRIM22", "ALAD", "DES", "ACSL1", "MEIS2
", "STUB1", "RND3", "CX3CL1") c("SLC1A3", "EIF4B", "VSNL1", "GAS1", "THSD7A", "TXNIP", "NFIB") c("PR
") c("MYCN", "DIO1", "NPDC1", "IGFBP1", "SAP30") c("MSX1", "AGAP1", "WBP4", "XIST", "UPF3A") c("E
"SQSTM1", "ABLIM1", "CHN2", "QDPR", "IL20RA", "STAC", "PTPN3", "BAG2", "LAMB1", "RND3", "GOLM
"PADI2", "OXA1L", "ISL1", "SNAPC1", "FGFR2") c("ZNF318", "FDPS", "C4BPA", "MCM2", "NUP43", "HMG
"SNRPD3", "CSDE1", "NINL", "NUAK1", "DCLRE1C", "ISL1") c("HYOU1", "CTSC", "TSPAN4", "SFRP1", "MC
3", "PAQR4", "WIF1", "OIP5", "PLA1A", "FZD10", "SLC30A1", "BSPRY", "OGFOD1", "ITGAE", "SCML2", "N
2C", "ESR1", "ALDH1A1", "SLCO4C1", "SOSTDC1", "GNE", "MT1F", "LGALS1", "CTSH", "DEGS1", "JAM3", "
", "GPSM2", "PLA1A", "HMGCS1", "UCHL1", "EBP", "ARL5A", "CGA", "PRIM2", "STEAP3", "SUSD4", "OGF
"CCDC92", "CCL18", "NUP43", "ANXA2P2", "RBPJ", "FKBP11", "IFI44L", "SPARCL1", "GALNT2", "GLIPR1",
CL10", "BCAT1", "UTP6") c("SMURF2", "HSP90AA1", "HSPA8", "CORO1C", "QKI", "CENPM", "MLXIP", "PC
"PIP4K2B", "CORO1C", "VAT1", "HHEX", "SATB2", "UCHL1", "EBP", "SNHG3", "SPC25", "LOR", "DNAJC10
", "VAT1", "PLA1A", "SATB2", "HMGCS1", "EBP", "SRPK1", "UBIAD1", "SPC25", "LOR", "DNAJC10", "NMU
"RGS5", "SERPINB9", "CP") c("MGLL", "PELI1", "PRSS23", "SCG5", "FARP1", "PIP4K2B", "ABCA3", "VAT1",
"CORO1C", "ANXA5", "OPN3", "GNAI2", "ELOVL5", "VAT1", "CALR", "CCDC88A", "RRP8", "PPIF", "CLIC4",
", "PELI1", "GNG11", "IGFBP4", "FARP1", "AXL", "CILP", "LAMC1", "FN1", "AKR1B1", "OPN3", "GNAI2", "
HG3", "SPC25", "NR3C1", "STEAP3", "OGFOD1", "RASSF1", "SSX1", "SHOX2", "HERC5", "MYBPC1", "IFIT1
MSMB") c("TFF1", "ZNF318", "NFAT5", "CUEDC1", "PLA1A", "TKTL1", "BET1", "RDH11", "NR3C1", "RASSF
TS2L", "PCMT1", "CCDC92", "MMD", "NFIC", "SDC2", "SERPINE2", "UCHL1", "NQO2", "TAGLN", "TUBB2
, "PELI1", "PRSS23", "HSP90AA1", "FARP1", "CORO1C", "PMP22", "VAT1", "PLA1A", "CCDC88A", "LRIG1"
5", "HLA-DRA", "RGS1", "CD14", "NR2F1", "SERPINF1") c("MGLL", "NCALD", "PELI1", "PRSS23", "SFRP1",
CAM", "FDFT1", "LSM2", "AGPAT5", "ABCA3", "RHOC", "MUC2", "TNC", "RDH11", "RFWD3", "TMEM147
"NUDT11", "DOCK10", "TKTL1", "ZDHHC11", "UBIAD1", "OPHN1", "CCNE2", "CD83", "HERC5", "SNAP25"
A", "SLCO2A1", "RGS5", "NR2F1", "SERPINF1") c("NCAM1", "NCALD", "FARP1", "FBN2", "FADS1", "CILP",
F1", "FCGR2A") c("MGLL", "S100P", "MELK", "CKS1B", "CORO1C", "TXNRD1", "MUC1", "PLAT", "SRPK1",

SL1", "DYNC1H1", "CCDC28A", "ZNF586") c("KIF2C", "OIP5", "CDCA3", "MT1H", "MT1M", "EED", "MCM2
1", "PPID", "FTO", "IFIT1", "PYCR1", "A2M", "TIMP3", "IKBKE", "GSTO1", "NT5E", "VAT1", "STEAP3", "MA
", "GABRP", "ATM", "COL7A1", "GRSF1", "STUB1", "SNAP25", "IL17RB", "PIK3IP1", "SEMA5A", "PRKACB"
"KCTD2", "CITED2", "PPP1R3C") c("NDUFA4", "SCGB2A1", "C3", "GRAMD1C", "LIMK2", "LSM4", "SAMM
LL1", "KCTD2", "JAM3", "RGL1", "GSDMB", "KIAA0232", "CYP2J2", "ACE2", "COX15", "GCC2", "PSD3", "G
"IFITM3", "TMEM63A", "IFITM2", "RUNDC3B", "IVD", "SELENBP1", "CLNS1A", "ZDHHC11", "MXRA5", "E
, "ZNF83", "LPAR1", "MGLL", "IFIT1", "IFITM3", "DMXL2", "SORBS2", "RUNDC3B", "IVD", "DYNLT3", "MR
GFBR2", "EXT1", "ABCC3", "NR2F1", "NRP1", "PYCR1", "SMAD2", "LGALS1", "DAB2", "TCTN1", "LPAR1",
3", "PMP22", "NR2F1", "YTHDF1", "SERPINE2", "DPYSL2", "SMAD2", "LPAR1", "AHR", "ZSCAN18", "QKI",
ND3", "LIMCH1", "TNNC1", "GABRP") c("POMZP3", "DES", "CYB561", "SPRR2B", "GALNT2", "SLC25A46", "
2E3", "NDUFS8", "ABCG2", "FCGBP", "NDUFC1", "SAMM50", "HIGD2A", "NUP43", "TPRKB", "FDPS", "CYB
R1B", "PKP1", "IFITM3", "TGFBR2", "NQO1", "THSD7A", "NCALD", "SF1", "HSD11B1", "DPP8", "ZNF580",
P25", "IVNS1ABP", "DYNC1H1", "BAG2", "MBTPS1", "SATB2", "PAK1", "ITCH", "FGFR4") c("SLC1A3", "KIF
, "CHKA", "TBP", "RGS4", "TCTA") c("NFATC3", "LPCAT4", "MYCN") c("ESR1", "JAG1", "CDKN1B", "CRISP3
IECAP2", "SLC24A3", "HMOX1", "SASH1", "CALML3", "HAUS4", "MED23", "UVRAG", "BLCAP", "SEMA5A"
"RGS4", "ALDH3A1", "URM1", "AZGP1", "HIGD2A", "SEC61B", "NDUFS8", "FKBP2", "DCTPP1", "POLR2G",
"PSIP1", "SMARCA2", "MAST2", "ZNF589", "QPRT", "ERMP1") c("UCHL1", "SPEN", "AHCY", "TLE1", "GPS
", "SNAP25", "KLHL21", "ZFP64", "NOTCH3", "MNT") c("MGLL", "RBMS1", "SIX2", "CPT1A", "ABCC3", "PF
1", "SOX4", "IRF4", "MED23", "SAR1B", "IFNGR1", "LPAR1", "TBX3", "PLA2G15", "TM4SF1", "ZKSCAN1", "
7A1", "SIX2", "ABCC3", "PRSS23", "COX8A", "OPHN1", "TLE1", "LRIG1", "IRF4", "TDP1", "AHR", "MYLIP",
FBR2", "MGLL", "SLC47A1", "ABCC3", "TIMP3", "NT5E", "PRSS23", "COX8A", "ITGB1", "RET", "STAP2", "
"ENPEP", "BEX4", "SCPEP1") c("TM7SF2", "FOXJ3", "THAP11", "MX2", "EED", "TMEM14A", "GLOD4", "W
TMEM14A", "SLC37A4", "ALDH3A1", "TXNL4B", "MCM2", "SRPK2", "GNPDA1", "GPX7", "C4BPA", "BTG1
NVL", "BTG1", "GLOD4", "SHROOM2", "URM1", "BANP", "NFYC") c("EFNA1", "NFATC3", "BANP", "NPC1"
", "NEO1", "APOD") c("AZGP1", "VSNL1", "SPRR2B", "ALDH3A2", "MT1H", "RRAGD", "TMEM14A", "CXAD
FML2A", "RRAGD", "FBXL5", "TMEM14A", "FCGBP", "IKZF1", "SCAND1", "POLR2G", "GPX7", "PYCARD", "
BTG1", "TRIM22", "BTG3", "MT1H", "ACSL1") c("HBD", "ALAS2", "TRIM22", "TCL1A", "IP6K2") c("MT1G"
RKACB", "ADH5", "UVRAG", "GABRP", "STUB1", "LIMCH1", "TTC3", "FBXO34", "HAUS4") c("CST3", "COF
25", "APOD", "ABLIM1", "PIR", "CST6", "TNNC1", "GABRP", "CNN3", "PLXNA2", "HAUS4") c("TRIM22", "
IL2", "UQCRB", "IGFBP2", "TRPM4", "ABCG2", "PCTP", "ACAD8", "SPRR2B", "RGS4", "NDUFA4", "GPX7",

V35" "V36" "V37" "V38" "V39" "V40" "V41" "V42" "V43" "V44" "V45" "V46" "V47" "V48" "V49" "V50" "V
MEIS2", "ZNF512B", "HOXA10", "KIAA0355", "WIF1", "NEBL", "SLCO3A1", "LAMA5", "LDB3", "IRX5", "C2
ARCA2", "SORT1", "TSC22D1", "TSPAN6", "UNG") c("APOL2", "CCDC28A", "CXADR", "DDX58", "EED", "FI
", "IKZF1", "NCOA2", "NDRG2", "NUP43", "OLFML2A", "PRKDC", "RAB25", "SCGB2A1", "TNFSF10", "TRIN
2", "EDNRA", "EPS8", "GRK5", "HERC5", "HHEX", "IFIT1", "IL1R1", "LAPTM4B", "OXA1L", "PMM1", "RGS5
TDC1", "DUSP2") c("FBXO21", "RAD51C", "ITGAE", "MT1H", "NMT1", "WNT5A", "KIF2C", "POP7", "OIP5
PC3") c("RRAGD", "POMZP3", "TXNIP", "COX7C", "RERE", "CCDC28A", "EIF4B", "RBBP4", "FOXJ3", "RTN3
ASAP3", "PTOV1", "PHYH", "TNNT3") c("NVL", "GPX7", "PSAT1") c("ASAP1", "B3GALNT1") c("RCBTB2", "
A10", "FKBP1B", "CAST", "MGLL", "GULP1", "PTPN3", "MUC5AC") c("RRAGD", "TSPAN7", "KATNB1", "UI
GLL", "PRDX6", "TSPAN13", "CRCT1") c("ACSL1", "C3", "ITGAE", "SCGB2A1", "DHCR24", "SPRR2B", "GDE1
LC", "MT1M", "BRD8", "ABCB6", "ACSL1", "TMEM14A", "CST3", "SLC1A3", "TBCB", "DNASE2", "ANKMY2
B", "IVNS1ABP", "SATB2", "UNG", "OPTN", "PAK1", "STAU2", "MBTPS1", "C5", "SCPEP1", "TMBIM6", "T
A8", "RNF10", "ACPP", "PEBP1", "GOLM1", "PER3", "CHPT1", "FLG", "DDX42", "PKP1", "KLHDC2", "ENO
SPA6") c("SLC39A6", "PRSS21", "C3", "NOVA1", "PTP4A1", "CLIC5", "OLFM1", "SFRP1", "NNMT", "TSPAN
200", "FZD10", "SLC30A1", "ITGAE", "OIP5", "LSM4", "GTF3A", "POSTN", "NEFM", "NR2F6", "AKAP8", "
L14", "CXCL12", "SFRP1", "ESR1", "BCL2", "CALML3", "SPINK2", "GLRX", "NUCB2", "FCGRT", "DEGS1", "H
MLLT11", "MCF2L") c("BCHE", "FAS", "RNF128", "RAD51C", "SPEN", "KRT10", "SERPINB9", "CGA", "KIF1
"ITM2A", "HSD11B1", "ZDHHC11", "SERPINB9", "NUCB2", "FGF13", "MST1", "USP18", "GPM6B", "SQSTN
BP1") c("SACS", "DDB2", "ZMAT3", "RET", "CDH2", "SHCBP1", "MGLL", "BIRC3", "DIP2C", "IFI44L", "IFI44
"KPNA4", "OLR1", "DLEU1", "RNASEH2A", "ABCE1", "SAMSN1", "CXADR", "SNX10", "CXCL11", "USP6NL"
"GRWD1", "DLAT", "HSP90AA1", "SIRT3", "SATB2", "CORO1C", "PUF60", "DSTN", "TIAM1", "CYC1", "LYZ
OR", "SPEN", "IFIT1", "CXCL12", "HSD11B1", "SERPINB9", "HSPA13", "DNAJC10", "GSTO1", "CKS1B", "PC
AS3", "GSTO1", "PMP22", "PCOLCE", "GRWD1", "LPAR1", "PTGDS", "MED27", "HSP90AA1", "NELL2", "N
"FKBP11", "SLC1A4", "DNAJC10", "CCDC47", "DIRAS3", "GSTO1", "TRIP13", "RFWD3", "CKS1B", "PCOLC
"DNAJC10", "DIRAS3", "GSTO1", "NMU", "NDRG4", "GRWD1", "DLAT", "YRDC", "CHORDC1", "SPC25", "N
PTGER3", "GMFG", "CPVL", "ZDHHC11", "FGF13", "MST1", "IGFBP4", "FKBP11", "HSPA13", "LGALS1", "T
", "SLCO2A1", "PRKCA", "XAF1", "CCDC59", "TPSB2", "PCP4", "COL6A1", "ABCA3", "IFITM2", "APP", "PXI
A1", "SPC25", "MACF1", "PARP8", "LGALS3BP", "CALR", "CXCL10", "IFIT5", "SATB2", "MRC2", "IFIT2", "FA
", "PON3", "TFPI", "CGA", "PEG3", "MFAP5", "HSPA13", "MUC5B", "TSPAN7", "GCLM", "DIRAS3", "MEF2
ML3", "SERPINB9", "MST1", "SERPINE2", "MFAP5", "SFRP4", "DNAJC10", "TSPAN7", "DIRAS3", "GSTO1", "
"ANGPTL2", "RNASE1", "TCF12", "PDSS2", "NOTCH1", "MUC2", "PNMA1", "MSR1", "LRRN3") c("SNAP2
1", "PCOLCE", "GRWD1", "LPAR1", "OSBPL3", "HSP90AA1", "SIRT3", "PRIM2", "NELL2", "USP13", "LGALS
"GSTO1", "PCOLCE", "GRWD1", "LPAR1", "ME1", "DLAT", "MTMR2", "HSP90AA1", "PRIM2", "NELL2", "I
"CPVL", "ANXA6", "ZDHHC11", "CHSY1", "NUCB2", "PODXL", "DPYSL2", "DNAJC10", "BET1", "FBLN5", "D
", "MGLL", "FANCL", "PPIF", "LOR", "IFIT1", "GAMT", "IFI6", "HERC6", "FSTL1", "HSD11B1", "IFI44", "PTG
FB", "PELI1") c("RET", "AKAP12", "RPS27L", "HERC5", "CDV3", "CDH6", "DIP2C", "PPIF", "NPY1R", "IFIT1"
L", "MGLL", "HERC5", "SPEN", "IFIT1", "SERPINB9", "RRP8", "CSTA", "GRWD1", "ME1", "HSP90AA1", "SP
RPINB5", "SCO2", "HSPA6") c("NR3C1", "ZMAT3", "TFF1", "HERC5", "NDUFA4L2", "SRD5A1", "BET1", "TN
C4", "OPN3", "PCOLCE", "GRWD1", "LPAR1", "ME1", "MTMR2", "SIRT3", "SPC25", "PRIM2", "MACF1", "I
", "BIRC3", "TGFBR3", "ZNF337", "RCN1", "CGA", "SERPINE2", "SLC7A1", "MFAP5", "HSPA13", "MUC5B"
1", "LGALS1", "HSP90B1", "CD34", "MYC", "CLIC4", "RASIP1", "NOC3L", "OPN3", "CKS1B", "TEX10", "CILI
CAM1", "PRSS23", "MGLL", "FANCL", "TGFBR3", "ZNF337", "LOR", "SPEN", "IFIT1", "SFRP1", "PCOLCE2",
"DNAJB2", "KIF18A", "IRS1", "LGALS1", "HOPX", "DNAJC10", "CCDC47", "DIRAS3", "GSTO1", "CLIC4", "C
N", "IFIT1", "ISG15", "HSD11B1", "IFIT3", "SERPINB9", "MST1", "DNAJC10", "DIRAS3", "GSTO1", "PMP22
C", "JMJD6", "FKBP11", "LGALS1", "BET1", "ISLR", "CLIC4", "OPN3", "CKS1B", "MED27", "CKB", "BRD2", "
3", "SERPINB9", "NUCB2", "NDUFA4L2", "SLC47A1", "MST1", "PHGDH", "CD34", "TSPAN7", "IGF1", "DIR
, "MREG", "GPX1", "MST1", "FLRT3", "FKBP11", "SLC34A2", "ETV4", "ENPP2", "IL13RA2", "CFH", "IDO1",

, "MAP1A", "MFAP4", "NR2F1", "RCAN2", "RPS6KA2", "H1F0", "COL9A3", "APOC1", "CD14", "TMEM176
5A", "COL4A6", "ROGDI", "SSX1", "NOTCH1", "PELI1") c("SNCA", "AOX1", "CXCL12", "NDUFA4L2", "RFW
L1X", "KHDRBS3", "PHACTR1", "WNT5A", "IFITM3", "OPHN1", "CYC1", "ASCC3", "TOX3", "CCNE2", "PELI1
, "SFRP4", "HOPX", "TSPAN7", "DIRAS3", "GSTO1", "PMP22", "MEF2C", "TRHDE", "LPAR1", "PTGDS", "Y
"OLFML3", "ZDHHC11", "PON3", "TFPI", "SERPINB9", "MST1", "PODXL", "SERPINE2", "DPYSL2", "SFRP4"
D1", "MELK", "SPC25", "CXADR", "USP13", "PHACTR1", "USP6NL", "SPRY2", "IFIT2", "MUC1", "CORO1C",
PR", "GPC3") c("ID2", "SMC2", "ABCB6", "BRD8", "OIP5", "GRAMD1C", "BTG1", "GAS1", "ANKMY2", "TM
ZHX2", "ZNF586", "ASMTL", "STAC", "ADH5", "SATB2", "UBB", "QDPR") c("GNPDA1", "RAB25", "TRIM22"
BCB6", "SLC37A4", "TNFSF10", "SRPK2", "SCAND1", "AZGP1", "FCGBP", "AKR7A3", "E2F2", "PRKDC", "SR
"IL11") c("C1S", "IL1R1", "MTHFD2", "SULF1", "IFIT1", "HERC5", "ADAMTS5", "HSPA13", "FADS1", "DAZ
", "SHOX2", "VAT1", "ALDH3B1", "EDNRA", "CXCL10", "LGALS1", "LTBP1", "SNAP25", "TKTL1", "TIMP3",
CDKN1B", "PSMD12", "NTRK2") c("RGS4", "NDC80") c("SLCO3A1", "FKBP5", "IRS2", "ATP2B1", "PELI1",
GD", "THAP11", "MEIS2", "GALNT2", "DES", "BCL11B", "CST3", "TRIM22", "RAB25", "ZBED1", "GAS1", "S
", "CD320", "FZD10", "ACSL1", "SPRR2B", "GALNT11", "RAMP3", "GPN3", "CST3", "COPS7A", "OIP5", "L
", "RNF130", "TM7SF2", "SMC2", "MITF", "MT1H", "GCLC", "CST3", "ALAD", "SLC1A3", "PIK3R1", "PLLP",
99", "UBL3", "BAG2", "RUNX1") c("AZGP1", "MEIS2", "RGS4", "NDRG2", "NVL", "CXADR", "ITGAE", "ALD
A", "AKAP12", "HERC5", "SIDT2", "DIRAS3", "FADS1", "FXYD5", "MYOT", "HHEX", "FUT8", "LTBP3", "LOC
LS1", "SIDT2", "UBR4", "MICAL1", "MEF2C", "NR2F2", "FN1", "TGFBR3", "ALDH3B1", "PDE4D", "TGFBR2"
NAP1L2", "ABCA5", "ARMCX6") c("PYCARD", "NVL", "SCGB2A1", "COL8A1") c("XIST", "UBE2G2", "ATP2B
H1", "GPC3", "TNNC1", "ABLIM1", "KIAA0040", "SLC24A3", "LMBRD1", "SNAP25", "APOD", "CST6", "CS
CB", "NVL", "DHCR24") c("PNN", "FAS", "POLR2A", "CD55", "EEF1E1", "PFDN5", "ADCY1", "DUSP2", "ZN
SSPN", "S100A8", "ARMCX2", "CRYAB", "CRISP3", "GNPDA1", "TRPS1", "WNT5A", "NDP", "AHNAK2", "N
", "RAB7A", "SLC1A3", "MT1H", "ITGAE") c("NFATC3", "CAV2", "SERPINA5", "TRIM22", "F2RL1", "MYCN"
RG1", "THSD7A") c("XIST", "OXA1L", "EIF3H", "TPD52", "UBR5", "AHNAK", "TRIM2", "SH3GLB2", "NEBL"
2F2", "SCGB2A1", "IKZF1") c("HSPA1A", "ERAP1", "XIST", "FKBP5", "RNF13", "IRS2") c("SLC1A3", "ERAP1"
1", "PLXNA2", "CNN3", "CD99", "UNG", "SNAP25", "LIMCH1", "CUL4B") c("MITF", "CTDSPL", "SMC2", "
B2", "RTN3", "PAQR4", "SCGB2A1", "KCND3", "DHCR7", "MCM6", "AZGP1", "OIP5", "P4HB", "SRPK1", "S
AM", "C5AR1", "PTP4A1", "IGFBP4", "OLFM1", "S100A4", "PCDH17", "PDIA6", "ASB13", "LGALS1", "ANX
EFM", "SEMA3C", "PKP1", "GAS1") c("NCAM1", "ZNF516", "SHCBP1", "RBBP8", "FBN2", "PCMT1", "LGAL
LASP1", "PCM1", "PSIP1", "TM2D3", "GLRX", "MAOA", "RAP2A", "NAV2", "LRIG1", "CXCL14", "TCF4", "S
OD1", "RASSF1", "SSX1", "BCHE", "SHOX2", "KIF18A", "MYBPC1", "UNG", "RAD51C", "PCYOX1", "SERPIN
"TDP1", "RNFT2", "PAPSS2", "EVI2A", "PDLIM5", "SACS", "CDH2", "ZNF281", "MSX1", "RET", "SLC44A4",
DXL", "SATB2", "DHRS9", "HMGCS1", "ZDHHC11", "SNHG3", "RRS1", "BET1", "HNRNPAB", "TEX10", "PP
", "GALNT2", "SORBS1", "PRIM2", "MAOB", "STEAP3", "PPID", "OGFOD1", "PCOLCE", "RASSF1", "AKT3",
", "SORBS1", "GALNT3", "MAOB", "SCD5", "STEAP3", "RASSF1", "OPHN1", "LTBP1", "SHOX2", "HERC5", "
, "RAP2A", "ZDHHC11", "SNHG3", "UBIAD1", "SPC25", "GALNT2", "SORBS1", "STEAP3", "PPID", "PCOLCE
, "SRPK1", "SPC25", "LOR", "GALNT2", "IQCG", "NR3C1", "ENO2", "PTPN3", "PCOLCE", "LTBP1", "LPAR1"
HL1", "CSNK1E", "HHEX", "GPM6B", "DKK3", "SATB2", "PCOLCE2", "PPIF", "CLIC4", "TKTL1", "ZDHHC11"
", "PCYOX1", "SERPINB9", "GBP1") c("SERPINA1", "HSPA6", "TUBB2A", "COL9A3", "CRELD2", "HSP90AA
1", "H2AFX", "SHOX2", "HERC5", "LPAR1", "NOTCH1", "HLTF", "PCYOX1") c("MGLL", "PRSS23", "HSP90A
B", "SLC7A1", "SPARC", "CGA", "SPARCL1", "GALNT3", "SEC61B", "PCOLCE", "WRB", "TDP1", "SCML2", "
, "SRPK1", "MRPS16", "UBIAD1", "LOR", "DNAJC10", "WFDC1", "COL15A1", "MAOB", "VEGFA", "OGFOD
"SCG5", "SLC47A1", "LGALS1", "CORO1C", "FN1", "QKI", "ABCA3", "GNAI2", "VAT1", "CCDC88A", "MICA
", "SH2D2A", "NCBP1", "PELO", "AKT3", "ITGAE", "SLC25A15", "PSRC1", "GREM1", "SEL1L", "WNT5A", "
, "KHDRBS3", "WNT5A", "OXA1L", "SULF1") c("SMC1A", "TCEA2", "TSPAN4", "IL6R", "FOXM1", "KCNN4",
, "DPYSL2", "CCDC92", "CD55", "EMP3", "PMP22", "SERPING1", "QKI", "PON3", "ABCA3", "GNAI2", "POD
"UBIAD1", "SPC25", "LOR", "SPRY2", "DNAJC10", "GALNT2", "ME2", "CACYBP", "SCML1", "CCNE2", "TFR

2", "SMC2", "TMEM14A", "GPN3", "TAF5", "MEOX2", "MEIS2", "RTN3", "IFIT1", "RNF130", "ITGAE", "CD3
ACF1", "CLNS1A", "GNAI2", "AIDA", "GYS1", "SERPINH1", "TFPI", "UGDH", "SIRT3", "TGFBR2", "EXT1", "L
", "TMBIM4", "BCAS1", "CHN2", "SATB2", "KIAA0232", "ADAP1", "ACE2") c("TXNIP", "GRAMD1C", "LIMK
150", "BTG1", "SLC25A11", "COL8A1", "TNFSF10", "REEP1", "AZGP1", "ENPEP", "PYCARD", "ZNF512B", "
SPT2") c("GSK3B", "NRP1", "HMGCS1", "SCUBE2", "ZNF83", "FASN", "ARFIP2", "SPINK1", "HMGB1", "G
DNRA", "STUB1", "SNAP25", "PELI1", "PCCA", "SIRT3", "KCNMA1", "GSTO1", "PTPRD", "OPHN1") c("SF3
C2", "FTO", "SNAP25", "PELI1", "PCCA", "HSP90AA1", "SIRT3", "KCNMA1", "GSTO1", "HSD11B1", "FGF1
"MGLL", "IFIT1", "QKI", "IFITM3", "IFITM2", "RUNDC3B", "IVD", "A2M", "TIMP3", "CLNS1A", "TCTN2", "
"RUNDC3B", "IVD", "BICC1", "MYH10", "SYNM", "VPS13C", "ARL6IP5", "ZDHHC11", "RARRES2", "DYNLT
"TMEM14A", "WNT5A", "PDE6D", "RTN3", "THSD7A", "IFIT1", "IFIT3", "ACSL1", "OAS1", "MT1H", "FAM5
561", "COX7B", "NDUFA9", "CHMP2A", "NDUFB7", "SPRR2B", "SPRR1B", "GALNT2", "NDUFA6", "SLC25
"GM2A", "IFIT1", "RET", "CXCL10", "LTBP1", "PCBP4", "IFIT2", "EIF2AK2", "SOD3", "PELI1", "MEG3", "PY
2C", "CDCA3", "NFIB", "MT1M", "ITGAE", "OIP5", "SMC2", "MT1H", "CD320", "COPS7A", "ACSL1", "MCM
", "HOXA10", "HOXB6", "MSX1", "CXCL14", "ZNF91", "LPIN1", "SESN1", "AHR", "LGR4") c("SCGB2A1", "
", "TSC22D1", "ADH5", "IL15RA", "PEX6", "TP53I3", "CD99", "CST6", "RUNX1", "KIAA0232", "ZNF586", "C
"PCTP", "TNFSF10", "DKFZP586I1420", "C3", "NVL", "APEX1", "CKAP2", "COMMD3", "PLA2G15", "IGFB
M2", "MYBPC1", "RAD51C", "ASF1A", "TFIP11", "SCUBE2", "ZNF589", "CCR2", "MLH3", "GPR143") c("RG
SS23", "CORO1C", "FAM174B", "OPN3", "LPAR1", "CXCL10", "NR3C1", "ACVR1B", "EIF3G", "MFAP3L", "
"NR3C1", "PPP1R10", "ZBED5", "DPF2", "SCUBE2", "ZNF277", "ELN", "POGZ", "PHF10", "ZFP64") c("CYP1
"SNAI2", "PTGDS", "FABP7", "LPAR1", "TSPAN7", "ZNF337", "PDGFA", "SCUBE2", "SH3BGRL", "PMP22",
CTPP1", "MAPK13", "CORO1C", "NR2F2", "LRIG1", "DPP8", "BAZ1A", "ARMCX3", "UBR4", "SAR1B", "LP
NT5A", "SMC2", "COPS7A", "FZD10", "MCM2", "GNPDA1", "TXNIP", "PIK3R1", "BTG1", "MT1H", "MYLIP
", "BBS1", "ENPEP", "BANP", "AZGP1", "CASP1", "MCM6", "GLCE", "TSPAN7") c("SLCO3A1", "CNOT4", "
", "SAP30", "TRIAP1", "NPAT") c("ARMCX6", "CDKN1B", "PSMD12", "UBR5", "COIL", "DDX42", "GLRX", "C
R", "ZNF589", "BTG3", "MT1M", "ANKMY2", "DES", "ACSL1", "TM7SF2", "GAS1", "DST", "ALAD", "TCFL5
TNS3", "ACSL1", "UBE2E3", "REPIN1", "RAMP3", "C3", "CLNS1A") c("XIST", "PLS3", "WDR61", "ZNF721",
, "CPB1", "OPTN", "EML3", "PRG2", "MOAP1", "FKBP1B", "TTC3", "DEGS1", "SLC7A8", "CXCL14") c("NVL
S7A", "IFIT1", "OAS1", "REEP1", "NFIB", "PLLP", "TCTA", "DST", "ACSL1", "NVL", "CXADR", "RNF130", "G
ACSL1", "TM7SF2", "AZGP1", "WNT5A", "GAS1", "IFIT1", "ALAD", "RASL11B", "IFIT3", "TCTA", "TXNIP", "
"FCGBP") c("XIST", "TUBB2A", "PLXNA1", "UBE2G2", "ADCY1", "RUNX1", "ZNF274", "KLHDC2", "IRS2", "

```
/51" "V52" "V53" "V54" "V55" "V56" "V57" "V58" "V59" "V60" "V61" "V62" "V63" "V64" "V65" "V66" "V
CD2", "PLXNB1", "XIST", "ELF5", "MKNK2", "ESR1", "AHNAK2") c("E2F2", "ZNF512B", "RAB25", "C4BPA",
3XO21", "FOXJ3", "GNPDA1", "ID2", "IFIT1", "KCNJ2", "MITF", "MX2", "NCOA2", "NDRG2", "OAS1", "RAB
M22", "TSPAN7") c("AMACR", "DCK", "DUSP2", "ERAP1", "ETNK1", "HBP1", "HSPA1A", "IRS2", "KCNK3", "
", "SLC17A5") c("ACO1", "ARMCX2", "CLNS1A", "COL4A6", "EDNRA", "IFIT1", "IKBKE", "LRIG1", "NOTCH1
", "FZD10", "CDCA3", "SMC2", "MCM2", "TXNIP", "TBX2", "THAP11", "BTG3", "PIK3R1") c("ITGA3", "PPIF
3", "MYLIP", "KIF2C", "PIK3R1", "CHKA", "ABCG1", "ANKMY2", "OAS1", "DDX58") c("NFATC3", "CAV2", "N
NRBF2", "COPS5") c("PCDH9", "MED17", "SIX2") c("PCDH9", "RBP4", "PNMA2", "STC2", "UNC119B") c("(
BE2E3") c("TUBB2A", "PELI1", "ASAP1", "RNF138", "MCOLN3") ACOT7 c("ALDH1A2", "SMC1A", "PCDH9"
1", "TMPRSS3", "DHCR7") c("SSBP2", "XIST", "PLS3", "ERGIC2", "PELI1", "FKBP5") c("LAP3", "ANGPT1", "F
", "MEOX2", "PLLP", "SMC2", "RAB25", "POMZP3", "DES", "BCOR", "BCL11B", "CHKA", "CBX4", "ROGDI",
RIM24", "GOLM1", "RGN", "DDX42", "GPC3", "PCBD1", "SALL2", "BAG2") c("BTG3", "DST", "AZGP1", "RA
2", "PADI2", "HSDL2", "LDB3", "DACH1", "KRT1", "COL4A5", "ROGDI", "NDN", "TCFL5", "EFHC2", "TRPS1
N4", "OLFML3", "MFAP3L", "KLF10", "CTSC", "MYO1B", "IGFBP4", "HSPA13", "THY1", "LGALS1", "TSPAN7
FBXO21", "RNF10", "GAS1", "HGF", "PKP1", "CA9", "PIGT", "ROGDI", "PSMA3", "MCF2L") c("NCAM1", "A
PGD", "HEATR1", "LGALS1", "IL20RA", "SPON1", "ISLR", "CCDC85B", "TCF4", "CXCL13", "CSTA", "TRHDE"
8A", "GGCT", "PRIM2", "MACF1", "USP13", "UNG", "MRC2", "ZNF83", "ADK", "CCR2", "ASCC3", "ARFIP2"
M1", "DNAJC10", "DIRAS3", "GSTO1", "TRIP13", "CLIC4", "DHCR7", "PCOLCE", "GRWD1", "LPAR1", "RPA3
", "MST1", "FLRT3", "FKBP11", "HSPA13", "TMPRSS3", "SLC34A2", "GSTO1", "TM4SF1", "AKAP11", "CSF2
, "CCDC59", "SELL", "ALG13", "MUC1", "STAT1", "LYZ", "NPC1", "HAPLN1", "PLEK", "ZYX", "MGAM", "CD
", "MAN2B1", "PPID", "TERF1", "SLC10A3", "PAK2", "ANKRD10", "KPNA3", "NOTCH1") c("TCN1", "RRM2
OLCE", "GRWD1", "ME1", "MED27", "KPNA4", "MTMR2", "SIRT3", "SPC25", "PRIM2", "MACF1", "USP13
MACF1", "USP13", "IFITM1", "IPO5", "IFIT5", "NEFM", "IFIT2", "SEMA3B", "ZNF83", "CORO1C", "IFITM3",
E", "GRWD1", "HSP90AA1", "SMC2", "TACC3", "UBE2C", "NEK2", "TPX2", "SMC4", "USP13", "PHACTR1",
MACF1", "USP13", "PPDPF", "SATB2", "UNG", "MRC2", "IFIT2", "ZNF83", "CORO1C", "IFITM3", "OPHN1",
MPRSS3", "FXYD5", "DIRAS3", "CLIC4", "CDH11", "ENPP2", "OPN3", "GRWD1", "COTL1", "KHDRBS3", "C
ON", "CD93", "DRAM1", "RGS5", "RCAN2", "CD14", "GLIPR1") c("PCNA", "ZMAT3", "RPS27L", "FAS", "PRS
ABP7", "ZNF83", "IFITM3", "ABCA3", "GPI", "DSTN", "LHFPL2", "CTNNB1", "U2AF1", "CYC1", "ASCC3", "G
C", "LPAR1", "TEK", "SDC2", "MTMR2", "CSF2RA", "SLCO2A1", "ZNF473", "TCF21", "TPSB2", "TPSAB1", "
'CLIC4", "CKS1B", "PMP22", "PCOLCE", "LPAR1", "CILP", "COX7A1", "ME1", "PTGDS", "MAN1A1", "ACTA
5", "RET", "RPS27L", "PRSS23", "MGLL", "HERC5", "ZNF337", "SPEN", "IFIT1", "ZDHHC11", "SERPINB9", "
3BP", "IPO5", "MRC2", "IFIT2", "ZNF83", "CORO1C", "IFITM3", "ABCA3", "PUF60", "GDPD5", "ASCC3", "D
MACF1", "USP13", "IFITM1", "CXCL10", "IFIT2", "ZNF83", "RBM26", "CORO1C", "IFITM3", "IFITM2", "EIF4
DIRAS3", "GSTO1", "CDC42EP3", "RASIP1", "ADD3", "FOLR1", "CKS1B", "KLHL21", "COL6A2", "ABCA8", "(
GER3", "MFAP3L", "ANXA6", "COL4A1", "MST1", "LGALS1", "GSTO1", "CLIC4", "OPN3", "RRP8", "PCOLCE"
, "GNG11", "IFI44L", "PCOLCE2", "FSTL1", "COCH", "EFHD1", "HSD11B1", "COL5A2", "ZDHHC11", "SERPI
C25", "NBPF10", "MRC2", "IFIT2", "ZNF83", "IFITM3", "GDPD5", "DSTN", "CTNNB1", "ASCC3", "HUWE1",
M4SF1", "LPAR1", "H2AFX", "HLTF", "PARP8", "PPDPF", "POLR1D", "SOX4", "CD81", "HGF", "XAB2", "CA9
PHACTR1", "IFITM1", "CXCL10", "MRC2", "CCL8", "CORO1C", "IFITM3", "ABCA3", "OPHN1", "GDPD5", "IF
, "THY1", "TSPAN7", "DIRAS3", "GSTO1", "PRUNE2", "PCOLCE", "MEF2C", "PAEP", "LPAR1", "YRDC", "TE
P", "COX7A1", "ADAMTS1", "HSP90AA1", "ID3", "DEPDC1", "AURKB", "SKA1", "EXOSC8", "KCNQ2", "CALI
"HSD11B1", "OLFML3", "PON3", "SERPINB9", "NUCB2", "SLC47A1", "MST1", "SFRP4", "LGALS1", "DNAJC
D83", "CKS1B", "PCOLCE", "GRWD1", "LPAR1", "IQCG", "MED27", "HSP90AA1", "TACC3", "FERMT2", "TN
", "TGM2", "PCOLCE", "GRWD1", "LPAR1", "ME1", "MED27", "HSP90AA1", "NELL2", "MACF1", "WFDC1"
'CALR", "IFIT5", "SATB2", "CBS", "IFIT2", "HEPH", "CCL8", "IFITM3", "ABCA3", "IFITM2", "ANXA5", "HSPA
RAS3", "CKS1B", "PCOLCE", "GRWD1", "LPAR1", "COX7A1", "PTGDS", "ADAMTS1", "NCBP1", "SIRT3", "CK
, "TBL1X", "C1QB", "RIPK2", "CXCL10", "SATB2", "HEPH", "ZNF281", "SLC14A1", "ZFPM2", "CSF1R", "IFITI
```

3", "GLIPR1", "MRPL15") c("SNAP25", "RET", "RPS27L", "PRSS23", "MGLL", "TGFBR3", "ZNF337", "LOR", "
D3", "AGPAT5", "NDRG4", "PAEP", "ITGAE", "NCBP1", "CIT", "APOA2", "TNC", "TOX", "TCTA", "PSRC1", "I
L") c("IL6R", "SMC1A", "KRT2", "GGH", "TSPAN4", "DSG3", "MPHOSPH9", "SFRP4", "CKAP4", "KIF15", "FC
RDC", "NBL1", "SDC2", "NEK2", "NCAPH", "TIMP3", "TUBB2B", "SLCO2A1", "PRKCA", "MRC2", "SAP18", "
", "HSPA13", "CD34", "DIRAS3", "GSTO1", "CLIC4", "RASIP1", "PMP22", "COL6A2", "TEX10", "LPAR1", "CI
, "IFITM3", "IFITM2", "PDE8B", "CYC1", "ASCC3", "TFRC", "NPC1", "GYS1", "HUWE1", "CD81", "S100P", "
EM14A", "BTG3", "KIF2C", "MEIS2", "MCM2", "KCNJ2", "POMZP3", "NCOA2", "COPS7A", "MT1M", "REE
", "CXADR", "COPS7A", "FOXJ3", "CST3", "ALDH3A2", "BTG1", "EED", "ABCB6", "TXNIP", "MX2", "IFIT1", "
EBF1", "CYB561", "RGS4", "ANAPC13", "HMGB2", "C3", "TMEM14A", "DHCR7", "BCL6", "ATP10B", "PYCA
AP2", "AKAP12", "GNS", "EDNRA", "CXCL10", "ALOX5", "SNAP25", "EPS8", "MATN2", "SATB2", "FKBP11",
"NELL2", "ZHX2", "TBC1D16", "TGFBR2", "A2M", "RRAGD", "NR2F1") c("FRY", "TOPBP1", "JAG1", "ANO1
"RNF138") c("IRAK3", "SAP30", "ALDH2", "ARNT2") c("PCDH9", "TARP", "ADD3", "IRS2", "PSMD12") c("L
MPDL3A", "CD320", "BTG3", "RAD51C", "ACSL1", "TAF5", "ANKMY2", "WNT5A") c("SAP30", "CAV2", "LII
EPROTL1", "CDCA3", "ABCB6", "VSNL1", "CYB561", "ITGAE") c("RAD23A", "KBTBD11", "TIMM8B", "HAMI
"TCTA") c("HOXB7", "SERPINA5", "NFATC3", "ZNF468", "LIF", "LGALS3", "PCSK6", "FTH1") c("HPGD", "S1
H3A2", "MT1M", "SLC27A3", "PPIH", "TCTA", "GAS1", "CST3", "TXNIP", "ABCG1", "MITF") c("F2RL1", "TC
283683", "APOE", "IGFBP1", "ANTXR1", "CDH11", "LAPTM4B", "ACTA2", "CCL18", "IFITM2", "SULF1") c("
", "SLC47A1", "IGFBP3", "MPDZ", "MACF1", "GSTO1", "IFITM2", "ODC1") c("CXCL12", "MAFB", "TOX", "IT
1", "PRNP", "RPS26", "IL1A", "TM4SF20", "ZNF721", "EGOT", "PLXNA1") PHACTR4 c("SMC1A", "GAS6", "I
P1") c("ITGAE", "BTG3", "MEIS2", "MT1M", "THAP11", "AZGP1", "UNC119B", "ZC3HAV1", "FOXJ3", "REI
F329", "PELI1", "ASXL2", "XIST", "HEBP2", "PERP", "ABHD6", "ATP11B") c("ALDH2", "ARHGEF3", "FASTKD
TRK2", "NEBL") c("SRPK2", "ALDH3A1", "SPATA20", "ITGAE", "GNPDA1", "RRAGD", "ACO1", "MCM6", "C
", "RAB5B") c("AZGP1", "RIMS3", "SH3BGRL", "CRYAB", "TPD52", "MAGEH1", "ACPP", "FBXL5", "GATM",
, "CHPT1", "DDX42", "CD36", "CRYAB", "PPL", "STUB1", "CSTA", "ITIH5", "PPP1R3C", "IGF1", "ZBED2", "L
", "SAP30", "PSMB8") c("MSX1", "TARP", "WBP4", "MAN1A1", "IRS2", "POLR1E") c("STC2", "POLR1E", "R
MCM2", "RTN3", "PAQR4", "WNK1", "AZGP1", "OIP5", "VSNL1", "CD320", "ALDH3A2", "FZD10", "SLC1A3"
PRR1B", "PSAT1", "EGF", "TBCB", "GALNT2", "KLHDC8A", "AGTPBP1", "DHCR24", "C3", "NDUFB6", "IKZF
A5", "MMD", "SERPINA3", "C1QB", "MLXIP", "OSBPL10", "TMEM158", "TUBB3", "PLAUR", "NOVA1", "PL
S1", "SGCB", "NFIC", "AZGP1", "HHEX", "GABBR1", "MDC1", "SLC7A8", "IGF1", "CD320", "ALDH3A2", "IG
ERPINI1", "CXCL13", "RHOB", "MAST2", "PTGDS", "COL15A1", "IL20RA", "SCD5", "SUSD4", "STAC", "ANG
B9") c("PELI1", "FARP1", "FADS1", "CCL18", "LAPTM4B", "C1QB", "GNAI2", "HHEX", "RAP2A", "GPM6B",
"ITIH5", "RGS5", "CREB1", "SULF1") c("PELI1", "LDB2", "FARP1", "DPY19L1", "FBN2", "RCBTB2", "LGALS
ID", "AHSA1", "SHOX2", "GPR37", "TIAM1", "CYP51A1", "SIRT3", "SLC25A37", "NOTCH1", "MYC", "GRAN
"SSX1", "OPHN1", "LTBP1", "SHOX2", "LMCD1", "PLTP", "ABCC3", "SIRT3", "NOTCH1", "SNAP25", "IFIT1"
PGR", "ABCC3", "NOTCH1", "UNG", "SNAP25", "IFIT1", "SEMA6A", "PCYOX1", "DCLRE1C", "ADI1", "SERP
", "RASSF1", "SSX1", "OPHN1", "LTBP1", "HERC5", "PGR", "NELL2", "LPAR1", "SIRT3", "SNAP25", "IFIT1",
, "ABCC3", "FSTL1", "IFIT1", "CAPRIN1", "CXCL10") c("PELI1", "PRSS23", "HSP90AA1", "PARP2", "OLFM1"
", "IFI44L", "PDLIM3", "GALNT2", "SORBS1", "RCAN2", "A2M", "KLF11", "PER2", "FBLN5", "TEX10", "PPID
1", "HSP90B1", "FARP1", "DPY19L1", "FBN2", "CILP", "MT1F", "JMJD6", "DPYSL2", "LGALS1", "CORO1C",
A1", "DNAJB4", "ITM2A", "FADS1", "CILP", "PIP4K2B", "CORO1C", "EMP3", "PMP22", "JAM3", "PON3", "
LTBP1", "PGR", "RCN1", "LPAR1", "PNMA1", "NEFM", "TCTN1", "MSX1", "MYH11", "SLCO2A1", "IGFBP3"
1", "LRRC37A3", "PCOLCE", "RASSF1", "LTBP1", "NELL2", "TGM2", "LPAR1", "ABCC3", "NOTCH1", "NRXN
AL1", "SERPINH1", "LRIG1", "TKTL1", "ZDHHC11", "SNHG3", "ODC1", "UBIAD1", "SPC25", "LOR", "DNAJC1
PHKA1") c("TCEA2", "TFF1", "RRM2", "PRSS23", "SCG5", "SERPINB2", "PBK", "SOSTDC1", "SLC47A1", "CTI
, "PLA1A", "SFRP4", "KIF15", "PCSK6", "GGH", "EXOSC4", "SGCE", "CKAP4", "MSMB") c("MGLL", "TFF1", "
OXL", "MICAL1", "SERPINE2", "DOCK10", "SFRP4", "UCHL1", "CLIC4", "ZDHHC11", "SNHG3", "RASIP1", "SI
RC", "SLC37A4", "SHOX2", "KIF18A", "ABCC3", "NOTCH1", "ADI1", "NGDN") c("MICB", "S100P", "MGAM",

320", "RGS4", "GAS1", "DES", "NPDC1", "RAB25", "ACSBG1", "LRP5", "CCDC28A", "CYB561", "CXADR", "T
OR", "DCTPP1") c("CIT", "PSRC1", "APOA2", "TIMM17A", "ITGAE", "JAG1", "NDUFA4L2", "RAI14", "TWF2
2", "LSM4", "BTG1", "PIK3R1", "COL8A1", "REEP1", "AZGP1", "GALNT11", "GAS1", "IFIT1", "CXADR", "NV
SLC16A6", "SPATA20", "FAM117A", "ACADVL", "ARID1A", "NVL", "C4BPA", "GMNN", "ACP6", "COMMD3
GCT", "MRC2", "RNF128", "MACF1", "BCHE", "CCR2", "PCCA", "CGA", "AHCY") c("C7", "LAPTM4B", "KHDF
A2", "GZMA", "GNS", "SNX11", "TBC1D12", "IFIT1", "IFITM3", "IFITM2", "SORBS2", "MYH10", "AHCYL2",
3", "MAP2K5", "NR1I2", "PROX1") c("PPP2R5E", "KHDRBS3", "PPP3CA", "TNFSF10", "RGL2", "MGLL", "TN
MRPL44", "NID1", "ZDHHC11", "DYNLT3", "MXRA5", "IKBKE", "CXCL10", "EDNRA", "IFITM1", "FTO", "MA
T3", "MRC2", "PDLIM3", "RGS5", "DDB2", "F2R", "TACC1", "AOC3", "PCP4", "WSB1", "EFHD1", "PRKCA", "
50A", "NVL", "CHKA", "IFIT2", "REEP1", "NTHL1", "ITGAE", "ABHD10", "MT1M") c("SERPINA5", "PON1", "
A46", "TMEM14A", "PDE6D", "C3", "RTN3", "APBA2", "ACSL1", "NVL", "DHCR7", "CHKA", "REEP1", "NTH
CR1", "GSTO1", "SLC47A1") c("TOX", "NDUFA4L2", "WNT5A", "SLC35A1", "MRPL2", "GREM1", "PPP1R10
12", "BTG3", "PIK3R1", "THAP11", "IFIT2") c("ZNF468", "NFATC3", "NPC1", "SLC7A5") c("MGLL", "MED21
D2", "PYCARD", "SIX1", "CHKA", "TNS3", "RGS4", "TCTA") c("NFKBIA", "PNN", "DUSP2", "SIGLEC15", "ZF
X3CL1", "BMI1", "GOLM1", "E2F5", "SMARCA2", "SMARCD3", "SNAP25", "STUB1", "LMBRD1", "PAK1") c
P2", "TCTA", "TSPAN7", "TMEM14A", "PEX6", "AASDHPPT", "LEPROTL1", "GDE1", "REEP1", "SIX1", "NCO
S5", "HHEX", "OXA1L", "UMOD", "ACTN1", "AKAP12", "FGF13", "EPS8", "RET", "SLC35A1", "IRF4", "ADA
SYCP2", "GDPD5", "PPOX", "PHF10", "MAF") c("JMJD6", "HSP90AA1", "HHEX", "LGALS3BP", "RBMS1", "
LB1", "UCHL1", "TFPI", "HSP90AA1", "SPEN", "DNAJB4", "XIST", "MGLL", "AHCY", "LGALS3BP", "ACSL4", "
"SNAP25", "PHF10", "NELL2", "MLH3") c("TCN1", "DNAJA1", "HSP90AA1", "SPEN", "OXA1L", "ADNP", "L
AR1", "TFIP11", "NCALD", "CXCL10", "C7", "ZNF337", "UBE4A", "UGDH", "MEF2C", "GDPD5", "FN1", "PP
", "BANP", "AZGP1", "IFIT1") c("F2RL1", "NFATC3", "DIO1", "PPIF", "EFNA1", "DYNC2LI1", "BANP", "NPA
WDR61", "PELI1", "RUFY2", "TOPBP1", "GADD45B", "HSPA1A", "DUSP2", "FKBP5", "HBP1", "INTS7", "PL
OXA1L", "ESR1", "CXCL14", "EIF3H", "ZBED5", "S100A8", "RXRA", "CALML5", "PTPN3", "ZNF91", "CALML3
", "REPIN1", "CST3", "ST13", "RAMP3", "MEIS2", "VWA1") c("F2RL1", "FTH1", "LGALS3", "CREB3", "COM
"SNRPE", "RPS26", "KCNK3", "SPAG16", "IRS2", "DUSP2", "LYST", "TMEM131", "INHBA", "ANKRD12", "L
", "BTG1", "TXNL4B", "DHCR7", "TRIM22", "ATP10B", "KCND3", "GPX7", "FAM117A", "ACSL1") c("KCNK3
AS1", "MRPL48", "BTG3") c("SERPINA5", "CSAD", "ERBB3", "LGALS3", "MYCN", "PCSK6", "NFATC3", "F2R
CST3", "SHROOM2", "CDIPT", "ACAD8", "TCFL5", "SPRR2B", "ZBED1", "RGS4", "MT1H", "MT1M", "SEC11
PERP", "CD55", "CGA", "TSG101", "ZNF277", "PELI1", "HBP1", "NINJ2") c("TRIM22", "LAP3", "ERBB3", "T

67" "V68" "V69" "V70" "V71" "V72" "V73" "V74" "V75" "V76" "V77" "V78" "V79" "V80" "V81" "V82" "V8
, "DCTPP1", "DHCR24", "GPX7", "P4HB", "FDPS", "SSH3", "SAC3D1") c("ZNF721", "RUFY2", "SLCO3A1", "
25", "RBBP4", "SALL2", "SMC2", "SMPDL3A", "TM7SF2", "TRIM22", "TXNIP", "ZC3HAV1") c("COMP", "EF
LRRN3", "NFKBIA", "NINJ2", "RUFY2", "SSBP2", "TOPBP1", "XIST", "YWHAB") c("ALDH2", "BBX", "CLDN7
1", "OGN", "PLTP", "PMM1", "PTEN", "SLC17A5", "SNTB2", "SPARCL1", "SPON1", "STC1", "TCEAL2", "TSP
", "SLC7A5", "KDSR", "KBTBD11", "TCL1A", "NFATC3", "HBD") c("CPB1", "MT1G", "GOLM1", "COL4A5", "
MYCN", "KBTBD11", "NPC1") c("SLC24A3", "NEBL", "PRR15L", "MKNK2", "TRPS1", "ESR1", "ENO2", "AHN
CSGALNACT2", "MCOLN1") c("LPAR1", "CEBPG", "ELF5", "RPS6KA2", "KCNK2", "ARMCX1", "PCOLCE2", "
) c("LRRFIP1", "PCDH9", "RBP4", "EIF5") c("MCOLN1", "CYCS", "EIF5") c("FKBP1B", "CEBPG", "CD209", "S
PHACTR4", "TMPRSS3") c("CLU", "PCDH9", "MRPS30", "GNAI1", "IL7R") c("PI3", "EIF5", "USP7", "LRRFIP1
, "LRP5", "MT1H", "WNT5A", "NVL", "TMPRSS3") c("HOXB7", "SLC7A5", "HAMP", "CAV2", "RB1CC1", "KD
AD51C", "POMZP3", "MCM2", "IFIT1", "IFIT3", "FZD10", "TMPRSS3", "CD320", "RGS4", "MT1M", "ITGAE"
", "XIST", "HOXB6") c("AZGP1", "C3", "SCGB2A1", "PSAT1", "ALDH3A1", "MCM2", "IKZF1", "SPRR1B", "T
", "IGF1", "NLRP2", "PDIA4", "ISLR", "TMEM158", "CLIC4", "RASIP1", "MCOLN1", "MARS", "TLR4", "DCN
AZGP1", "AHNAK2", "KRT2", "SHCBP1", "RBBP8", "AKR7A2", "FANCL", "IGF2BP3", "CALML3", "CTSG", "C
, "LMO4", "LIMCH1", "PTGDS", "AUTS2", "HLTF", "ITM2C", "PARP8", "RNASET2", "PRKACB", "PHACTR1",
", "HUWE1", "MYBPC1", "FLG", "STEAP3", "SPINK1", "TFCP2L1", "SSX1") c("SNAP25", "RET", "ALOX5", "A
", "C7", "ACTA2", "SMC2", "OIP5", "MRTO4", "TUBB", "SKA1", "PPP2R1B", "PRIM1", "PPDPF", "IVNS1AB
2RA", "PAPSS2", "POLR1D", "CBS", "ZNF281", "NR2F6", "LSP1", "MATN2", "HUWE1", "RGS5", "SYNE1", "
2", "ATP2A3", "PAK2", "SSX1", "CCT4", "KYNU", "SFPQ") c("SLC39A6", "RET", "NOVA1", "RPS27L", "FHO
", "PRSS23", "LOR", "IFIT1", "IFI6", "PCOLCE2", "HSD11B1", "OLFML3", "PON3", "TFPI", "SERPINB9", "SC
", "CXCL10", "SATB2", "MRC2", "MPHOSPH8", "IFIT2", "HEPH", "ZNF83", "CORO1C", "IFITM3", "OPHN1"
"ABCA3", "GDPD5", "IFITM2", "PDE4D", "TRAK2", "ANXA5", "PLTP", "MATN2", "HUWE1", "IVD", "CD81"
"IFITM1", "CALR", "CXCL10", "IFIT2", "CORO1C", "IFITM3", "GDPD5", "IFITM2", "ASCC3", "PXDN", "ACO
"CSF1R", "SCD5", "CYC1", "ASCC3", "TRAK2", "GYS1", "HUWE1", "SOBP", "IVD", "CLIP2", "STEAP3", "RB
1QB", "PHACTR1", "IFITM1", "CXCL10", "IFIT5", "CBS", "HEPH", "IFITM3", "GDPD5", "LHFPL2", "IFITM2",
SS23", "MGLL", "LOR", "SPEN", "IFIT1", "GAMT", "SERPINB9", "DNAJC10", "GSTO1", "PCOLCE", "GRWD1"
YS1", "HUWE1", "PPID", "IVD", "SYCP2", "NOTCH1", "PELI1") c("SLC39A6", "RET", "RPS27L", "PRSS23", "
PDCD4", "ZFPM2", "LTBP3", "ZNF331", "LSP1", "SMAD4", "TRAK2", "GLDC", "HUWE1", "SOBP", "TAGLN"
2", "FERMT2", "NELL2", "TUBB2B", "SLCO2A1", "KCNQ2", "RIPK2", "PRKCA", "IFITM1", "IPO5", "NEFM", "
MST1", "DIRAS3", "GSTO1", "PCOLCE", "GRWD1", "LPAR1", "MTMR2", "SIRT3", "SPC25", "NELL2", "USP1
DMXL2", "IVD", "PROX1", "MAOB", "LONP1", "PDSS2", "SSX1", "ELOVL5", "NOTCH1", "PELI1") c("TCN1",
4EBP1", "CYC1", "ASCC3", "GYS1", "HUWE1", "SOBP", "IVD", "SLIT2", "MAOB", "COL4A6", "ROGDI", "ELC
CILP", "LMO4", "DLEU1", "DIAPH2", "GNPTAB", "KHDRBS3", "ITM2C", "CDC42BPB", "LGALS3BP", "SATB2
", "LPAR1", "ME1", "IQCG", "SPC25", "NBPF10", "IFITM1", "CALR", "CXCL10", "IFIT2", "CORO1C", "IFITM
NB9", "GPX1", "COL4A1", "FGF13", "IGFBP4", "HSPA13", "GPM6B", "FBLN5", "GSTO1", "CLIC4", "CDH11
, "MYBPC1", "IVD", "STEAP3", "SSX1", "PELI1") c("TUBB2A", "SLC39A6", "CDH2", "MKI67", "DIP2C", "SER
", "NOTCH1") c("ZMAT3", "RET", "PRSS23", "MGLL", "ZNF337", "SPEN", "IFIT1", "ITM2A", "HSD11B1", "
FITM2", "ASCC3", "PXDN", "PLTP", "MATN2", "GYS1", "HUWE1", "PPID", "IVD", "EDNRA", "ROGDI", "SYC
K", "SDC2", "NQO2", "TUBB2B", "SLCO2A1", "IFITM1", "NEFM", "SEMA3B", "COL6A1", "PDCD4", "IFITM
R", "IPO5", "DKK3", "CBS", "IFIH1", "CCL8", "HNRNPM", "CORO1C", "IFITM3", "ABCA3", "CTSK", "GDPD5"
10", "CD34", "TSPAN7", "DIRAS3", "GSTO1", "RASIP1", "PMP22", "PCOLCE", "LPAR1", "PTGDS", "YRDC",
MPO", "SKA1", "TMED2", "USP13", "PHACTR1", "LGALS3BP", "IFITM1", "CALR", "CXCL10", "IPO5", "IFIT2"
, "USP13", "PPDPF", "NEFM", "IFIT2", "ZNF83", "CORO1C", "IFITM3", "GDPD5", "DSTN", "IFITM2", "TGF
5", "ENG", "EDNRA", "GLI2", "CLIP2", "SORBS3", "LAPTM4B", "COL4A6", "ROGDI", "PDSS2", "SYCP2") c("
B", "TBL1X", "NELL2", "EXOSC8", "PARP8", "PPDPF", "PHACTR1", "LGALS3BP", "ELOVL4", "POSTN", "WN
M2", "CST3", "LSP1", "GYS1", "RGS5", "ACD", "ENG", "GLI2", "LAPTM4B", "NLGN4X", "CTSB") c("TCN1", "

"SPEN", "IFIT1", "SFRP1", "HSD11B1", "ZDHHC11", "TFPI", "SERPINB9", "SLC47A1", "CITED2", "MST1", "L
RSL1D1", "WNT5A", "ABCA3", "PHKA1", "DDHD2", "MCAM", "HGF", "COL14A1", "ENAH", "PPT1", "KPNA
)XM1", "MACF1", "PARP8", "PDE8B", "MSMB", "SLIT2", "PRRX1", "STAP1", "GATAD2A", "THBS2", "MCF2
PAK1", "PCP4", "SERPINF1", "OPHN1", "LTBP3", "TCEAL2", "TGFBR2", "MPDZ", "TRAK2", "PLP2", "PLTP",
LP", "COX7A1", "PTGDS", "C7", "YRDC", "NBL1", "ACTA2", "SIRT3", "PRIM2", "KCNQ2", "PRKCA", "MRC2
CCNE2", "NOTCH1") c("MICB", "CDV3", "RFC2", "CD200", "RPL36", "PCDH9", "GAB2", "SLPI", "RAB31", "
P1", "NDRG2", "CDCA3", "SLC1A3", "PSMC3IP") c("CLDN4", "CAV2", "HOXB7", "SLC7A5", "GRAMD1C", "
OAS1", "AZGP1", "TM7SF2", "RBBP4", "CYB561", "EIF4B", "RGS4", "BTG3", "SLC1A3", "TMEM14A", "WN
ARD", "SIX1", "NVL", "NDUFA6", "ASMTL", "ITGAE", "RRAGD", "ADI1", "SPATA20") c("NFKBIA", "GADD45
"EYA1", "CDH11", "COPS4", "RGS5", "C1QB", "OXA1L") c("NQO1", "CLNS1A", "UCHL1", "SFRP4", "MTHF
0", "CXCL12", "SLC5A6", "TBX3", "ABCA3", "TMEM147", "AOX1", "TRIM29", "CBX6", "HNRNPH3", "WNT
RRFIP1", "PCDH9", "EIF5") c("EIF5", "MAGEA6", "CYCS", "CPVL") c("ERP44", "FOSB", "FKBP1B", "PSMD12
F", "F2RL1", "TRIM22", "RAB25", "ZBED1", "COMP", "LGALS3", "IGF2BP2") c("CAST", "CITED2", "PPP1R3
P", "BZW1") c("AHI1", "TSPAN13", "FOXO4", "BCHE", "SOCS2", "CDKN1B", "PRG2", "ESR1", "XIST", "ME1
100A8", "CRISP3", "PLCB4", "CITED2", "DHRS2", "BCHE", "WNT5A", "PPP1R3C", "MEIS2", "MT1G", "MKN
L1A", "FTH1", "IGFBP1", "DIO1", "PCSK6", "SERPINA5", "HAMP") c("S100A8", "AZGP1", "XIST", "DAAM2"
"MAOB", "COL4A6", "NQO1", "HSD11B1", "USP13", "LPAR1", "GM2A", "SGCE", "SFRP4", "SERPINE2", "L
GAE", "PAEP", "TCTA", "COL14A1", "STAT5A", "RDH11", "DNMBP", "VGLL4", "CTPS2") c("CXCL10", "TCN
RBPJ", "FTO", "AFF1", "WSB1") c("USP7", "CTBP2", "CD83", "STC2", "COL9A3", "RPS26", "FTO", "AFF1", "
RE", "CYB561", "MITF", "BTG1", "TXNIP", "IFIT1", "TRIM22", "CXADR", "OAS1", "IFIT3", "ID2", "COPS7A",
1", "TMED5", "PSMB8", "PSMA6", "ZDHHC13", "PPP1CC", "ALG3", "UBE3B", "TRIM22", "ID2", "TAP1", "
4BPA", "BTG1") c("HBP1", "SSBP2", "HSPA1A", "RUNX1", "ETNK1", "AMACR", "RAB9A") character(0) c("
"TCEAL2", "LMO4", "MEIS2", "GADD45G", "PER3", "CRISP3", "ZMYM2", "DACH1", "JAK2", "PADI2", "SLC
AMA5", "SSPN", "AHNAK2", "GATM", "ZNF512B", "LPIN1", "CX3CL1", "KCTD3", "PGRMC2", "RXRA", "CA
BP4") c("IGF2BP3", "ZNF124") c("TF", "EIF3E", "ESF1", "MAN1A1", "RHOBTB1", "PCOLCE2") c("SERPINA
", "TBCB", "GALNT2", "POP7", "BTG3", "MT1H", "RASSF1", "ITGAE", "CDCA3", "TCFL5", "MEIS2", "SPRR2
1", "SEC61B", "ITGAE", "POLA2", "PSRC1", "SLC37A4", "HMMR", "CDC7", "GMNN", "SPRR2B", "ABCG2",
OD1", "LPL", "UCHL1", "CLIC4", "IGF1", "DIO2", "RASIP1", "HNRNPAB", "WFDC1", "CSRP1", "CHN2", "C3"
F2BP3", "HOMER2", "KLF11", "LRRC37A3", "ZNF721", "NT5E", "DDAH1", "CD34", "NELL2", "C2CD2", "D
PT1", "ISOC1", "GOLM1", "SPINK2", "PTN", "ZNF532", "ZNF395", "CDH2", "C2CD2", "WDR7", "ZBTB20",
"SATB2", "DOCK10", "CLIC4", "FKBP11", "ZDHHC11", "UBIAD1", "SPC25", "ALOX5", "SORBS1", "FXYD5",
1", "CTSH", "LAMC1", "ANXA5", "COTL1", "FN1", "AKR1B1", "LAPTM4B", "OPN3", "LAT2", "LASP1", "GN
MD1B", "SERINC5", "CDC123", "PUF60", "HSPA9", "B3GNT2") c("RRM2", "PRSS23", "CTDSPL", "CILP", "PI
", "SERPINB9", "CXCL10") c("MGLL", "NCAM1", "PRSS23", "HSP90AA1", "FBN2", "CTDSPL", "LGALS1", "C
INB9", "GBP1", "FCGR2A", "LCP2") c("MGLL", "PRSS23", "CORO1C", "VAT1", "SATB2", "EBP", "SRPK1", "
"RET", "PCYOX1", "SERPINB9", "GBP1") c("TFF1", "SERPINB2", "TGFA", "FARP1", "DTL", "PLOD2", "ZWIN
", "HPRT1", "FARP1", "FBN2", "FADS1", "SMC2", "JMJD6", "DPYSL2", "CORO1C", "CD55", "EMP3", "ANXA
", "BICC1", "COCH", "HERC5", "COL6A2", "LMCD1", "GPX1", "KCNQ2", "INHBB", "FSTL1", "IGFBP1", "SLC
"LRP4", "EMP3", "LAMC1", "ANXA5", "SPTLC2", "IGFBP6", "OPN3", "PCDH9", "ABCA3", "FKBP1B", "GNA
ABCA3", "CXCL9", "GNAI2", "UCHL1", "SLC7A8", "CLIC4", "TKTL1", "RASIP1", "SPARCL1", "DNAJC10", "P
, "GAS1", "CP", "SULF1") c("SMURF2", "HSP90AA1", "HSP90B1", "OLFM1", "HSPA8", "RFC2", "SPTLC2", "
3", "NEFM", "SNAP25", "IFIT1", "TMEM47", "PCYOX1", "ITIH5", "IGFBP3", "ADI1", "SERPINB9") c("MGLL
10", "GALNT2", "SORBS1", "RCAN2", "PRIM2", "A2M", "KLF11", "STEAP3", "PPID", "TCTN2", "OGFOD1", "
DSPL", "PIP4K2B", "CORO1C", "RRM1", "PMP22", "QKI", "PON3", "WIF1", "MCM6", "CXCL9", "VAT1", "C
"PRSS23", "HSP90AA1", "SOSTDC1", "NTRK2", "PIP4K2B", "CORO1C", "VAT1", "CALR", "MICAL1", "DOCK
PARCL1", "PDLIM3", "SORBS1", "SYNM", "PTGDS", "RCAN2", "PRIM2", "UBE2L3", "KLF11", "TEX10", "RA
, "MELK", "RAB31", "SEC14L1", "CD200", "RFC2", "PRDX1", "LTB", "GAB2", "MUC1", "PCDH9", "TPBG", "

'BCB", "NVL") c("NSL1", "SLC7A5", "SAP30", "NPDC1", "F2RL1", "RAB25") c("HLF", "DLEU1", "ESPL1", "M
", "PPP1CA", "MBOAT7", "AGPAT5", "VGLL4", "EIF3J") c("PBK", "GRWD1", "RRM2", "TRIP13", "PRIM2", "
'L", "BTG3", "COPS7A", "ABCG1", "RGS4", "SLC1A3", "TRIM22", "NCOA2", "PLLP", "MYLIP", "HOXB13", "L
", "CLNS1A", "RGS4", "TRIM22", "NCOA2", "ARL6IP5", "PIGT", "IKZF1", "ATP10B", "NDRG2", "FCGBP", "C
RBS3", "IL33", "L1CAM", "GNS", "ADAMTS5", "ITGAM", "IFIT1", "IFITM3", "IFITM2", "RUNDC3B", "CCL18"
"MUC1", "SPINK1", "LIMA1", "NID1", "PLN", "IFITM1", "HAPLN1", "RNF128", "PXDN", "PCCA", "VPS37B"
MEM63A", "RGS1", "VASH1", "CXADR", "MUC1", "ETV1", "JAKMIP2", "PLAT", "ITM2A", "TOX3", "TCN1", "
CF1", "SNAP25", "PELI1", "PCCA", "RCAN2", "SIRT3", "KCNMA1", "GSTO1", "HSD11B1", "MEG3", "PTPRD
'RCAN2", "SIRT3", "GSTO1", "BTAF1", "PTPRD", "EMP3", "FBN2", "NRXN3", "ACTR1A", "TFPI") c("ABCC3"
IGFBP1", "CYP2C18", "CSNK2B", "CAV2", "THSD7A", "HBD", "OMG", "ERBB3") c("ESR1", "AHNAK", "AHN
L1", "E2F2", "AP1B1", "ITGAE", "ABHD10", "PSAT1") c("HEBP2", "TPRKB", "PERP", "ZFAND1", "ERGIC2", "
", "HTRA1", "UPK1B", "MCAM", "IPO7", "SNCA", "JAG1", "ITGAE", "ABHD10", "TIMM17A", "AP3M2", "A
", "LMO4", "ESPL1", "CLU", "S100A8", "XIST", "CXCL14", "MT1G", "NTRK2", "KRT1", "PTPN3", "CRYAB", "
P36", "RAB9A", "FKBP5", "TAF4", "ZNF721", "PLXNA1") c("ID2", "FOXC1", "PSMB8", "LPCAT4", "TCTA") c
("RERE", "RGS4", "GLOD4", "KCNJ2", "CST3", "URM1", "AZGP1", "CORO1B", "PLLP", "ARL2", "CD320", "F
A2", "FBXO21", "E2F2", "ABCG2") c("NFKBIA", "XIST", "DUSP2", "PRNP", "SPAG16", "B3GALNT1", "ABHD
MTS5", "TOX3", "ARMCX3", "FUT8", "HELZ", "ACTA2", "TFIP11", "CXCL10", "L1CAM", "C7", "LAPTM4B", "
ANXA1", "CTBP2", "SYNM", "SIX2", "EMCN", "ABCC3", "ZNF706", "PRSS23", "LYZ", "ITGB1", "STAP2", "DC
'ABCC3", "ZNF706", "PRSS23", "COX8A", "OPHN1", "RET", "CORO1C", "IRF4", "TDP1", "SNAI2", "LPAR1",
GALS3BP", "TOP1", "COL15A1", "PXDN", "ABCC3", "PRSS23", "COX8A", "LRRC8D", "DCTPP1", "CORO1C"
OX", "SNAP25", "PHF10", "ZNF589", "PAK1", "NELL2", "MLH3") c("MCAM", "MUC2", "CXCL12", "AKT3", "
T") c("SLCO3A1", "GATM", "DSC1", "ARMCX6", "KRT2", "FLG", "LOR", "TM7SF2", "KIAA0355", "RNF5", "A
XNA1", "KLHDC2", "ETNK1", "DCK", "BNIP3", "SSBP2", "INHBA", "PANK3") c("ALDH2", "CKS2", "PPIC", "N
", "HS2ST1", "UBAP2L", "DEGS1") c("APEX1", "OLFML2A", "SIX1", "NVL", "SCAND1", "BTG1", "URM1", "
IP", "HBD", "CHRNA3", "CSNK2B") c("XIST", "CSTA", "CRYAB", "CAST", "AZGP1", "CALML3", "ZNF512B", "
RRN3", "ASXL2") c("CEACAM6", "ARHGEF3", "GYG1", "RBPMS", "ALDH2", "REPIN1") c("ZMAT3", "CLU",
", "INHBA") c("TRIM22", "ALDH2") c("IL7R", "CHD9") GPATCH8 c("MCOLN1", "LGALS3BP", "GNA11", "YN
L1", "FTH1") c("GULP1", "SH3BGRL", "GATM", "OXR1", "ZNF91", "SLC24A3", "SESN1", "TSPAN13", "TRPS
A", "VSNL1") c("TRIM22", "LGALS3", "ERBB3", "SLC7A5", "COMP", "HOXB7", "HBD", "CAV2", "ZBED1") c
CTA", "TM9SF2", "TAP1", "B2M", "CEACAM6", "HOXB7", "ZBED1", "NDUFA4", "ALDH2", "LST1") c("IL7R"

33" "V84" "V85" "V86" "V87" "V88" "V89" "V90" "V91" "V92" "V93" "V94" "V95" "V96" "V97" "V98" "V9
RUNX1", "XIST", "PLAGL2") c("LDB3", "GGCT", "SSH3", "SAC3D1") c("GNAI1", "DVL3") c("EIF5", "LRRFIP1"
NA1", "NFATC3", "RAB25", "TRIM22") c("AHNAK", "AHNAK2", "AKAP11", "ALDH6A1", "ANG", "AR", "AR
", "ERAP1", "ID2", "MLF2", "RCBTB2", "TRIM22", "ZDHHC13") c("ADD3", "CD302", "CLU", "HLTF", "IRS2"
AN7", "ZNF3") c("ACO1", "AKAP12", "ARMCX2", "BIRC3", "C7", "CYB5A", "EDNRA", "FABP7", "GIMAP6",
NCAM1", "NQO1", "STUB1", "MUC5AC", "KLK11", "CRABP2", "RBBP8", "S100A8", "GATM", "CSTA", "AL
AK2", "PRTN3", "IQSEC1", "RMND5B", "XIST", "GOLM1", "SCNN1A", "MAP3K1", "FKBP1B", "EIF3H", "SC
TNNT3", "RHOBTB1") c("GNAS", "PXMP2", "PNMA2", "CHFR") c("ITGBL1", "RBP4", "PDE4DIP", "DCN", "C
FRP1") c("GNAS", "ACP5", "AKAP1", "ACVR1", "GRB14") c("GREM1", "AGPAT5", "EIF5A", "STAG3", "CEP
", "RANBP6", "PCDH9") c("EIF5", "MDFIC", "CPVL", "CYCS", "IGF2BP3", "IL7R", "HSPA6") c("UCHL1", "SL
SR", "RAB25", "KBTBD11", "ROGDI", "NECAB3", "EFNA1", "TMPRSS3") c("CSTA", "S100A8", "CRISP3", "A
, "SMC2", "KIF2C", "CDCA3", "OIP5", "TCTA", "ABCG1", "LSM4", "MT1H", "NVL", "CXADR", "CYB561", "C
PRKB", "ENPEP", "TMPRSS3", "TSPAN7", "DHCR7", "RGS4", "ITGAE", "NDC80", "KIF2C", "OIP5", "HMGB2
", "TGM2", "LPL", "PCOLCE", "C7", "LRRC32", "DLEU1", "TIMP3", "FERMT2", "CHI3L2", "AIMP2", "WFDC1
TED2", "IL18", "MST1", "THY1", "LGALS1", "CD34", "IGF1", "NT5E", "CD320", "HOMER2", "CCDC85B", "Y
, "FGF9", "MTA1", "QPRT", "SCG5", "MPHOSPH8", "PDE9A", "MT1F", "PSIP1", "SERPINF1", "MDK", "CYP
KAP12", "RPS27L", "HERC5", "IFIT1", "FSTL1", "COCH", "HSD11B1", "PTGER3", "ZDHHC11", "FGF13", "M
P", "PHACTR1", "IPO5", "IFIT2", "FABP7", "IFITM3", "ABCA3", "OPHN1", "GDPD5", "IFITM2", "LTBP3",
SPARCL1", "DPY19L2P2", "ESM1", "XIST", "GLIPR1") c("SLC39A6", "RET", "ALOX5", "RPS27L", "HERC5", "
D3", "PRSS23", "AHSA1", "GINS2", "SPEN", "SFRP1", "HSD11B1", "FSCN1", "ADA", "ANXA6", "ZDHHC11",
ML1", "COL4A1", "PTER", "MST1", "SLC7A1", "YARS", "CD34", "PDIA4", "CCDC47", "GSTO1", "TRIP13", "
, "GDPD5", "DSTN", "LHFPL2", "IFITM2", "CYC1", "ASCC3", "PLTP", "HUWE1", "RNF10", "PPID", "IVD", "C

M15", "KLHDC2", "MAOB", "COL4A6", "NOTCH1", "PELI1") c("SNAP25", "ZMAT3", "PRSS23", "MGLL", "F
"CTNNB1", "ANXA5", "ACOT9", "GYS1", "PPID", "IVD", "EDNRA", "DPY19L1", "CLIP2", "IGLL1", "HLA-DPA
, "ME1", "DLAT", "MED27", "SIRT3", "PRIM2", "MACF1", "USP13", "MRC2", "IFIT2", "CORO1C", "IFITM3
HERC5", "IFIT1", "FSTL1", "HSD11B1", "PTGER3", "ADA", "ANXA6", "ZDHHC11", "PON3", "FGF13", "MST1
", "RGS5", "SYNE1", "PDE4A", "SPARCL1", "NR2F1", "RCAN2", "CD14", "GLIPR1", "SNCAIP") c("PCNA", "B

13", "PHACTR1", "SCG5", "IFIT5", "IFIT2", "SEMA3B", "ZNF83", "IFITM3", "ABCA3", "OPHN1", "GDPD5", "
"TGFA", "PLAC8", "AREG", "AKAP12", "PDZD8", "PLAT", "MGLL", "HERC5", "AOX1", "CXCL12", "COCH", "
VL5", "PELI1") c("PCNA", "SNAP25", "ZMAT3", "PRSS23", "MGLL", "FEN1", "FANCL", "ZNF337", "LOR", "
", "CBS", "RABAC1", "AHI1", "ASS1", "KIAA0753", "UBA1", "ABCA3", "CTSK", "DSTN", "CTSH", "ACOT9",
3", "GDPD5", "IFITM2", "CTNNB1", "EIF4EBP1", "ANXA5", "GYS1", "HUWE1", "CLIP2", "CPT1A", "ESM1",
", "OPN3", "COL6A2", "TEX10", "CNN3", "CILP", "COX7A1", "ADAMTS1", "C7", "AKR1B1", "ACTA2", "CD5
PINA1", "IFIT1", "IGFBP6", "IFI44L", "FSTL1", "HSD11B1", "PTGER3", "ADA", "DSC3", "ANXA6", "ZDHHC1
EIF2S3", "PON3", "TFPI", "SERPINB9", "MST1", "HSPA13", "DNAJC10", "CD34", "DIRAS3", "GSTO1", "CLIC
P2", "SSX1", "ELOVL5", "PELI1") c("SNAP25", "RPS27L", "PRSS23", "MGLL", "FANCL", "LOR", "SPEN", "IFI
2", "APP", "PXDN", "RBM38", "TSPAN1", "SPATS2L", "RNF10", "TAGLN", "GAS1", "SPARCL1", "PLXND1", "

"NBL1", "NELL2", "SLCO2A1", "USP13", "IFITM1", "IPO5", "NEFM", "IFIT2", "FABP7", "IFITM3", "ABCA3"

BR2", "ASCC3", "PXDN", "MATN2", "GYS1", "HUWE1", "RNF10", "IVD", "EDNRA", "MFAP4", "MAOB", "C
SERPINB9", "CGA", "KIF18A", "IRS1", "SPON1", "TCF4", "PCOLCE", "GRWD1", "PRIM2", "MACF1", "WFD
T5A", "MRC2", "MPHOSPH8", "IFIT2", "MT1X", "ABCA3", "OPHN1", "LTBP3", "SCD5", "ZNF331", "NAV2"
DDB2", "NCAM1", "TGFA", "CXCL5", "CXCL6", "WT1", "RBP1", "ITGA3", "PDGFC", "RRM2", "GBE1", "CE

.GALS1", "DNAJC10", "CD34", "NT5E", "GSTO1", "PCOLCE", "MEF2C", "GRWD1", "LPAR1", "ODC1", "ME1
.3", "EIF3J", "MUC2") c("TCN1", "SNAP25", "NR3C1", "TFF1", "NOVA1", "RPS27L", "RRM2", "PRSS23", "R.
2L") c("SNAP25", "ZMAT3", "TFF1", "PRSS23", "MGLL", "FANCL", "ZNF337", "LOR", "SPEN", "IFIT1", "GAM

", "PCP4", "ABCA3", "DSTN", "LTBP3", "PDE4D", "LGALS3", "WSB1", "TRAK2", "PLTP", "MATN2", "TBC1D
TLR4", "CHI3L1", "MELK", "DONSON", "TOX", "TCTA", "CXADR", "SHMT2", "SUPT3H", "MUC1", "HNRNPA
SAP30", "HBD", "KBTBD11") c("CSTA", "LIG1", "CRISP3", "TNFSF9", "S100A8", "CLIC3", "ESPL1", "TSPAN1
IT5A", "MT1M", "GAS1", "THAP11", "SLC27A3", "NVL", "KCNJ2", "ITGAE", "SMPDL3A", "RRAGD") c("RAB
B", "INHBA", "TOPBP1", "DCK", "PANK3", "CD55", "XIST", "DUSP2", "HSPA1A", "RUFY2", "KCNK3", "WDF
D2", "IFIT1", "IFITM1", "ISG20L2", "MXRA5", "PLTP", "OGT", "ABCA3", "RAB4A", "SHOX2", "PTEN", "VAT
5A", "MUC2", "FAM114A1", "APIP", "ITGAE", "MAFB", "PELO", "NDUFA4L2", "CRYL1") c("S100A11", "UC
2") ACVR1 c("EIF5A", "SORBS3", "NUP93", "GREM1", "MBNL1", "ODC1", "ATP2B1", "STAG3", "NTRK2", "
C", "GATM", "MGLL", "DDX42", "RNF128", "CUEDC1", "OGFRL1", "MEIS2", "CALML5", "ZFP36L1", "CRISP
", "NUDT11", "PRTN3", "CRABP2", "LPIN1", "PLD3", "YRDC", "MTA1", "SLC7A8", "IRX5", "S100A8", "TRP
IK2", "RNF128", "ZMYM2", "MGLL", "GATM", "NISCH", "KLHDC2", "GAMT", "AOX1", "TM7SF2", "LOR", "
", "CLU", "MFAP2", "SORT1", "COL4A5", "MEIS2", "CRYAB", "RAB40B", "ALDH6A1", "TECR", "COG5", "PL
GALS1", "UCHL1", "MICAL1", "DIRAS3", "PMP22", "EMP3", "RHOBTB1", "PDE4D", "TIMP1", "SLC47A1", "
1", "CXCL9", "SORBS1", "SCG5", "COL4A6", "SPARC", "NR3C1", "USP13", "SIRT3", "LPAR1", "GM2A", "PB
'WSB1") c("GAS6", "MFAP2", "CD83", "RBPJ", "SOAT1", "WSB1") c("CTBP2", "GAS6", "ENO2", "P2RX4", "
"BCL11B", "DES", "TBCB", "GAS1", "NVL", "KDM5A") c("CLDN4", "COMP", "TCL1A", "TRIM22", "EFNA1",
SERPINA5", "CEACAM6") c("ADD3", "P4HTM", "ANKRD28", "PCDH9", "PTX3", "BIRC2", "STK32B", "CLU",
GNAI1", "TFDP2", "ERCC1", "FEZ1") c("STC2", "FEZ1") c("HSPA6", "NOTCH2") c("ARMCX1", "TFDP2", "PC
O3A1", "HNRNPU", "MT1G", "PRG2", "ACAA2", "BEX4", "OSBPL1A", "ACVR1B", "CX3CL1", "CXCL14") c("
ST") c("TNFSF10", "C3", "CASP1", "DHCR24", "FCGBP", "IKZF1", "UBE2E3", "ZNF512B") c("XIST", "RUNX1
1", "AKAP1", "ANKRD27", "NFE2") c("EIF5A", "XK", "NOP56", "MACF1", "AGPAT5", "PTCH1", "RBP4", "CC
B", "THAP11", "IFIT1", "TBX2", "STAT5B", "RAD51C", "CYB561", "WNT5A", "GAS1") c("SLC7A5", "ITGA3",
"TNFSF10", "OLFML2A", "SKP2", "TRPM4", "CYB561", "FCGBP", "ADI1", "APEX1") c("HSPA1A", "TUBB2A
', "STAC", "NNMT", "PCOLCE", "AKT3", "RER1", "DLEU1", "BCCIP", "CHN1", "PGR", "GOLM1", "TGM2", "(
HRS2", "MSX1", "PKP1", "SLC44A4", "ITIH5", "ARHGEF18", "PADI2", "DCLRE1C", "OXA1L", "CREB1", "ISL1

"KLF11", "PPID", "COCH", "HERC5", "LMCD1", "FSTL1", "IGFBP1", "SNAP25", "IFIT1", "RET", "KHDRBS3",
AI2", "HHEX", "RAP2A", "SATB2", "PPIF", "CLIC4", "HBEGF", "ZDHHC11", "UBIAD1", "SPC25", "ETV5", "AL
P4K2B", "CORO1C", "EPHX1", "PMP22", "FN1", "MAPRE2", "ITPR3", "TPX2", "JAM3", "GSPT1", "PON3", "
ORO1C", "EMP3", "ANXA5", "PMP22", "SGCB", "PON3", "WIF1", "ABCA3", "VAT1", "HHEX", "CCDC88A",
LOR", "DNAJC10", "PRIM2", "MAOB", "PPID", "OGFOD1", "PCOLCE", "RASSF1", "OPHN1", "LTBP1", "SHO
IT", "MUC1", "KIF11", "RAP2A", "UBIAD1", "SF3A2", "SSX1", "SLC37A4", "HERC5", "CHI3L2", "IFIT1", "ML
\5", "PMP22", "SERPING1", "QKI", "DHCR7", "ABCA3", "GNAI2", "PODXL", "HHEX", "CALR", "CCDC88A", "
35F2", "CNN3", "IGFBP5", "IFIT1", "ALDH1A2", "RET", "RGS5", "SERPINB9", "CD53", "NXF1", "CP", "CXCL
\I2", "TK1", "ELOVL5", "ADCY9", "HHEX", "EPB41L2", "GPM6B", "CCDC88A", "DKK3", "HOPX", "MFHAS1",
RIM2", "KLF11", "PPID", "RASSF1", "BICC1", "OPHN1", "TDP1", "LTBP1", "CD34", "SHOX2", "PLTP", "PDE
NFE2L1", "QKI", "C1QB", "MLXIP", "PODXL", "TPBG", "SDC3", "PLOD1", "IRS2", "ZDHHC11", "SNHG3", "\
", "PELI1", "PRSS23", "HSP90AA1", "HPRT1", "FARP1", "FADS1", "CORO1C", "ANXA5", "ABCA3", "ELOVL5
'PCOLCE", "RASSF1", "SSX1", "NT5E", "LTBP1", "CD34", "SHOX2", "NELL2", "PLTP", "PDE4A", "LPAR1", "A
ALR", "TRIP13", "MICAL1", "SERPINH1", "NOVA1", "LRIG1", "SFRP4", "UCHL1", "SPARC", "SQLE", "SNHG
10", "UCHL1", "LOR", "DNAJC10", "MAOB", "OGFOD1", "PCOLCE", "RASSF1", "LTBP1", "PRKD3", "SHOX2
SSF1", "BICC1", "TBC1D1", "CD34", "COL6A2", "SGCE", "NBL1", "PGR", "PLTP", "LPAR1", "ABCC3", "KCNC
CHI3L1", "PER2", "DONSON", "SLPI", "LRRC37A3", "TLR4", "SEL1L", "MB", "TWIST1", "RGS5", "MS4A6A",

T1G", "NSL1", "PHGDH", "GOLM1", "MPHOSPH8", "DHTKD1", "MEIS2", "PSPH", "NUCB2", "HSDL2", "RT
"MCM6", "RRM1", "UCHL1", "IFI6", "TIMP1", "USP13", "PMP22", "PPID", "SFRP4", "SPARC", "IKBKE", "SE
LRRC1", "PIGT", "CST3", "DES", "NDRG2", "OAS1", "NPDC1", "SMPDL3A") c("NFATC3", "GRAMD1C", "ITG
ASP3") c("MRPL40", "AMACR", "B3GALNT1", "KLHDC2", "COMMD3", "IRS2", "UBE2G2", "DCK", "ZNF72
", "TOX3", "ZDHHC11", "BST2", "CXCL10", "EDNRA", "RGS5", "SNAP25", "PELI1", "HEPH", "GPM6B", "EY
) c("LTBP1", "TGFBR2", "VIM", "MYH11", "LTBP2", "IL33", "CP", "PMP22", "CCL19", "FHOD3", "TGFB1I1"
"TBX3", "KLK3", "NUPR1", "EDNRA", "APOA2", "HGF", "HAPLN1", "CXCL12", "GPM6B", "VPS37B", "SAT1"
", "MAP2K5", "PROX1", "CITED2", "TFPI") c("GREM1", "COL14A1", "LSM2", "VASH1", "MBOAT7", "TME
, "CYC1", "NRP1", "SPRY2", "MGLL", "IFITM3", "IFITM2", "TACSTD2", "CXADR", "CLNS1A", "MUC1", "PL
AK2", "NUCB2", "GOLM1", "MGLL", "DECR1", "SLC2A10", "SCNN1A", "TPD52", "ZBED2", "MYL6B", "CYB
"TUBB2A", "ABHD6", "HIGD1A", "IRS2", "LRRN3", "MRPL40", "UBE2E1", "PLS3", "POLR1D", "FKBP5", "C
POA2") c("SGCE", "QPRT", "SERPINB9", "PON3", "ERCC1", "SPON1", "DIRAS3", "SAR1B", "SERPINB2", "U
PHGDH", "NASP", "CALML3", "TRHDE", "ESR1", "MSX1", "EIF3H", "NCAM1", "GATM", "TSPAN13", "NQ
("MSX1", "ZFP36L2", "BCL10", "PTX3", "KIAA1324") c("STC2", "GDPD3") c("BCL10", "YTHDF1", "SKAP2",
RAD51C", "FOXJ3", "TAF5", "THAP11", "TBP", "NVL", "MYLIP", "SHROOM2", "COPS7A", "HELZ", "TCTA", "
6", "IRS2", "TAF4", "JARID2", "HSPA1A", "PANK3", "GGNBP2", "TFEC", "COMMD3", "PLAGL2", "SIAH1", "
"ALOX5", "COCH", "INPP4B", "SYCP2", "TRIAP1", "GDPD5", "FKBP11", "SNAP25", "PHF10", "ZFP64") c("U
TPP1", "CORO1C", "IRF4", "DPYSL2", "ZCCHC24", "PARP2", "MED13", "ZCCHC2", "EIF4ENIF1", "IFI44L", "
"ACTA2", "TFIP11", "CXCL10", "JAM3", "CXCL9", "ZNF337", "TMEM43", "PDGFA", "ACOT9", "MEF2C", "
, "CCNE2", "FOXJ3", "DNAJC12", "TAOK3", "IRF4", "IFI44L", "SNAI2", "LPAR1", "TFIP11", "CXCL10", "MA
"COL14A1", "SEL1L", "NARS", "PPP1CA", "FDFT1", "ARL2", "GREM1", "SLC35A1", "TNC", "PELO", "UPF1",
KAP7", "TECR", "JAG1", "WNT5A", "OXA1L", "TST", "CNNM3", "TMED3", "SLC7A8", "GNPDA1", "GOLGB
DUFA4", "PECR", "RBPMS", "MLF2", "EHD3", "BBX", "NPAT", "GLCE", "LAP3") c("RIF1", "CLU", "SIX2", "C
BANP") c("NFKBIA", "HSPA1A", "ZNF277", "PANK3", "THOC2", "CD55", "SIGLEC15") c("ARHGEF3", "NRB
PRTN3", "MAGEH1", "ALDH3A2", "PRG2", "CLU", "BEX4", "CALML5", "FBXL5", "COG5", "JAG1", "PKP1",
"FTO", "PLCB4", "PDE8B", "PTX3", "ZFP36L2", "KIAA1324", "IL7R", "ALDH1A2", "SMC1A", "IRS2", "LYST",
ME1L1", "GPA33", "IL7R", "SERINC1", "MS4A1") c("GNA11", "EML3", "OCA2", "UCHL1", "PLCD1", "FKBP1
1", "NEBL", "SH3GLB2", "RAB40B", "SYNJ2BP", "CLU", "PACSIN2", "AHR", "CRISP3", "S100A8", "CPB1", "
("STUB1", "CXCL14", "XIST", "ZNF91", "ARMCX2", "TM7SF2", "HOXB6", "CCNG1", "NEBL", "AZGP1", "W
, "DVL3", "ZMAT3", "SMC1A", "KLF2", "CLU", "IRS2", "AFF1", "CTSB", "PTX3") c("STC2", "UPK3B", "PI3", "

)9" "V100" "V101" "V102" "V103" "V104" "V105" "V106" "V107" "V108" "V109" "V110" "V111" "V112" '
") c("MAGEA6", "AMIGO2", "TLR2", "EIF5", "FKBP14", "ADAM17", "IGF2BP3", "YTHDF1") c("SFRP1", "DL(
MCX2", "ARMCX6", "BEX4", "BNIP3L", "C2CD2", "CCDC28A", "CD36", "CLU", "COG5", "COL4A5", "CROT"
, "MSX1", "SORL1", "WSB1", "ZFP36L2", "ZFPM2") c("EIF5", "LDOC1", "RAD54B", "WSB1", "WWOX") c("I
"GRK5", "GSTM3", "HERC5", "IFIT1", "ITM2A", "NID1", "PARP2", "PDGFRL", "PMM1", "PPP2R1B", "RGS!
DH3B1", "WNT5A", "NTRK2", "PSMD12", "NEBL", "TRHDE", "LIG1", "PRTN3", "FAM173A", "ALDH5A1", "
ARB2", "NUDT11", "GULP1", "CCDC28A", "CLU", "RTN3", "SOSTDC1", "AR", "PHACTR2", "CUTA", "ERMP1
CSGALNACT2", "SARS", "CDC14B", "NFIB", "PSAT1", "DSP", "KCNK2") c("PDE4DIP", "CEBPG", "CD1B", "ELI
68", "RBP4") c("CD1B", "STAG3", "CEBPG", "GRB14", "DIAPH2") c("KLF11", "HOXA10", "FGFR1", "CDO1",
T2") c("S100A9", "GRB14", "PTDSS1", "SERPINA1") c("UCHL1", "ITGAE", "HIP1", "SLIT2", "HSPA6", "CAV1
AZGP1", "MSX1", "TSPAN13", "BCHE", "NCAM1", "DNAJB2", "MTA1", "FOXO4", "PRTN3", "MED21", "BLV
)AS1", "WNT5A", "IFIT2", "MX2", "RIMS3", "CST3", "SLC25A46", "FBXO21", "ID2", "RNF130", "EIF4B", "N
", "TCTA", "PSRC1", "HMMR", "LSM4", "NVL", "CYB561", "CDC7", "FCGBP", "PMS1", "SLC25A11", "SLC25
L", "MS4A4A", "HEY1", "KHDRBS3", "C1QB", "HYOU1", "RABAC1", "MRC2", "HEPH", "SERPINF1", "CTSK",
RDC", "CRNN", "MDC1", "CSF2RA", "NELL2", "GTF3A", "AHI1", "PDCD4", "ZFPM2", "PRTN3", "AKAP8", "I
4B1", "SCD5", "TCEAL2", "WDR7", "CACNA2D3", "NAV2", "FBP1", "CTSH", "METTL7A", "SORL1",
ST1", "FKBP11", "HSPA13", "GPM6B", "FXYD5", "DIRAS3", "CLIC4", "CDH11", "GRWD1", "C7", "ACTA2",

ETV5", "PPIF", "IFIT1", "FSTL1", "COL5A2", "PTGER3", "ANXA6", "ZDHHC11", "FGF13", "MST1", "LGALS1"
"SERPINB9", "KIF18A", "FGF13", "JMJD6", "PODXL", "HSP90B1", "DNAJC10", "CLIC4", "OPN3", "HSPH1",
CLIC4", "RASIP1", "ENPP2", "MICAL2", "PMP22", "COL6A2", "CXCL9", "PCOLCE", "LPAR1", "CILP", "ME1",

ANCL", "ZNF337", "LOR", "SPEN", "IFIT1", "GAMT", "HSD11B1", "SERPINB9", "DNAJC10", "GSTO1", "PCOI

", "GDPD5", "ASCC3", "PPID", "EDNRA", "STEAP3", "SPINK1", "HGF", "COL21A1", "MAOB", "SSX1") c("SN
.", "DPYSL2", "DNAJC10", "FXYD5", "GSTO1", "CLIC4", "CDH11", "ENPP2", "OPN3", "PCOLCE", "GRWD1",
CHE", "KRT2", "AKR7A2", "FANCL", "LOR", "GAMT", "ITM2A", "SERPINB9", "CITED2", "IGF1", "PCOLCE", '

'IFITM2", "CPOX", "MATN2", "HUWE1", "PPID", "IVD", "EDNRA", "STEAP3", "ROGDI", "PITX2", "SSX1", "P
'ITM2A", "NDUFA4L2", "SCML1", "SERPINB2", "GPM6B", "BET1", "NDRG4", "PPP2R5E", "KLK6", "VPS37B
SPEN", "IFIT1", "GAMT", "HSD11B1", "SERPINB9", "THY1", "DNAJC10", "DIRAS3", "GSTO1", "PCOLCE", "G

"ENO2", "ATP2A3", "ROGDI", "SYCP2", "ELOVL5") c("RPS27L", "PRSS23", "OLFM1", "IFIT1", "ANXA1", "IF
3", "A2M", "FHL1", "IGFBP1", "KCNQ2", "CXCL10", "IPO5", "DKK3", "SATB2", "IFIT2", "HEPH", "IFITM3",
1", "PCDH9", "JMJD6", "DPYSL2", "USP18", "FKBP11", "GPM6B", "LGALS1", "HOPX", "HSP90B1", "CD34"
4", "RASIP1", "PMP22", "CXCL9", "MEF2C", "GRWD1", "LPAR1", "CILP", "YRDC", "ACTA2", "MTMR2", "H
T1", "GAMT", "HSD11B1", "SERPINB9", "DNAJC10", "DIRAS3", "GSTO1", "CLIC4", "PCOLCE", "GRWD1", "I
'SPRY4", "CNR1", "STC1", "PNMA1") c("HSPA8", "WT1", "IRS2", "FHOD3", "OLFM1", "RFC2", "SDC3", "SP

, "OPHN1", "IFITM2", "TCEAL2", "PDE4D", "TRAK2", "PLTP", "MATN2", "HUWE1", "IVD", "RGS5", "CD81'

OL4A6", "PITX2", "NOTCH1", "PELI1", "LRRN3") c("SNAP25", "RET", "RPS27L", "PRSS23", "MGLL", "AHSA1
C1", "USP13", "UNG", "MT1F", "EIF4EBP1", "PXDN", "MLEC", "MAN2B1", "CD81", "SPINK1", "XAB2", "AD

ACAM5", "CDT1", "NPY1R", "IFI6", "PCOLCE2", "ADA", "OLFML3", "KRT17", "DSC3", "PON3", "TFPI", "SRP

.", "C7", "MED27", "SIRT3", "TIMP3", "SPC25", "A2M", "PRIM2", "NELL2", "MACF1", "USP13", "PHACTR1
AD51C", "PBK", "LOR", "SPEN", "IFI6", "WIF1", "OLFML3", "PON3", "SERPINB9", "SLC47A1", "SERPINB2",
MT", "SERPINB9", "NTRK2", "DNAJC10", "SPON1", "DIRAS3", "GSTO1", "PCOLCE", "GRWD1", "LPAR1", "M

A2B1", "NPC1", "HSPE1", "HAPLN1", "MB", "SET", "RGS5", "S100P", "MGAM", "TPBG", "RBM22", "NDUF
.3", "PHGDH", "MKNK2", "SESN1", "PPP1R3C", "DHRS2", "HOXA10", "BNIP3L", "CCNG1", "GATM", "NCA
25", "TRIM22", "HOXB7", "KDSR", "COMP", "MYCN", "FTH1", "CHRNA3", "F2RL1") c("GNPDA1", "BEX4",
R61", "ERAP1", "POLR1D", "DSC2", "TUBB2A", "HBP1", "BNIP3", "FKBP5") c("TRIM22", "ALDH2", "HLA-A
1", "EDNRA", "LGALS1", "SCUBE2", "LTBP1", "SNAP25", "CBX6", "SPON1", "TIMP1", "FERMT2", "NRXN2
HL1", "SFRP4", "MTHFD2", "SCG5", "IFITM1", "TCN1", "LXN", "CXCL9", "MXRA5", "PLTP", "OGT", "IL1RL
NDFIP1") c("SORBS3", "UPF3A", "STAG3", "PSMD12") c("HOXA10", "TARP", "CD40", "ERP44", "PPP3CA",
3", "ACPP", "LOR", "ISL1", "IGF1", "ESR1", "XIST", "CXCL14", "DACH1", "GOLM1", "LRBA", "CROT", "COL
S1", "DACH1", "CPB1", "CSTA", "CXCL14", "CD36", "JAG1", "SOSTDC1", "ARMCX6", "ZNF512B", "PKP1", "
FBXL5", "CD36", "CST6", "MEIS1", "SLC24A3", "TRHDE", "NDP", "CLU", "NQO1", "NR1D2", "PADI2", "AG
D3", "KLK11", "CRISP3", "ERP44", "ANGPT1", "MEIS1", "NTRK2", "MPHOSPH8", "DYNLT1", "NQO1", "AC
MACF1", "GSTO1", "IFITM2") c("CLIC4", "IVNS1ABP", "GBP1", "RGS5", "KCNMB1", "ITM2A", "ZDHHC11"
K", "MRPL18", "SGCE", "SFRP4", "UCHL1", "MICAL1", "TNNC2", "DIRAS3", "PMP22", "RHOBTB1", "TIMP
SLIT2", "RBPJ", "IRX5", "KCNK2", "HGSNAT") c("STAT3", "CCL20", "ETS2", "CA10") c("TARBP1", "ATP2B1
"HAMP", "F2RL1", "SERPINA5", "KDSR") c("BEX4", "CKB", "OXA1L", "PHACTR2", "RAB4A", "COL4A5", "C
"FTO", "SMC1A", "DVL3", "SORL1", "ZMAT3", "RBM15", "PDE8B", "CTSB", "MSX1", "TARP") c("PI3", "PC
OLCE2", "NME4", "NOTCH2", "CEBPG", "BTBD3", "ANXA9") c("STAT3", "SELENBP1") c("HSPA6", "ITGAE",
TNFSF10", "AZGP1", "TSPAN7", "TRIM22", "FBXL5", "ACSL1", "PSAT1", "IKZF1", "EGF", "ITGAE", "TNS3")
", "SIGLEC15", "NINJ2", "ZFAND1", "SSBP2", "CD55", "IRS2", "LRRN3", "KCNK3", "FOXO1") c("SLC1A3", "
L21A1") c("ESF1", "UPF3A", "RHOBTB1", "QKI") c("PFKFB3", "LEFTY1", "TARP", "MAN1A1", "MDN1", "P
"NFATC3", "PPIF", "PCSK6", "CAV2", "SERPINA5", "LIF", "SAP30", "IGFBP1", "LGALS3") c("MGLL", "ZNF3
.", "PELI1", "ADCY1", "PHIP", "MALT1", "MTF2", "GADD45B", "CD55", "THOC2", "RAPGEF2", "BNIP3", "IR

.") c("ASRGL1", "STAR", "CPM", "MLXIP", "ACTG2", "FILIP1L", "RCAN2", "BICD2", "PPID", "CA4", "MMP1"

"RGS5", "OXA1L", "CXCL10", "SULF1", "FCGR2A") c("TCEA2", "MGLL", "PRSS23", "HSP90AA1", "SOSTDC
OX5", "GALNT2", "SORBS1", "SYNM", "FXYD5", "PPID", "PTPN3", "ENPP2", "LTBP1", "HERC5", "LMCD1",
CXCL9", "GNAI2", "OIP5", "TRIP13", "TIPIN", "PCOLCE2", "CCDC47", "MICAL2", "SLC7A1", "CLIC4", "RAS
"SFRP4", "UCHL1", "LOR", "DNAJC10", "PTGDS", "MAOB", "KLF11", "PCOLCE", "RASSF1", "BICC1", "LTBP
X2", "ABCC3", "NOTCH1", "SNAP25", "IFIT1", "SERPINB9") c("MICB", "PELI1", "PRSS23", "SCG5", "FARP1
LT3", "RET", "IDE", "PLN", "FCGR2A") c("NCAM1", "PRSS23", "SFRP1", "MCAM", "S100A4", "CCDC92", "E
TTF2", "NFASC", "SNRPA", "G3BP1", "CLIC4", "SPARC", "ZDHHC11", "SNHG3", "IFI44L", "DNAJC10", "PD

, "CLIC4", "FKBP11", "ZDHHC11", "UBIAD1", "SCO2", "IFI44L", "CSRP1", "SYNM", "COL15A1", "SERPINB5
A", "LPAR1", "ABCC3", "LGALS3BP", "SIRT3", "NOTCH1", "TNS1", "IFIT1", "RET", "SERPINB9", "CXCL10")
VT1", "TESC", "DDX10", "TEX10", "SUSD4", "ENO2", "DPH2", "OPHN1", "SIVA1", "PTP4A2", "PLTP", "KC
", "PODXL", "VAT1", "CALR", "SATB2", "HMGCS1", "G3BP1", "EBP", "CLIC4", "ZDHHC11", "SNHG3", "UB

3", "LOR", "DNAJC10", "SORBS1", "PTGDS", "PRIM2", "RANBP1", "NR3C1", "STEAP3", "PPID", "OGFOD1",
", "TMPO", "NELL2", "LPAR1", "ABCC3", "ZNF22", "NOTCH1", "UNG", "NEFM", "SNAP25", "IFIT1", "SERP

"SETD5") c("NPY1R", "LBR", "GLDC", "HPRT1", "FADS1", "MARCKS", "DPYSL2", "S100A10", "EMP2", "AC

N3", "NTRK2", "LMO4", "TSPAN13", "GLRX", "KLK11", "NCAM1", "TAX1BP1", "BCHE", "JAG1", "CAST", "C
ERPINB2", "SQLE", "GSTO1", "DIRAS3", "EIF4EBP1", "VAT1", "TCN1", "STEAP3", "MACF1", "PON3", "CALR
A3", "MMRN1", "CSAD", "TRIM22", "HOXB7", "NPDC1", "RAB5B") c("MEIS1", "CLU", "DACH1", "HPGD",
1", "PELI1", "THOC2", "LRRN3", "ADCY1", "CGA", "XIST", "PERP", "PLS3", "KIAA0232", "ANKRD12") c("ND
A1", "KCNMA1", "HSD11B1", "SATB2", "FGF13", "PTPRD") c("SLIT2", "LTBP1", "L1CAM", "CP", "ABCC3", "
", "NRP1", "MMRN1", "NR3C1", "PCLO", "GNS", "PPP3CA", "RCN1", "SMAD2", "REEP1", "VCAN", "DAB2"
", "NSF", "KYNU", "KCNMA1", "AREG", "TNFAIP8", "PDZD8", "PRDX3", "MAP2K5", "PINK1") c("SLIT2", "LT
M140", "SNCA", "PTMS", "CIT", "IPO7", "TNC", "TBX3", "MCAM", "PAEP", "PPT1", "APOA2", "HGF", "MU
AT", "TCN1", "DOCK9", "PCMTD2", "RNF128", "NDUFS7", "PCCA", "KCNMA1", "HSD11B1") c("TLR4", "SN
561", "CRISP3", "FOXO4", "MSX1", "CLIC3", "S100A8", "CALML5", "DYNLT1", "CLU", "C2CD2", "PEX3", "C
GA", "HNRNPH1", "ZNF721", "PELI1", "COMMD8", "COMMD10", "XIST", "RSL24D1") c("NDUFA9", "SERPI
CHL1", "IFITM3", "CXCL9", "STC1", "SFRP4", "ZNF580", "GM2A", "SNAI2", "CXCL10", "IFI6", "S100A11", "
D1", "PRG2", "CITED2", "CPB1", "NR1D2", "VIPR1", "MBTPS1", "CHPT1", "TRPS1", "ISOC1", "WIF1", "IRX5
"IGF2BP3") c("PPARG", "SFRP1", "GDPD3", "KLF4") c("SERPINA1", "HNRNPA1", "SERTAD3") c("NOTCH1"
"SLC1A3", "DST", "TMEM14A", "LEPROTL1", "GDE1", "REEP1", "ABCG1", "LRRC1", "NCOA2", "MT1M", "M
"LRRN3", "RAB9A", "FOXO1", "AMACR", "FKBP5", "TMEM43", "SIGLEC15", "SGMS1", "RUNX1", "KIAA023
UCHL1", "HSP90AA1", "MGLL", "SLC47A1", "SIX2", "EMCN", "ABCC3", "PRSS23", "COX8A", "DCTPP1", "CC
"SLC20A1", "SNAI2", "PTGDS", "LPAR1", "ATP6V1C1", "ACTA2", "SERPING1", "C3AR1", "TMEM176B", "L1
SCUBE2", "PMP22", "ITM2A", "PHF10", "MLH3") c("MGLL", "SIX2", "PXDN", "CLTC", "ABCC3", "PRSS23",
DB", "UBE4A", "VEGFA", "ACOT9", "NUSAP1", "CD83", "SNAP25", "PHF10", "ZNF589", "MLH3") c("HSP90
, "NUP62", "DKFZP586I1420", "SNX5", "TMEM11", "TBX3", "TCTA", "PRCC", "PPP1R10", "TRIM29", "RSL
1", "HBP1", "ENO2", "ARMCX2", "SHMT1", "AHR", "DDAH1", "DHRS2", "CLU", "KLHDC2", "ESR1", "CSTA"
AMKK2", "GNAI1") c("PTPN13", "TMEM134", "GTF2I", "CTBP2", "WWOX", "STC2") c("PTPN13", "HSPA6"
F2", "PCGF1", "STAM", "LST1", "CKS2", "TSPAN14", "PTPRO", "SAP30", "NPAT") c("ZFP36L2", "PSMD12",
"CPB1", "MOAP1", "CRISP3", "PLCB4", "TSPAN13", "CKB", "CDKN1B", "OGFRL1", "PHYH", "NEBL", "NQO
, "SORL1", "DVL3", "CHD9", "GNAI1", "PCDH9", "FEZ1", "SEMA6A", "ZFPM2") c("RNF144A", "STC2", "FTO
LB", "SLC7A8") c("SELENBP1", "TLK2", "ETS2", "TNFRSF17") c("GNA11", "GGH", "CHP2", "UCHL1", "CCDC
"DHRS2", "MEIS1", "C2CD2", "FGGY", "DACH1", "BCHE", "VPS37C", "FOXO4", "HOXB6", "HPGD", "KIF1B",
NT5A", "CITED2", "IRX5", "DDX42", "TNPO2", "UBR5", "EXPH5", "KLHDC2", "S100A8", "CLU", "CLIC3", "TS
"AFF1", "RNF144A", "CTBP2", "PTPN13", "RNH1") c("LGALS3BP", "IL7R", "CITED2", "ZMAT3", "HSPA6", "

"V113" "V114" "V115" "V116" "V117" "V118" "V119" "V120" "V121" "V122" "V123" "V124" "V125" "V1
G5", "IRX5", "MFAP3L", "ELF5", "PPARG") c("CCNE2", "STAT3", "POLR1B") c("CDC25B", "TAOK1") c("TGF
, "CXCL14", "ELF5", "ENO2", "ERLIN2", "ESR1", "GATM", "GNPDA1", "HBP1", "HLF", "HOXA5", "HPGD", "
EIF5", "GINS1", "HOXA5", "HSPA6", "IL6R", "LGALS3BP", "MCOLN1", "PTGER4", "RBM4", "RSBN1", "SKAI
5", "SMC2", "SPARCL1", "TKTL1", "ZSCAN18") c("AKAP11", "ARL6IP5", "BIRC3", "BTK", "CLIC2", "GRK5", "
MGLL", "XIST", "MEIS1", "PHGDH", "LOR", "DAAM2", "DLEU1", "ESPL1", "CRYAB", "MSX1", "JAG1", "SOS
L", "NUMA1", "LRRC37A3", "SETD2", "CNNM3", "TSPAN13", "CHPT1", "MFAP2", "MGLL", "CX3CL1", "GLI
F5", "NFIB", "USP13", "RHOBTB1") c("PTN", "CYP1B1", "ERCC2", "KLF6") c("DLC1", "PDGFRA", "DCN", "C
, "KLF6") c("VAT1", "HMOX1", "AKAP12", "IGFBP5", "PDZRN3", "DLC1", "GNAQ", "ARG2", "CEP68", "COL
L") c("USP7", "GRB14", "IL6ST") PTN c("HMOX1", "HLA-DPB1", "CPVL", "MLLT3", "HSPA6", "MRPS10", "T
/RB", "GATM", "MYL6B", "DACH1", "ALDH6A1", "PPP1R3C", "NUCB1", "HPGD", "CLU", "MT1G", "CD36",
PDC1", "TBX2", "GAS1", "SMPDL3A", "XAB2", "PIGT", "ROGDI", "SALL2", "TCFL5") c("CAV2", "ITGA3", "SI
5A46", "AGTPBP1", "IGFBP2", "E2F2", "FBXO21", "ID2", "ABCG2", "P4HB", "PCTP", "NDUFA4", "NDUFA6"

PW", "RNF10", "SLIT2", "EDNRA", "CHPT1", "PRRX1", "PKP1", "KLHDC2", "PADI2", "HSDL2", "ROGDI", "T

"SPC25", "IGFBP1", "KHDRBS3", "C1QB", "PHACTR1", "CXCL10", "SATB2", "IFIT2", "HEPH", "IFITM3", "GD

, "FXYD5", "ISLR", "ARHGEF2", "CLIC4", "CDH11", "ENPP2", "OPN3", "GRWD1", "CFH", "ME1", "AKR1B1"
, "PCOLCE", "GRWD1", "LPAR1", "ODC1", "PTGDS", "KPNA4", "HSP90AA1", "SIRT3", "SPC25", "EXOSC9",
, "YRDC", "VRK1", "ACTA2", "MTMR2", "TPX2", "OIP5", "FERMT2", "PRIM2", "TBL1X", "RPF1", "IFITM1",

LCE", "GRWD1", "ME1", "DLAT", "PRIM2", "MACF1", "USP13", "SATB2", "MRC2", "IFIT2", "ZNF83", "COR

IAP25", "RET", "RPS27L", "PRSS23", "MGLL", "SPEN", "IFIT1", "HSD11B1", "SERPINB9", "PODXL", "DNAJC
"LPAR1", "CFH", "COX7A1", "AKR1B1", "ACTA2", "KPNA4", "COTL1", "SPC25", "C1QB", "KCNQ2", "PHAC
"ME1", "ZNF22", "MED27", "GNAS", "DLEU1", "RRP1B", "NELL2", "DIAPH2", "MACF1", "USP13", "NEFM"

PELI1") c("TGFA", "TFF1", "RET", "DTL", "RNF128", "HERC5", "ALDH3A1", "ALB", "IFIT1", "IFI44", "IFIT3", "
", "RGS1", "APOA2", "PPP2R1B", "CXADR", "KHDRBS3", "PPDPF", "GCH1", "USP6NL", "NAMPT", "MUC1"
GRWD1", "LPAR1", "ME1", "MED27", "HSP90AA1", "UBE2C", "NELL2", "MACF1", "USP13", "CALR", "IPO5

I44L", "HSD11B1", "PTGER3", "ADA", "OLFML3", "EIF2S3", "ZDHHC11", "SERPINB9", "JMJD6", "PODXL",

, "FBLN5", "DIRAS3", "GSTO1", "CLIC4", "MFHAS1", "OPN3", "DYRK4", "COL6A2", "TEX10", "CSTA", "CILF
SP90AA1", "SIRT3", "PRIM2", "TBL1X", "LGALS3BP", "CXCL10", "MRC2", "IFIT2", "CORO1C", "IFITM3", "A
LPAR1", "ODC1", "ME1", "MTMR2", "XYLT1", "SPC25", "CXCL10", "IPO5", "MRC2", "IFIT2", "ZNF83", "CO
PEN", "DDX10", "HSD11B1", "LGALS2", "ZDHHC11", "SERPINB9", "PODXL", "SND1", "HSP90B1", "SLC34A2

1", "SPEN", "IFIT1", "HSD11B1", "ZDHHC11", "SERPINB9", "PODXL", "DNAJC10", "GSTO1", "CLIC4", "PCOI
IPOQ", "MAOB") c("TCN1", "ZMAT3", "TGFA", "PLAC8", "C3", "CDH2", "RBBP8", "PRSS23", "AKR7A2", "N

PX", "MST1", "PEG3", "TSKU", "SERPINB2", "MFAP5", "SFRP4", "GCLM", "ENPP2", "PPP2R5E", "TGM2", "I

", "IFITM1", "CXCL10", "SCG5", "IFIT2", "SEMA3B", "PAK1", "TCTN2", "CORO1C", "IFITM3", "ABCA3", "GI
"SFRP4", "DNAJC10", "SPON1", "DIRAS3", "GSTO1", "TRIP13", "PMP22", "CXCL9", "PCOLCE", "GRWD1",
1E1", "ZNF22", "MED27", "MTMR2", "HSP90AA1", "TMPO", "NELL2", "MACF1", "USP13", "PHACTR1", "C

A7", "TMED5") c("IFI27", "CDKN2C", "RNF128", "IFI16", "CYP4F3", "NPY1R", "ECHDC3", "ADA", "SLC9A3F
M1", "ERLIN2", "SKP1", "SH3BGRL", "MXI1", "JAG1", "FAM162A", "ALDH5A1", "BNIP3", "WIF1", "SLC7A8
"CXCL14", "ESR1", "NQO1", "ALDH3A2", "CSTA", "GOLGA7", "CALML3", "SLC7A8", "HOXA5", "S100A8",
", "MLF2", "CLDN7", "NRBF2", "ARHGEF3", "HOXB7", "BBX", "PSMA6", "TSPAN31", "ARL8B", "MGST3", "
", "NELL2", "ZHX2", "SIX2", "SGCE", "TBC1D16", "TUBB2B", "EMCN", "PON3", "RHOBTB1") c("CYB5A", "P
1", "NPEPL1", "SHOX2", "PTEN", "VAT1", "RRM2", "NR3C1", "CXCL10", "SCUBE2", "LTBP1", "SNAP25", "S
"SLC12A2", "PSMD12") c("DLC1", "TOB1", "ZNF423", "GNAQ", "CPVL", "ELANE", "ENTPD4", "COL6A2", "
4A5", "C2CD2", "NTRK2", "HPGD", "SESN1", "PDCD4", "CPB1", "RUNDC3B", "AR", "TMED3", "SSPN", "UB
NTRK2", "PSMD12", "RBBP8", "PGS1", "STUB1", "SLCO3A1", "HOXB6", "NQO1", "DHRS2", "SEMA3F", "C
AP1", "BMPR1A", "RPAP2", "PRTN3", "COL4A5", "NCAM1", "NEBL", "CSTA", "TRIM2", "SESN1", "CX3CL1"
SF2", "ALDH3A2", "FLG", "SLC27A3", "DEGS1", "ZBED2", "NOL3", "TACC2", "CSTA", "CRAT", "BCHE", "PH
", "TBC1D8", "HSD11B1", "PCOLCE2", "SLC27A3", "LPAR1", "GM2A", "MRPL18", "AKAP12", "HERC5", "SII
1", "SLC47A1", "MACF1", "GSTO1", "IFITM2") c("SELENBP1", "ZDHHC11", "TOX3", "GM2A", "HERC5", "SI
", "SLIT2", "FTO", "SAV1", "KCNK2", "WSB1", "ITGBL1") c("USP7", "ETV1", "CTBP2", "GAS6", "DIAPH2", "
PB1", "UBR5", "LAMA5", "CCNG1", "MEIS2", "NPAS3", "ARMCX2", "ARMCX6", "S100A8", "PPP1R3C", "C
DH9", "IBSP", "UNC119B", "FTO", "COL9A3", "PNMA2", "KDSR") c("YME1L1", "MDFIC", "CCNC", "HSPA6"
"GSTA4", "FOXO3", "GPNMB", "NTRK2") c("PRDX2", "TGFBR3", "NOTCH2", "CEBPG", "ANXA9") c("CYP1
c("SGMS1", "ETNK1", "PART1", "GJB1", "TUBB2A", "SLCO3A1", "GADD45B") c("SERPINA5", "TRIM22", "
FOXC1", "B2M", "TBC1D15", "RBPMS") c("ZFP36L2", "PCDH9", "CTSB", "SMC1A", "ITGA5", "IRS2", "DMX
OLR1E") c("LGMN", "SPTBN1", "MAN1A1", "SNHG3", "QKI", "RBBP5", "ZNF124") c("SEH1L", "ESM1", "ITC
18", "NCAM1", "ESR1", "RBBP8", "ESPL1", "SOSTDC1", "NTRK2", "MUC5AC", "PCMT1", "DEGS1", "RTN3"
S2", "PRMT3", "CGA", "SLCO3A1", "EEF1E1", "PLS3", "LSM3", "FKBP5", "INHBA", "PRNP", "ZNF721", "PL

", "BGN", "NXN", "SNAPC1") c("TFPI2", "COLEC11", "FDFT1", "MED1", "CXCL13", "VSNL1", "SCD", "HIRA",

1", "SLC47A1", "CTDSPL", "PIP4K2B", "LGALS1", "CORO1C", "EMP3", "PMP22", "SGCB", "PON3", "ABCA3
"INHBB", "FSTL1", "IGFBP1", "ATP6V1D", "IFIT1", "RET", "CSGALNACT1", "ADAM28", "BASP1", "ISLR", "C
IP1", "LOR", "GALNT2", "SYNM", "PRIM2", "BICD2", "RANBP1", "NUP155", "NME1", "SLBP", "PPID", "PCC
1", "PGR", "NELL2", "PLTP", "LPAR1", "ABCC3", "NOTCH1", "NEFM", "SNAP25", "IFIT1", "SLC22A5", "PKP
", "FBN2", "PIP4K2B", "QKI", "ABCA3", "GNAI2", "PODXL", "CALR", "RAP2A", "MICAL1", "SATB2", "SNRP
EMP3", "LAMC1", "PMP22", "COL6A3", "IGFBP6", "MSH6", "NFIC", "PON3", "THOC2", "CLEC3B", "SDC2",
LIM3", "SYNM", "PTGDS", "BICD2", "TMEM176B", "KLF11", "SLC20A1", "NDUFB6", "PPID", "PCOLCE", "R

c("MGLL", "PELI1", "PRSS23", "FARP1", "PIP4K2B", "CORO1C", "OPN3", "ABCA3", "GNAI2", "ELOVL5", "V
NQ2", "SIRT3", "NOTCH1", "LGALS3", "IGFBP5", "PUF60", "ADI1", "RNH1", "SERPINB9", "NXF1") c("THBS1
AD1", "SPC25", "DNAJC10", "MAOB", "PCOLCE", "RASSF1", "SSX1", "LTBP1", "AHSA1", "SHOX2", "LPAR1

, "PCOLCE", "RASSF1", "LTBP1", "SHOX2", "SGCE", "PGR", "NELL2", "PLTP", "LPAR1", "ABCC3", "ZNF22", "
INB9", "NGDN") c("C5AR1", "SPRED2", "ECT2", "PRKAR2B", "NUP43", "SERPING1", "HLA-DMB", "PLAT",

ACA", "PLOD2", "ZWINT", "IFI27", "HMGA1", "GCLM", "ADORA2B", "EGR2", "G3BP1", "GINS1", "UBIAD1

STA", "SSPN", "LIG1", "VAPB", "HOXB6", "CCNG1", "AHNAK2", "MSX1", "CCDC28A", "GATM", "MFAP2",
", "SERPINH1", "SIRT3", "ZNF22", "LOR", "MRPL18") c("NSL1", "CDH19", "COLEC12", "EYA1", "VAT1", "Z
"PLCB4", "S100A8", "WWC1", "SCARB2", "CSTA", "CELSR2", "ISL1", "PRG2", "ZFP36L1", "PKP4", "SORT1"
UFA4", "GRAMD1C", "GZMA", "FANCG", "ANGPT1", "MGST3", "SLC1A3", "TRIM22", "HOXB7", "GGCT",
PMP22", "PIK3R1", "SERPINE2", "SCUBE2", "LGALS1", "SPON1", "VCAN", "LPAR1", "MGLL", "IFIT1", "IFIT
, "LPAR1", "SPON2", "DMXL2", "COLEC12", "SORBS2", "BICC1", "TIMP3", "DRAM1", "SELENBP1", "SNCA"
BP1", "ABCC3", "PIK3R1", "PCLO", "PYCR1", "ZNF83", "SH3BGRL", "MGLL", "FAM149A", "IFIT1", "IFITM3
2", "CXCL12", "MAP2K5", "RALY", "ANKRD12") c("SLIT2", "LTBP1", "L1CAM", "ABCC3", "PMP22", "PIK3
RPG", "SLPI", "RFC2", "ST7", "PNKP", "RAB31", "CHI3L1", "CD200", "TPBG", "LRRC37A3", "CXADR", "SEC
XCL14", "MUC5AC", "CRCT1", "PKP1", "DYNC1LI2", "MIOS", "WNT5A", "NDN", "NTRK2", "CST6", "CHPT1
NA5", "PSMA6", "ZCCHC10", "PDE6D", "PLS1", "PHACTR4", "FOXC1", "ANGPT1", "COMMD10", "STAM",
LTBP1", "GSTO1", "TCN1", "WIF1", "SLC47A1", "NDUFB3", "RHOBTB1") c("TOX3", "TBL1X", "EYA1", "WN
", "ANGPT1") c("KIF2C", "NDC80", "HMMR", "ITGAE", "OIP5", "PSRC1", "C3", "IGFBP2", "SPRR1B", "SCG
, "SNAI2", "DIDO1") c("NOTCH1", "NCALD", "GDPD3", "LTBP1", "THAP11", "TGFBR3") c("HOXA10", "BIR
MEIS2", "OAS1", "STAT5B", "FBXO21", "CXADR", "ZNF589") c("NPC1", "FOXK2", "NPAT", "HOXB7", "MYC
2", "TUBB2A", "ASXL2", "SYCP2", "MORC4", "ZNF277", "DCK", "CNOT4", "SSBP2", "RAPGEF2", "MNT") c
ORO1C", "PTEN", "LRIG1", "IRF4", "CP", "SNAI2", "PTGDS", "LPAR1", "TFIP11", "SPON1", "MAOB", "TSPAN
LCAM", "RNASE1", "GDPD5", "CHMP6", "PMP22", "PODXL", "PHF10", "TLR7") c("IGFBP4", "A2M", "RGS5
"OPHN1", "RET", "STAP2", "CORO1C", "MTPAP", "OPN3", "SNAI2", "LPAR1", "TFIP11", "CXCL10", "C3AR
AA1", "SPEN", "TGFBR2", "MGLL", "COL15A1", "PXDN", "ABCC3", "PRSS23", "COX8A", "DCTPP1", "TGM
1D1", "ARL4A", "ZBED5", "JAG1", "SNCA", "MBOAT7", "TMEM140", "PSMB4") c("UCHL1", "TCN1", "SERP
, "HLF", "MXD4", "MXI1", "HOXA5", "MT1G", "AR", "BCHE", "TSPAN13", "BBS1", "IFT122", "CRABP2", "I
, "FOXN3", "IGF2BP3", "MDFIC", "SETX", "GTF2I", "ING3", "RIF1", "HOXA5", "SOAT1", "MS4A1", "MAGE
"USP3", "PHF20", "CEBPZ", "RBM15", "SIX2", "UBAP2L") character(0) c("HSPA6", "HIST1H2AC", "AMIGO
1", "LAMA5", "HSPB6", "AKAP7", "PHGDH", "CXCL14", "DDAH2", "ZMYM2", "MEIS1", "MGLL", "FLG", "T
, "PI3", "RPS26", "SPRY1", "GPATCH8", "CTBP2", "LDOC1", "PCDH9", "FEZ1", "COL9A3", "WTAP") c("M
47", "GREM1") c("EML3", "MED13", "TGFBR3", "PRDX2", "KCNK3") c("DTX4", "MAOA", "CD40", "MED13
, "STUB1", "ZNF512B", "TTC3", "CUTA", "OGFRL1", "ELF5", "TMED3", "TSC2") c("TNFSF10", "TSPAN7", "
PAN13", "MPHOSPH8", "PHACTR2", "CHPT1", "TST", "CSTA", "ME1", "NCAM1", "ERMP1", "SEMA4D", "
CYCS", "DET1", "SEMA4D", "SKAP2", "YTHDF1", "ZMYM2", "MCOLN1", "IGF2BP3", "PTPN13", "MAGEA6"

BR3", "CRYBG3", "QKI", "ELF5") c("HOXA10", "COMT", "FKBP14") c("EFEMP1", "TLR2", "CSF1R", "QKI", "
IGF1", "INSIG2", "IRX5", "ISOC1", "ITIH5", "JAG1", "KIAA0355", "MOAP1", "MPHOSPH8", "MSX1", "MXI1
2", "SLC25A24", "WSB1") c("AKAP11", "ANG", "ANXA9", "ARMCX1", "CD302", "CPS1", "CRNKL1", "ELF5"
IFI44", "IFI44L", "ITIH5", "MSX1", "NUP43", "RGS5", "RSBN1", "SPARCL1", "XIST") c("ARMCX2", "ARMCX
TDC1", "ITIH5", "SCARB2", "NR1D2", "LGR4", "TNPO2", "CUEDC1", "RPA2", "DYNC1LI2") c("FBXO21", "E
RX", "PEX3", "TRIM2", "COIL", "HBP1", "TULP4", "ZBED5") c("PIK3C2B", "COMMD3", "MCM6", "RRAGD",
RISPLD2", "CSGALNACT2", "RPS6KA2", "PRELP", "KCNK2", "MED22") c("NRP1", "TNNC2") c("RBP4", "PDE
6A2", "PDE4A", "CRISPLD2") c("PAPSS1", "SEH1L", "AFM") c("TSPAN7", "COL14A1", "GNAQ", "STAG3", "
FPI", "IGFBP5") character(0) c("SNAP25", "UCHL1", "USP7", "SPARCL1", "SGCE", "IL6ST", "SLIT2", "HSPA
"SLC2A10", "SKP1", "ALDH5A1", "PRG2", "MUC5AC", "ABCA5", "NUCB2", "PKP1", "MFAP2", "CKB", "FAM
ERPINA5", "LIF", "PPIF", "F2RL1", "NFATC3", "SLC7A5", "FTH1", "TMPRSS3", "IGFBP1", "EFNA1", "FCHSD
, "OSBPL11", "PIGT", "ROGDI", "EGF", "KLHDC8A") c("TUBB2A", "IRS2", "KCNK3", "FAS", "BNIP3", "RUN

RPS1", "XIST") c("RRAD", "ASRGL1", "STAR", "SNAPC1", "SLC34A2", "TPPP3", "NXN", "CPM", "QPRT", "C

PD5", "LHFPL2", "IFITM2", "LTBP3", "CTNNB1", "MATN2", "GYS1", "HUWE1", "PPID", "RGS5", "ENG", "E

, "ATP6V1D", "ACTA2", "COTL1", "SPC25", "IGFBP1", "DDX60", "IFITM1", "CXCL10", "SATB2", "CBS", "IFI
"AIMP2", "EXOSC8", "PHACTR1", "CXCL10", "MRC2", "FABP7", "SEMA3B", "CCL8", "NME4", "CORO1C",
"CXCL10", "ITPR3", "COL6A1", "CORO1C", "SLC38A1", "IFITM3", "EPHX1", "OPHN1", "IFITM2", "CYC1",

O1C", "IFITM3", "OPHN1", "GDPD5", "IFITM2", "CYC1", "ASCC3", "HUWE1", "PPID", "IVD", "CD81", "EDN

10", "GSTO1", "PCOLCE", "GRWD1", "DLAT", "RAPGEF5", "EREG", "KPNA4", "HSP90AA1", "HMG20B", "S
TR1", "DDX60", "IFITM1", "CALR", "CXCL10", "SATB2", "IFIT2", "HEPH", "TPSB2", "IFITM3", "ABCA3", "L
, "AHI1", "MSMB", "FLVCR2", "IVD", "SLIT2", "EDNRA", "ADIPOQ") c("SNAP25", "HERC5", "ATAD2", "ND

SERPINB2", "GRWD1", "VPS37B", "KIF11", "CHI3L2", "PHACTR1", "DDX60", "IFITM1", "USP6NL", "IFIT2",
, "NR2F6", "ABCA3", "GDPD5", "LMO3", "SCD5", "FXYD6", "PLP2", "NPC1", "HAPLN1", "EDNRA", "DDHD
", "NEFM", "IFIT2", "ZNF83", "CORO1C", "IFITM3", "ABCA3", "GDPD5", "DSTN", "IFITM2", "ASCC3", "HU

"DPYSL2", "USP18", "DNAJC10", "DIRAS3", "GSTO1", "CLIC4", "NFASC", "DHCR7", "PMP22", "PCOLCE", "

", "COX7A1", "ADAMTS1", "ACTA2", "HSP90AA1", "ID3", "A2M", "KCNQ2", "IPO5", "DKK3", "CBS", "HEP
BCA3", "OPHN1", "IFITM2", "PDE4D", "CYC1", "ASCC3", "PLTP", "ACOT9", "MATN2", "HUWE1", "PPID",
RO1C", "IFITM3", "ABCA3", "GDPD5", "DSTN", "IFITM2", "ASCC3", "ANXA5", "HUWE1", "IVD", "EDNRA",
", "TEX10", "DPH2", "COX7A1", "MAN1A1", "HSP90AA1", "SIRT3", "EIF5B", "C1QB", "KCNQ2", "NBPF10"

LCE", "GRWD1", "LPAR1", "KPNA4", "HSP90AA1", "SIRT3", "SPC25", "USP13", "LGALS3BP", "CALR", "CXC
MUC16", "PPP1R15A", "FANCL", "ZNF337", "RUNX1", "LOR", "RUNX3", "SLC6A14", "IFIT1", "CXCL12", "G

KLK6", "TRHDE", "VPS37B", "LPAR1", "ELMO1", "NBL1", "SDC2", "RECK", "RGS1", "CKS2", "KIF11", "SPOC

, "LPAR1", "PTGDS", "ZNF22", "MTMR2", "SIRT3", "RRM1", "PRIM2", "NELL2", "MACF1", "USP13", "IFITM
ALR", "NEFM", "UNG", "IFIT2", "SEMA3B", "ZNF83", "CORO1C", "IFITM3", "IFITM2", "PDE4D", "ASCC3", "

R1", "FGF13", "DPYSL2", "SLPI", "TMPRSS3", "EGR2", "GCLM", "CREM", "TLR4", "IMPA1", "ELK3", "KPNA4
3", "CDKN1B", "BCL11A", "MEIS2", "CLU", "CITED2", "NUDT11", "MT1G", "TTC17", "CD36", "NEBL", "SLC2
"SCNN1A", "JAG1", "MAX", "PEX5", "CCDC6", "ANG", "SHMT1", "CRNN", "XIST", "DACH1", "GATM", "CP
FOXC1", "MCTS1", "SLC1A3", "GAD1", "ERAP1", "CKS2", "COX7A2", "NCF4", "LAP3") c("ZFP36L2", "SMC1
PP2R1B", "SLC37A4", "MTHFD2", "SULF1", "CDS2", "FABP7", "IFIT1", "HERC5", "GSTM3", "EPS8L3", "PD(
PON1", "NOVA1", "TIMP1", "SQLE", "RNASE1", "IFI6", "NELL2", "ZHX2", "SIX2", "SGCE", "TBC1D16", "PB
"OSBPL9") character(0) c("ZNF423", "CASP10", "SORBS3", "GNAQ", "IRS2", "ELANE", "ENTPD4", "COL6A2
L3", "FOSL2", "WIF1", "MIOS", "WNT5A") c("RRAGD", "TNFSF10", "E2F2", "C3", "GALNT2", "BCL11B", "F
YB561", "WIF1", "SCNN1A", "GATM", "NDN", "ACPP", "PMM1", "TCEAL2") c("SCAND1", "RGS4", "CBX4"
, "CRYAB", "OXA1L") c("SCGB2A1", "RGS4", "RASGRP1", "ACSL1", "SLC25A11", "FBXL5", "TNS3", "GCLC",
YH", "NCAM1", "SEMA4D", "GOLM1", "KLHDC2", "DHRS2", "AHNAK", "IGF1", "ALDH3B1", "SNAPC1", "ZN
DT2", "ZSCAN18", "DIRAS3", "FADS1", "TRIM44", "MYOT", "LTBP3", "FAM169A", "ACTA2", "GSTO1", "GS
IDT2", "MYOT", "SULF1") c("DSG3", "FOXM1", "SMC1A", "ARF1", "SGCE", "SFRP4", "MSMB", "NBAS", "TH
RBPJ", "RAB27A", "HGSNAT") c("CD40", "CTBP2", "KLF6", "RPS26", "RBPJ", "FTO") c("ABCG1", "SOAT1", "
RISP3", "NTRK2", "TSPAN13", "AZGP1", "SSPN", "ZNF329", "HBS1L", "CLU", "GULP1", "HOXA5", "ZNF211
", "CYTIP", "HIST1H2AC", "SKAP2", "AMIGO2", "MAGEA6", "IGF2BP3", "HOXA5", "MS4A1", "ZMAT3", "M
B1", "MAOA", "PFKFB3", "CFI") c("PLTP", "C1R", "IL1R1", "HSPA6", "HLA-DPB1", "MTHFD2", "SLC16A4",
RCBTB2", "GYG1", "SLC1A3", "ACTR2") c("PRF1", "GNAI1", "SMC1A", "ADD3", "MAN1A1") c("BPHL", "ST(
L2") c("PCDH9", "GRINA", "SPRY1", "EIF5", "PTPN13") c("GPA33", "GRINA", "ZNF281", "PDE4D", "EIF5", "
GB3") c("ESF1", "MYRIP", "BMP6", "MAN1A1", "ATP8B1", "IRS2", "PCOLCE2", "RBP4", "ZNF124") c("MSX
, "NASP", "NFIC", "CUEDC1", "IRX5", "WIF1", "KRT1", "FKBP1B", "AZGP1", "GLRX", "GABBR1", "SCNN1A"
XNA1", "ABHD6", "AMACR", "HIGD1A", "DCK", "RNF138") c("CKS2", "ACOT7", "RCBTB2", "CDK2AP1", "T

, "NEK2", "VLDLR", "MKI67", "H2AFX", "NEFM", "RAD51C", "HLA-A") c("COLEC11", "TSPAN5", "FBN2", "D

", "VAT1", "CCDC88A", "MICAL1", "LRIG1", "SERPINE2", "SFRP4", "UCHL1", "TUBB2B", "LOR", "SPARCL1"
CXCL10", "FCGR2A") c("MGAM", "ADM", "SNX10", "SERPINB2", "DNAJB4", "INSIG1", "DNAJB1", "KCTD5"
OLCE", "OPHN1", "ENPP2", "SCML1", "LTBP1", "CD34", "COL6A2", "VRK1", "PLTP", "PDE4A", "LPAR1", "A
1", "SERPINB9", "CP") c("MGLL", "HSPA6", "RRM2", "PRSS23", "HSP90AA1", "UBE2C", "CKS1B", "FARP1"
A", "CLIC4", "SPARC", "ZDHHC11", "SNHG3", "SPC25", "SCD", "POLR2I", "GALNT2", "SORBS1", "PTGDS", "
"SERPINH1", "UCHL1", "NQO2", "TAGLN", "TUBB2B", "CCL19", "SPARC", "VIM", "FKBP11", "TKTL1", "RD

VAT1", "MICAL1", "PCOLCE2", "CLIC4", "ZDHHC11", "SNHG3", "UBIAD1", "SPC25", "DNAJC10", "GALNT2"
", "ANPEP", "CA12", "COL6A3", "PON3", "CXCL9", "CLEC3B", "PCOLCE2", "UCHL1", "CGA", "GALNT2", "S
", "ABCC3", "LGALS3BP", "SIRT3", "NOTCH1", "SNAP25", "IFIT1", "RET", "SERPINB9", "GBP1", "CXCL10")

"TAGLN", "TESC", "RCAN2", "DUSP7", "RFTN1", "H2AFX", "RGS2", "ZNF281", "TCTN1", "SEMA6A", "NRN

", "PRDX4", "P2RY13", "GLIPR1", "SLPI", "APOBEC3B", "FANCI", "IMPA1", "CDKN2C", "ECHDC3", "KNTC1

"AHNAK", "CALML5", "CYB561", "TPD52", "GAMT", "PEBP1", "CXCL14", "ENO2", "PRG2", "RBBP8", "CKB
MAT3", "CCNE2") c("FOXM1", "KIF15", "ARHGAP11A", "GGH", "SMC1A", "SFRP4", "THBS2", "MACF1", "T
", "SYNJ2BP", "GATM", "AZGP1", "DDAH1", "SH3BGRL", "DEGS1", "AHR", "MGLL", "AHNAK", "MTA1", "TI
"RCBTB2", "LAP3", "PTPRO", "LDB3") c("ZFPM2", "CLU", "SEMA6A", "PLCB4", "MDM4", "KIAA1324", "TA
TM3", "DMXL2", "IFITM2", "RUNDC3B", "IVD", "CLNS1A", "VPS13C", "MXRA5", "IKBKE", "EDNRA", "IFITM
", "PFKM", "LAMC1", "KCTD12", "NID1", "MCAM", "DYNLT3", "MXRA5", "SPARC", "PLN", "PXDN", "RELN
3", "RUNDC3B", "IVD", "ENO2", "FABP7", "CLNS1A", "DYNLT3", "CXCL10", "EDNRA", "FTO", "SNAP25", "C
R1", "NR3C1", "DAAM1", "SCUBE2", "IFI6", "SPON1", "ZNF83", "VCAN", "LPAR1", "RRM2", "QKI", "IFITM
14L1", "MUC1", "PTMS", "RGS5", "HAPLN1", "GPM6A", "HIST1H2BG", "RAB5B") c("NDUFA4", "TLR4", "G
L", "RTN3", "ZMYM2", "ELAVL1", "NQO1", "BCHE", "FGFR2", "NISCH", "LMO4", "CSTA", "PEBP1", "TIMM
"ERBB3", "EHD3", "PECR", "ALDH2") c("EMP3", "CACYBP", "FTO", "ITGA5", "KIAA1324", "ERCC1", "MRP
NT5A", "SLC35A1", "IFITM3", "GM2A", "OXA1L", "OPHN1", "EIF2AK2", "PELI1", "NUDT11", "CDH19") c("K
B2A1", "TSPAN7", "TNFSF10", "HMGB2", "C4BPA", "ACSL1", "MCM2", "FAM117A", "MCM6", "IKZF1", "C
C3", "ZFP36L2") c("HMOX1", "AKAP12", "PIAS3") c("BIRC3", "LCP2") c("SFRP1", "SGCE", "ATP8B1", "LTBP
N", "LPCAT4", "SERPINA5", "F2RL1", "FCHSD2", "TRIAP1", "SAP30") c("CXCL14", "BEX4", "ARMCX6", "NT
c("TSPAN14", "CKS2", "ARHGEF3", "NPAT", "HOXB7", "PCGF1", "NUP153", "LPCAT4", "TCTA", "SLC1A3", "
N7", "L1CAM", "ZNF337", "UBE4A", "OGN", "SCUBE2", "GDPD5", "PMP22", "SNAP25", "PHF10", "ZNF58
5", "ADAMTS1", "HHEX", "NPY1R", "UMOD", "SIX2", "COL6A2", "EMCN", "CPT1A", "AKAP12", "GNG11", "
1", "L1CAM", "UBE4A", "SYCP2", "SCUBE2", "GDPD5", "PPOX", "SNAP25", "PHF10") c("SPEN", "MGLL", "
2", "CORO1C", "LRIG1", "TOB2", "IRF4", "LPAR1", "TFIP11", "TSC22D1", "MAOB", "LRRN3", "L1CAM", "F1
PINB2", "SPEN", "TFF1", "SLC47A1", "SIX2", "ABCC3", "ZNF706", "PRSS23", "STAP2", "CORO1C", "RAD51C
TIH5", "INSIG2", "PPP1R3C", "AOX1", "IRX5", "CXCL14", "BEX4", "RNF128", "AZGP1", "MUC5AC", "BNIP3
A6", "NOTCH2", "ACSL4", "IL11", "MFAP2") c("PTPN13", "PCSK5", "BLMH", "DSC1", "GSTA1", "CNOT4", "
2", "CUL2", "ZNF281", "CNOT7", "FBXO28", "YTHDF1") c("PSMD12", "ARMCX1", "FAM102A", "UBAP2L")
M7SF2", "EXPH5", "TACC2", "LOR", "DACH1", "CST6", "TCFL5", "AHNAK2", "PER3", "CD36", "LGR4", "GU
AGEA6", "ZMAT3", "PTGER4", "NOTCH2", "IL6R", "YTHDF1", "HIST1H2AC", "IL7R", "GPA33", "ZMYM2", "
", "TEX2") c("TCF4", "HMOX1", "CD81") c("TCL1A", "PRMT2") c("GNA11", "ACAT1", "CHP2", "TNS1", "AK
REEP1", "C3", "LONP2", "ENPEP", "TCTA", "ACSL1", "RAB17", "NVL", "SPRR1B", "MRPL48", "IKZF1", "GPX
SOCS2", "DACH1", "CALML5", "MGLL", "ZMYM2", "KLK11", "PPDPF", "CST6", "HPGD", "MKNK2", "DUSP2
", "PASK", "MFAP2") c("IRX5", "NME4", "TBC1D4", "CHPT1", "RPS6KA2", "PPARG", "COBL", "NUAK1", "C

CFD", "RBM5") NRP1 c("TGFBR3", "SFRP1", "SNAP25", "FHL1", "CCND2", "ATP8B1", "FKBP14", "ZC3HAV
.", "NCAM1", "NTRK2", "OXA1L", "OXR1", "PDCD4", "PDK3", "PHYH", "PMM1", "PPP1R3C", "PRR15L", "R
", "ENO2", "FBXO21", "FOXF1", "GINS1", "IRX5", "MXI1", "NME4", "OXR1", "PCSK5", "TCEAL2", "TMEM1
3", "BASP1", "BST2", "C1S", "CAMK2N1", "CFH", "CLNS1A", "COG5", "COL4A6", "CPA3", "CTNNB1", "CTS
2F2", "ITGAE", "ALDH3A1", "NMT1", "FCGBP", "PSAT1", "APBA2", "P4HB", "HMGB2", "SLC16A6", "KIF2C
, "FDPS", "DHCR7", "TNFSF10", "TRPM4", "TPRKB", "ENPEP", "RTN3", "SREBF1", "ARID1A", "KIF2C", "CHK
E4DIP", "DCN", "DEGS1", "FHL1", "CSGALNACT2", "ELF5", "ANK2", "NFIB", "RPS6KA2", "PRELP", "SLC38A
RBP4", "JAM3", "FGFR1", "COL6A2", "GULP1", "MCOLN3", "SFRP1", "FHL1") c("MCOLN1", "SERPINA3", "
5", "BMP6") c("BCAT1", "LGALS1", "C3", "UCHL1", "CTSC", "CD93", "LPL", "NNMT", "THY1", "PDIA4", "C1
M162A", "BCOR", "ARMCX6", "CXCL14", "SCNN1A", "LRBA", "ROGDI", "TRHDE", "GULP1", "HOXA10", "ES
2", "LGALS3", "NPC1", "NPDC1", "ROGDI", "HBD") c("LIG1", "NCAM1", "BCHE", "KLK11", "AZGP1", "CELSR
K1", "ABHD6", "PLS3", "TPRKB", "CGA", "SIPA1L1", "PRNP", "FKBP5", "ZNF274", "DCK", "GADD45A", "AD

A4", "TPSB2", "NR2F6", "LYZ", "FBXL12", "PPID", "RGN", "BGN", "RAMP1", "RCAN2", "STAP1", "ASTN2") c

EDNRA", "GLI2", "CLIP2", "IGLL1", "TOX3", "LAPTM4B", "SYCP2", "PELI1") c("SNAP25", "ZMAT3", "PRSS23

T2", "HEPH", "IFITM3", "LHFPL2", "IFITM2", "CTNNB1", "CTSH", "FN1", "ANXA5", "GYS1", "HSPA5", "PPID
"IFITM3", "ABCA3", "PUF60", "GDPD5", "ANXA5", "ERAP2", "MATN2", "HSPA5", "TBC1D1",

NRA", "HGF", "MAOB", "COL4A6", "NOTCH1") c("ME2", "RET", "MICB", "RPS27L", "PRSS23", "COBL", "SP

IRT3", "SPC25", "PRIM2", "CXADR", "MACF1", "USP13", "IL24", "LGALS3BP", "CALR", "CXCL10", "SATB2"

UFA4L2", "BET1", "ITGAE", "TOX", "CD69", "HLTF", "PARP8", "PHACTR1", "WNT5A", "ABCA3", "OPHN1",

, "MUC1", "IFITM3", "IFITM2", "METTL7A", "PXDN", "ACOT9", "HAPLN1", "GYS1", "RNF10", "B2M", "SF3A
D2", "GLI2", "HGF", "TOX3", "TNFAIP8", "LONP1", "PDSS2", "NOTCH1", "KYNU") c("SNAP25", "ZMAT3", "F
WE1", "ANXA2", "IVD", "CD81", "EDNRA", "MAOB", "COL4A6", "PDSS2", "NOTCH1", "PELI1") c("SACS", "I

GRWD1", "LAP3", "LPAR1", "COX7A1", "SLC20A1", "PTGDS", "MAN1A1", "ACTA2", "HSP90AA1", "SMC2"

"MAOB", "COL4A6", "SSX1", "ELOVL5") c("RET", "AKAP12", "RPS27L", "DIP2C", "FANCL", "PPIF", "GNG1
", "TPSB2", "RNH1", "OPHN1", "PUF60", "ZBTB10", "LGALS3", "TESC", "U2AF1", "PLTP", "YWHAE", "UST"

L10", "IPO5", "SATB2", "MRC2", "IFIT2", "ZNF83", "CORO1C", "IFITM3", "ABCA3", "GDPD5", "GPI", "DSTI
AMT", "ITM2A", "CGA", "GABRP", "SPP1", "SCML1", "KRT7", "MUC5B", "LGALS1", "PCOLCE", "VPS37B", "

K1", "TIMP3", "CXADR", "C1QB", "SLCO2A1", "TCF21", "MAPKAPK5", "NAMPT", "QPRT", "MFGE8", "PCP

M1", "CALR", "CXCL10", "QPRT", "SCG5", "NEFM", "MRC2", "SEMA3B", "ZNF83", "PAK1", "CORO1C", "IFIT
"HUWE1", "IVD", "CD81", "SLIT2", "EDNRA", "MAOB", "TIMM9", "NOTCH1", "THBS2") c("CXCL6", "PLAT

4", "GINS1", "DEPDC1", "LBR", "TOX", "PRDX4", "APOBEC3B", "KNTC1", "PHACTR1", "GCH1", "CCDC59",
24A3", "CRYAB", "ZMYM2", "FGGY", "FBXL5", "BCHE", "NDN", "PRTN3", "GIN1", "C2CD2", "ELF5") c("HM
B1", "NUDT11", "SLC2A10", "COG5", "SORT1", "AZGP1", "INSIG2", "NTRK2", "TST", "PDK3", "NEBL", "TM
1A", "BCL10", "ALDH1A2", "WSB1", "PDE8B", "CLU", "SIX2", "STAC", "GAS6", "GNAI1") c("TMEM134", "G
GFRL", "FADS1", "ZSCAN18", "OGT", "ABCA3", "AKAP12", "AHCY", "EDNRA", "SCUBE2", "SNAP25", "DHC
K", "PON3", "RHOBTB1") c("TRAPPC4", "MTHFD2", "SULF1", "MAX", "NUDT11", "HERC5", "ADAMTS5", "
2", "STAG3", "MYRIP", "NTRK2", "OSBPL9") c("TLR4", "CLIC4", "CSGALNACT1", "ICAM2", "HYOU1", "CYP2
CGBP", "CHGA", "OLFML2A", "HNRNPUL1", "SLC37A4", "RASGRP1", "TRIM22", "ADI1", "RAB25", "IGFBP
, "TNFSF10", "TXNL4B", "E2F2", "DHCR24", "SPRR1B", "DHCR7", "ACSL1", "SPRR2B", "ZNF512B", "COX7E
, "BCKDK", "FCGBP", "USP46", "SPATA20", "P4HB", "HMMR", "IL13RA1", "TCTA", "C4BPA") c("INHBA", "I
NF211", "NBAS", "ZNF512B", "MUC5AC", "UBL3", "DHTKD1", "MSX1", "CLIC3", "DDAH1") c("EGF", "AZGF
TM3", "IFITM2", "SULF1") c("XIST", "RGS5", "NQO1", "BTK", "RBPJ", "CDH2", "SYNE1", "TRIM44", "MYO
HBS2", "HIST1H2BK", "PCSK6", "MACF1", "EML4", "TSPAN4", "LRRC2") c("MAOB", "NTRK2", "ZNF365", "
"PDGFRA", "KCNK2", "SMURF2", "HGSNAT") c("NRP1", "SLC39A6", "ESM1") c("USP7", "DEGS1", "GAS6",
.", "XIST", "CYB561", "CAST", "BCHE", "JAG1", "MFAP2", "DHRS2", "SLC24A3", "MSX1", "AHNAK2", "CXCL
ICOLN1", "LGALS3BP", "MFAP2", "PTGER4", "YTHDF1") c("CKB", "RAB4A", "PPARG", "F2R", "UCHL1", "ZS
"HMOX1", "CDC42SE1", "C1S", "PPFIBP1") CP c("PLTP", "GPM6B", "IL1R1", "ARMCX1", "HSPA6", "MTHF
C2", "RNF144A") c("AMIGO2", "IGF2BP3", "ZMYM2", "BPHL") c("TF", "SFRP1", "ABCA8", "FOSB", "TCEAL
"PTPN13") c("EIF3E", "FAM102A", "TF", "P2RX4", "CHPT1", "PLEKHB1", "PAQR6", "PLAGL1", "PLCD1", "P
1", "SSR3", "MAP1A", "CSGALNACT1", "HEPH", "FERMT2", "SOX17", "NES", "ESM1", "ITGB3") GTF3A c("
, "TSPAN13", "CX3CL1", "BNIP3", "NUDT11", "CXCL14", "SQSTM1", "SLC7A8", "KLK11", "IGF1", "ALDH3A
IMELESS", "HLA-DMB", "EXO1", "SLC1A3", "DDX10", "ANGPT1", "PFDN2", "PSRC1", "SERPINA5", "CEACA

DHCR7", "PLA1A", "LPL", "SLC1A3", "SLC30A1", "NEFM", "IFIT1", "SEMA3C", "GAS1") c("COLEC11", "TSPA

, "DNAJC10", "PTGDS", "MAOB", "PCOLCE", "RASSF1", "LTBP1", "SHOX2", "SGCE", "PGR", "NELL2", "PLT
, "CA12", "SLC7A7", "ZWINT", "MUC1", "CD2", "PLAT", "SAMSN1", "NCAPD2", "G3BP1", "MICAL2", "PCS

, "FJX1", "NUSAP1", "FADS1", "SMC2", "SLC39A14", "CORO1C", "LRP4", "MCM2", "LAPTM4B", "TPX2", "
"PRIM2", "ME2", "PRIM1", "STEAP3", "PPID", "PCOLCE", "RASSF1", "BICC1", "SSX1", "LTBP1", "SLC37A4"
H11", "DNAJC10", "UBE2L3", "KLF11", "KIF5C", "NR3C1", "CAV1", "RASSF1", "BICC1", "TBC1D1", "LTBP1

, "SORBS1", "PRIM2", "FANCA", "PPID", "PCOLCE", "RASSF1", "BICC1", "SSX1", "OPHN1", "LTBP1", "HER
ORBS1", "PRIM2", "MT1H", "SHQ1", "CRYAB", "LUM", "SACS", "CRIP1", "PKD1P1", "DHRS2", "RAD51C",
c("FGFR3", "PLAT", "BIRC5", "CPS1", "NARS2", "TMPO", "LAGE3", "KCNQ2", "MDFIC", "NOTCH1", "LGAL

1", "MYH11", "G0S2", "CANX", "SLC46A3", "RGS5", "ALDH7A1", "LTBP2", "CD14", "VRK2") c("GRIA2", "E

.", "TLR4", "SEL1L", "SELL", "SLC1A5", "FZD6", "MLLT3", "HOXB2", "BCAT1") c("ZNF318", "CXCL12", "GNG

", "TACC2", "LOR", "GULP1") c("KIF2C", "HMMR", "OIP5", "C4BPA", "NDC80", "MCM6", "HMGB2", "GMI
TP53", "EXOSC4", "ZNF587") c("GRWD1", "TMPO", "UCHL1", "TIMP1", "NDFIP1", "USP13", "NTRK2", "FT
RPS1", "RAB40B", "IRX5", "ZNF512B", "CD36", "SH3GLB2", "CUEDC1", "ARMCX6", "RUNDC3B", "PADI2",
RP", "USP3", "DMXL2", "FEZ1", "PRSS21", "IRS2", "PTER", "ADD3", "FTO", "WSB1", "HLTF", "RBPJ", "EMI
I1", "FTO", "MACF1", "SNAP25", "PCCA", "HSP90AA1", "TIMP1", "KCNMA1", "GSTO1", "HSD11B1", "PTP
", "PCP4", "MAN1C1", "SDC2", "PRKCA", "THOC2", "DNAJC15", "LRRN3", "LPP", "TIMP1", "XIST", "GOT1"
KB", "PCCA", "AHCY", "KCNMA1", "GSTO1", "HSD11B1", "EMP3", "MAP2K5", "OPHN1", "PINK1") c("LAP
3", "DMXL2", "IFITM2", "IVD", "NPEPL1", "QPRT", "MRPL44", "TCN1", "CXCL9", "MRC2", "MXRA5", "IKBK
3BP1", "SLPI", "APOBEC3B", "DPYSL2", "SELL", "TBC1D12", "ATXN7", "COLEC12", "VCAM1", "HOXB2", "
22", "CUTA", "OXA1L", "STUB1", "PHGDH", "DACH1", "MT1G", "ISL1", "PRG2", "OGFRL1", "HLF", "JAG1",
S30", "MSX1", "CLU", "KLF2", "PCDH9", "IRS2", "SF1", "EIF1", "GNAI1", "SMC1A", "ZFP36L2", "PTX3") c("
CNN4", "SGCE", "TSPAN4", "ATOX1", "PBX1", "PLA1A", "MSMB", "DDA1", "DSG3", "EXOSC4", "G6PD", "
OSBPL11", "SLC25A12", "DHCR7") c("XIST", "PELI1", "PRMT3", "MTF2", "ERGIC2", "PLS3", "HSPA1A", "RA
1", "TGFBR3", "MRC2", "SLC38A2") c("SFRP1", "KLF10", "MSX1", "PYCARD", "SNAI2", "YTHDF1", "PCOLC
RK2", "XIST", "OXA1L", "MGLL", "THUMPD1", "ESR1", "PSMD12", "SLC2A10", "HPGD", "LGR4", "TSPAN1
"FAF2", "PEX6", "GZMA", "SERPINA5", "NCF2", "SAP30", "ARNT2") c("KLF2", "CREBBP", "ZFP36L2", "PSM
", "NELL2") c("RGS5", "CDS2", "ABAT", "MGLL", "AHCY", "AKAP12", "ABCC3", "FGF13", "PRSS23", "COX
FGF13", "EPS8", "LYZ", "DIP2C", "ITGB1", "RET", "STAP2", "CSNK1E", "CD37", "TAF1D", "INHBB", "ADAM
RBMS1", "COL15A1", "EMCN", "ABCC3", "PRSS23", "TLE1", "STAP2", "DCTPP1", "CORO1C", "TAF5", "SN
L3A1", "UBE4A", "VEGFA", "UGDH", "GDPD5", "MRPS16", "PMP22", "PPOX", "SNAP25", "PHF10", "ZNF5
", "PTEN", "LRIG1", "HAUS4", "REV3L", "NGDN", "SAR1B", "SNAI2", "PTGDS", "LPAR1", "TFIP11", "CXCL1

PRKCZ", "MBIP", "ANXA9", "SLIT2", "UCHL1", "COPS7A", "SLC7A8", "GNB5", "ENO2", "RIF1", "NME4", "C
) c("TLK2", "HNRNPA1", "S100A9", "CHMP7") c("HSPA6", "FOXO3", "PHF20", "SFRP4", "ADNP", "H2AFV"

EPB41L4B", "BICC1", "PDE4D") c("GSTA1", "RPS6KA2", "ZSCAN18", "PLEKHB1", "F2R", "CEBPG", "NOTCH
T3", "TERF2", "UCHL1", "CCDC47", "KIF5C", "TGFBR3", "RETSAT", "DEGS1") c("MCOLN1", "SERPINF1", "
7", "OLFML2A", "ZNF512B", "SKP2", "HDHD3", "C4BPA") c("AMACR", "RAB9A", "ABHD6", "TUBB2A", "P
2", "C2CD2", "CRABP2", "GULP1", "SH3BGRL", "TCFL5", "JAG1", "BCHE", "PTOV1", "OXA1L", "MFAP2", "
TBP2", "PTPN13", "TMEM204", "SFRP1", "UCHL1", "ANG", "PDLIM5", "PLEKHB1") c("SERPINA1", "VAMP

", "TSPAN7", "OLFML2A", "EGF", "OIP5", "DHCR7", "C4BPA", "PSRC1", "HMMR", "MCM2
A", "C4BPA") c("COMMD3", "HNRNPH1", "FOXO1", "WDR61", "ERGIC2", "ABHD6", "S

R1", "ACPP", "TPD52", "WNT5A", "ISOC1", "ESPL1", "LGR4", "XIST", "IRX5", "ELAVL1", "N

IGB2", "ID2", "PSAT1", "FCGBP", "NDC80", "ABCB6", "BRD8", "HMMR", "FAM111A",

PATCH8", "ISG20L2", "ADNP2", "WSB1", "KDSR", "RNF144A") c("GINS1", "HOXA5"

'SELENBP1", "OGT", "VAT1", "CCNE2", "SNAP25", "WNT5A", "TKTL1", "COLEC12"

3", "CDC7", "NDUFA6", "NDUFB6", "RAMP3", "GPN3", "OSBPL11", "C4BPA", "CASP1

P1", "RGS4", "SRPK2", "NDRG2", "RASGRP1", "NVL", "TRPM4", "ITGAE", "IGFBP2
T", "F2R", "DENND5A", "MSX1", "CCL18", "GSTO1", "ARHGAP26", "SULF1") c("CL

SCAN18", "ENPP1", "PCOLCE2", "NME4", "FOSB", "RPRD2", "ZNF551", "MAGEF1
FD2", "PCOLCE2", "ANK2", "TGFBR3", "GSTA4", "FOXO3", "GPNMB", "ATP8B1", "NT

AM6", "SAP30", "HLA-DQB1", "NCF2", "HLA-A", "UBE3B") c("SMC1A", "PRSS23",

NN", "PSRC1", "MCM2", "E2F2", "TMEM14A", "GPN3", "PSAT1", "ALDH3A1", "TNFSF10

P3", "SMC1A", "MAN1A1", "IL7R") c("PNMA2", "SPRY1", "LDOC1", "FEZ1", "PRSS21

3", "CRYAB", "S100A8", "PHYH", "AZGP1", "SQSTM1", "FGFR2", "SLC24A3", "RXRA"
ID12", "MRPS30", "PTX3", "SIX2", "KIAA1324", "MED17", "PRSS23", "IRS2", "AFF1",

ITS5", "CRY1", "TEX10", "TOX3", "ARMCX3", "RPP38", "MED23", "WEE1", "MED

TMEM158", "ADAMDEC1", "AKT3", "UCHL1", "PRMT2", "MCAM", "SERPINA3") c(
